## Supplementary material for "Completing the BASEL phage collection to unlock hidden diversity for systematic exploration of phage-host interactions": Data S2: 64.html

FANPEZAQ\_CDS\_0064


Return to summary | Go to previous | Go to next

|  |  |
| --- | --- |
| FANPEZAQ\_CDS\_0064  Page creation date: 02 Sep 2024, 12:00  Project folder: n/a  Input sequences file: Escherichia\_virus\_HeidiAbel.gb | lysozyme endolysin t4 receptor hydrolase e c glycoside fragment chimera phage type muramidase phage\_related glucagon\_like peptide a membrane lysis b factor in the baseplate complex calcium domain\_containing gh24 yes mitochondrial alpha putative escherichia coli engineered and pesticin so4 fusion phosphatidylinositol 4\_kinase adrenergic enterobacteria organism\_taxid gene expressed expression\_system\_taxid mainly arch orthogonal |

### Sequence information

|  |  |
| --- | --- |
| Name | FANPEZAQ\_CDS\_0064  64\_FANPEZAQ\_CDS\_0064 (pipeline id) |
| Imported annotations | Escherichia\_virus\_HeidiAbel Bas97 |
| Protein sequence | MKISSNGIAVLKYFEDCHLKAYPDPATGGAPWTIGWGHTGPEVKRGLVWTQKQADDALVS DLARFESAVSAAVRVPLNQGQFDALVSFTYNLGEGNLKSSTLLKMVNAGNFAGAAEQFKR WNKANGKTMRGLTRRRAAEQCLFLGMGGASAIERGVAAA |
| Number of residues | 159 |
| Molecular weight (Da) | 17019.22 |
| Output files | ../../query\_sequences/64\_FANPEZAQ\_CDS\_0064.fasta |

### Putative domain architecture and protein family

#### Search results (HHblits)^1^

|  |  |
| --- | --- |
| Domain family databases searched | Pfam, Ncbi-cd, Cath, Phrogs |
| Results, scheme(s)  (Top layers only; threshold 1.00e-03 (evalue)) | xml version="1.0" encoding="utf-8" standalone="no"?       2024-09-02T21:08:26.346943 image/svg+xml   Matplotlib v3.7.2, https://matplotlib.org/ |
| Results, table  (E-value ≤ 1.00e-03 (evalue)) | db \| id \| prob \| evalue \| pvalue \| score \| cols \| query \| query\_len \| template \| template\_len \| name \| description; pfam \| PF00959 \| 98.5 \| 9.9e-12 \| 1.7e-15 \| 70.8 \| 106 \| (32, 137) \| 159 \| (2, 107) \| 108 \| Phage\_lysozyme \| Phage lysozyme; pfam \| PF16754 \| 97.3 \| 1.2e-07 \| 2.2e-11 \| 55.9 \| 104 \| (9, 112) \| 159 \| (5, 149) \| 149 \| Pesticin \| Bacterial toxin homologue of phage lysozyme, C-term; ncbi-cd \| cd16900 \| 99.2 \| 9.1e-16 \| 1.6e-19 \| 88.5 \| 137 \| (4, 144) \| 159 \| (4, 142) \| 142 \| endolysin\_R21-like \| cd16900 endolysin\_R21-like; endolysin R21-like proteins. Unlike T4 E phage lysozyme, the endolysin R21 from Enterobacteria phage P21 has an N-terminal SAR (signal-arrest-release) domain that anchors the endolysin to the membrane in an inactive form, which act to prevent premature lysis of the infected bacterium.; ncbi-cd \| cd16901 \| 99.2 \| 2.1e-15 \| 3.5e-19 \| 86.1 \| 138 \| (3, 144) \| 159 \| (2, 140) \| 140 \| lyz\_P1 \| cd16901 lyz\_P1; P1 lysozyme Lyz-like proteins. Enterobacteria phage P1 lysozyme Lyz is secreted to the Escherichia coli periplasm where it is membrane bound and inactive.; ncbi-cd \| cd00737 \| 98.6 \| 8.1e-12 \| 1.4e-15 \| 69.4 \| 131 \| (9, 143) \| 159 \| (3, 135) \| 136 \| lyz\_endolysin\_autolysin \| cd00737 lyz\_endolysin\_autolysin; endolysin and autolysin. The dsDNA phages of eubacteria use endolysins or muralytic enzymes in conjunction with hollin, a small membrane protein, to degrade the peptidoglycan found in bacterial cell walls.; ncbi-cd \| cd00735 \| 98.2 \| 1.6e-10 \| 2.8e-14 \| 65.2 \| 128 \| (8, 144) \| 159 \| (3, 140) \| 146 \| T4-like\_lys \| cd00735 T4-like\_lys; bacteriophage T4-like lysozymes. Bacteriophage T4-like lysozymes hydrolyze the beta-1,4-glycosidic bond between N-acetylmuramic acid (MurNAc) and N-acetylglucosamine (GlcNAc) in peptidoglycan heteropolymers of prokaryotic cell walls.; ncbi-cd \| cd16904 \| 96.8 \| 1.8e-06 \| 3e-10 \| 47.3 \| 86 \| (4, 93) \| 159 \| (2, 138) \| 138 \| pesticin\_lyz-like \| cd16904 pesticin\_lyz-like; pesticin C-terminal-like domain of uncharacterized proteins. This subfamily is composed of uncharacterized proteins containing a lysozyme-like domain similar to the C-terminal domain of pesticin.; cath \| 3hdeC00 \| 99.4 \| 3e-17 \| 3.7e-21 \| 98.5 \| 134 \| (10, 146) \| 159 \| (30, 163) \| 165 \| Lysozyme \| CATHCODE: 1.10.530.40 NAME: Lysozyme. Chain: a, b, c, d. Synonym: lysis protein, muramidase, endolysin. Engineered: yes SOURCE: Enterobacteria phage p21. Bacteriophage 21. Organism\_taxid: 10711. Gene: r. Expressed in: escherichia coli. Expression\_system\_taxid: 562. CLASS: Mainly Alpha, ARCH: Orthogonal Bundle, TOPOL: Lysozyme, HOMOL: Lysozyme; cath \| 1xjtA00 \| 99.4 \| 3.5e-17 \| 4.3e-21 \| 101.1 \| 141 \| (3, 147) \| 159 \| (30, 182) \| 191 \| Lysozyme \| CATHCODE: 1.10.530.40 NAME: Lysozyme. Chain: a. Synonym: lysis protein, muramidase, endolysin, protein gp17. Engineered: yes SOURCE: Enterobacteria phage p1. Organism\_taxid: 10678. Gene: 17, lysa, lyz. Expressed in: escherichia coli. Expression\_system\_taxid: 562. CLASS: Mainly Alpha, ARCH: Orthogonal Bundle, TOPOL: Lysozyme, HOMOL: Lysozyme; cath \| 6h9dA00 \| 99.3 \| 1.3e-16 \| 1.6e-20 \| 93.2 \| 139 \| (3, 145) \| 159 \| (6, 148) \| 149 \| Lysozyme \| CATHCODE: 1.10.530.40 NAME: Lysozyme. Chain: a, b, c. Synonym: muramidase domain of spmx. Engineered: yes SOURCE: Asticcacaulis excentricus. Organism\_taxid: 78587. Gene: astex\_1112. Expressed in: escherichia coli bl21(de3). Expression\_system\_taxid: 469008. Expression\_system\_variant: ril. Expression\_system\_vector\_type: pet CLASS: Mainly Alpha, ARCH: Orthogonal Bundle, TOPOL: Lysozyme, HOMOL: Lysozyme; cath \| 2anvA00 \| 99.2 \| 1.7e-15 \| 2e-19 \| 87.6 \| 139 \| (2, 144) \| 159 \| (3, 145) \| 146 \| Lysozyme \| CATHCODE: 1.10.530.40 NAME: Lysozyme. Chain: a, b. Synonym: lysis protein, muramidase, endolysin. Engineered: yes. Mutation: yes SOURCE: Enterobacteria phage p22. Organism\_taxid: 10754. Gene: 19. Expressed in: escherichia coli. Expression\_system\_taxid: 562 CLASS: Mainly Alpha, ARCH: Orthogonal Bundle, TOPOL: Lysozyme, HOMOL: Lysozyme; cath \| 1jtmA00 \| 99.2 \| 2.5e-15 \| 3.1e-19 \| 91.0 \| 130 \| (7, 146) \| 159 \| (3, 156) \| 178 \| Lysozyme \| CATHCODE: 1.10.530.40 NAME: Lysozyme. Chain: a. Synonym: lysis protein. Engineered: yes. Mutation:yes SOURCE: Enterobacteria phage t4. Organism\_taxid: 10665. Gene: e. Expressed in: escherichia coli. Expression\_system\_taxid: 562. CLASS: Mainly Alpha, ARCH: Orthogonal Bundle, TOPOL: Lysozyme, HOMOL: Lysozyme; cath \| 2o7aA00 \| 99.0 \| 6.1e-14 \| 7.4e-18 \| 80.4 \| 92 \| (50, 148) \| 159 \| (1, 100) \| 124 \| Lysozyme \| CATHCODE: 1.10.530.40 NAME: Lysozyme. Chain: a. Synonym: lysis protein, muramidase, endolysin. Engineered: yes. Mutation: yes SOURCE: Enterobacteria phage t4. Organism\_taxid: 10665. Gene: e. Expressed in: escherichia coli. Expression\_system\_taxid: 562. CLASS: Mainly Alpha, ARCH: Orthogonal Bundle, TOPOL: Lysozyme, HOMOL: Lysozyme; cath \| 4evxA00 \| 98.9 \| 2.6e-13 \| 3.2e-17 \| 74.6 \| 94 \| (3, 100) \| 159 \| (7, 101) \| 106 \| Putative phage endolysin \| CATHCODE: 1.10.1740.240 NAME: Putative phage endolysin. Chain: a, b. Engineered: yes SOURCE: Salmonella enterica subsp. Enterica serovar typhimurium. Organism\_taxid: 99287. Strain: lt2. Gene: stm3605. Expressed in: escherichia coli.Expression\_system\_taxid: 469008. CLASS: Mainly Alpha, ARCH: Orthogonal Bundle, TOPOL: Rna Polymerase Sigma Factor; Chain: A, HOMOL: Rna Polymerase Sigma Factor; Chain: A; phrogs \| 7 \| 99.9 \| 1.2e-33 \| 1.6e-37 \| 221.0 \| 131 \| (8, 152) \| 159 \| (8, 145) \| 226 \| endolysin \| endolysin; Category: lysis; NC\_001629\_p14; phrogs \| 33083 \| 99.9 \| 2e-27 \| 2.2e-31 \| 187.5 \| 136 \| (8, 148) \| 159 \| (8, 145) \| 456 \| endolysin \| endolysin; Category: lysis; p415817 VI\_06217; phrogs \| 28234 \| 99.8 \| 7.3e-25 \| 8.2e-29 \| 170.5 \| 142 \| (8, 150) \| 159 \| (65, 216) \| 359 \| endolysin \| endolysin; Category: lysis; NC\_015465\_p23; phrogs \| 10457 \| 99.8 \| 3.4e-24 \| 3.9e-28 \| 143.3 \| 98 \| (47, 145) \| 159 \| (1, 99) \| 99 \| endolysin \| endolysin; Category: lysis; p305180 VI\_10042; phrogs \| 3617 \| 99.6 \| 5.7e-20 \| 6.3e-24 \| 142.8 \| 136 \| (8, 144) \| 159 \| (251, 404) \| 405 \| NA \| NA; Category: unknown function; KJ019096\_p210; phrogs \| 13211 \| 98.9 \| 1.2e-13 \| 1.4e-17 \| 115.7 \| 103 \| (8, 115) \| 159 \| (745, 867) \| 894 \| baseplate hub subunit and tail lysozyme \| baseplate hub subunit and tail lysozyme; Category: tail; NC\_019401\_p237; phrogs \| 16615 \| 98.7 \| 1.9e-12 \| 2.2e-16 \| 103.6 \| 134 \| (8, 152) \| 159 \| (200, 349) \| 519 \| peptidase \| peptidase; Category: other; KY996523\_p13; phrogs \| 3572 \| 98.6 \| 6.8e-12 \| 7.5e-16 \| 100.9 \| 131 \| (8, 148) \| 159 \| (268, 406) \| 653 \| endolysin \| endolysin; Category: lysis; KJ019101\_p13; phrogs \| 15410 \| 98.6 \| 8.1e-12 \| 9.2e-16 \| 95.4 \| 90 \| (18, 110) \| 159 \| (181, 280) \| 308 \| NA \| NA; Category: unknown function; NC\_025447\_p525; phrogs \| 7910 \| 98.3 \| 1.9e-10 \| 2.2e-14 \| 64.4 \| 40 \| (108, 147) \| 159 \| (2, 41) \| 42 \| endolysin \| endolysin; Category: lysis; p276551 VI\_10444; phrogs \| 10075 \| 97.6 \| 2.7e-08 \| 3e-12 \| 83.1 \| 105 \| (8, 113) \| 159 \| (859, 977) \| 1205 \| NA \| NA; Category: unknown function; KX349298\_p12; phrogs \| 6543 \| 96.7 \| 3.6e-06 \| 4.1e-10 \| 49.7 \| 30 \| (8, 41) \| 159 \| (28, 58) \| 62 \| NA \| NA; Category: unknown function; p415893 VI\_08530; phrogs \| 33534 \| 96.1 \| 2.7e-05 \| 3.1e-09 \| 48.3 \| 51 \| (74, 125) \| 159 \| (11, 71) \| 88 \| NA \| NA; Category: unknown function; p192930 VI\_05576; phrogs \| 35759 \| 95.2 \| 0.00024 \| 2.6e-08 \| 34.1 \| 24 \| (124, 147) \| 159 \| (2, 25) \| 26 \| NA \| NA; Category: unknown function; p377996 VI\_10733 |
| Top keywords  (threshold 1.00e-03 (evalue)) | **Lysozyme, endolysin, phage, lysis, in, a, yes, the, escherichia, coli** |
| Output files | ../../domain\_architecture/64\_FANPEZAQ\_CDS\_0064\_cath.hhr ../../domain\_architecture/64\_FANPEZAQ\_CDS\_0064\_merged.svg ../../domain\_architecture/64\_FANPEZAQ\_CDS\_0064\_ncbi-cd.hhr ../../domain\_architecture/64\_FANPEZAQ\_CDS\_0064\_pfam.hhr ../../domain\_architecture/64\_FANPEZAQ\_CDS\_0064\_phrogs.hhr |

### Identical protein sequences/structures

#### Search results

|  |  |
| --- | --- |
| Protein sequence databases searched | Pdb, Swissprot, Refseq |
| Identical proteins found | -- |
| Top keywords | -- |
| Output files | -- |

### Similar protein sequences/structures

#### Sequence similarity search results (HHblits)^1^

|  |  |
| --- | --- |
| Sequence databases searched | Uniclust, Pdb70 |
| Results, scheme(s)  (Top layers only, threshold 1.00e-03 (evalue)) | xml version="1.0" encoding="utf-8" standalone="no"?       2024-09-02T21:08:55.208612 image/svg+xml   Matplotlib v3.7.2, https://matplotlib.org/ |
| Results, table(s)  (threshold 1.00e-03 (evalue)) | db \| id \| prob \| evalue \| pvalue \| score \| cols \| query \| query\_len \| template \| template\_len \| name \| description; uniclust \| UniRef100\_A0A068Q6I2 \| 100.0 \| 1.1e-52 \| 2.1e-58 \| 282.7 \| 149 \| (1, 152) \| 159 \| (28, 176) \| 218 \| Endolysin \| Endolysin; uniclust \| UniRef100\_A0A0K8MD99 \| 100.0 \| 1.6e-51 \| 3.1e-57 \| 268.9 \| 146 \| (1, 149) \| 159 \| (13, 159) \| 172 \| Lysozyme \| Lysozyme; uniclust \| UniRef100\_A0A075MK28 \| 100.0 \| 7.1e-51 \| 1.4e-56 \| 275.6 \| 146 \| (1, 150) \| 159 \| (30, 175) \| 203 \| Lysozyme \| Lysozyme; uniclust \| UniRef100\_A0A024LSA6 \| 100.0 \| 4.3e-50 \| 8.3e-56 \| 283.4 \| 148 \| (1, 151) \| 159 \| (14, 165) \| 300 \| Lysozyme \| Lysozyme; uniclust \| UniRef100\_A0A014M974 \| 100.0 \| 6e-50 \| 1.2e-55 \| 265.7 \| 151 \| (1, 154) \| 159 \| (20, 174) \| 178 \| Lysozyme \| Lysozyme; uniclust \| UniRef100\_A0A2A4YRT5 \| 100.0 \| 1e-49 \| 2e-55 \| 280.4 \| 143 \| (1, 147) \| 159 \| (9, 155) \| 256 \| Lysozyme \| Lysozyme; uniclust \| UniRef100\_A0A0F4VJS5 \| 100.0 \| 1.7e-49 \| 3.3e-55 \| 270.5 \| 152 \| (1, 155) \| 159 \| (37, 188) \| 211 \| Lysozyme \| Lysozyme; uniclust \| UniRef100\_A0A0J1D667 \| 100.0 \| 1.2e-48 \| 2.2e-54 \| 256.4 \| 143 \| (1, 147) \| 159 \| (19, 161) \| 175 \| Lysozyme \| Lysozyme; uniclust \| UniRef100\_A0A009YLP0 \| 100.0 \| 3.6e-48 \| 6.8e-54 \| 273.8 \| 147 \| (1, 150) \| 159 \| (59, 206) \| 280 \| Lysozyme \| Lysozyme; uniclust \| UniRef100\_A0A013U6E4 \| 100.0 \| 3.9e-48 \| 7.4e-54 \| 269.4 \| 147 \| (1, 150) \| 159 \| (38, 187) \| 237 \| Lysozyme \| Lysozyme; uniclust \| UniRef100\_A0A060H344 \| 100.0 \| 6.3e-48 \| 1.2e-53 \| 266.3 \| 153 \| (1, 156) \| 159 \| (46, 198) \| 231 \| Lysozyme \| Lysozyme; uniclust \| UniRef100\_A0A009F987 \| 100.0 \| 1e-47 \| 2e-53 \| 267.5 \| 152 \| (1, 155) \| 159 \| (81, 236) \| 255 \| Lysozyme (Fragment) \| Lysozyme (Fragment); uniclust \| UniRef100\_A0A077XNR0 \| 100.0 \| 1.6e-47 \| 3.1e-53 \| 254.7 \| 148 \| (1, 151) \| 159 \| (16, 169) \| 193 \| Lysozyme \| Lysozyme; uniclust \| UniRef100\_A0A0C1Y3N8 \| 100.0 \| 6.1e-47 \| 1.1e-52 \| 258.1 \| 146 \| (2, 150) \| 159 \| (92, 241) \| 245 \| Lysozyme \| Lysozyme; uniclust \| UniRef100\_A0A0J1IN41 \| 100.0 \| 7.6e-47 \| 1.4e-52 \| 260.7 \| 148 \| (1, 149) \| 159 \| (27, 174) \| 292 \| Lysozyme \| Lysozyme; uniclust \| UniRef100\_A0A147I3J2 \| 100.0 \| 3.5e-46 \| 6.7e-52 \| 253.3 \| 149 \| (1, 152) \| 159 \| (29, 189) \| 211 \| Lysozyme \| Lysozyme; uniclust \| UniRef100\_A0A060BQW9 \| 100.0 \| 1.5e-45 \| 2.8e-51 \| 234.3 \| 145 \| (1, 150) \| 159 \| (5, 150) \| 152 \| Lysozyme (Fragment) \| Lysozyme (Fragment); uniclust \| UniRef100\_A0A0R2IIP4 \| 100.0 \| 1.5e-45 \| 2.8e-51 \| 248.9 \| 145 \| (1, 149) \| 159 \| (16, 160) \| 241 \| Lysozyme \| Lysozyme; uniclust \| UniRef100\_A0A010J5Z2 \| 100.0 \| 1.9e-45 \| 3.5e-51 \| 242.9 \| 144 \| (2, 148) \| 159 \| (52, 199) \| 205 \| Lysozyme \| Lysozyme; uniclust \| UniRef100\_A0A0Q4KN18 \| 100.0 \| 3.1e-45 \| 5.7e-51 \| 230.5 \| 142 \| (4, 149) \| 159 \| (7, 148) \| 150 \| Lysozyme \| Lysozyme; uniclust \| UniRef100\_A0A1C7HE45 \| 100.0 \| 3.3e-45 \| 6.1e-51 \| 246.5 \| 147 \| (1, 151) \| 159 \| (57, 203) \| 222 \| Lysozyme \| Lysozyme; uniclust \| UniRef100\_A0A258A958 \| 100.0 \| 1.7e-44 \| 3.2e-50 \| 252.3 \| 145 \| (1, 148) \| 159 \| (54, 202) \| 308 \| Lysozyme \| Lysozyme; uniclust \| UniRef100\_A0A5Q0BKX1 \| 100.0 \| 1.8e-44 \| 3.4e-50 \| 232.4 \| 151 \| (2, 155) \| 159 \| (4, 169) \| 173 \| Lysozyme \| Lysozyme; uniclust \| UniRef100\_A0A512IKZ3 \| 100.0 \| 3.7e-44 \| 7.1e-50 \| 253.0 \| 145 \| (1, 148) \| 159 \| (10, 157) \| 314 \| Lysozyme \| Lysozyme; uniclust \| UniRef100\_A0A1M3I817 \| 100.0 \| 5.3e-44 \| 1e-49 \| 239.0 \| 146 \| (2, 149) \| 159 \| (54, 200) \| 211 \| Lysozyme \| Lysozyme; uniclust \| UniRef100\_A0A2A2JXF0 \| 100.0 \| 5.5e-44 \| 1e-49 \| 229.3 \| 151 \| (1, 151) \| 159 \| (3, 154) \| 159 \| lysozyme \| lysozyme; uniclust \| UniRef100\_A0A0K1JXH1 \| 100.0 \| 1.2e-43 \| 2.3e-49 \| 233.6 \| 145 \| (1, 148) \| 159 \| (21, 170) \| 179 \| Lysozyme \| Lysozyme; uniclust \| UniRef100\_A0A293MYF4 \| 100.0 \| 1.3e-43 \| 2.4e-49 \| 249.1 \| 144 \| (2, 149) \| 159 \| (153, 296) \| 299 \| Lysozyme \| Lysozyme; uniclust \| UniRef100\_A0A0A1VBW2 \| 100.0 \| 1.8e-43 \| 3.4e-49 \| 244.2 \| 145 \| (1, 149) \| 159 \| (18, 162) \| 258 \| Lysozyme \| Lysozyme; uniclust \| UniRef100\_A0A8J7JSY3 \| 100.0 \| 1e-42 \| 1.9e-48 \| 245.7 \| 143 \| (2, 148) \| 159 \| (206, 348) \| 352 \| Lysozyme \| Lysozyme; uniclust \| UniRef100\_A0A0F3QFX2 \| 100.0 \| 2.1e-42 \| 4.1e-48 \| 229.2 \| 148 \| (2, 152) \| 159 \| (15, 167) \| 173 \| Lysozyme \| Lysozyme; uniclust \| UniRef100\_A0A066PZ20 \| 100.0 \| 2.2e-42 \| 4.1e-48 \| 237.7 \| 143 \| (3, 148) \| 159 \| (45, 192) \| 222 \| Lysozyme \| Lysozyme; uniclust \| UniRef100\_A0A1F3A5L4 \| 100.0 \| 2.5e-42 \| 4.6e-48 \| 240.2 \| 149 \| (1, 152) \| 159 \| (1, 153) \| 319 \| Lysozyme \| Lysozyme; uniclust \| UniRef100\_A0A0F2HCE7 \| 100.0 \| 3.1e-42 \| 5.9e-48 \| 233.1 \| 144 \| (3, 149) \| 159 \| (20, 170) \| 202 \| Lysozyme \| Lysozyme; uniclust \| UniRef100\_A0A0A0RR05 \| 100.0 \| 3.3e-42 \| 6.2e-48 \| 231.1 \| 142 \| (2, 146) \| 159 \| (82, 227) \| 231 \| Endolysin \| Endolysin; uniclust \| UniRef100\_A0A075DXC6 \| 100.0 \| 3.8e-42 \| 7.1e-48 \| 228.2 \| 143 \| (2, 147) \| 159 \| (68, 214) \| 216 \| Endolysin \| Endolysin; uniclust \| UniRef100\_A0A2E1XDB6 \| 100.0 \| 4.3e-42 \| 8.1e-48 \| 225.5 \| 150 \| (1, 154) \| 159 \| (5, 158) \| 181 \| Lysozyme \| Lysozyme; uniclust \| UniRef100\_A0A271LNT6 \| 100.0 \| 5e-42 \| 9.5e-48 \| 228.1 \| 147 \| (1, 150) \| 159 \| (3, 157) \| 211 \| Lysozyme \| Lysozyme; uniclust \| UniRef100\_A0A0M4LIC1 \| 100.0 \| 5.8e-42 \| 1.1e-47 \| 225.9 \| 148 \| (2, 152) \| 159 \| (3, 154) \| 220 \| Lysozyme \| Lysozyme; uniclust \| UniRef100\_A0A1Q6UX15 \| 100.0 \| 1e-41 \| 2e-47 \| 234.6 \| 137 \| (1, 148) \| 159 \| (4, 144) \| 240 \| Lysozyme \| Lysozyme; uniclust \| UniRef100\_A0A073CH50 \| 100.0 \| 2.9e-41 \| 5.3e-47 \| 235.2 \| 145 \| (2, 150) \| 159 \| (98, 242) \| 319 \| Lysozyme \| Lysozyme; uniclust \| UniRef100\_Q28PX4 \| 100.0 \| 3.7e-41 \| 6.8e-47 \| 236.4 \| 144 \| (1, 147) \| 159 \| (1, 148) \| 340 \| Lysozyme \| Lysozyme; uniclust \| UniRef100\_A0A1V3II14 \| 100.0 \| 4.5e-41 \| 8.4e-47 \| 226.4 \| 149 \| (1, 152) \| 159 \| (6, 158) \| 269 \| Lysozyme \| Lysozyme; uniclust \| UniRef100\_A0A1F6ZTS5 \| 100.0 \| 5.3e-41 \| 1e-46 \| 218.6 \| 145 \| (1, 149) \| 159 \| (8, 153) \| 161 \| lysozyme \| lysozyme; uniclust \| UniRef100\_A0A0A1VPW0 \| 100.0 \| 8.2e-41 \| 1.5e-46 \| 248.3 \| 144 \| (1, 147) \| 159 \| (40, 191) \| 553 \| Lysozyme \| Lysozyme; uniclust \| UniRef100\_A0A023ZU91 \| 100.0 \| 1e-40 \| 2e-46 \| 227.4 \| 144 \| (5, 152) \| 159 \| (47, 190) \| 203 \| Lysozyme \| Lysozyme; uniclust \| UniRef100\_A0A1X7FA58 \| 100.0 \| 1.3e-40 \| 2.5e-46 \| 225.4 \| 143 \| (3, 149) \| 159 \| (16, 162) \| 242 \| Lysozyme \| Lysozyme; uniclust \| UniRef100\_A0A059ZQR8 \| 100.0 \| 1.4e-40 \| 2.6e-46 \| 230.5 \| 146 \| (1, 151) \| 159 \| (36, 191) \| 233 \| Lysozyme \| Lysozyme; uniclust \| UniRef100\_A0A1G6VI05 \| 100.0 \| 1.5e-40 \| 2.8e-46 \| 226.2 \| 139 \| (1, 147) \| 159 \| (4, 148) \| 235 \| Lysozyme \| Lysozyme; uniclust \| UniRef100\_A0A126R939 \| 100.0 \| 2.3e-40 \| 4.4e-46 \| 220.0 \| 157 \| (1, 157) \| 159 \| (46, 204) \| 207 \| Lysozyme \| Lysozyme; uniclust \| UniRef100\_A0A3B0S578 \| 100.0 \| 2.5e-40 \| 4.6e-46 \| 224.5 \| 143 \| (1, 146) \| 159 \| (37, 183) \| 268 \| lysozyme \| lysozyme; uniclust \| UniRef100\_A0A1F1J0Y5 \| 100.0 \| 2.7e-40 \| 5e-46 \| 213.9 \| 144 \| (1, 148) \| 159 \| (24, 170) \| 172 \| Lysozyme \| Lysozyme; uniclust \| UniRef100\_A0A0N9SGQ7 \| 100.0 \| 3.6e-40 \| 6.7e-46 \| 222.9 \| 147 \| (1, 150) \| 159 \| (7, 154) \| 248 \| Endolysin \| Endolysin; uniclust \| UniRef100\_A0A241W0V3 \| 100.0 \| 4.1e-40 \| 7.6e-46 \| 219.7 \| 146 \| (1, 146) \| 159 \| (80, 227) \| 228 \| Lysozyme \| Lysozyme; uniclust \| UniRef100\_A0A167C5R0 \| 100.0 \| 1.2e-39 \| 2.3e-45 \| 222.1 \| 140 \| (1, 149) \| 159 \| (21, 164) \| 248 \| Lysozyme \| Lysozyme; uniclust \| UniRef100\_UPI0004B90056 \| 100.0 \| 1.9e-39 \| 3.6e-45 \| 215.7 \| 143 \| (1, 146) \| 159 \| (1, 144) \| 224 \| lysozyme \| lysozyme; uniclust \| UniRef100\_A0A078LD01 \| 100.0 \| 2.1e-39 \| 4.1e-45 \| 223.8 \| 139 \| (8, 149) \| 159 \| (58, 196) \| 250 \| Lysozyme \| Lysozyme; uniclust \| UniRef100\_A0A222G881 \| 100.0 \| 2.2e-39 \| 4.2e-45 \| 216.9 \| 141 \| (2, 146) \| 159 \| (84, 224) \| 225 \| Lysozyme \| Lysozyme; uniclust \| UniRef100\_UPI001EF6EE9B \| 100.0 \| 2.5e-39 \| 4.7e-45 \| 206.9 \| 142 \| (1, 146) \| 159 \| (44, 185) \| 203 \| lysozyme \| lysozyme; uniclust \| UniRef100\_A0A109BL19 \| 100.0 \| 2.8e-39 \| 5.4e-45 \| 223.4 \| 147 \| (1, 151) \| 159 \| (15, 168) \| 262 \| Lysozyme \| Lysozyme; uniclust \| UniRef100\_A0A086D2W4 \| 100.0 \| 5e-39 \| 9.4e-45 \| 220.8 \| 143 \| (7, 152) \| 159 \| (48, 191) \| 237 \| Lysozyme \| Lysozyme; uniclust \| UniRef100\_A0A0Q6SQZ2 \| 100.0 \| 5.5e-39 \| 1.1e-44 \| 228.3 \| 140 \| (2, 144) \| 159 \| (97, 244) \| 289 \| Lysozyme \| Lysozyme; uniclust \| UniRef100\_A0A1I0J1Q9 \| 100.0 \| 5.8e-39 \| 1.1e-44 \| 203.5 \| 145 \| (1, 148) \| 159 \| (1, 151) \| 183 \| Lysozyme \| Lysozyme; uniclust \| UniRef100\_UPI0007C7D1D0 \| 100.0 \| 7.2e-39 \| 1.3e-44 \| 218.2 \| 152 \| (1, 152) \| 159 \| (14, 169) \| 273 \| glycoside hydrolase family protein \| glycoside hydrolase family protein; uniclust \| UniRef100\_A0A1S9AW89 \| 100.0 \| 9.7e-39 \| 1.8e-44 \| 200.9 \| 142 \| (1, 145) \| 159 \| (5, 148) \| 149 \| Lysozyme \| Lysozyme; uniclust \| UniRef100\_A0A0Q9I3N9 \| 100.0 \| 1.3e-38 \| 2.5e-44 \| 224.4 \| 142 \| (1, 152) \| 159 \| (36, 182) \| 284 \| Lysozyme \| Lysozyme; uniclust \| UniRef100\_A0A351L291 \| 100.0 \| 1.3e-38 \| 2.5e-44 \| 203.5 \| 147 \| (1, 150) \| 159 \| (1, 156) \| 158 \| Lysozyme (Fragment) \| Lysozyme (Fragment); uniclust \| UniRef100\_A0A7Y0FFV2 \| 100.0 \| 1.9e-38 \| 3.5e-44 \| 215.1 \| 144 \| (7, 150) \| 159 \| (71, 231) \| 233 \| Lysozyme \| Lysozyme; uniclust \| UniRef100\_A0A011LWT5 \| 100.0 \| 1.9e-38 \| 3.6e-44 \| 218.6 \| 147 \| (2, 152) \| 159 \| (63, 210) \| 224 \| Lysozyme \| Lysozyme; uniclust \| UniRef100\_A0A060C0Y3 \| 100.0 \| 2.2e-38 \| 4.1e-44 \| 212.6 \| 135 \| (13, 150) \| 159 \| (48, 182) \| 210 \| Lysozyme (Fragment) \| Lysozyme (Fragment); uniclust \| UniRef100\_A0A060CFF5 \| 100.0 \| 2.1e-38 \| 4.1e-44 \| 221.7 \| 145 \| (4, 152) \| 159 \| (58, 202) \| 232 \| Lysozyme (Fragment) \| Lysozyme (Fragment); uniclust \| UniRef100\_A0A371R7Q5 \| 100.0 \| 2.3e-38 \| 4.2e-44 \| 224.4 \| 146 \| (1, 149) \| 159 \| (8, 168) \| 374 \| Lysozyme \| Lysozyme; uniclust \| UniRef100\_A0A1M6MED4 \| 100.0 \| 2.7e-38 \| 5.1e-44 \| 214.8 \| 143 \| (1, 149) \| 159 \| (3, 145) \| 229 \| Lysozyme \| Lysozyme; uniclust \| UniRef100\_A0A522A0K7 \| 100.0 \| 3.9e-38 \| 7.3e-44 \| 218.1 \| 143 \| (1, 147) \| 159 \| (17, 163) \| 249 \| Lysozyme \| Lysozyme; uniclust \| UniRef100\_A0A1C3FDG7 \| 100.0 \| 4.2e-38 \| 7.9e-44 \| 197.8 \| 125 \| (33, 157) \| 159 \| (1, 129) \| 136 \| Lysozyme \| Lysozyme; uniclust \| UniRef100\_A0A031JXV8 \| 100.0 \| 4.5e-38 \| 8.4e-44 \| 214.4 \| 147 \| (2, 150) \| 159 \| (81, 235) \| 242 \| Lysozyme \| Lysozyme; uniclust \| UniRef100\_A0A023VTC7 \| 100.0 \| 4.7e-38 \| 8.9e-44 \| 209.4 \| 154 \| (1, 155) \| 159 \| (20, 174) \| 191 \| Endolysin \| Endolysin; uniclust \| UniRef100\_A0A840NNX4 \| 100.0 \| 5e-38 \| 9.3e-44 \| 206.4 \| 148 \| (2, 152) \| 159 \| (5, 156) \| 222 \| Lysozyme (Fragment) \| Lysozyme (Fragment); uniclust \| UniRef100\_A0A930XJW8 \| 100.0 \| 5.6e-38 \| 1e-43 \| 215.8 \| 142 \| (2, 146) \| 159 \| (215, 356) \| 357 \| Lysozyme \| Lysozyme; uniclust \| UniRef100\_A0A931BXL2 \| 100.0 \| 1e-37 \| 1.9e-43 \| 205.6 \| 146 \| (1, 149) \| 159 \| (48, 197) \| 225 \| Lysozyme \| Lysozyme; uniclust \| UniRef100\_A0A1C5W0G4 \| 100.0 \| 1.5e-37 \| 2.8e-43 \| 223.8 \| 136 \| (1, 144) \| 159 \| (24, 164) \| 323 \| Lysozyme \| Lysozyme; uniclust \| UniRef100\_A0A072T9I2 \| 100.0 \| 1.7e-37 \| 3.2e-43 \| 210.4 \| 148 \| (2, 152) \| 159 \| (31, 185) \| 208 \| Glycoside hydrolase family 24 \| Glycoside hydrolase family 24; uniclust \| UniRef100\_A0A017SCH5 \| 100.0 \| 1.8e-37 \| 3.4e-43 \| 211.2 \| 147 \| (3, 152) \| 159 \| (68, 223) \| 236 \| lysozyme \| lysozyme; uniclust \| UniRef100\_A0A858RDI0 \| 100.0 \| 1.8e-37 \| 3.4e-43 \| 199.9 \| 142 \| (2, 148) \| 159 \| (41, 187) \| 193 \| Lysozyme \| Lysozyme; uniclust \| UniRef100\_A0A8J7K7G8 \| 100.0 \| 2.2e-37 \| 4e-43 \| 215.3 \| 142 \| (2, 146) \| 159 \| (251, 392) \| 393 \| Lysozyme \| Lysozyme; uniclust \| UniRef100\_A0A067NBT7 \| 100.0 \| 2.4e-37 \| 4.4e-43 \| 212.3 \| 147 \| (3, 152) \| 159 \| (100, 255) \| 260 \| lysozyme \| lysozyme; uniclust \| UniRef100\_A0A370N746 \| 100.0 \| 3.4e-37 \| 6.4e-43 \| 210.3 \| 151 \| (1, 151) \| 159 \| (74, 270) \| 274 \| Lysozyme \| Lysozyme; uniclust \| UniRef100\_A0A840I6B0 \| 100.0 \| 3.7e-37 \| 6.9e-43 \| 197.3 \| 144 \| (1, 148) \| 159 \| (3, 150) \| 192 \| Lysozyme \| Lysozyme; uniclust \| UniRef100\_A0A140K4G7 \| 100.0 \| 6.1e-37 \| 1.2e-42 \| 211.0 \| 147 \| (2, 151) \| 159 \| (78, 231) \| 237 \| Lysozyme \| Lysozyme; uniclust \| UniRef100\_A0A4D7CU28 \| 100.0 \| 6.3e-37 \| 1.2e-42 \| 201.3 \| 145 \| (1, 149) \| 159 \| (1, 145) \| 227 \| Lysozyme \| Lysozyme; uniclust \| UniRef100\_A0A949XFQ5 \| 100.0 \| 6.8e-37 \| 1.3e-42 \| 198.2 \| 145 \| (2, 149) \| 159 \| (5, 152) \| 185 \| Lysozyme \| Lysozyme; uniclust \| UniRef100\_A0A1A6FHX7 \| 100.0 \| 7.1e-37 \| 1.3e-42 \| 215.1 \| 143 \| (1, 151) \| 159 \| (4, 150) \| 316 \| Lysozyme \| Lysozyme; uniclust \| UniRef100\_A0A6J5LS01 \| 100.0 \| 7.3e-37 \| 1.4e-42 \| 197.0 \| 148 \| (1, 151) \| 159 \| (6, 178) \| 181 \| Endolysin \| Endolysin; uniclust \| UniRef100\_A0A653CSV6 \| 100.0 \| 8e-37 \| 1.5e-42 \| 201.5 \| 141 \| (1, 145) \| 159 \| (112, 252) \| 252 \| lysozyme \| lysozyme; uniclust \| UniRef100\_A0A397TNV2 \| 100.0 \| 8.3e-37 \| 1.5e-42 \| 196.4 \| 142 \| (2, 146) \| 159 \| (28, 173) \| 176 \| lysozyme \| lysozyme; uniclust \| UniRef100\_A0A0H5BHU8 \| 100.0 \| 1.2e-36 \| 2.2e-42 \| 209.9 \| 144 \| (1, 149) \| 159 \| (4, 161) \| 262 \| Lysozyme \| Lysozyme; uniclust \| UniRef100\_A0A1X7GNQ4 \| 100.0 \| 1.3e-36 \| 2.3e-42 \| 204.4 \| 150 \| (3, 152) \| 159 \| (4, 170) \| 254 \| Lysozyme \| Lysozyme; uniclust \| UniRef100\_A0A060QH24 \| 100.0 \| 1.6e-36 \| 3.1e-42 \| 205.1 \| 137 \| (6, 145) \| 159 \| (43, 189) \| 223 \| Lysozyme \| Lysozyme; uniclust \| UniRef100\_A0A1F9BL94 \| 100.0 \| 2.1e-36 \| 4e-42 \| 203.3 \| 148 \| (1, 150) \| 159 \| (11, 170) \| 193 \| Lysozyme \| Lysozyme; uniclust \| UniRef100\_A0A8J7EUD0 \| 100.0 \| 2.5e-36 \| 4.7e-42 \| 212.1 \| 142 \| (2, 146) \| 159 \| (285, 426) \| 429 \| Lysozyme \| Lysozyme; uniclust \| UniRef100\_A0A085GNC0 \| 100.0 \| 2.5e-36 \| 4.7e-42 \| 198.3 \| 148 \| (2, 153) \| 159 \| (33, 180) \| 198 \| Lysozyme \| Lysozyme; uniclust \| UniRef100\_A0A073IWM7 \| 100.0 \| 2.5e-36 \| 4.7e-42 \| 211.7 \| 144 \| (5, 152) \| 159 \| (147, 290) \| 299 \| Lysozyme \| Lysozyme; uniclust \| UniRef100\_A0A0P7Y7A2 \| 100.0 \| 2.9e-36 \| 5.5e-42 \| 211.8 \| 140 \| (1, 149) \| 159 \| (24, 167) \| 273 \| Lysozyme \| Lysozyme; uniclust \| UniRef100\_A0A1D9G253 \| 100.0 \| 3.2e-36 \| 6e-42 \| 203.5 \| 140 \| (2, 145) \| 159 \| (118, 264) \| 265 \| Lysozyme \| Lysozyme; uniclust \| UniRef100\_A0A285CW41 \| 100.0 \| 3.6e-36 \| 6.7e-42 \| 192.5 \| 149 \| (1, 152) \| 159 \| (1, 154) \| 179 \| Lysozyme \| Lysozyme; uniclust \| UniRef100\_A0A943WD90 \| 100.0 \| 4.7e-36 \| 8.6e-42 \| 202.6 \| 140 \| (2, 145) \| 159 \| (164, 303) \| 303 \| Lysozyme \| Lysozyme; uniclust \| UniRef100\_E0WSU1 \| 100.0 \| 5.9e-36 \| 1.1e-41 \| 197.2 \| 147 \| (3, 152) \| 159 \| (63, 209) \| 214 \| Lysozyme \| Lysozyme; uniclust \| UniRef100\_A0A2U3PSF2 \| 100.0 \| 6e-36 \| 1.1e-41 \| 203.4 \| 145 \| (2, 149) \| 159 \| (106, 257) \| 265 \| Lysozyme \| Lysozyme; uniclust \| UniRef100\_A0A4P6YRS8 \| 100.0 \| 6.1e-36 \| 1.1e-41 \| 201.6 \| 143 \| (1, 151) \| 159 \| (5, 147) \| 249 \| Lysozyme \| Lysozyme; uniclust \| UniRef100\_A0A067NV59 \| 100.0 \| 8.6e-36 \| 1.6e-41 \| 214.2 \| 147 \| (3, 152) \| 159 \| (122, 277) \| 349 \| lysozyme \| lysozyme; uniclust \| UniRef100\_A0A0R2IQS1 \| 100.0 \| 9.2e-36 \| 1.7e-41 \| 202.0 \| 148 \| (1, 152) \| 159 \| (6, 153) \| 312 \| Lysozyme \| Lysozyme; uniclust \| UniRef100\_A0A291BV27 \| 100.0 \| 9.9e-36 \| 1.8e-41 \| 201.8 \| 146 \| (1, 150) \| 159 \| (6, 151) \| 285 \| Lysozyme \| Lysozyme; uniclust \| UniRef100\_A0A1V4BM01 \| 100.0 \| 1.3e-35 \| 2.4e-41 \| 214.3 \| 146 \| (1, 149) \| 159 \| (6, 153) \| 473 \| Lysozyme \| Lysozyme; uniclust \| UniRef100\_A0A0L9Y629 \| 100.0 \| 1.4e-35 \| 2.6e-41 \| 208.5 \| 146 \| (3, 151) \| 159 \| (134, 285) \| 289 \| Lysozyme \| Lysozyme; uniclust \| UniRef100\_A0A0K1NGA9 \| 100.0 \| 1.8e-35 \| 3.3e-41 \| 206.0 \| 144 \| (1, 147) \| 159 \| (1, 144) \| 394 \| Lysozyme \| Lysozyme; uniclust \| UniRef100\_A0A1F9BXF9 \| 100.0 \| 1.9e-35 \| 3.5e-41 \| 209.9 \| 145 \| (1, 148) \| 159 \| (87, 231) \| 424 \| Lysozyme \| Lysozyme; uniclust \| UniRef100\_A0A412YY68 \| 100.0 \| 2.2e-35 \| 4e-41 \| 210.6 \| 140 \| (2, 145) \| 159 \| (341, 480) \| 480 \| Lysozyme \| Lysozyme; uniclust \| UniRef100\_A0A2A2JXU9 \| 100.0 \| 2.4e-35 \| 4.5e-41 \| 199.2 \| 138 \| (1, 142) \| 159 \| (1, 138) \| 302 \| lysozyme \| lysozyme; uniclust \| UniRef100\_A0A0A0CLL3 \| 100.0 \| 2.4e-35 \| 4.5e-41 \| 199.6 \| 149 \| (1, 149) \| 159 \| (59, 219) \| 223 \| Lysozyme \| Lysozyme; uniclust \| UniRef100\_UPI000EF53BDC \| 100.0 \| 2.5e-35 \| 4.6e-41 \| 208.7 \| 146 \| (1, 149) \| 159 \| (3, 155) \| 432 \| LysM peptidoglycan-binding domain-containing protein \| LysM peptidoglycan-binding domain-containing protein; uniclust \| UniRef100\_UPI001FD78962 \| 100.0 \| 2.5e-35 \| 4.7e-41 \| 200.2 \| 144 \| (1, 148) \| 159 \| (6, 149) \| 316 \| lysozyme \| lysozyme; uniclust \| UniRef100\_A0A8T4P871 \| 100.0 \| 2.5e-35 \| 4.7e-41 \| 192.9 \| 148 \| (2, 152) \| 159 \| (64, 211) \| 215 \| lysozyme \| lysozyme; uniclust \| UniRef100\_A0A117T1Q6 \| 100.0 \| 2.7e-35 \| 5e-41 \| 195.0 \| 144 \| (2, 151) \| 159 \| (4, 151) \| 216 \| Lysozyme \| Lysozyme; uniclust \| UniRef100\_A0A1U7IY75 \| 100.0 \| 3.8e-35 \| 7e-41 \| 212.5 \| 144 \| (1, 147) \| 159 \| (3, 158) \| 440 \| Lysozyme \| Lysozyme; uniclust \| UniRef100\_A0A1G3PUD1 \| 100.0 \| 3.8e-35 \| 7.1e-41 \| 195.7 \| 149 \| (1, 149) \| 159 \| (30, 187) \| 190 \| Lysozyme \| Lysozyme; uniclust \| UniRef100\_A0A0N0JEL9 \| 100.0 \| 3.9e-35 \| 7.3e-41 \| 200.9 \| 146 \| (3, 151) \| 159 \| (8, 180) \| 262 \| Lysozyme \| Lysozyme; uniclust \| UniRef100\_A0A0T9RE44 \| 100.0 \| 4.3e-35 \| 8e-41 \| 183.5 \| 142 \| (9, 154) \| 159 \| (2, 143) \| 152 \| Lysozyme \| Lysozyme; uniclust \| UniRef100\_A0A3P1XHR8 \| 100.0 \| 4.8e-35 \| 8.8e-41 \| 216.2 \| 144 \| (1, 147) \| 159 \| (350, 493) \| 722 \| Lysozyme \| Lysozyme; uniclust \| UniRef100\_A0A1C5P9H3 \| 100.0 \| 6.2e-35 \| 1.2e-40 \| 201.6 \| 138 \| (1, 147) \| 159 \| (22, 169) \| 276 \| Lysozyme \| Lysozyme; uniclust \| UniRef100\_A0A067T7N7 \| 100.0 \| 7.4e-35 \| 1.4e-40 \| 202.3 \| 147 \| (3, 152) \| 159 \| (151, 306) \| 310 \| lysozyme \| lysozyme; uniclust \| UniRef100\_A0A1H1FH24 \| 100.0 \| 1e-34 \| 1.9e-40 \| 186.8 \| 141 \| (1, 149) \| 159 \| (3, 147) \| 173 \| Lysozyme \| Lysozyme; uniclust \| UniRef100\_A0A8T3SPI5 \| 100.0 \| 1.1e-34 \| 2.1e-40 \| 189.2 \| 139 \| (2, 144) \| 159 \| (65, 205) \| 206 \| Lysozyme \| Lysozyme; uniclust \| UniRef100\_A0A1X2I864 \| 100.0 \| 1.2e-34 \| 2.3e-40 \| 200.6 \| 141 \| (2, 145) \| 159 \| (117, 262) \| 266 \| lysozyme \| lysozyme; uniclust \| UniRef100\_A0A127MKM6 \| 100.0 \| 1.3e-34 \| 2.5e-40 \| 193.9 \| 140 \| (6, 149) \| 159 \| (26, 177) \| 198 \| Lysozyme \| Lysozyme; uniclust \| UniRef100\_UPI000B336A28 \| 100.0 \| 1.9e-34 \| 3.4e-40 \| 195.9 \| 145 \| (1, 149) \| 159 \| (6, 150) \| 314 \| lysozyme \| lysozyme; uniclust \| UniRef100\_A0A1V0DCW7 \| 100.0 \| 2e-34 \| 3.6e-40 \| 186.9 \| 143 \| (3, 148) \| 159 \| (15, 164) \| 200 \| Lysozyme \| Lysozyme; uniclust \| UniRef100\_UPI001C04A56E \| 100.0 \| 2.3e-34 \| 4.3e-40 \| 177.8 \| 130 \| (14, 146) \| 159 \| (1, 134) \| 141 \| lysozyme \| lysozyme; uniclust \| UniRef100\_A0A1S8NI04 \| 100.0 \| 2.5e-34 \| 4.7e-40 \| 203.7 \| 148 \| (2, 152) \| 159 \| (139, 288) \| 292 \| Lysozyme \| Lysozyme; uniclust \| UniRef100\_A0A1M3KMN9 \| 100.0 \| 2.9e-34 \| 5.3e-40 \| 185.2 \| 147 \| (1, 147) \| 159 \| (2, 154) \| 196 \| Lysozyme \| Lysozyme; uniclust \| UniRef100\_A0A2D0K3Q9 \| 100.0 \| 3.4e-34 \| 6.4e-40 \| 174.5 \| 112 \| (1, 112) \| 159 \| (2, 117) \| 119 \| Lysozyme \| Lysozyme; uniclust \| UniRef100\_A0A927EH04 \| 100.0 \| 3.7e-34 \| 6.8e-40 \| 210.0 \| 144 \| (3, 150) \| 159 \| (442, 585) \| 587 \| Lysozyme \| Lysozyme; uniclust \| UniRef100\_A0A6H9GT10 \| 100.0 \| 4.1e-34 \| 7.4e-40 \| 189.2 \| 145 \| (1, 148) \| 159 \| (1, 153) \| 254 \| Lysozyme \| Lysozyme; uniclust \| UniRef100\_A0A1Q4S2E5 \| 100.0 \| 4.4e-34 \| 8.1e-40 \| 192.2 \| 145 \| (2, 149) \| 159 \| (102, 254) \| 258 \| Lysozyme \| Lysozyme; uniclust \| UniRef100\_A0A2W4WM75 \| 100.0 \| 4.7e-34 \| 8.8e-40 \| 175.8 \| 110 \| (1, 114) \| 159 \| (12, 121) \| 124 \| Lysozyme (Fragment) \| Lysozyme (Fragment); uniclust \| UniRef100\_A0A6P1B8L0 \| 100.0 \| 5.4e-34 \| 1e-39 \| 200.1 \| 147 \| (1, 149) \| 159 \| (5, 169) \| 295 \| Lysozyme \| Lysozyme; uniclust \| UniRef100\_A0A0P0NZY8 \| 100.0 \| 5.5e-34 \| 1.1e-39 \| 213.4 \| 145 \| (1, 149) \| 159 \| (23, 171) \| 455 \| Lysozyme \| Lysozyme; uniclust \| UniRef100\_A0A2A4Z0X3 \| 100.0 \| 5.8e-34 \| 1.1e-39 \| 180.9 \| 142 \| (2, 147) \| 159 \| (3, 149) \| 188 \| Lysozyme \| Lysozyme; uniclust \| UniRef100\_A0A239Q0D5 \| 100.0 \| 5.9e-34 \| 1.1e-39 \| 193.1 \| 141 \| (2, 147) \| 159 \| (88, 228) \| 229 \| Lysozyme \| Lysozyme; uniclust \| UniRef100\_UPI001F5504E3 \| 100.0 \| 6.3e-34 \| 1.2e-39 \| 202.0 \| 143 \| (3, 149) \| 159 \| (302, 444) \| 447 \| lysozyme \| lysozyme; uniclust \| UniRef100\_A0A9E8ZDG3 \| 100.0 \| 6.7e-34 \| 1.2e-39 \| 206.5 \| 144 \| (2, 149) \| 159 \| (441, 584) \| 587 \| Lysozyme \| Lysozyme; uniclust \| UniRef100\_A0A0K1H0L4 \| 100.0 \| 8e-34 \| 1.5e-39 \| 170.1 \| 103 \| (43, 145) \| 159 \| (3, 105) \| 106 \| Lysozyme \| Lysozyme; uniclust \| UniRef100\_A0A022G329 \| 100.0 \| 1.1e-33 \| 2.1e-39 \| 195.8 \| 139 \| (7, 150) \| 159 \| (73, 211) \| 231 \| Lysozyme \| Lysozyme; uniclust \| UniRef100\_A0A2W5V772 \| 100.0 \| 1.5e-33 \| 2.7e-39 \| 204.8 \| 144 \| (3, 149) \| 159 \| (74, 217) \| 590 \| Lysozyme \| Lysozyme; uniclust \| UniRef100\_A0A5S9RTX0 \| 100.0 \| 1.7e-33 \| 3.1e-39 \| 189.9 \| 144 \| (2, 151) \| 159 \| (65, 224) \| 225 \| Lysozyme \| Lysozyme; uniclust \| UniRef100\_A0A5K7ZI99 \| 100.0 \| 1.9e-33 \| 3.6e-39 \| 197.2 \| 146 \| (2, 151) \| 159 \| (4, 155) \| 383 \| Lysozyme \| Lysozyme; uniclust \| UniRef100\_A0A093XAS9 \| 100.0 \| 2.1e-33 \| 4e-39 \| 201.6 \| 144 \| (3, 149) \| 159 \| (138, 290) \| 377 \| lysozyme \| lysozyme; uniclust \| UniRef100\_A0A926ZD07 \| 100.0 \| 2.2e-33 \| 4.1e-39 \| 190.3 \| 144 \| (3, 149) \| 159 \| (70, 217) \| 308 \| Lysozyme \| Lysozyme; uniclust \| UniRef100\_A0A1X7MCZ0 \| 100.0 \| 2.8e-33 \| 5.4e-39 \| 172.3 \| 102 \| (47, 148) \| 159 \| (11, 112) \| 113 \| Lysozyme \| Lysozyme; uniclust \| UniRef100\_A0A5C7LRY7 \| 100.0 \| 3.3e-33 \| 6e-39 \| 188.0 \| 143 \| (1, 146) \| 159 \| (59, 205) \| 288 \| Lysozyme \| Lysozyme; uniclust \| UniRef100\_A0A0F9RCT0 \| 100.0 \| 3.6e-33 \| 6.5e-39 \| 181.9 \| 149 \| (1, 152) \| 159 \| (1, 158) \| 225 \| lysozyme \| lysozyme; uniclust \| UniRef100\_A0A1H2JQ13 \| 100.0 \| 4e-33 \| 7.5e-39 \| 194.9 \| 149 \| (1, 149) \| 159 \| (113, 273) \| 276 \| Lysozyme \| Lysozyme; uniclust \| UniRef100\_A0A7X9HPJ2 \| 100.0 \| 4.4e-33 \| 8e-39 \| 181.4 \| 145 \| (2, 149) \| 159 \| (59, 217) \| 224 \| Lysozyme \| Lysozyme; uniclust \| UniRef100\_A0A059DT11 \| 100.0 \| 4.7e-33 \| 8.8e-39 \| 206.4 \| 145 \| (1, 149) \| 159 \| (28, 177) \| 428 \| Lysozyme \| Lysozyme; uniclust \| UniRef100\_A0A1B1KLF1 \| 100.0 \| 5.4e-33 \| 1e-38 \| 177.5 \| 143 \| (2, 148) \| 159 \| (27, 169) \| 171 \| Lysozyme \| Lysozyme; uniclust \| UniRef100\_A0A077FK68 \| 100.0 \| 5.7e-33 \| 1.1e-38 \| 171.6 \| 115 \| (43, 157) \| 159 \| (3, 117) \| 119 \| Lysozyme \| Lysozyme; uniclust \| UniRef100\_UPI00215FDBC0 \| 100.0 \| 6.4e-33 \| 1.2e-38 \| 181.8 \| 123 \| (20, 146) \| 159 \| (112, 234) \| 235 \| phage portal protein \| phage portal protein; uniclust \| UniRef100\_A0A3E2N5A0 \| 100.0 \| 8e-33 \| 1.5e-38 \| 183.7 \| 136 \| (1, 145) \| 159 \| (97, 238) \| 239 \| Lysozyme \| Lysozyme; uniclust \| UniRef100\_A0A1G4TH47 \| 100.0 \| 8e-33 \| 1.5e-38 \| 217.0 \| 147 \| (2, 152) \| 159 \| (7, 157) \| 832 \| Lysozyme \| Lysozyme; uniclust \| UniRef100\_A0A3S0BYI5 \| 100.0 \| 8.5e-33 \| 1.6e-38 \| 178.1 \| 143 \| (3, 149) \| 159 \| (6, 148) \| 207 \| Lysozyme \| Lysozyme; uniclust \| UniRef100\_A0A1E5TA89 \| 100.0 \| 9.1e-33 \| 1.7e-38 \| 188.3 \| 145 \| (2, 152) \| 159 \| (84, 244) \| 245 \| Lysozyme \| Lysozyme; uniclust \| UniRef100\_A0A1B1IHU6 \| 100.0 \| 9.3e-33 \| 1.7e-38 \| 183.6 \| 145 \| (1, 148) \| 159 \| (4, 153) \| 233 \| Lysozyme \| Lysozyme; uniclust \| UniRef100\_A0A6I5R3J3 \| 100.0 \| 1e-32 \| 1.9e-38 \| 200.5 \| 145 \| (1, 148) \| 159 \| (244, 392) \| 456 \| Lysozyme \| Lysozyme; uniclust \| UniRef100\_A0A271J3C8 \| 100.0 \| 1.1e-32 \| 2.1e-38 \| 181.6 \| 145 \| (2, 149) \| 159 \| (10, 155) \| 227 \| Lysozyme \| Lysozyme; uniclust \| UniRef100\_A0A350KNR5 \| 100.0 \| 1.4e-32 \| 2.6e-38 \| 176.5 \| 140 \| (3, 145) \| 159 \| (25, 165) \| 167 \| Lysozyme \| Lysozyme; uniclust \| UniRef100\_A0A0S3PZQ8 \| 100.0 \| 1.4e-32 \| 2.7e-38 \| 184.0 \| 138 \| (8, 149) \| 159 \| (21, 160) \| 220 \| Lysozyme \| Lysozyme; uniclust \| UniRef100\_A0A073J6L3 \| 100.0 \| 1.4e-32 \| 2.7e-38 \| 197.7 \| 144 \| (5, 152) \| 159 \| (145, 293) \| 327 \| Lysozyme \| Lysozyme; uniclust \| UniRef100\_A0A0C9WJX8 \| 100.0 \| 1.5e-32 \| 2.9e-38 \| 213.0 \| 147 \| (3, 152) \| 159 \| (476, 631) \| 635 \| lysozyme \| lysozyme; uniclust \| UniRef100\_A0A7J6JRX3 \| 100.0 \| 1.7e-32 \| 3.1e-38 \| 186.8 \| 143 \| (2, 147) \| 159 \| (6, 148) \| 316 \| lysozyme \| lysozyme; uniclust \| UniRef100\_A0A7W0JH81 \| 100.0 \| 1.7e-32 \| 3.2e-38 \| 175.5 \| 144 \| (2, 148) \| 159 \| (3, 153) \| 197 \| Lysozyme \| Lysozyme; uniclust \| UniRef100\_A0A1W1XZS6 \| 100.0 \| 1.7e-32 \| 3.3e-38 \| 175.7 \| 135 \| (18, 153) \| 159 \| (5, 147) \| 154 \| Lysozyme \| Lysozyme; uniclust \| UniRef100\_A0A226E4N2 \| 100.0 \| 2.1e-32 \| 3.9e-38 \| 189.1 \| 141 \| (4, 147) \| 159 \| (50, 201) \| 290 \| lysozyme \| lysozyme; uniclust \| UniRef100\_A0A261KX80 \| 100.0 \| 2.5e-32 \| 4.7e-38 \| 199.6 \| 144 \| (1, 147) \| 159 \| (1, 152) \| 497 \| Lysozyme (Fragment) \| Lysozyme (Fragment); uniclust \| UniRef100\_A0A2U2P948 \| 99.9 \| 2.9e-32 \| 5.4e-38 \| 182.6 \| 140 \| (1, 144) \| 159 \| (96, 239) \| 247 \| Lysozyme (Fragment) \| Lysozyme (Fragment); uniclust \| UniRef100\_UPI00200AF8DA \| 99.9 \| 3.5e-32 \| 6.5e-38 \| 179.3 \| 146 \| (1, 149) \| 159 \| (1, 153) \| 244 \| glycoside hydrolase family protein \| glycoside hydrolase family protein; uniclust \| UniRef100\_A0A258GXH3 \| 99.9 \| 4.4e-32 \| 8.2e-38 \| 185.0 \| 140 \| (2, 146) \| 159 \| (98, 241) \| 245 \| Lysozyme \| Lysozyme; uniclust \| UniRef100\_A0A2D6VSD9 \| 99.9 \| 4.5e-32 \| 8.5e-38 \| 204.3 \| 148 \| (1, 148) \| 159 \| (80, 231) \| 490 \| lysozyme \| lysozyme; uniclust \| UniRef100\_A0A381GPR2 \| 99.9 \| 5.4e-32 \| 1e-37 \| 183.8 \| 143 \| (2, 148) \| 159 \| (123, 265) \| 266 \| Lysozyme \| Lysozyme; uniclust \| UniRef100\_A0A158G2K7 \| 99.9 \| 5.9e-32 \| 1.1e-37 \| 162.2 \| 102 \| (46, 147) \| 159 \| (3, 104) \| 106 \| Lysozyme \| Lysozyme; uniclust \| UniRef100\_L0EV97 \| 99.9 \| 6.4e-32 \| 1.2e-37 \| 176.7 \| 145 \| (2, 149) \| 159 \| (10, 158) \| 230 \| Lysozyme \| Lysozyme; uniclust \| UniRef100\_A0A0N7CEK9 \| 99.9 \| 6.2e-32 \| 1.2e-37 \| 172.3 \| 126 \| (21, 152) \| 159 \| (3, 137) \| 138 \| Lysozyme \| Lysozyme; uniclust \| UniRef100\_A0A014PYD0 \| 99.9 \| 7e-32 \| 1.3e-37 \| 176.1 \| 145 \| (2, 150) \| 159 \| (37, 181) \| 197 \| Lysozyme \| Lysozyme; uniclust \| UniRef100\_A0A7W5EKN2 \| 99.9 \| 7.1e-32 \| 1.3e-37 \| 175.0 \| 127 \| (7, 145) \| 159 \| (26, 152) \| 154 \| Lysozyme \| Lysozyme; uniclust \| UniRef100\_A0A0H4TH14 \| 99.9 \| 7.4e-32 \| 1.4e-37 \| 179.5 \| 145 \| (2, 150) \| 159 \| (30, 182) \| 194 \| Endolysin \| Endolysin; uniclust \| UniRef100\_A0A858RGZ4 \| 99.9 \| 7.7e-32 \| 1.4e-37 \| 181.1 \| 142 \| (3, 149) \| 159 \| (131, 277) \| 282 \| Lysozyme \| Lysozyme; uniclust \| UniRef100\_A0A8S5N3I7 \| 99.9 \| 8.7e-32 \| 1.6e-37 \| 188.6 \| 148 \| (1, 151) \| 159 \| (10, 162) \| 389 \| Endolysin \| Endolysin; uniclust \| UniRef100\_A0A1F8IHC7 \| 99.9 \| 8.6e-32 \| 1.6e-37 \| 208.2 \| 142 \| (1, 149) \| 159 \| (27, 168) \| 624 \| Lysozyme \| Lysozyme; uniclust \| UniRef100\_A0A239PWS9 \| 99.9 \| 9.9e-32 \| 1.9e-37 \| 190.6 \| 145 \| (1, 149) \| 159 \| (17, 165) \| 339 \| Lysozyme \| Lysozyme; uniclust \| UniRef100\_F8WVD9 \| 99.9 \| 1.1e-31 \| 2e-37 \| 170.2 \| 139 \| (2, 143) \| 159 \| (38, 181) \| 183 \| Lysozyme \| Lysozyme; uniclust \| UniRef100\_A0A1Q3Q7Z8 \| 99.9 \| 1.2e-31 \| 2.3e-37 \| 178.5 \| 145 \| (1, 149) \| 159 \| (5, 173) \| 187 \| Lysozyme \| Lysozyme; uniclust \| UniRef100\_A0A7C4STG1 \| 99.9 \| 1.3e-31 \| 2.5e-37 \| 185.9 \| 145 \| (2, 149) \| 159 \| (205, 363) \| 370 \| Lysozyme \| Lysozyme; uniclust \| UniRef100\_A0A0A2W020 \| 99.9 \| 1.7e-31 \| 3.1e-37 \| 193.5 \| 139 \| (4, 145) \| 159 \| (421, 560) \| 561 \| lysozyme \| lysozyme; uniclust \| UniRef100\_A0A9E7MUM9 \| 99.9 \| 1.9e-31 \| 3.4e-37 \| 176.4 \| 146 \| (2, 150) \| 159 \| (15, 160) \| 249 \| Lysozyme \| Lysozyme; uniclust \| UniRef100\_A0A0C9W3W6 \| 99.9 \| 1.9e-31 \| 3.7e-37 \| 200.1 \| 146 \| (4, 152) \| 159 \| (364, 518) \| 522 \| lysozyme \| lysozyme; uniclust \| UniRef100\_A0A2E6Y944 \| 99.9 \| 3e-31 \| 5.6e-37 \| 175.5 \| 149 \| (1, 152) \| 159 \| (1, 161) \| 225 \| Lysozyme \| Lysozyme; uniclust \| UniRef100\_A0A926PFU6 \| 99.9 \| 3.3e-31 \| 6e-37 \| 176.6 \| 141 \| (2, 146) \| 159 \| (120, 262) \| 263 \| Lysozyme \| Lysozyme; uniclust \| UniRef100\_A0A060H052 \| 99.9 \| 3.5e-31 \| 6.6e-37 \| 178.5 \| 142 \| (4, 150) \| 159 \| (54, 197) \| 208 \| Lysozyme \| Lysozyme; uniclust \| UniRef100\_A0A060DST8 \| 99.9 \| 3.9e-31 \| 7.2e-37 \| 179.8 \| 140 \| (5, 148) \| 159 \| (19, 175) \| 288 \| Lysozyme \| Lysozyme; uniclust \| UniRef100\_UPI0007659C9C \| 99.9 \| 4e-31 \| 7.3e-37 \| 163.9 \| 123 \| (1, 126) \| 159 \| (2, 126) \| 146 \| lysozyme \| lysozyme; uniclust \| UniRef100\_A0A6M0G4D1 \| 99.9 \| 4.4e-31 \| 8.1e-37 \| 192.4 \| 146 \| (1, 149) \| 159 \| (5, 161) \| 474 \| Lysozyme \| Lysozyme; uniclust \| UniRef100\_A0A410DNY0 \| 99.9 \| 4.8e-31 \| 8.9e-37 \| 185.3 \| 145 \| (2, 149) \| 159 \| (168, 317) \| 320 \| Lysozyme \| Lysozyme; uniclust \| UniRef100\_A0A2D5M4Q0 \| 99.9 \| 4.8e-31 \| 9e-37 \| 170.0 \| 137 \| (4, 144) \| 159 \| (45, 181) \| 182 \| Lysozyme \| Lysozyme; uniclust \| UniRef100\_A0A2E4S589 \| 99.9 \| 4.8e-31 \| 9.2e-37 \| 200.7 \| 147 \| (2, 152) \| 159 \| (13, 163) \| 502 \| Lysozyme \| Lysozyme; uniclust \| UniRef100\_UPI000979CF0A \| 99.9 \| 5.1e-31 \| 9.4e-37 \| 176.6 \| 142 \| (2, 150) \| 159 \| (4, 151) \| 245 \| lysozyme \| lysozyme; uniclust \| UniRef100\_A0A0Q9KM09 \| 99.9 \| 5.6e-31 \| 1e-36 \| 189.0 \| 146 \| (1, 149) \| 159 \| (60, 215) \| 408 \| Lysozyme \| Lysozyme; uniclust \| UniRef100\_UPI0015658B88 \| 99.9 \| 5.7e-31 \| 1e-36 \| 169.3 \| 146 \| (3, 149) \| 159 \| (5, 151) \| 202 \| lysozyme \| lysozyme; uniclust \| UniRef100\_A0A087M4E4 \| 99.9 \| 5.7e-31 \| 1.1e-36 \| 181.3 \| 142 \| (6, 151) \| 159 \| (58, 209) \| 236 \| Lysozyme \| Lysozyme; uniclust \| UniRef100\_A0A6J5M318 \| 99.9 \| 7.1e-31 \| 1.3e-36 \| 173.2 \| 141 \| (3, 146) \| 159 \| (59, 213) \| 214 \| Endolysin \| Endolysin; uniclust \| UniRef100\_A0A7C3SVB1 \| 99.9 \| 7.2e-31 \| 1.3e-36 \| 170.6 \| 144 \| (2, 148) \| 159 \| (62, 216) \| 217 \| Lysozyme \| Lysozyme; uniclust \| UniRef100\_A0A841WQ48 \| 99.9 \| 7.3e-31 \| 1.3e-36 \| 172.6 \| 143 \| (2, 148) \| 159 \| (5, 147) \| 237 \| Lysozyme \| Lysozyme; uniclust \| UniRef100\_A0A1V5YWE3 \| 99.9 \| 8.6e-31 \| 1.6e-36 \| 169.1 \| 147 \| (1, 152) \| 159 \| (1, 148) \| 207 \| Lysozyme \| Lysozyme; uniclust \| UniRef100\_A0A929A6N4 \| 99.9 \| 8.9e-31 \| 1.6e-36 \| 201.3 \| 144 \| (3, 149) \| 159 \| (754, 901) \| 987 \| Lysozyme \| Lysozyme; uniclust \| UniRef100\_A0A158I7H1 \| 99.9 \| 1e-30 \| 1.9e-36 \| 177.2 \| 147 \| (2, 156) \| 159 \| (37, 201) \| 204 \| Lysozyme \| Lysozyme; uniclust \| UniRef100\_UPI001BD4AB69 \| 99.9 \| 1e-30 \| 1.9e-36 \| 190.3 \| 147 \| (1, 150) \| 159 \| (1, 155) \| 511 \| peptidoglycan amidohydrolase family protein \| peptidoglycan amidohydrolase family protein; uniclust \| UniRef100\_A0A098RJQ8 \| 99.9 \| 1e-30 \| 1.9e-36 \| 192.9 \| 146 \| (1, 150) \| 159 \| (58, 219) \| 393 \| Lysozyme \| Lysozyme; uniclust \| UniRef100\_UPI001E4B06EF \| 99.9 \| 1.1e-30 \| 2e-36 \| 174.0 \| 149 \| (3, 151) \| 159 \| (52, 212) \| 213 \| lysozyme \| lysozyme; uniclust \| UniRef100\_A0A7V8YU39 \| 99.9 \| 1.2e-30 \| 2.3e-36 \| 173.8 \| 141 \| (3, 146) \| 159 \| (113, 261) \| 262 \| Lysozyme \| Lysozyme; uniclust \| UniRef100\_A0A348HCF7 \| 99.9 \| 1.7e-30 \| 3e-36 \| 176.2 \| 148 \| (3, 150) \| 159 \| (4, 152) \| 300 \| Lysozyme \| Lysozyme; uniclust \| UniRef100\_A0A1M5TXL6 \| 99.9 \| 1.6e-30 \| 3.1e-36 \| 193.4 \| 149 \| (1, 153) \| 159 \| (31, 187) \| 433 \| Lysozyme \| Lysozyme; uniclust \| UniRef100\_A0A7Y4UQW0 \| 99.9 \| 1.8e-30 \| 3.3e-36 \| 165.3 \| 144 \| (2, 148) \| 159 \| (7, 163) \| 167 \| Lysozyme \| Lysozyme; uniclust \| UniRef100\_A0A9D1LFB1 \| 99.9 \| 1.9e-30 \| 3.5e-36 \| 165.5 \| 137 \| (8, 148) \| 159 \| (46, 182) \| 190 \| Lysozyme \| Lysozyme; uniclust \| UniRef100\_A0A0Q7TTY5 \| 99.9 \| 2e-30 \| 3.7e-36 \| 176.7 \| 150 \| (3, 152) \| 159 \| (65, 220) \| 239 \| Lysozyme \| Lysozyme; uniclust \| UniRef100\_A0A3Q9BKE1 \| 99.9 \| 2.2e-30 \| 4.1e-36 \| 192.0 \| 148 \| (1, 152) \| 159 \| (6, 153) \| 715 \| Lysozyme \| Lysozyme; uniclust \| UniRef100\_A0A5E4NVC9 \| 99.9 \| 3.1e-30 \| 5.7e-36 \| 172.1 \| 142 \| (3, 148) \| 159 \| (5, 146) \| 264 \| Lysozyme \| Lysozyme; uniclust \| UniRef100\_A0A0F9HV13 \| 99.9 \| 3.4e-30 \| 6.4e-36 \| 174.4 \| 137 \| (7, 147) \| 159 \| (89, 227) \| 229 \| lysozyme \| lysozyme; uniclust \| UniRef100\_A0A8J5ASL3 \| 99.9 \| 3.9e-30 \| 7.3e-36 \| 165.3 \| 141 \| (3, 146) \| 159 \| (36, 180) \| 183 \| lysozyme \| lysozyme; uniclust \| UniRef100\_A0A9D8PN11 \| 99.9 \| 4.6e-30 \| 8.5e-36 \| 178.3 \| 154 \| (1, 154) \| 159 \| (1, 163) \| 325 \| Lysozyme \| Lysozyme; uniclust \| UniRef100\_A0A2I5ARC6 \| 99.9 \| 4.5e-30 \| 8.5e-36 \| 188.3 \| 143 \| (6, 148) \| 159 \| (104, 257) \| 404 \| Lysozyme \| Lysozyme; uniclust \| UniRef100\_A0A0B5DZ45 \| 99.9 \| 4.7e-30 \| 8.7e-36 \| 178.1 \| 137 \| (7, 147) \| 159 \| (137, 274) \| 275 \| Lysozyme \| Lysozyme; uniclust \| UniRef100\_A0A7R9KK34 \| 99.9 \| 5.2e-30 \| 9.6e-36 \| 178.2 \| 145 \| (2, 150) \| 159 \| (19, 166) \| 341 \| lysozyme \| lysozyme; uniclust \| UniRef100\_A0A1E5TEK2 \| 99.9 \| 5.6e-30 \| 1e-35 \| 190.7 \| 145 \| (1, 152) \| 159 \| (322, 476) \| 476 \| Lysozyme \| Lysozyme; uniclust \| UniRef100\_A0A258FCB1 \| 99.9 \| 5.8e-30 \| 1.1e-35 \| 172.7 \| 143 \| (3, 150) \| 159 \| (4, 150) \| 244 \| Lysozyme \| Lysozyme; uniclust \| UniRef100\_A0A843K231 \| 99.9 \| 6e-30 \| 1.1e-35 \| 180.1 \| 146 \| (2, 150) \| 159 \| (28, 183) \| 413 \| lysozyme \| lysozyme; uniclust \| UniRef100\_A0A376CQV8 \| 99.9 \| 5.9e-30 \| 1.1e-35 \| 157.4 \| 131 \| (13, 147) \| 159 \| (2, 132) \| 134 \| Lysozyme \| Lysozyme; uniclust \| UniRef100\_A0A8S5NUM4 \| 99.9 \| 7.3e-30 \| 1.4e-35 \| 173.6 \| 145 \| (1, 152) \| 159 \| (1, 148) \| 258 \| Endolysin \| Endolysin; uniclust \| UniRef100\_A0A7R9M799 \| 99.9 \| 7.5e-30 \| 1.4e-35 \| 168.5 \| 140 \| (2, 145) \| 159 \| (101, 243) \| 243 \| lysozyme \| lysozyme; uniclust \| UniRef100\_A0A0U3A7J9 \| 99.9 \| 8.8e-30 \| 1.6e-35 \| 187.1 \| 141 \| (7, 147) \| 159 \| (320, 469) \| 470 \| Lysozyme \| Lysozyme; uniclust \| UniRef100\_A0A1F2Z7P5 \| 99.9 \| 8.8e-30 \| 1.6e-35 \| 162.6 \| 141 \| (4, 147) \| 159 \| (18, 159) \| 169 \| Lysozyme \| Lysozyme; uniclust \| UniRef100\_A0A2W5P4F4 \| 99.9 \| 9.3e-30 \| 1.7e-35 \| 183.2 \| 146 \| (2, 147) \| 159 \| (282, 428) \| 433 \| Lysozyme \| Lysozyme; uniclust \| UniRef100\_A0A2G2MWX1 \| 99.9 \| 9.8e-30 \| 1.8e-35 \| 177.5 \| 143 \| (1, 146) \| 159 \| (1, 147) \| 381 \| Lysozyme \| Lysozyme; uniclust \| UniRef100\_A0A0H0XNY4 \| 99.9 \| 9.6e-30 \| 1.8e-35 \| 172.2 \| 146 \| (1, 150) \| 159 \| (50, 202) \| 221 \| Lysozyme \| Lysozyme; uniclust \| UniRef100\_A0A151FIE5 \| 99.9 \| 1.1e-29 \| 2e-35 \| 172.5 \| 138 \| (8, 151) \| 159 \| (55, 197) \| 211 \| Lysozyme \| Lysozyme; uniclust \| UniRef100\_UPI0018DC8388 \| 99.9 \| 1.2e-29 \| 2.1e-35 \| 181.1 \| 142 \| (2, 147) \| 159 \| (4, 149) \| 469 \| glycoside hydrolase family protein \| glycoside hydrolase family protein; uniclust \| UniRef100\_A0A1H9I7C8 \| 99.9 \| 1.5e-29 \| 2.8e-35 \| 174.0 \| 149 \| (1, 150) \| 159 \| (5, 159) \| 336 \| Lysozyme \| Lysozyme; uniclust \| UniRef100\_A0A060BVC9 \| 99.9 \| 1.7e-29 \| 3.2e-35 \| 156.6 \| 125 \| (20, 148) \| 159 \| (6, 130) \| 132 \| Lysozyme (Fragment) \| Lysozyme (Fragment); uniclust \| UniRef100\_A0A060CM67 \| 99.9 \| 1.8e-29 \| 3.4e-35 \| 168.7 \| 139 \| (8, 150) \| 159 \| (50, 196) \| 207 \| Lysozyme (Fragment) \| Lysozyme (Fragment); uniclust \| UniRef100\_A0A0F9U0N7 \| 99.9 \| 1.8e-29 \| 3.4e-35 \| 177.0 \| 149 \| (1, 149) \| 159 \| (142, 302) \| 304 \| lysozyme \| lysozyme; uniclust \| UniRef100\_UPI001F2D85D0 \| 99.9 \| 2.1e-29 \| 3.9e-35 \| 185.8 \| 145 \| (3, 150) \| 159 \| (344, 497) \| 666 \| D-Ala-D-Ala carboxypeptidase family metallohydrolase \| D-Ala-D-Ala carboxypeptidase family metallohydrolase; uniclust \| UniRef100\_A0A2P7EBL1 \| 99.9 \| 2.1e-29 \| 3.9e-35 \| 177.4 \| 146 \| (1, 152) \| 159 \| (1, 148) \| 414 \| Lysozyme \| Lysozyme; uniclust \| UniRef100\_A0A2Z4PEP3 \| 99.9 \| 2.3e-29 \| 4.3e-35 \| 153.8 \| 94 \| (1, 97) \| 159 \| (12, 109) \| 117 \| Lysozyme \| Lysozyme; uniclust \| UniRef100\_A0A940SYQ5 \| 99.9 \| 2.7e-29 \| 4.9e-35 \| 184.6 \| 149 \| (1, 153) \| 159 \| (6, 155) \| 637 \| Lysozyme \| Lysozyme; uniclust \| UniRef100\_A0A0Q9QM53 \| 99.9 \| 2.8e-29 \| 5.3e-35 \| 192.7 \| 142 \| (1, 148) \| 159 \| (92, 233) \| 612 \| Lysozyme \| Lysozyme; uniclust \| UniRef100\_A0A0F4VLV0 \| 99.9 \| 2.9e-29 \| 5.4e-35 \| 155.2 \| 137 \| (6, 145) \| 159 \| (4, 142) \| 143 \| Lysozyme \| Lysozyme; uniclust \| UniRef100\_A0A6B3UKC5 \| 99.9 \| 3.7e-29 \| 6.9e-35 \| 182.7 \| 145 \| (1, 149) \| 159 \| (5, 153) \| 442 \| Lysozyme (Fragment) \| Lysozyme (Fragment); uniclust \| UniRef100\_A0A257RRU2 \| 99.9 \| 3.9e-29 \| 7.2e-35 \| 169.5 \| 141 \| (4, 147) \| 159 \| (7, 157) \| 255 \| Lysozyme \| Lysozyme; uniclust \| UniRef100\_A0A0K2M126 \| 99.9 \| 4.1e-29 \| 7.6e-35 \| 173.3 \| 147 \| (6, 152) \| 159 \| (124, 279) \| 282 \| Lysozyme \| Lysozyme; uniclust \| UniRef100\_A0A933Z3Z4 \| 99.9 \| 4.4e-29 \| 8.1e-35 \| 166.0 \| 142 \| (3, 147) \| 159 \| (7, 157) \| 255 \| Lysozyme \| Lysozyme; uniclust \| UniRef100\_A0A146F1I5 \| 99.9 \| 4.6e-29 \| 8.5e-35 \| 184.9 \| 150 \| (1, 153) \| 159 \| (17, 170) \| 609 \| Lysozyme \| Lysozyme; uniclust \| UniRef100\_A0A1A9WZ64 \| 99.9 \| 4.7e-29 \| 8.7e-35 \| 168.3 \| 138 \| (9, 149) \| 159 \| (143, 280) \| 285 \| lysozyme \| lysozyme; uniclust \| UniRef100\_A0A7W0G9B2 \| 99.9 \| 4.8e-29 \| 8.8e-35 \| 178.8 \| 140 \| (3, 145) \| 159 \| (322, 469) \| 470 \| Lysozyme \| Lysozyme; uniclust \| UniRef100\_A0A318AUS2 \| 99.9 \| 5.6e-29 \| 1e-34 \| 172.4 \| 145 \| (1, 149) \| 159 \| (5, 153) \| 297 \| Lysozyme (Fragment) \| Lysozyme (Fragment); uniclust \| UniRef100\_A0A060C1D9 \| 99.9 \| 5.7e-29 \| 1e-34 \| 149.9 \| 111 \| (1, 111) \| 159 \| (2, 117) \| 118 \| Lysozyme (Fragment) \| Lysozyme (Fragment); uniclust \| UniRef100\_A0A2S9WZ50 \| 99.9 \| 6e-29 \| 1.1e-34 \| 148.1 \| 105 \| (1, 108) \| 159 \| (3, 107) \| 107 \| Lysozyme (Fragment) \| Lysozyme (Fragment); uniclust \| UniRef100\_A0A7W0Z8H7 \| 99.9 \| 7.4e-29 \| 1.4e-34 \| 169.0 \| 147 \| (1, 151) \| 159 \| (69, 223) \| 309 \| Lysozyme \| Lysozyme; uniclust \| UniRef100\_A0A1V0EBQ3 \| 99.9 \| 7.3e-29 \| 1.4e-34 \| 167.2 \| 148 \| (1, 151) \| 159 \| (13, 214) \| 224 \| Lysozyme \| Lysozyme; uniclust \| UniRef100\_A0A158BDW6 \| 99.9 \| 8.8e-29 \| 1.7e-34 \| 166.6 \| 142 \| (1, 149) \| 159 \| (67, 211) \| 212 \| Lysozyme \| Lysozyme; uniclust \| UniRef100\_A0A0P1FJR3 \| 99.9 \| 9.5e-29 \| 1.8e-34 \| 181.0 \| 145 \| (1, 151) \| 159 \| (41, 190) \| 355 \| Lysozyme \| Lysozyme; uniclust \| UniRef100\_A0A8T5QLT9 \| 99.9 \| 1e-28 \| 1.9e-34 \| 171.3 \| 145 \| (2, 150) \| 159 \| (167, 312) \| 317 \| lysozyme \| lysozyme; uniclust \| UniRef100\_A0A444LK34 \| 99.9 \| 1.1e-28 \| 2e-34 \| 177.8 \| 145 \| (1, 148) \| 159 \| (33, 185) \| 376 \| Lysozyme \| Lysozyme; uniclust \| UniRef100\_A0A7R9KK34 \| 99.9 \| 1.1e-28 \| 2e-34 \| 171.8 \| 140 \| (2, 145) \| 159 \| (194, 340) \| 341 \| lysozyme \| lysozyme; uniclust \| UniRef100\_A0A099KGY8 \| 99.9 \| 1.3e-28 \| 2.5e-34 \| 167.0 \| 129 \| (2, 133) \| 159 \| (24, 165) \| 189 \| Lysozyme \| Lysozyme; uniclust \| UniRef100\_A0A1D2MI83 \| 99.9 \| 1.7e-28 \| 3.1e-34 \| 172.7 \| 140 \| (4, 146) \| 159 \| (73, 224) \| 406 \| lysozyme (Fragment) \| lysozyme (Fragment); uniclust \| UniRef100\_A0A1B1ALJ1 \| 99.9 \| 1.7e-28 \| 3.1e-34 \| 178.4 \| 145 \| (1, 149) \| 159 \| (25, 173) \| 363 \| Lysozyme \| Lysozyme; uniclust \| UniRef100\_A0A976EUN2 \| 99.9 \| 1.8e-28 \| 3.3e-34 \| 179.3 \| 147 \| (2, 151) \| 159 \| (100, 254) \| 596 \| Lysozyme \| Lysozyme; uniclust \| UniRef100\_A0A4R2P729 \| 99.9 \| 1.9e-28 \| 3.5e-34 \| 166.6 \| 144 \| (1, 147) \| 159 \| (95, 244) \| 257 \| Lysozyme \| Lysozyme; uniclust \| UniRef100\_A0A0P0DLD7 \| 99.9 \| 1.9e-28 \| 3.6e-34 \| 184.7 \| 141 \| (1, 147) \| 159 \| (27, 167) \| 482 \| Lysozyme \| Lysozyme; uniclust \| UniRef100\_A0A0Q4N8X0 \| 99.9 \| 2.1e-28 \| 3.9e-34 \| 154.1 \| 140 \| (4, 146) \| 159 \| (7, 169) \| 171 \| Lysozyme \| Lysozyme; uniclust \| UniRef100\_UPI001F1FE970 \| 99.9 \| 2.4e-28 \| 4.3e-34 \| 179.6 \| 149 \| (3, 155) \| 159 \| (248, 399) \| 629 \| glycoside hydrolase family protein \| glycoside hydrolase family protein; uniclust \| UniRef100\_A0A0K8MB38 \| 99.9 \| 2.5e-28 \| 4.5e-34 \| 155.3 \| 138 \| (10, 150) \| 159 \| (1, 139) \| 183 \| Lysozyme \| Lysozyme; uniclust \| UniRef100\_UPI0013DE3C73 \| 99.9 \| 2.5e-28 \| 4.5e-34 \| 167.7 \| 147 \| (2, 151) \| 159 \| (130, 284) \| 328 \| glycoside hydrolase family protein \| glycoside hydrolase family protein; uniclust \| UniRef100\_A0A9D8X297 \| 99.9 \| 3e-28 \| 5.4e-34 \| 173.8 \| 135 \| (1, 145) \| 159 \| (319, 460) \| 460 \| N-acetylmuramoyl-L-alanine amidase \| N-acetylmuramoyl-L-alanine amidase; uniclust \| UniRef100\_A0A0F9UX49 \| 99.9 \| 3.3e-28 \| 6.1e-34 \| 184.6 \| 140 \| (2, 146) \| 159 \| (611, 765) \| 770 \| lysozyme \| lysozyme; uniclust \| UniRef100\_A0A2G1Y8D8 \| 99.9 \| 3.4e-28 \| 6.5e-34 \| 183.6 \| 145 \| (2, 150) \| 159 \| (11, 159) \| 466 \| Lysozyme \| Lysozyme; uniclust \| UniRef100\_A0A7J7BR61 \| 99.9 \| 3.8e-28 \| 7e-34 \| 165.9 \| 139 \| (2, 143) \| 159 \| (171, 313) \| 313 \| lysozyme \| lysozyme; uniclust \| UniRef100\_A0A968UEN8 \| 99.9 \| 4.2e-28 \| 7.7e-34 \| 146.0 \| 108 \| (3, 114) \| 159 \| (20, 127) \| 128 \| Lysozyme (Fragment) \| Lysozyme (Fragment); uniclust \| UniRef100\_A0A285RXB2 \| 99.9 \| 4.1e-28 \| 7.7e-34 \| 160.8 \| 145 \| (1, 150) \| 159 \| (4, 164) \| 202 \| Lysozyme \| Lysozyme; uniclust \| UniRef100\_A0A2T2RGS0 \| 99.9 \| 4.2e-28 \| 7.7e-34 \| 198.5 \| 142 \| (2, 147) \| 159 \| (4131, 4272) \| 4673 \| Lysozyme \| Lysozyme; uniclust \| UniRef100\_A0A023Q0S2 \| 99.9 \| 4e-28 \| 7.7e-34 \| 166.4 \| 122 \| (4, 132) \| 159 \| (23, 149) \| 185 \| Lysozyme \| Lysozyme; uniclust \| UniRef100\_A0A172Y625 \| 99.9 \| 4.9e-28 \| 9.2e-34 \| 167.9 \| 143 \| (1, 147) \| 159 \| (31, 173) \| 296 \| Lysozyme \| Lysozyme; uniclust \| UniRef100\_A0A090N7E4 \| 99.9 \| 5.1e-28 \| 9.5e-34 \| 154.8 \| 144 \| (1, 149) \| 159 \| (14, 169) \| 176 \| Lysozyme \| Lysozyme; uniclust \| UniRef100\_A0A0D6B817 \| 99.9 \| 5.2e-28 \| 9.8e-34 \| 165.4 \| 147 \| (2, 151) \| 159 \| (66, 218) \| 231 \| Lysozyme \| Lysozyme; uniclust \| UniRef100\_A0A847GG99 \| 99.9 \| 6.4e-28 \| 1.2e-33 \| 150.7 \| 119 \| (31, 149) \| 159 \| (1, 127) \| 141 \| Lysozyme \| Lysozyme; uniclust \| UniRef100\_A0A7C9TZG6 \| 99.9 \| 6.5e-28 \| 1.2e-33 \| 178.9 \| 148 \| (2, 152) \| 159 \| (79, 234) \| 696 \| Lysozyme \| Lysozyme; uniclust \| UniRef100\_A0A161SEG4 \| 99.9 \| 6.5e-28 \| 1.2e-33 \| 151.7 \| 137 \| (4, 145) \| 159 \| (5, 153) \| 155 \| Lysozyme \| Lysozyme; uniclust \| UniRef100\_UPI000A6534B1 \| 99.9 \| 6.6e-28 \| 1.2e-33 \| 169.3 \| 147 \| (2, 152) \| 159 \| (226, 381) \| 396 \| lysozyme \| lysozyme; uniclust \| UniRef100\_A0A0J7Y672 \| 99.9 \| 6.6e-28 \| 1.2e-33 \| 160.8 \| 143 \| (4, 148) \| 159 \| (9, 160) \| 234 \| Lysozyme \| Lysozyme; uniclust \| UniRef100\_A0A062GS02 \| 99.9 \| 7.2e-28 \| 1.4e-33 \| 181.6 \| 146 \| (1, 152) \| 159 \| (435, 595) \| 595 \| Lysozyme \| Lysozyme; uniclust \| UniRef100\_A0A0P7AX87 \| 99.9 \| 8.8e-28 \| 1.6e-33 \| 165.2 \| 146 \| (3, 151) \| 159 \| (133, 287) \| 292 \| lysozyme \| lysozyme; uniclust \| UniRef100\_A0A9D9SU57 \| 99.9 \| 8.9e-28 \| 1.6e-33 \| 160.9 \| 149 \| (1, 150) \| 159 \| (1, 165) \| 267 \| Lysozyme \| Lysozyme; uniclust \| UniRef100\_A0A7V8DKX2 \| 99.9 \| 8.8e-28 \| 1.6e-33 \| 169.8 \| 146 \| (1, 150) \| 159 \| (4, 153) \| 358 \| Lysozyme \| Lysozyme; uniclust \| UniRef100\_A0A8J7WEN4 \| 99.9 \| 9.1e-28 \| 1.7e-33 \| 181.6 \| 148 \| (2, 152) \| 159 \| (365, 531) \| 885 \| Lysozyme \| Lysozyme; uniclust \| UniRef100\_A0A7I8ZS27 \| 99.9 \| 9.7e-28 \| 1.8e-33 \| 158.9 \| 133 \| (13, 148) \| 159 \| (33, 165) \| 244 \| Lysozyme \| Lysozyme; uniclust \| UniRef100\_A0A3B0MJC2 \| 99.9 \| 1e-27 \| 1.9e-33 \| 157.2 \| 136 \| (9, 147) \| 159 \| (79, 214) \| 218 \| Lysozyme \| Lysozyme; uniclust \| UniRef100\_A0A3A8ZW54 \| 99.9 \| 1.1e-27 \| 2e-33 \| 173.0 \| 144 \| (2, 154) \| 159 \| (4, 150) \| 473 \| Lysozyme \| Lysozyme; uniclust \| UniRef100\_A0A3M0SWM4 \| 99.9 \| 1.2e-27 \| 2.2e-33 \| 160.2 \| 145 \| (3, 150) \| 159 \| (5, 155) \| 243 \| Lysozyme \| Lysozyme; uniclust \| UniRef100\_A0A0U3TF44 \| 99.9 \| 1.2e-27 \| 2.3e-33 \| 168.2 \| 149 \| (1, 149) \| 159 \| (120, 283) \| 285 \| Lysozyme \| Lysozyme; uniclust \| UniRef100\_A0A501W9K6 \| 99.9 \| 1.3e-27 \| 2.4e-33 \| 151.3 \| 141 \| (2, 145) \| 159 \| (9, 157) \| 159 \| Lysozyme \| Lysozyme; uniclust \| UniRef100\_UPI0014737934 \| 99.9 \| 1.3e-27 \| 2.4e-33 \| 157.0 \| 143 \| (1, 150) \| 159 \| (1, 143) \| 230 \| DUF5776 domain-containing protein \| DUF5776 domain-containing protein; uniclust \| UniRef100\_A0A1V5ITR2 \| 99.9 \| 1.4e-27 \| 2.5e-33 \| 152.5 \| 124 \| (24, 151) \| 159 \| (2, 125) \| 187 \| Lysozyme \| Lysozyme; uniclust \| UniRef100\_A0A0M1IT78 \| 99.9 \| 1.6e-27 \| 2.9e-33 \| 167.0 \| 147 \| (3, 152) \| 159 \| (4, 156) \| 338 \| Lysozyme \| Lysozyme; uniclust \| UniRef100\_UPI001C2C21C7 \| 99.9 \| 1.9e-27 \| 3.4e-33 \| 160.3 \| 146 \| (3, 151) \| 159 \| (5, 156) \| 278 \| glycoside hydrolase family protein \| glycoside hydrolase family protein; uniclust \| UniRef100\_A0A0F8Z6L9 \| 99.9 \| 2e-27 \| 3.7e-33 \| 163.7 \| 142 \| (2, 146) \| 159 \| (180, 331) \| 334 \| lysozyme (Fragment) \| lysozyme (Fragment); uniclust \| UniRef100\_A0A074MCX7 \| 99.9 \| 2e-27 \| 3.8e-33 \| 168.0 \| 146 \| (1, 150) \| 159 \| (146, 298) \| 312 \| Lysozyme \| Lysozyme; uniclust \| UniRef100\_A0A1M3KKZ9 \| 99.9 \| 2.2e-27 \| 4e-33 \| 154.0 \| 140 \| (4, 146) \| 159 \| (9, 158) \| 209 \| Lysozyme \| Lysozyme; uniclust \| UniRef100\_A0A4Q2QRI9 \| 99.9 \| 2.6e-27 \| 4.9e-33 \| 176.0 \| 144 \| (2, 151) \| 159 \| (375, 530) \| 531 \| Lysozyme (Fragment) \| Lysozyme (Fragment); uniclust \| UniRef100\_A0A2V2RWC3 \| 99.9 \| 2.7e-27 \| 5e-33 \| 179.1 \| 152 \| (1, 152) \| 159 \| (113, 275) \| 639 \| Lysozyme \| Lysozyme; uniclust \| UniRef100\_A0A1Y1J2A2 \| 99.9 \| 2.7e-27 \| 5.2e-33 \| 148.1 \| 118 \| (31, 149) \| 159 \| (3, 128) \| 129 \| Phage-related lysozyme \| Phage-related lysozyme; uniclust \| UniRef100\_A0A966W4D6 \| 99.9 \| 2.9e-27 \| 5.4e-33 \| 158.0 \| 139 \| (3, 144) \| 159 \| (118, 256) \| 260 \| Lysozyme \| Lysozyme; uniclust \| UniRef100\_A0A1G8AZ06 \| 99.9 \| 3e-27 \| 5.5e-33 \| 180.6 \| 146 \| (2, 150) \| 159 \| (230, 388) \| 831 \| Lysozyme \| Lysozyme; uniclust \| UniRef100\_A0A961ZTK2 \| 99.9 \| 3.1e-27 \| 5.6e-33 \| 158.1 \| 137 \| (9, 148) \| 159 \| (14, 157) \| 262 \| Lysozyme \| Lysozyme; uniclust \| UniRef100\_A0A843LAG6 \| 99.9 \| 4.4e-27 \| 8.1e-33 \| 180.3 \| 143 \| (2, 147) \| 159 \| (28, 180) \| 1047 \| lysozyme \| lysozyme; uniclust \| UniRef100\_A0A2W6IFF7 \| 99.9 \| 4.4e-27 \| 8.1e-33 \| 157.4 \| 134 \| (8, 146) \| 159 \| (73, 218) \| 245 \| Lysozyme \| Lysozyme; uniclust \| UniRef100\_UPI001565CE07 \| 99.9 \| 5.1e-27 \| 9.4e-33 \| 159.1 \| 148 \| (2, 149) \| 159 \| (84, 243) \| 245 \| lysozyme \| lysozyme; uniclust \| UniRef100\_A0A017HBD8 \| 99.9 \| 5e-27 \| 9.5e-33 \| 162.2 \| 139 \| (1, 151) \| 159 \| (8, 157) \| 243 \| Phage-related lysozyme \| Phage-related lysozyme; uniclust \| UniRef100\_A0A0A8K6Z9 \| 99.9 \| 5.1e-27 \| 9.5e-33 \| 170.4 \| 141 \| (1, 151) \| 159 \| (5, 151) \| 384 \| Lysozyme \| Lysozyme; uniclust \| UniRef100\_A0A3P2RCD3 \| 99.9 \| 5.2e-27 \| 9.6e-33 \| 157.6 \| 143 \| (1, 147) \| 159 \| (1, 145) \| 270 \| Lysozyme \| Lysozyme; uniclust \| UniRef100\_A0A0H3H043 \| 99.9 \| 5.3e-27 \| 1e-32 \| 154.5 \| 137 \| (2, 149) \| 159 \| (22, 159) \| 165 \| Lysozyme \| Lysozyme; uniclust \| UniRef100\_A0A346MWA4 \| 99.9 \| 5.5e-27 \| 1e-32 \| 156.6 \| 141 \| (7, 150) \| 159 \| (31, 184) \| 204 \| Lysozyme \| Lysozyme; uniclust \| UniRef100\_A0A0R3DQ56 \| 99.9 \| 6.4e-27 \| 1.2e-32 \| 155.5 \| 141 \| (8, 150) \| 159 \| (22, 162) \| 210 \| Lysozyme \| Lysozyme; uniclust \| UniRef100\_A0A085FNZ0 \| 99.9 \| 7e-27 \| 1.3e-32 \| 158.2 \| 141 \| (2, 150) \| 159 \| (73, 218) \| 239 \| Lysozyme \| Lysozyme; uniclust \| UniRef100\_A0A246G718 \| 99.9 \| 7.1e-27 \| 1.3e-32 \| 164.2 \| 145 \| (2, 152) \| 159 \| (177, 337) \| 337 \| Lysozyme \| Lysozyme; uniclust \| UniRef100\_A0A099FC71 \| 99.9 \| 7.4e-27 \| 1.4e-32 \| 163.9 \| 141 \| (1, 151) \| 159 \| (14, 165) \| 282 \| Lysozyme \| Lysozyme; uniclust \| UniRef100\_A0A0A2V9H7 \| 99.9 \| 8e-27 \| 1.5e-32 \| 163.4 \| 136 \| (9, 147) \| 159 \| (17, 152) \| 335 \| lysozyme \| lysozyme; uniclust \| UniRef100\_A0A078KPY1 \| 99.9 \| 9.4e-27 \| 1.7e-32 \| 156.6 \| 147 \| (1, 149) \| 159 \| (96, 245) \| 246 \| Lysozyme \| Lysozyme; uniclust \| UniRef100\_A0A6J5LRB2 \| 99.9 \| 1e-26 \| 1.9e-32 \| 147.4 \| 148 \| (1, 148) \| 159 \| (2, 156) \| 159 \| Lysozyme \| Lysozyme; uniclust \| UniRef100\_A0A3D5P936 \| 99.9 \| 1.1e-26 \| 2.1e-32 \| 158.5 \| 145 \| (2, 149) \| 159 \| (156, 303) \| 306 \| Lysozyme \| Lysozyme; uniclust \| UniRef100\_A0A6J7WL85 \| 99.9 \| 1.2e-26 \| 2.2e-32 \| 148.7 \| 143 \| (1, 149) \| 159 \| (1, 152) \| 191 \| Endolysin \| Endolysin; uniclust \| UniRef100\_UPI000AED14A8 \| 99.9 \| 1.2e-26 \| 2.3e-32 \| 167.9 \| 139 \| (6, 147) \| 159 \| (7, 169) \| 523 \| glycoside hydrolase family protein \| glycoside hydrolase family protein; uniclust \| UniRef100\_A0A143HSA8 \| 99.9 \| 1.3e-26 \| 2.4e-32 \| 147.8 \| 125 \| (21, 148) \| 159 \| (1, 127) \| 131 \| Lysozyme \| Lysozyme; uniclust \| UniRef100\_A0A840XX44 \| 99.9 \| 1.3e-26 \| 2.4e-32 \| 154.3 \| 147 \| (2, 149) \| 159 \| (5, 170) \| 237 \| Lysozyme \| Lysozyme; uniclust \| UniRef100\_A0A0K1S2G2 \| 99.9 \| 1.4e-26 \| 2.7e-32 \| 159.9 \| 144 \| (2, 146) \| 159 \| (86, 238) \| 253 \| Lysozyme \| Lysozyme; uniclust \| UniRef100\_A0A6G9YUS1 \| 99.9 \| 1.5e-26 \| 2.7e-32 \| 178.6 \| 147 \| (3, 152) \| 159 \| (594, 747) \| 1148 \| Lysozyme \| Lysozyme; uniclust \| UniRef100\_A0A1C6AN06 \| 99.9 \| 1.5e-26 \| 2.7e-32 \| 169.9 \| 144 \| (1, 148) \| 159 \| (38, 190) \| 495 \| Lysozyme \| Lysozyme; uniclust \| UniRef100\_A0A0F9C0T9 \| 99.9 \| 1.5e-26 \| 2.8e-32 \| 160.8 \| 132 \| (4, 138) \| 159 \| (45, 187) \| 206 \| lysozyme \| lysozyme; uniclust \| UniRef100\_A0A3B0JB27 \| 99.9 \| 1.6e-26 \| 3e-32 \| 139.7 \| 117 \| (32, 148) \| 159 \| (2, 118) \| 124 \| Lysozyme \| Lysozyme; uniclust \| UniRef100\_A0A317CEU3 \| 99.9 \| 1.7e-26 \| 3.1e-32 \| 156.9 \| 147 \| (1, 150) \| 159 \| (2, 168) \| 276 \| Lysozyme \| Lysozyme; uniclust \| UniRef100\_A0A0A1TCW0 \| 99.9 \| 1.7e-26 \| 3.1e-32 \| 166.6 \| 146 \| (4, 150) \| 159 \| (25, 180) \| 425 \| lysozyme \| lysozyme; uniclust \| UniRef100\_A0A349NKU6 \| 99.9 \| 1.7e-26 \| 3.2e-32 \| 136.6 \| 113 \| (1, 117) \| 159 \| (1, 113) \| 114 \| Lysozyme (Fragment) \| Lysozyme (Fragment); uniclust \| UniRef100\_A0A8J6UXX9 \| 99.9 \| 1.9e-26 \| 3.5e-32 \| 157.6 \| 143 \| (6, 151) \| 159 \| (92, 254) \| 269 \| Lysozyme \| Lysozyme; uniclust \| UniRef100\_A0A0F4VMI7 \| 99.9 \| 2e-26 \| 3.7e-32 \| 144.7 \| 95 \| (1, 99) \| 159 \| (23, 119) \| 128 \| Lysozyme \| Lysozyme; uniclust \| UniRef100\_A0A3S3SML5 \| 99.9 \| 2.2e-26 \| 4e-32 \| 146.9 \| 142 \| (2, 146) \| 159 \| (5, 146) \| 185 \| lysozyme \| lysozyme; uniclust \| UniRef100\_UPI001B3C9599 \| 99.9 \| 2.2e-26 \| 4.1e-32 \| 151.4 \| 143 \| (2, 147) \| 159 \| (7, 150) \| 230 \| lysozyme \| lysozyme; uniclust \| UniRef100\_A0A069IM11 \| 99.9 \| 2.2e-26 \| 4.2e-32 \| 151.2 \| 139 \| (2, 147) \| 159 \| (10, 148) \| 198 \| Lysozyme (Fragment) \| Lysozyme (Fragment); uniclust \| UniRef100\_A0A1Y1VV99 \| 99.9 \| 2.3e-26 \| 4.2e-32 \| 157.1 \| 144 \| (3, 150) \| 159 \| (115, 267) \| 273 \| lysozyme \| lysozyme; uniclust \| UniRef100\_A0A538JLK8 \| 99.9 \| 2.3e-26 \| 4.3e-32 \| 161.6 \| 138 \| (5, 145) \| 159 \| (187, 338) \| 339 \| Lysozyme \| Lysozyme; uniclust \| UniRef100\_A0A845WUP1 \| 99.9 \| 2.4e-26 \| 4.4e-32 \| 146.9 \| 135 \| (6, 146) \| 159 \| (50, 186) \| 186 \| Lysozyme \| Lysozyme; uniclust \| UniRef100\_A0A143XLM0 \| 99.9 \| 2.4e-26 \| 4.4e-32 \| 153.1 \| 138 \| (2, 149) \| 159 \| (3, 151) \| 229 \| Lysozyme \| Lysozyme; uniclust \| UniRef100\_A0A4Q2QUC8 \| 99.9 \| 2.8e-26 \| 5.3e-32 \| 173.1 \| 143 \| (2, 151) \| 159 \| (493, 644) \| 645 \| Lysozyme (Fragment) \| Lysozyme (Fragment); uniclust \| UniRef100\_A0A081NGH9 \| 99.9 \| 3.2e-26 \| 6.1e-32 \| 157.3 \| 147 \| (1, 150) \| 159 \| (27, 189) \| 206 \| Lysozyme \| Lysozyme; uniclust \| UniRef100\_A0A6B8KGM4 \| 99.9 \| 3.5e-26 \| 6.4e-32 \| 153.8 \| 144 \| (2, 148) \| 159 \| (3, 152) \| 271 \| Lysozyme \| Lysozyme; uniclust \| UniRef100\_UPI0011247FAC \| 99.9 \| 3.8e-26 \| 6.9e-32 \| 152.9 \| 143 \| (1, 150) \| 159 \| (1, 143) \| 260 \| glycoside hydrolase family protein \| glycoside hydrolase family protein; uniclust \| UniRef100\_A0A0Q5MFL0 \| 99.9 \| 3.8e-26 \| 7.1e-32 \| 167.2 \| 140 \| (2, 147) \| 159 \| (55, 194) \| 423 \| Lysozyme \| Lysozyme; uniclust \| UniRef100\_A0A0N4VYY9 \| 99.9 \| 4.5e-26 \| 8.3e-32 \| 155.1 \| 139 \| (6, 149) \| 159 \| (79, 220) \| 297 \| lysozyme \| lysozyme; uniclust \| UniRef100\_UPI0013DFCEBC \| 99.9 \| 4.8e-26 \| 8.8e-32 \| 158.3 \| 146 \| (1, 149) \| 159 \| (1, 157) \| 355 \| glycoside hydrolase family protein \| glycoside hydrolase family protein; uniclust \| UniRef100\_A0A6J4RNY5 \| 99.9 \| 5.3e-26 \| 9.7e-32 \| 175.3 \| 146 \| (2, 150) \| 159 \| (623, 778) \| 1117 \| Lysozyme \| Lysozyme; uniclust \| UniRef100\_A0A3D3IVC7 \| 99.9 \| 6.9e-26 \| 1.3e-31 \| 169.3 \| 145 \| (3, 150) \| 159 \| (13, 164) \| 735 \| Lysozyme \| Lysozyme; uniclust \| UniRef100\_UPI001184926A \| 99.9 \| 7.4e-26 \| 1.4e-31 \| 159.4 \| 148 \| (2, 150) \| 159 \| (4, 168) \| 398 \| glycoside hydrolase family protein \| glycoside hydrolase family protein; uniclust \| UniRef100\_A0A384UH61 \| 99.9 \| 7.6e-26 \| 1.4e-31 \| 142.0 \| 126 \| (18, 147) \| 159 \| (2, 127) \| 164 \| Endolysin \| Endolysin; uniclust \| UniRef100\_A0A6C2CEI5 \| 99.9 \| 7.7e-26 \| 1.4e-31 \| 166.6 \| 144 \| (1, 147) \| 159 \| (463, 620) \| 622 \| Lysozyme \| Lysozyme; uniclust \| UniRef100\_A0A0C1NIH4 \| 99.9 \| 7.9e-26 \| 1.5e-31 \| 159.1 \| 149 \| (3, 152) \| 159 \| (139, 297) \| 310 \| Lysozyme \| Lysozyme; uniclust \| UniRef100\_UPI001CC4FB84 \| 99.9 \| 8.3e-26 \| 1.5e-31 \| 165.3 \| 145 \| (2, 149) \| 159 \| (383, 538) \| 539 \| peptidoglycan-binding protein \| peptidoglycan-binding protein; uniclust \| UniRef100\_A0A4P7CYW0 \| 99.9 \| 8.5e-26 \| 1.6e-31 \| 152.2 \| 142 \| (7, 148) \| 159 \| (84, 253) \| 255 \| Lysozyme \| Lysozyme; uniclust \| UniRef100\_A3Z6C9 \| 99.9 \| 8.6e-26 \| 1.6e-31 \| 150.9 \| 144 \| (3, 147) \| 159 \| (5, 150) \| 256 \| Lysozyme \| Lysozyme; uniclust \| UniRef100\_A0A095FJ13 \| 99.9 \| 9.2e-26 \| 1.7e-31 \| 156.3 \| 148 \| (9, 156) \| 159 \| (84, 278) \| 280 \| Lysozyme \| Lysozyme; uniclust \| UniRef100\_A0A850BAY3 \| 99.9 \| 9.8e-26 \| 1.8e-31 \| 171.5 \| 145 \| (3, 150) \| 159 \| (740, 899) \| 919 \| Lysozyme (Fragment) \| Lysozyme (Fragment); uniclust \| UniRef100\_A0A3T0D2Y2 \| 99.9 \| 1.1e-25 \| 2e-31 \| 154.6 \| 148 \| (3, 153) \| 159 \| (109, 280) \| 292 \| Lysozyme \| Lysozyme; uniclust \| UniRef100\_R9JLH4 \| 99.9 \| 1.1e-25 \| 2e-31 \| 160.8 \| 136 \| (2, 146) \| 159 \| (4, 142) \| 451 \| Lysozyme \| Lysozyme; uniclust \| UniRef100\_A0A0A8TLB0 \| 99.9 \| 1.1e-25 \| 2.2e-31 \| 152.4 \| 123 \| (5, 134) \| 159 \| (32, 159) \| 186 \| Lysozyme \| Lysozyme; uniclust \| UniRef100\_A0A0F9BCT7 \| 99.9 \| 1.1e-25 \| 2.2e-31 \| 150.5 \| 126 \| (5, 133) \| 159 \| (10, 149) \| 172 \| lysozyme \| lysozyme; uniclust \| UniRef100\_A0A1G6ZEY4 \| 99.9 \| 1.4e-25 \| 2.5e-31 \| 143.2 \| 142 \| (3, 147) \| 159 \| (6, 152) \| 160 \| Lysozyme \| Lysozyme; uniclust \| UniRef100\_A0A562PQL7 \| 99.9 \| 1.4e-25 \| 2.6e-31 \| 174.0 \| 144 \| (2, 152) \| 159 \| (585, 742) \| 742 \| Lysozyme \| Lysozyme; uniclust \| UniRef100\_A0A009PDS4 \| 99.9 \| 1.4e-25 \| 2.6e-31 \| 151.0 \| 137 \| (8, 147) \| 159 \| (35, 184) \| 189 \| Lysozyme \| Lysozyme; uniclust \| UniRef100\_UPI002174E5E2 \| 99.9 \| 1.4e-25 \| 2.6e-31 \| 143.7 \| 135 \| (7, 144) \| 159 \| (15, 152) \| 189 \| lysozyme \| lysozyme; uniclust \| UniRef100\_A0A2G6ENY7 \| 99.9 \| 1.6e-25 \| 2.9e-31 \| 156.9 \| 141 \| (6, 149) \| 159 \| (6, 150) \| 377 \| Lysozyme \| Lysozyme; uniclust \| UniRef100\_A0A7G8Q4E4 \| 99.9 \| 1.9e-25 \| 3.5e-31 \| 152.1 \| 143 \| (3, 148) \| 159 \| (6, 168) \| 296 \| Lysozyme \| Lysozyme; uniclust \| UniRef100\_A0A535XI64 \| 99.9 \| 2e-25 \| 3.7e-31 \| 144.2 \| 137 \| (2, 148) \| 159 \| (55, 199) \| 200 \| Lysozyme \| Lysozyme; uniclust \| UniRef100\_D3F184 \| 99.9 \| 2.1e-25 \| 3.9e-31 \| 156.9 \| 141 \| (2, 146) \| 159 \| (230, 389) \| 391 \| Lysozyme \| Lysozyme; uniclust \| UniRef100\_A0A2D7IXU0 \| 99.9 \| 2.1e-25 \| 3.9e-31 \| 171.0 \| 144 \| (2, 149) \| 159 \| (6, 153) \| 686 \| Lysozyme \| Lysozyme; uniclust \| UniRef100\_A0A6G9YUS1 \| 99.9 \| 2.4e-25 \| 4.5e-31 \| 172.1 \| 145 \| (2, 149) \| 159 \| (394, 545) \| 1148 \| Lysozyme \| Lysozyme; uniclust \| UniRef100\_A0A2E3PQX9 \| 99.9 \| 2.6e-25 \| 4.7e-31 \| 152.2 \| 145 \| (1, 150) \| 159 \| (1, 161) \| 302 \| Lysozyme \| Lysozyme; uniclust \| UniRef100\_UPI0009B89992 \| 99.9 \| 2.8e-25 \| 5.2e-31 \| 147.7 \| 135 \| (11, 149) \| 159 \| (26, 160) \| 245 \| lysozyme \| lysozyme; uniclust \| UniRef100\_A0A0V8BQ91 \| 99.9 \| 2.8e-25 \| 5.3e-31 \| 159.5 \| 143 \| (1, 149) \| 159 \| (1, 148) \| 388 \| Lysozyme \| Lysozyme; uniclust \| UniRef100\_A0A127VHU8 \| 99.9 \| 3e-25 \| 5.5e-31 \| 148.2 \| 146 \| (2, 152) \| 159 \| (9, 160) \| 221 \| Lysozyme \| Lysozyme; uniclust \| UniRef100\_UPI0003638F79 \| 99.9 \| 3e-25 \| 5.5e-31 \| 144.3 \| 109 \| (3, 114) \| 159 \| (68, 178) \| 193 \| lysozyme \| lysozyme; uniclust \| UniRef100\_A0A832GZF3 \| 99.9 \| 3.1e-25 \| 5.7e-31 \| 153.5 \| 125 \| (30, 155) \| 159 \| (2, 129) \| 337 \| Lysozyme \| Lysozyme; uniclust \| UniRef100\_A0A940QJJ3 \| 99.9 \| 3.1e-25 \| 5.8e-31 \| 145.0 \| 147 \| (2, 152) \| 159 \| (39, 194) \| 217 \| Lysozyme (Fragment) \| Lysozyme (Fragment); uniclust \| UniRef100\_A0A011MKX9 \| 99.9 \| 3.6e-25 \| 6.7e-31 \| 162.6 \| 145 \| (3, 150) \| 159 \| (282, 434) \| 460 \| Lysozyme \| Lysozyme; uniclust \| UniRef100\_A0A068Q6B6 \| 99.9 \| 3.7e-25 \| 7e-31 \| 151.3 \| 132 \| (8, 144) \| 159 \| (31, 192) \| 218 \| Putative endolysin \| Putative endolysin; uniclust \| UniRef100\_K9XRH0 \| 99.9 \| 3.9e-25 \| 7.2e-31 \| 159.0 \| 114 \| (2, 119) \| 159 \| (67, 180) \| 481 \| Lysozyme \| Lysozyme; uniclust \| UniRef100\_A0A0D0KR74 \| 99.9 \| 3.9e-25 \| 7.3e-31 \| 154.1 \| 141 \| (9, 152) \| 159 \| (91, 241) \| 258 \| Lysozyme \| Lysozyme; uniclust \| UniRef100\_UPI0021BA9689 \| 99.9 \| 4.3e-25 \| 8e-31 \| 178.2 \| 148 \| (2, 152) \| 159 \| (1932, 2087) \| 2569 \| S8 family serine peptidase \| S8 family serine peptidase; uniclust \| UniRef100\_A0A147GQV9 \| 99.9 \| 4.4e-25 \| 8.2e-31 \| 142.9 \| 153 \| (1, 155) \| 159 \| (1, 176) \| 181 \| Lysozyme \| Lysozyme; uniclust \| UniRef100\_A0A3C1W3S6 \| 99.9 \| 4.7e-25 \| 8.7e-31 \| 140.5 \| 146 \| (1, 149) \| 159 \| (2, 161) \| 163 \| Lysozyme \| Lysozyme; uniclust \| UniRef100\_A0A1Q6RIL6 \| 99.9 \| 4.9e-25 \| 9.1e-31 \| 162.9 \| 151 \| (1, 152) \| 159 \| (31, 196) \| 536 \| Lysozyme \| Lysozyme; uniclust \| UniRef100\_UPI0008499688 \| 99.9 \| 5e-25 \| 9.1e-31 \| 146.2 \| 141 \| (7, 151) \| 159 \| (19, 162) \| 241 \| glycoside hydrolase family protein \| glycoside hydrolase family protein; uniclust \| UniRef100\_A0A261KVX8 \| 99.9 \| 5.2e-25 \| 9.6e-31 \| 129.2 \| 99 \| (1, 102) \| 159 \| (1, 103) \| 103 \| Lysozyme (Fragment) \| Lysozyme (Fragment); uniclust \| UniRef100\_A0A443J0A0 \| 99.8 \| 5.7e-25 \| 1.1e-30 \| 151.2 \| 142 \| (1, 152) \| 159 \| (11, 163) \| 269 \| Lysozyme \| Lysozyme; uniclust \| UniRef100\_A0A0E0UJC6 \| 99.8 \| 6.1e-25 \| 1.1e-30 \| 164.6 \| 146 \| (1, 151) \| 159 \| (39, 188) \| 627 \| Lysozyme \| Lysozyme; uniclust \| UniRef100\_UPI0009FEF08B \| 99.8 \| 6.3e-25 \| 1.2e-30 \| 151.1 \| 148 \| (2, 149) \| 159 \| (128, 287) \| 289 \| lysozyme \| lysozyme; uniclust \| UniRef100\_A0A1G5SFW3 \| 99.8 \| 6.3e-25 \| 1.2e-30 \| 147.7 \| 137 \| (4, 145) \| 159 \| (118, 265) \| 267 \| Lysozyme \| Lysozyme; uniclust \| UniRef100\_A0A081C6S4 \| 99.8 \| 6.4e-25 \| 1.2e-30 \| 155.7 \| 145 \| (1, 148) \| 159 \| (1, 159) \| 419 \| Lysozyme \| Lysozyme; uniclust \| UniRef100\_A0A368HDJ6 \| 99.8 \| 6.5e-25 \| 1.2e-30 \| 143.7 \| 114 \| (1, 118) \| 159 \| (5, 118) \| 218 \| Lysozyme \| Lysozyme; uniclust \| UniRef100\_UPI002107C0D7 \| 99.8 \| 7e-25 \| 1.3e-30 \| 130.0 \| 112 \| (33, 144) \| 159 \| (2, 113) \| 114 \| lysozyme \| lysozyme; uniclust \| UniRef100\_A0A359LSI1 \| 99.8 \| 7e-25 \| 1.3e-30 \| 146.8 \| 144 \| (2, 148) \| 159 \| (59, 211) \| 215 \| Lysozyme \| Lysozyme; uniclust \| UniRef100\_UPI000405D4F2 \| 99.8 \| 8.2e-25 \| 1.5e-30 \| 158.0 \| 131 \| (10, 144) \| 159 \| (367, 497) \| 498 \| glycoside hydrolase family protein \| glycoside hydrolase family protein; uniclust \| UniRef100\_A0A0L1KIF7 \| 99.8 \| 8.1e-25 \| 1.5e-30 \| 154.6 \| 145 \| (2, 150) \| 159 \| (147, 298) \| 312 \| Lysozyme \| Lysozyme; uniclust \| UniRef100\_A0A829Z1I7 \| 99.8 \| 8.4e-25 \| 1.6e-30 \| 134.2 \| 100 \| (53, 152) \| 159 \| (2, 101) \| 132 \| Lysozyme \| Lysozyme; uniclust \| UniRef100\_A0A5C7USV4 \| 99.8 \| 8.9e-25 \| 1.6e-30 \| 161.6 \| 148 \| (2, 152) \| 159 \| (44, 204) \| 637 \| Lysozyme \| Lysozyme; uniclust \| UniRef100\_UPI001BD066A5 \| 99.8 \| 9.6e-25 \| 1.8e-30 \| 143.4 \| 136 \| (8, 147) \| 159 \| (85, 222) \| 223 \| glycoside hydrolase family protein \| glycoside hydrolase family protein; uniclust \| UniRef100\_A0A7K1TL27 \| 99.8 \| 9.9e-25 \| 1.8e-30 \| 150.3 \| 141 \| (1, 144) \| 159 \| (171, 323) \| 324 \| Lysozyme \| Lysozyme; uniclust \| UniRef100\_UPI001F43C1FD \| 99.8 \| 1e-24 \| 1.9e-30 \| 139.3 \| 140 \| (7, 146) \| 159 \| (37, 180) \| 183 \| lysozyme \| lysozyme; uniclust \| UniRef100\_A0A1U7J078 \| 99.8 \| 1e-24 \| 1.9e-30 \| 163.4 \| 149 \| (3, 151) \| 159 \| (49, 206) \| 474 \| Lysozyme \| Lysozyme; uniclust \| UniRef100\_A0A380ZDW0 \| 99.8 \| 1.2e-24 \| 2.2e-30 \| 142.0 \| 107 \| (46, 152) \| 159 \| (3, 109) \| 175 \| Lysozyme \| Lysozyme; uniclust \| UniRef100\_A0A4S2MJL7 \| 99.8 \| 1.2e-24 \| 2.2e-30 \| 149.0 \| 138 \| (3, 148) \| 159 \| (123, 266) \| 271 \| lysozyme \| lysozyme; uniclust \| UniRef100\_A0A932PB96 \| 99.8 \| 1.3e-24 \| 2.4e-30 \| 151.3 \| 98 \| (52, 149) \| 159 \| (142, 252) \| 285 \| Lysozyme \| Lysozyme; uniclust \| UniRef100\_UPI001EEE7D44 \| 99.8 \| 1.4e-24 \| 2.5e-30 \| 141.3 \| 138 \| (9, 149) \| 159 \| (21, 159) \| 208 \| lysozyme \| lysozyme; uniclust \| UniRef100\_UPI000A7BED71 \| 99.8 \| 1.5e-24 \| 2.7e-30 \| 156.7 \| 140 \| (1, 148) \| 159 \| (1, 148) \| 440 \| glycoside hydrolase family protein \| glycoside hydrolase family protein; uniclust \| UniRef100\_A0A7W9YX61 \| 99.8 \| 1.5e-24 \| 2.7e-30 \| 147.6 \| 142 \| (1, 152) \| 159 \| (1, 147) \| 291 \| Lysozyme \| Lysozyme; uniclust \| UniRef100\_A0A7Y3RIZ1 \| 99.8 \| 1.5e-24 \| 2.7e-30 \| 160.9 \| 142 \| (6, 150) \| 159 \| (435, 576) \| 661 \| Lysozyme \| Lysozyme; uniclust \| UniRef100\_A0A1C7D850 \| 99.8 \| 1.7e-24 \| 3.1e-30 \| 150.1 \| 146 \| (1, 150) \| 159 \| (121, 273) \| 287 \| Lysozyme \| Lysozyme; uniclust \| UniRef100\_A0A177N7A2 \| 99.8 \| 1.8e-24 \| 3.4e-30 \| 139.9 \| 148 \| (2, 149) \| 159 \| (12, 173) \| 177 \| Lysozyme \| Lysozyme; uniclust \| UniRef100\_A0A252DI96 \| 99.8 \| 1.9e-24 \| 3.4e-30 \| 134.2 \| 105 \| (45, 149) \| 159 \| (7, 111) \| 132 \| Lysozyme \| Lysozyme; uniclust \| UniRef100\_A0A3A1Y9X2 \| 99.8 \| 1.9e-24 \| 3.5e-30 \| 138.1 \| 138 \| (5, 145) \| 159 \| (35, 176) \| 183 \| Lysozyme \| Lysozyme; uniclust \| UniRef100\_A0A022PFR1 \| 99.8 \| 1.9e-24 \| 3.5e-30 \| 149.2 \| 148 \| (2, 149) \| 159 \| (147, 306) \| 308 \| Lysozyme \| Lysozyme; uniclust \| UniRef100\_A0A192H186 \| 99.8 \| 1.9e-24 \| 3.5e-30 \| 160.9 \| 142 \| (1, 145) \| 159 \| (538, 687) \| 687 \| Lysozyme \| Lysozyme; uniclust \| UniRef100\_UPI0009EAD2D6 \| 99.8 \| 2e-24 \| 3.6e-30 \| 129.6 \| 105 \| (2, 109) \| 159 \| (3, 107) \| 122 \| lysozyme \| lysozyme; uniclust \| UniRef100\_UPI0007CCFC1A \| 99.8 \| 2.1e-24 \| 3.8e-30 \| 151.5 \| 146 \| (2, 154) \| 159 \| (216, 370) \| 378 \| lysozyme \| lysozyme; uniclust \| UniRef100\_A0A8H7CL31 \| 99.8 \| 2.2e-24 \| 4.1e-30 \| 146.2 \| 147 \| (3, 150) \| 159 \| (104, 262) \| 282 \| lysozyme \| lysozyme; uniclust \| UniRef100\_UPI00191D0E7B \| 99.8 \| 2.3e-24 \| 4.2e-30 \| 148.4 \| 154 \| (1, 155) \| 159 \| (89, 251) \| 320 \| lysozyme \| lysozyme; uniclust \| UniRef100\_A0A8A1MB86 \| 99.8 \| 2.4e-24 \| 4.5e-30 \| 144.6 \| 143 \| (3, 148) \| 159 \| (33, 182) \| 249 \| lysozyme \| lysozyme; uniclust \| UniRef100\_A0A1F3AD95 \| 99.8 \| 2.4e-24 \| 4.5e-30 \| 143.7 \| 145 \| (1, 149) \| 159 \| (1, 149) \| 250 \| Lysozyme \| Lysozyme; uniclust \| UniRef100\_A0A2T7Q7E6 \| 99.8 \| 2.7e-24 \| 4.9e-30 \| 138.9 \| 138 \| (2, 144) \| 159 \| (53, 196) \| 197 \| Lysozyme \| Lysozyme; uniclust \| UniRef100\_A0A357WVB9 \| 99.8 \| 2.8e-24 \| 5.2e-30 \| 144.9 \| 138 \| (2, 149) \| 159 \| (68, 210) \| 221 \| Lysozyme (Fragment) \| Lysozyme (Fragment); uniclust \| UniRef100\_J5TH48 \| 99.8 \| 2.8e-24 \| 5.2e-30 \| 161.6 \| 142 \| (3, 147) \| 159 \| (558, 713) \| 738 \| lysozyme \| lysozyme; uniclust \| UniRef100\_A0A8H9NPM1 \| 99.8 \| 3e-24 \| 5.6e-30 \| 146.1 \| 135 \| (9, 147) \| 159 \| (28, 162) \| 291 \| Lysozyme \| Lysozyme; uniclust \| UniRef100\_A0A242NX27 \| 99.8 \| 3e-24 \| 5.6e-30 \| 168.8 \| 145 \| (2, 152) \| 159 \| (977, 1132) \| 1132 \| Lysozyme \| Lysozyme; uniclust \| UniRef100\_A0A0T9R1Y9 \| 99.8 \| 3.1e-24 \| 5.8e-30 \| 168.8 \| 145 \| (2, 152) \| 159 \| (867, 1023) \| 1023 \| Lysozyme \| Lysozyme; uniclust \| UniRef100\_A0A0F5ALB6 \| 99.8 \| 3.1e-24 \| 5.8e-30 \| 141.9 \| 139 \| (6, 149) \| 159 \| (17, 162) \| 202 \| Lysozyme \| Lysozyme; uniclust \| UniRef100\_UPI001FAEC223 \| 99.8 \| 3.6e-24 \| 6.5e-30 \| 141.1 \| 136 \| (5, 145) \| 159 \| (76, 223) \| 227 \| lysozyme \| lysozyme; uniclust \| UniRef100\_UPI001DFEEB7A \| 99.8 \| 3.7e-24 \| 6.8e-30 \| 152.0 \| 145 \| (3, 150) \| 159 \| (87, 240) \| 418 \| uncharacterized protein \| uncharacterized protein; uniclust \| UniRef100\_A0A2S0U5Q1 \| 99.8 \| 4.1e-24 \| 7.5e-30 \| 161.1 \| 148 \| (2, 152) \| 159 \| (494, 648) \| 654 \| Lysozyme \| Lysozyme; uniclust \| UniRef100\_A0A7W8Q238 \| 99.8 \| 4.1e-24 \| 7.7e-30 \| 138.0 \| 132 \| (12, 151) \| 159 \| (18, 152) \| 153 \| Lysozyme \| Lysozyme; uniclust \| UniRef100\_A3WB87 \| 99.8 \| 4.2e-24 \| 7.8e-30 \| 149.7 \| 146 \| (1, 150) \| 159 \| (173, 325) \| 329 \| Lysozyme \| Lysozyme; uniclust \| UniRef100\_UPI001C0C1628 \| 99.8 \| 4.3e-24 \| 7.9e-30 \| 144.8 \| 136 \| (1, 151) \| 159 \| (1, 137) \| 264 \| lysozyme \| lysozyme; uniclust \| UniRef100\_UPI000D51B1FE \| 99.8 \| 4.7e-24 \| 8.6e-30 \| 146.3 \| 136 \| (2, 141) \| 159 \| (27, 162) \| 309 \| Rz-like lysis system protein LysB \| Rz-like lysis system protein LysB; uniclust \| UniRef100\_A0A1C4CVR3 \| 99.8 \| 4.6e-24 \| 8.7e-30 \| 147.2 \| 145 \| (1, 150) \| 159 \| (45, 232) \| 235 \| Lysozyme \| Lysozyme; uniclust \| UniRef100\_A0A661CVB8 \| 99.8 \| 4.8e-24 \| 8.8e-30 \| 151.1 \| 142 \| (3, 148) \| 159 \| (244, 405) \| 410 \| Lysozyme (Fragment) \| Lysozyme (Fragment); uniclust \| UniRef100\_A0A942F8L9 \| 99.8 \| 4.9e-24 \| 9.1e-30 \| 161.0 \| 148 \| (1, 151) \| 159 \| (222, 382) \| 807 \| Lysozyme \| Lysozyme; uniclust \| UniRef100\_A0A1X2GML7 \| 99.8 \| 5.5e-24 \| 1e-29 \| 147.0 \| 145 \| (5, 152) \| 159 \| (119, 272) \| 283 \| lysozyme \| lysozyme; uniclust \| UniRef100\_A0A071LUB0 \| 99.8 \| 5.4e-24 \| 1e-29 \| 148.2 \| 128 \| (5, 135) \| 159 \| (38, 181) \| 217 \| Lysozyme \| Lysozyme; uniclust \| UniRef100\_J0QIK5 \| 99.8 \| 6.1e-24 \| 1.1e-29 \| 155.6 \| 122 \| (31, 152) \| 159 \| (374, 499) \| 565 \| Lysozyme \| Lysozyme; uniclust \| UniRef100\_A0A031IWU6 \| 99.8 \| 5.9e-24 \| 1.1e-29 \| 147.5 \| 146 \| (4, 150) \| 159 \| (51, 214) \| 230 \| Prophage PSPPH06 lysozyme \| Prophage PSPPH06 lysozyme; uniclust \| UniRef100\_UPI000D8E9A34 \| 99.8 \| 6.1e-24 \| 1.1e-29 \| 143.0 \| 145 \| (2, 149) \| 159 \| (117, 263) \| 265 \| SH3 domain-containing protein \| SH3 domain-containing protein; uniclust \| UniRef100\_UPI001FF05F52 \| 99.8 \| 6.7e-24 \| 1.2e-29 \| 132.0 \| 102 \| (48, 149) \| 159 \| (2, 103) \| 139 \| lysozyme \| lysozyme; uniclust \| UniRef100\_A0A8E7EIT5 \| 99.8 \| 6.7e-24 \| 1.2e-29 \| 158.6 \| 148 \| (1, 151) \| 159 \| (159, 314) \| 710 \| lysozyme \| lysozyme; uniclust \| UniRef100\_UPI0009E45E4A \| 99.8 \| 7e-24 \| 1.3e-29 \| 153.7 \| 146 \| (1, 149) \| 159 \| (3, 154) \| 510 \| glycoside hydrolase family protein \| glycoside hydrolase family protein; uniclust \| UniRef100\_A0A8J2RB09 \| 99.8 \| 7.5e-24 \| 1.4e-29 \| 143.5 \| 143 \| (3, 148) \| 159 \| (120, 271) \| 280 \| lysozyme \| lysozyme; uniclust \| UniRef100\_A0A928PRK0 \| 99.8 \| 7.7e-24 \| 1.4e-29 \| 156.0 \| 142 \| (2, 146) \| 159 \| (453, 605) \| 605 \| Lysozyme \| Lysozyme; uniclust \| UniRef100\_H0EY40 \| 99.8 \| 8.1e-24 \| 1.5e-29 \| 143.9 \| 135 \| (3, 140) \| 159 \| (135, 278) \| 280 \| lysozyme \| lysozyme; uniclust \| UniRef100\_A0A6B8LZY5 \| 99.8 \| 8.4e-24 \| 1.6e-29 \| 128.0 \| 116 \| (31, 151) \| 159 \| (1, 120) \| 128 \| Lysozyme \| Lysozyme; uniclust \| UniRef100\_UPI00177A9A85 \| 99.8 \| 8.5e-24 \| 1.6e-29 \| 144.4 \| 144 \| (2, 151) \| 159 \| (141, 296) \| 297 \| lysozyme \| lysozyme; uniclust \| UniRef100\_UPI001FDFC480 \| 99.8 \| 1e-23 \| 1.9e-29 \| 153.0 \| 144 \| (3, 149) \| 159 \| (350, 510) \| 514 \| peptidoglycan DD-metalloendopeptidase family protein \| peptidoglycan DD-metalloendopeptidase family protein; uniclust \| UniRef100\_A0A5N6J8L9 \| 99.8 \| 1.1e-23 \| 2e-29 \| 139.4 \| 140 \| (3, 144) \| 159 \| (77, 231) \| 232 \| lysozyme \| lysozyme; uniclust \| UniRef100\_A0A0D8TUV2 \| 99.8 \| 1.1e-23 \| 2e-29 \| 138.5 \| 134 \| (13, 149) \| 159 \| (32, 165) \| 187 \| Lysozyme \| Lysozyme; uniclust \| UniRef100\_A0A0C4K550 \| 99.8 \| 1.1e-23 \| 2.1e-29 \| 144.3 \| 150 \| (1, 151) \| 159 \| (1, 155) \| 223 \| Lysozyme \| Lysozyme; uniclust \| UniRef100\_A0A066T2U5 \| 99.8 \| 1.3e-23 \| 2.4e-29 \| 142.1 \| 138 \| (9, 149) \| 159 \| (37, 185) \| 210 \| Lysozyme \| Lysozyme; uniclust \| UniRef100\_A0A2X1B257 \| 99.8 \| 1.3e-23 \| 2.4e-29 \| 144.2 \| 134 \| (10, 147) \| 159 \| (2, 135) \| 262 \| Phage-related lysozyme (Muraminidase) \| Phage-related lysozyme (Muraminidase); uniclust \| UniRef100\_A0A252EDH3 \| 99.8 \| 1.4e-23 \| 2.6e-29 \| 130.1 \| 100 \| (7, 109) \| 159 \| (9, 113) \| 121 \| Lysozyme (Fragment) \| Lysozyme (Fragment); uniclust \| UniRef100\_A0A1G4WUT2 \| 99.8 \| 1.4e-23 \| 2.6e-29 \| 149.7 \| 143 \| (2, 146) \| 159 \| (202, 345) \| 433 \| Lysozyme \| Lysozyme; uniclust \| UniRef100\_A0A2V2CCV5 \| 99.8 \| 1.4e-23 \| 2.7e-29 \| 155.5 \| 148 \| (3, 154) \| 159 \| (230, 386) \| 477 \| Lysozyme \| Lysozyme; uniclust \| UniRef100\_A0A7W4A0H5 \| 99.8 \| 1.5e-23 \| 2.7e-29 \| 136.4 \| 147 \| (3, 149) \| 159 \| (35, 193) \| 195 \| Lysozyme \| Lysozyme; uniclust \| UniRef100\_A0A1S8L4X7 \| 99.8 \| 1.5e-23 \| 2.8e-29 \| 150.1 \| 144 \| (3, 149) \| 159 \| (4, 154) \| 450 \| Lysozyme \| Lysozyme; uniclust \| UniRef100\_A0A919S1J4 \| 99.8 \| 1.6e-23 \| 2.9e-29 \| 146.9 \| 147 \| (1, 149) \| 159 \| (214, 364) \| 370 \| Lysozyme \| Lysozyme; uniclust \| UniRef100\_A0A1H1WLZ3 \| 99.8 \| 1.7e-23 \| 3.2e-29 \| 147.7 \| 148 \| (2, 152) \| 159 \| (231, 388) \| 395 \| Lysozyme \| Lysozyme; uniclust \| UniRef100\_A0A447J4W7 \| 99.8 \| 1.7e-23 \| 3.2e-29 \| 127.2 \| 100 \| (50, 149) \| 159 \| (3, 102) \| 109 \| Lysozyme \| Lysozyme; uniclust \| UniRef100\_A0A1Y3P0S9 \| 99.8 \| 2e-23 \| 3.7e-29 \| 135.3 \| 132 \| (2, 145) \| 159 \| (59, 199) \| 200 \| lysozyme \| lysozyme; uniclust \| UniRef100\_A0A976QVJ7 \| 99.8 \| 2e-23 \| 3.8e-29 \| 131.6 \| 139 \| (2, 146) \| 159 \| (21, 159) \| 166 \| Endolysin \| Endolysin; uniclust \| UniRef100\_A0A952LVR1 \| 99.8 \| 2.1e-23 \| 3.8e-29 \| 141.0 \| 145 \| (2, 150) \| 159 \| (114, 266) \| 272 \| Lysozyme \| Lysozyme; uniclust \| UniRef100\_A0A094EPJ7 \| 99.8 \| 2.1e-23 \| 3.8e-29 \| 147.6 \| 138 \| (5, 145) \| 159 \| (246, 392) \| 400 \| lysozyme \| lysozyme; uniclust \| UniRef100\_A0A0Q5GY79 \| 99.8 \| 2e-23 \| 3.8e-29 \| 150.4 \| 146 \| (1, 150) \| 159 \| (3, 165) \| 350 \| Peptidoglycan binding-like domain-containing protein \| Peptidoglycan binding-like domain-containing protein; uniclust \| UniRef100\_A0A430BCR4 \| 99.8 \| 2.1e-23 \| 3.9e-29 \| 144.4 \| 138 \| (8, 148) \| 159 \| (131, 269) \| 270 \| Lysozyme \| Lysozyme; pdb70 \| 6ET6\_A \| 99.9 \| 7.8e-34 \| 5.8e-38 \| 190.1 \| 141 \| (3, 146) \| 159 \| (52, 196) \| 196 \| Aminotransferase (E.C.2.6.1.-) \| 6ET6\_A Aminotransferase (E.C.2.6.1.-) muramidase, Lysozyme, Acinetobacter baumannii, ANTIMICROBIAL; pdb70 \| 3HDF\_A \| 99.9 \| 1.2e-33 \| 9.2e-38 \| 180.5 \| 138 \| (7, 147) \| 159 \| (2, 139) \| 140 \| Lysozyme (E.C.3.2.1.17) \| 3HDF\_A Lysozyme (E.C.3.2.1.17) lysozyme-like, Antimicrobial, Bacteriolytic enzyme, Glycosidase HET: NO3, GOL; pdb70 \| 6H9D\_A \| 99.9 \| 5.7e-33 \| 4.2e-37 \| 179.1 \| 140 \| (2, 145) \| 159 \| (5, 148) \| 149 \| Lysozyme (E.C.3.2.1.17) \| 6H9D\_A Lysozyme (E.C.3.2.1.17) lysozyme, peptidoglycan hydrolase, cell morphogenesis HET: GOL; pdb70 \| 1XJU\_A \| 99.9 \| 2.4e-32 \| 1.8e-36 \| 178.6 \| 145 \| (2, 150) \| 159 \| (1, 157) \| 163 \| Lysozyme (E.C.3.2.1.17) \| 1XJU\_A Lysozyme (E.C.3.2.1.17) secreted inactive conformation, HYDROLASE HET: SO4; pdb70 \| 3HDE\_B \| 99.9 \| 1.1e-31 \| 8.4e-36 \| 175.7 \| 134 \| (11, 147) \| 159 \| (31, 164) \| 165 \| Lysozyme (E.C.3.2.1.17) \| 3HDE\_B Lysozyme (E.C.3.2.1.17) lysozyme-like, Antimicrobial, Bacteriolytic enzyme, Glycosidase; pdb70 \| 3HDE\_D \| 99.9 \| 1.1e-31 \| 8.4e-36 \| 175.7 \| 134 \| (11, 147) \| 159 \| (31, 164) \| 165 \| Lysozyme (E.C.3.2.1.17) \| 3HDE\_D Lysozyme (E.C.3.2.1.17) lysozyme-like, Antimicrobial, Bacteriolytic enzyme, Glycosidase; pdb70 \| 1XJT\_A \| 99.9 \| 1.3e-31 \| 9.3e-36 \| 179.0 \| 144 \| (2, 149) \| 159 \| (29, 184) \| 191 \| Lysozyme (E.C.3.2.1.17) \| 1XJT\_A Lysozyme (E.C.3.2.1.17) open conformation, HYDROLASE HET: CIT; pdb70 \| 2ANX\_A \| 99.9 \| 4.4e-31 \| 3.3e-35 \| 169.5 \| 141 \| (1, 145) \| 159 \| (2, 146) \| 146 \| Lysozyme (E.C.3.2.1.17) \| 2ANX\_A Lysozyme (E.C.3.2.1.17) phage lysozyme, Sm-SAD, atomic resolution HET: SO4; pdb70 \| 4E97\_B \| 99.9 \| 1.2e-29 \| 8.7e-34 \| 169.1 \| 139 \| (2, 150) \| 159 \| (21, 183) \| 187 \| Lysozyme (E.C.3.2.1.17) \| 4E97\_B Lysozyme (E.C.3.2.1.17) Hydrolase, Alkylation with 2-mercaptoethanol HET: SO4, BME; pdb70 \| 6K69\_B \| 99.9 \| 5.9e-29 \| 4.4e-33 \| 167.8 \| 139 \| (2, 150) \| 159 \| (18, 202) \| 206 \| Protein A, 3LRH intrabody \| 6K69\_B Protein A, 3LRH intrabody antibody, protein design, STRUCTURAL PROTEIN; pdb70 \| 5JDT\_A \| 99.9 \| 7.1e-29 \| 5.3e-33 \| 162.1 \| 135 \| (6, 150) \| 159 \| (2, 160) \| 164 \| Endolysin (E.C.3.2.1.17) \| 5JDT\_A Endolysin (E.C.3.2.1.17) NITROXIDE, SPIN LABEL, T4 LYSOZYME HET: BME, MTN; pdb70 \| 4PK0\_A \| 99.9 \| 6.4e-28 \| 4.7e-32 \| 158.5 \| 137 \| (6, 152) \| 159 \| (2, 162) \| 171 \| TEICOPLANIN-A2-2, LYSOZYME (E.C.3.2.1.17) \| 4PK0\_A TEICOPLANIN-A2-2, LYSOZYME (E.C.3.2.1.17) site-selective catalyst, carrier protein approach HET: GHP, 3FG, 3MY, GCS, DAL, T55, NAG, OMY, MAN, MHS; pdb70 \| 4YX7\_C \| 99.9 \| 6.8e-28 \| 5e-32 \| 161.6 \| 139 \| (2, 150) \| 159 \| (31, 193) \| 197 \| Surface presentation of antigens protein \| 4YX7\_C Surface presentation of antigens protein Type III Secretion System, PROTEIN; pdb70 \| 5JEA\_K \| 99.9 \| 4.4e-27 \| 3.2e-31 \| 164.1 \| 138 \| (3, 150) \| 159 \| (114, 275) \| 279 \| Exosome complex component RRP45, Exosome \| 5JEA\_K Exosome complex component RRP45, Exosome exosome, Ski7, nuclease, RNA degradation HET: MPD; pdb70 \| 4EPI\_A \| 99.9 \| 6.7e-27 \| 4.9e-31 \| 165.7 \| 137 \| (3, 149) \| 159 \| (165, 325) \| 330 \| Pesticin, Lysozyme (E.C.3.2.1.17) \| 4EPI\_A Pesticin, Lysozyme (E.C.3.2.1.17) bacterial toxin, TOXIN, HYDROLASE; pdb70 \| 5EWX\_B \| 99.9 \| 7.2e-27 \| 5.3e-31 \| 159.1 \| 136 \| (6, 151) \| 159 \| (2, 224) \| 227 \| Endolysin (E.C.3.2.1.17),Immunoglobulin G-binding protein A \| 5EWX\_B Endolysin (E.C.3.2.1.17),Immunoglobulin G-binding protein A Fusion, EY-CBS, alpha helix, cross-linker HET: EYC; pdb70 \| 4XSJ\_A \| 99.9 \| 9.9e-27 \| 7.4e-31 \| 162.2 \| 136 \| (6, 151) \| 159 \| (2, 161) \| 262 \| lysozyme, Calcium uniporter protein, mitochondrial \| 4XSJ\_A lysozyme, Calcium uniporter protein, mitochondrial Membrane protein, Calcium channel, Mitochondria HET: SO4; pdb70 \| 4TN3\_B \| 99.9 \| 1.3e-26 \| 9.5e-31 \| 170.2 \| 139 \| (3, 151) \| 159 \| (235, 397) \| 400 \| Trim5alpha \| 4TN3\_B Trim5alpha Trim protein Coiled-coil scaffold retroviral; pdb70 \| 5BZ6\_A \| 99.8 \| 1.7e-26 \| 1.2e-30 \| 161.0 \| 136 \| (6, 151) \| 159 \| (2, 161) \| 262 \| Lysozyme,Calcium uniporter protein, mitochondrial (E.C.3.2.1.17) \| 5BZ6\_A Lysozyme,Calcium uniporter protein, mitochondrial (E.C.3.2.1.17) Membrane protein, Calcium channel, Mitochondria HET: SO4; pdb70 \| 5B2G\_G \| 99.8 \| 1.9e-26 \| 1.4e-30 \| 167.5 \| 140 \| (2, 151) \| 159 \| (4, 167) \| 364 \| Endolysin,Claudin-4 (E.C.3.2.1.17), Heat-labile enterotoxin B \| 5B2G\_G Endolysin,Claudin-4 (E.C.3.2.1.17), Heat-labile enterotoxin B Membrane protein, Complex, Cell-free protein HET: MSE; pdb70 \| 4TN3\_A \| 99.8 \| 2.1e-26 \| 1.6e-30 \| 169.0 \| 139 \| (3, 151) \| 159 \| (235, 397) \| 400 \| Trim5alpha \| 4TN3\_A Trim5alpha Trim protein Coiled-coil scaffold retroviral; pdb70 \| 4HTT\_B \| 99.8 \| 3.2e-26 \| 2.4e-30 \| 169.0 \| 140 \| (3, 152) \| 159 \| (243, 406) \| 418 \| Sec-independent protein translocase protein TatC \| 4HTT\_B Sec-independent protein translocase protein TatC Twin arginine translocase receptor, Membrane; pdb70 \| 2QB0\_D \| 99.8 \| 4.4e-26 \| 3.3e-30 \| 157.2 \| 136 \| (4, 149) \| 159 \| (79, 238) \| 241 \| E80-TELSAM domain, Lysozyme (E.C.3.2.1.17) \| 2QB0\_D E80-TELSAM domain, Lysozyme (E.C.3.2.1.17) Helical polymer, HYDROLASE REGULATOR; pdb70 \| 5B2G\_C \| 99.8 \| 5.3e-26 \| 4e-30 \| 165.1 \| 141 \| (2, 152) \| 159 \| (4, 168) \| 364 \| Endolysin,Claudin-4 (E.C.3.2.1.17), Heat-labile enterotoxin B \| 5B2G\_C Endolysin,Claudin-4 (E.C.3.2.1.17), Heat-labile enterotoxin B Membrane protein, Complex, Cell-free protein HET: MSE; pdb70 \| 5YQR\_A \| 99.8 \| 1.2e-25 \| 8.8e-30 \| 157.8 \| 134 \| (6, 149) \| 159 \| (3, 160) \| 276 \| T4 lysozyme/Membrane-anchored lipid-binding protein LAM6 \| 5YQR\_A T4 lysozyme/Membrane-anchored lipid-binding protein LAM6 ligand binding domain, sterol, lipid HET: 2PE; pdb70 \| 5VBA\_B \| 99.8 \| 1.4e-25 \| 1e-29 \| 166.6 \| 142 \| (2, 153) \| 159 \| (17, 182) \| 437 \| ESX-3 secretion-associated protein EspG3 \| 5VBA\_B ESX-3 secretion-associated protein EspG3 ESX-1, type VII secretion system HET: CL; pdb70 \| 5VBA\_A \| 99.8 \| 1.6e-25 \| 1.2e-29 \| 166.1 \| 141 \| (2, 152) \| 159 \| (17, 181) \| 437 \| ESX-3 secretion-associated protein EspG3 \| 5VBA\_A ESX-3 secretion-associated protein EspG3 ESX-1, type VII secretion system HET: CL; pdb70 \| 6A9E\_B \| 99.8 \| 5.4e-25 \| 4e-29 \| 161.2 \| 135 \| (6, 150) \| 159 \| (2, 160) \| 389 \| Endolysin,Autophagy-related protein 2 (E.C.3.2.1.17) \| 6A9E\_B Endolysin,Autophagy-related protein 2 (E.C.3.2.1.17) lipid transport HET: MSE; pdb70 \| 6A9J\_A \| 99.8 \| 5.4e-25 \| 4e-29 \| 161.2 \| 135 \| (6, 150) \| 159 \| (2, 160) \| 389 \| Endolysin,Autophagy-related protein 2 (E.C.3.2.1.17) \| 6A9J\_A Endolysin,Autophagy-related protein 2 (E.C.3.2.1.17) LIPID TRANSPORT HET: 9TL; pdb70 \| 4WTV\_B \| 99.8 \| 6.7e-25 \| 5e-29 \| 165.2 \| 138 \| (3, 150) \| 159 \| (74, 235) \| 514 \| Chimera protein of Phosphatidylinositol 4-kinase \| 4WTV\_B Chimera protein of Phosphatidylinositol 4-kinase lipid kinase, phosphatidyl inositol, transferase HET: ATP; pdb70 \| 4IAP\_A \| 99.8 \| 3.3e-24 \| 2.5e-28 \| 148.9 \| 137 \| (2, 148) \| 159 \| (16, 176) \| 260 \| Lysozyme, Oxysterol-binding protein homolog 3 \| 4IAP\_A Lysozyme, Oxysterol-binding protein homolog 3 PH domain, beta sandwitch, targeting HET: SO4; pdb70 \| 5I0N\_A \| 99.8 \| 4.2e-24 \| 3.1e-28 \| 161.8 \| 139 \| (2, 150) \| 159 \| (101, 263) \| 556 \| Phosphatidylinositol 4-kinase type 2-alpha,Lysozyme,Phosphatidylinositol 4-kinase \| 5I0N\_A Phosphatidylinositol 4-kinase type 2-alpha,Lysozyme,Phosphatidylinositol 4-kinase kinase, calcium, Transferase HET: ATP; pdb70 \| 5ZBQ\_A \| 99.8 \| 8.3e-24 \| 6.3e-28 \| 158.4 \| 138 \| (5, 150) \| 159 \| (365, 524) \| 525 \| Neuropeptide Y Y1 receptor in \| 5ZBQ\_A Neuropeptide Y Y1 receptor in G Protein-Coupled Receptor, Receptor Inhibitor HET: 9AO; pdb70 \| 6A73\_B \| 99.8 \| 9.2e-24 \| 6.8e-28 \| 149.9 \| 136 \| (6, 151) \| 159 \| (136, 295) \| 304 \| COP9 signalosome complex subunit 2,Endolysin \| 6A73\_B COP9 signalosome complex subunit 2,Endolysin subunit2 of COP9 Signalosome (CSN). HET: IHP, SO4; pdb70 \| 5NDD\_A \| 99.8 \| 1.4e-23 \| 1e-27 \| 160.6 \| 121 \| (3, 126) \| 159 \| (21, 165) \| 619 \| Lysozyme,Proteinase-activated receptor 2,Soluble cytochrome b562,Proteinase-activated \| 5NDD\_A Lysozyme,Proteinase-activated receptor 2,Soluble cytochrome b562,Proteinase-activated MEMBRANE PROTEIN, GPCR, 7TM HET: 8TZ; pdb70 \| 4EVX\_A \| 99.8 \| 2.4e-23 \| 1.8e-27 \| 126.3 \| 101 \| (2, 110) \| 159 \| (6, 106) \| 106 \| Putative phage endolysin \| 4EVX\_A Putative phage endolysin Structural Genomics, PSI-Biology, Program for; pdb70 \| 5IV5\_YA \| 99.8 \| 2.5e-23 \| 1.9e-27 \| 158.6 \| 137 \| (3, 149) \| 159 \| (172, 337) \| 575 \| Baseplate wedge protein gp6, Baseplate \| 5IV5\_YA Baseplate wedge protein gp6, Baseplate T4, baseplate-tail tube complex, pre-attachment; pdb70 \| 5IV5\_YB \| 99.8 \| 2.5e-23 \| 1.9e-27 \| 158.6 \| 137 \| (3, 149) \| 159 \| (172, 337) \| 575 \| Baseplate wedge protein gp6, Baseplate \| 5IV5\_YB Baseplate wedge protein gp6, Baseplate T4, baseplate-tail tube complex, pre-attachment; pdb70 \| 5IV5\_YC \| 99.8 \| 2.5e-23 \| 1.9e-27 \| 158.6 \| 137 \| (3, 149) \| 159 \| (172, 337) \| 575 \| Baseplate wedge protein gp6, Baseplate \| 5IV5\_YC Baseplate wedge protein gp6, Baseplate T4, baseplate-tail tube complex, pre-attachment; pdb70 \| 6MXT\_A \| 99.8 \| 2.5e-23 \| 1.9e-27 \| 154.1 \| 118 \| (6, 126) \| 159 \| (2, 143) \| 472 \| Fusion protein of Lysozyme and \| 6MXT\_A Fusion protein of Lysozyme and G protein-coupled receptor, adrenergic receptor HET: OLA, K5Y, OLC, P33, HTO, YCM; pdb70 \| 6AJH\_A \| 99.8 \| 2.9e-23 \| 2.1e-27 \| 166.2 \| 135 \| (6, 150) \| 159 \| (764, 922) \| 943 \| Drug exporters of the RND \| 6AJH\_A Drug exporters of the RND membrane protein, transporter, RND family HET: MHA, 9ZF, L6T; pdb70 \| 2Z6B\_A \| 99.7 \| 4.3e-23 \| 3.2e-27 \| 157.2 \| 134 \| (3, 146) \| 159 \| (172, 334) \| 584 \| Tail-associated lysozyme (E.C.3.2.1.17), Baseplate structural \| 2Z6B\_A Tail-associated lysozyme (E.C.3.2.1.17), Baseplate structural Protein containing metal complexes, Antimicrobial; pdb70 \| 1WTH\_A \| 99.7 \| 5.8e-23 \| 4.3e-27 \| 156.6 \| 135 \| (3, 147) \| 159 \| (172, 335) \| 584 \| Tail-associated lysozyme (E.C.3.2.1.17)/Baseplate structural protein \| 1WTH\_A Tail-associated lysozyme (E.C.3.2.1.17)/Baseplate structural protein Triple-stranded beta-helix, OB fold, pseudohexamer; pdb70 \| 6QAJ\_B \| 99.7 \| 6.9e-23 \| 5.2e-27 \| 155.0 \| 121 \| (1, 124) \| 159 \| (21, 163) \| 544 \| Endolysin,Transcription intermediary factor 1-beta (E.C.3.2.1.17,2.3.2.27) \| 6QAJ\_B Endolysin,Transcription intermediary factor 1-beta (E.C.3.2.1.17,2.3.2.27) Transcriptional repressor epigenetic silencing; pdb70 \| 6AJG\_A \| 99.7 \| 7.3e-23 \| 5.5e-27 \| 163.8 \| 134 \| (6, 149) \| 159 \| (764, 921) \| 943 \| Drug exporters of the RND \| 6AJG\_A Drug exporters of the RND membrane protein, transporter, RND family HET: 3RX, LMT, MHA; pdb70 \| 6AJF\_A \| 99.7 \| 9.3e-23 \| 6.9e-27 \| 163.2 \| 135 \| (6, 150) \| 159 \| (764, 922) \| 943 \| Drug exporters of the RND \| 6AJF\_A Drug exporters of the RND membrane protein, transporter, RND family HET: MHA, L6T; pdb70 \| 6BG5\_A \| 99.7 \| 1.2e-22 \| 9.1e-27 \| 148.1 \| 128 \| (2, 132) \| 159 \| (20, 171) \| 384 \| Endolysin, DCN1-like protein 1 chimera \| 6BG5\_A Endolysin, DCN1-like protein 1 chimera E3 Ligase, HYDROLASE, LIGASE-INHIBITOR complex HET: DQD; pdb70 \| 6BG3\_A \| 99.7 \| 1.3e-22 \| 9.3e-27 \| 148.0 \| 128 \| (2, 132) \| 159 \| (20, 171) \| 384 \| Endolysin, DCN1-like protein 1 chimera \| 6BG3\_A Endolysin, DCN1-like protein 1 chimera E3 Ligase, HYDROLASE, LIGASE-INHIBITOR complex HET: DOJ; pdb70 \| 6P5V\_A \| 99.7 \| 1.3e-22 \| 9.3e-27 \| 148.0 \| 128 \| (2, 132) \| 159 \| (20, 171) \| 384 \| Lysozyme,DCN1-like protein 1 fusion (E.C.3.2.1.17) \| 6P5V\_A Lysozyme,DCN1-like protein 1 fusion (E.C.3.2.1.17) E3 Ligase, LIGASE HET: MPD, O37; pdb70 \| 6IIH\_A \| 99.7 \| 1.5e-22 \| 1.1e-26 \| 151.7 \| 121 \| (3, 126) \| 159 \| (7, 151) \| 493 \| Probable septum site-determining protein MinC \| 6IIH\_A Probable septum site-determining protein MinC calcium binding protein mitochondrial; pdb70 \| 1K28\_A \| 99.7 \| 1.7e-22 \| 1.3e-26 \| 153.9 \| 138 \| (3, 150) \| 159 \| (172, 338) \| 584 \| TAIL-ASSOCIATED LYSOZYME, BASEPLATE STRUCTURAL PROTEIN \| 1K28\_A TAIL-ASSOCIATED LYSOZYME, BASEPLATE STRUCTURAL PROTEIN Triple-stranded beta-helix, OB fold, pseudohexamer; pdb70 \| 4LDE\_A \| 99.7 \| 2.3e-22 \| 1.7e-26 \| 148.8 \| 118 \| (1, 121) \| 159 \| (11, 150) \| 469 \| Lysozyme, Beta-2 adrenergic receptor (E.C.3.2.1.17) \| 4LDE\_A Lysozyme, Beta-2 adrenergic receptor (E.C.3.2.1.17) G protein coupled receptor, MEMBRANE HET: P0G, 1WV; pdb70 \| 4QKX\_A \| 99.7 \| 2.9e-22 \| 2.2e-26 \| 148.2 \| 119 \| (1, 122) \| 159 \| (11, 151) \| 469 \| Beta-2 adrenergic receptor (E.C.3.2.1.17), R9 \| 4QKX\_A Beta-2 adrenergic receptor (E.C.3.2.1.17), R9 7-transmembrane helices, Signal transduction, G HET: 35V; pdb70 \| 4ARM\_A \| 99.7 \| 8.3e-22 \| 6.1e-26 \| 140.6 \| 140 \| (5, 151) \| 159 \| (168, 355) \| 359 \| PESTICIN \| 4ARM\_A PESTICIN TOXIN, MURAMIDASE; pdb70 \| 5W0P\_C \| 99.7 \| 9.5e-22 \| 7.1e-26 \| 156.3 \| 130 \| (7, 139) \| 159 \| (2, 155) \| 906 \| T4 lysozyme-human rhodopsin-mouse visual arrestin \| 5W0P\_C T4 lysozyme-human rhodopsin-mouse visual arrestin rhodopsin, GPCR, arrestin, GRK, phosphorylation HET: TPO, SEP, NAG; pdb70 \| 4ARM\_B \| 99.7 \| 1e-21 \| 7.5e-26 \| 140.2 \| 142 \| (4, 152) \| 159 \| (167, 356) \| 359 \| PESTICIN \| 4ARM\_B PESTICIN TOXIN, MURAMIDASE; pdb70 \| 6FW2\_A \| 99.7 \| 1.2e-21 \| 8.6e-26 \| 145.9 \| 136 \| (4, 149) \| 159 \| (81, 240) \| 459 \| Mitochondrial amidoxime-reducing component 1,Endolysin,Mitochondrial amidoxime-reducing \| 6FW2\_A Mitochondrial amidoxime-reducing component 1,Endolysin,Mitochondrial amidoxime-reducing mARC, MOSC, molybdenum cofactor, Moco HET: B3P, MOO, MTE; pdb70 \| 4XES\_A \| 99.6 \| 9.8e-21 \| 7.4e-25 \| 141.7 \| 134 \| (9, 150) \| 159 \| (381, 536) \| 541 \| neurotensin receptor subtype 1, lysozyme \| 4XES\_A neurotensin receptor subtype 1, lysozyme membrane protein, G protein-coupled receptor HET: PEG, GOL, CIT, EPE; pdb70 \| 5I14\_A \| 99.5 \| 6e-19 \| 4.5e-23 \| 108.9 \| 99 \| (44, 149) \| 159 \| (10, 116) \| 121 \| mutated and truncated T4 lysozyme \| 5I14\_A mutated and truncated T4 lysozyme T4 lysozyme, hydrolase; pdb70 \| 4AQN\_A \| 99.4 \| 2.5e-17 \| 1.8e-21 \| 117.2 \| 132 \| (12, 150) \| 159 \| (175, 354) \| 357 \| PESTICIN \| 4AQN\_A PESTICIN TOXIN, BACTERIOCIN, COLICIN, THREE DOMAINS; pdb70 \| 2O7A\_A \| 99.4 \| 4e-17 \| 2.9e-21 \| 100.9 \| 93 \| (50, 150) \| 159 \| (1, 101) \| 124 \| Lysozyme (E.C.3.2.1.17) \| 2O7A\_A Lysozyme (E.C.3.2.1.17) Protein folding, protein stability, lysozyme; pdb70 \| 4N9N\_A \| 99.3 \| 1.7e-16 \| 1.3e-20 \| 117.2 \| 139 \| (7, 153) \| 159 \| (125, 285) \| 438 \| Sterol uptake control protein 2 \| 4N9N\_A Sterol uptake control protein 2 fungal nuclear receptor-like transcription factor HET: MSE; pdb70 \| 6D9M\_A \| 98.9 \| 1.4e-13 \| 1e-17 \| 97.2 \| 111 \| (6, 119) \| 159 \| (2, 134) \| 337 \| Endolysin,Response receiver sensor diguanylate cyclase \| 6D9M\_A Endolysin,Response receiver sensor diguanylate cyclase Cyclic dinucleotide, GGDEF, HYDROLASE HET: GMP, GTP; pdb70 \| 4Z9G\_A \| 98.3 \| 2.1e-10 \| 1.5e-14 \| 81.8 \| 114 \| (9, 125) \| 159 \| (122, 259) \| 443 \| Human Corticotropin-Releasing Factor Receptor Type \| 4Z9G\_A Human Corticotropin-Releasing Factor Receptor Type 7TM, GPCR, FAMILY B, SIGNALING HET: 1Q5, SO4, OLA; pdb70 \| 5EE7\_A \| 98.3 \| 2.9e-10 \| 2.1e-14 \| 81.2 \| 113 \| (10, 125) \| 159 \| (125, 261) \| 452 \| Glucagon receptor \| 5EE7\_A Glucagon receptor GPCR, signaling protein, 7TM HET: PE5, OLA, TLA, 5MV; pdb70 \| 5VEW\_A \| 97.9 \| 5e-09 \| 3.7e-13 \| 75.1 \| 113 \| (10, 125) \| 159 \| (129, 265) \| 455 \| Glucagon-like peptide 1 receptor,Endolysin chimera \| 5VEW\_A Glucagon-like peptide 1 receptor,Endolysin chimera GPCR, class B, 7TM domain HET: CSD, OLC, OLA, 97Y; pdb70 \| 5VEW\_B \| 97.9 \| 5e-09 \| 3.7e-13 \| 75.1 \| 113 \| (10, 125) \| 159 \| (129, 265) \| 455 \| Glucagon-like peptide 1 receptor,Endolysin chimera \| 5VEW\_B Glucagon-like peptide 1 receptor,Endolysin chimera GPCR, class B, 7TM domain HET: CSD, OLA, OLC, 97Y; pdb70 \| 5YQZ\_R \| 97.0 \| 1.1e-06 \| 8.2e-11 \| 65.2 \| 112 \| (11, 125) \| 159 \| (240, 375) \| 575 \| Glucagon receptor,Endolysin (E.C.3.2.1.17), Glucagon analogue \| 5YQZ\_R Glucagon receptor,Endolysin (E.C.3.2.1.17), Glucagon analogue Human GCGR receptor, Class B HET: OLC, NAG; pdb70 \| 5XEZ\_A \| 96.8 \| 4.1e-06 \| 3e-10 \| 62.4 \| 110 \| (12, 124) \| 159 \| (240, 373) \| 574 \| Glucagon receptor,Endolysin (E.C.3.2.1.17), Antibody, mAb1 \| 5XEZ\_A Glucagon receptor,Endolysin (E.C.3.2.1.17), Antibody, mAb1 Human GCGR receptor, Class B HET: 97V, NAG; pdb70 \| 5XF1\_A \| 96.8 \| 4.1e-06 \| 3e-10 \| 62.4 \| 110 \| (12, 124) \| 159 \| (240, 373) \| 574 \| Glucagon receptor,Endolysin (E.C.3.2.1.17), Antibody mAb1 \| 5XF1\_A Glucagon receptor,Endolysin (E.C.3.2.1.17), Antibody mAb1 Human GCGR receptor, Class B HET: NAG, 97V |
| Top keywords  (threshold 1.00e-03 (evalue)) | **Lysozyme, Endolysin, E, C, Fragment, hydrolase, receptor, glycoside, Membrane, complex** |
| Output files | ../../similar\_sequences/64\_FANPEZAQ\_CDS\_0064\_merged.svg ../../similar\_sequences/64\_FANPEZAQ\_CDS\_0064\_pdb70.a3m ../../similar\_sequences/64\_FANPEZAQ\_CDS\_0064\_pdb70.hhr ../../similar\_sequences/64\_FANPEZAQ\_CDS\_0064\_uniclust.a3m ../../similar\_sequences/64\_FANPEZAQ\_CDS\_0064\_uniclust.hhr |

#### Structure prediction (AlphaFold)^2^

|  |  |
| --- | --- |
| Stats | xml version="1.0" encoding="utf-8" standalone="no"?       2024-09-02T21:09:59.129907 image/svg+xml   Matplotlib v3.7.2, https://matplotlib.org/ |
| Predicted structure | **NGL Viewer Controls:**  - Center: *Left-Click* - Rotate: *Left-Click + Drag* - Translate: *Right-Click + Drag* - Zoom: *Shift + Left-Click + Drag* |
| Output files | ../../predicted\_structures/64\_FANPEZAQ\_CDS\_0064/features.pkl ../../predicted\_structures/64\_FANPEZAQ\_CDS\_0064/ranked\_0.pdb ../../predicted\_structures/64\_FANPEZAQ\_CDS\_0064/ranked\_0\_plots.svg ../../predicted\_structures/64\_FANPEZAQ\_CDS\_0064/result\_model\_1\_ptm\_pred\_0.pkl |

#### Structure similarity search results (Foldseek)^3^

|  |  |
| --- | --- |
| Structure databases searched | Pdb, Afdb-proteome, Afdb-uniprot50 |
| Results, scheme(s)  (Top layers only, threshold 1.00e-02 (evalue)) | xml version="1.0" encoding="utf-8" standalone="no"?       2024-09-02T21:11:33.967636 image/svg+xml   Matplotlib v3.7.2, https://matplotlib.org/ |
| Results, table  (threshold 1.00e-02 (evalue)) | db \| id \| prob \| evalue \| bits \| fident \| alnlen \| mismatch \| gapopen \| qstart \| qend \| tstart \| tend \| name \| description; pdb \| 6ET6\_A \| 1.0 \| 1.504e-12 \| 515 \| 0.486 \| 150 \| 70 \| 3 \| 1 \| 146 \| 3 \| 149 \| Lysozyme \| Lysozyme; pdb \| 8APP\_D \| 1.0 \| 1.633e-11 \| 468 \| 0.443 \| 151 \| 73 \| 6 \| 1 \| 143 \| 38 \| 185 \| Endolysin \| Endolysin; pdb \| 3HDE\_A \| 1.0 \| 1.868e-10 \| 444 \| 0.351 \| 131 \| 82 \| 2 \| 14 \| 144 \| 34 \| 161 \| Lysozyme \| Lysozyme; pdb \| 7M5I\_B \| 1.0 \| 8.002e-11 \| 443 \| 0.407 \| 152 \| 80 \| 5 \| 1 \| 145 \| 5 \| 153 \| Endolysin \| Endolysin; pdb \| 8HP8\_A \| 1.0 \| 1.868e-10 \| 436 \| 0.329 \| 158 \| 104 \| 2 \| 1 \| 157 \| 3 \| 159 \| Endolysin mtEC340M \| Endolysin mtEC340M; pdb \| 6H9D\_B \| 1.0 \| 2.309e-10 \| 435 \| 0.393 \| 150 \| 83 \| 5 \| 1 \| 146 \| 3 \| 148 \| Lysozyme \| Lysozyme; pdb \| 2ANV\_B \| 1.0 \| 4.136e-10 \| 425 \| 0.429 \| 149 \| 75 \| 6 \| 1 \| 144 \| 2 \| 145 \| Lysozyme \| Lysozyme; pdb \| 4ZPU\_A \| 1.0 \| 7.811e-10 \| 408 \| 0.319 \| 144 \| 90 \| 3 \| 6 \| 144 \| 21 \| 161 \| Lysozyme RrrD \| Lysozyme RrrD; pdb \| 8B2S\_B \| 1.0 \| 2.027e-09 \| 372 \| 0.388 \| 152 \| 81 \| 4 \| 1 \| 143 \| 79 \| 227 \| GH24 family muramidase \| GH24 family muramidase; pdb \| 1XJT\_A \| 1.0 \| 1.931e-07 \| 310 \| 0.23 \| 156 \| 104 \| 5 \| 1 \| 144 \| 20 \| 171 \| Lysozyme \| Lysozyme; pdb \| 3HDE\_D \| 1.0 \| 1.264e-07 \| 301 \| 0.282 \| 145 \| 89 \| 4 \| 6 \| 145 \| 21 \| 155 \| Lysozyme \| Lysozyme; pdb \| 3HDF\_B \| 1.0 \| 9.743e-06 \| 215 \| 0.338 \| 133 \| 79 \| 4 \| 15 \| 143 \| 4 \| 131 \| Lysozyme \| Lysozyme; pdb \| 4EVX\_A \| 1.0 \| 0.0007918 \| 167 \| 0.25 \| 92 \| 66 \| 2 \| 2 \| 93 \| 1 \| 89 \| Putative phage endolysin \| Putative phage endolysin; pdb \| 7RGR\_A \| 1.0 \| 0.0003976 \| 164 \| 0.186 \| 177 \| 108 \| 9 \| 1 \| 153 \| 1 \| 165 \| Artificial protein L056 \| Artificial protein L056; pdb \| 201L\_B \| 1.0 \| 0.0005465 \| 159 \| 0.203 \| 172 \| 104 \| 9 \| 6 \| 153 \| 2 \| 164 \| T4 LYSOZYME \| T4 LYSOZYME; pdb \| 1XJU\_A \| 1.0 \| 0.0002893 \| 158 \| 0.177 \| 163 \| 100 \| 6 \| 2 \| 144 \| 1 \| 149 \| Lysozyme \| Lysozyme; pdb \| 3FI5\_C \| 1.0 \| 0.0002105 \| 158 \| 0.205 \| 170 \| 99 \| 9 \| 6 \| 150 \| 2 \| 160 \| Lysozyme \| Lysozyme; pdb \| 137L\_B \| 1.0 \| 0.0003217 \| 157 \| 0.207 \| 159 \| 96 \| 9 \| 6 \| 140 \| 2 \| 154 \| T4 LYSOZYME \| T4 LYSOZYME; pdb \| 1L65\_A \| 1.0 \| 0.0005465 \| 156 \| 0.207 \| 169 \| 100 \| 11 \| 6 \| 150 \| 2 \| 160 \| LYSOZYME \| LYSOZYME; pdb \| 7Z36\_A \| 1.0 \| 0.0005183 \| 156 \| 0.192 \| 177 \| 110 \| 9 \| 1 \| 153 \| 3 \| 170 \| Endolysin,Transcription intermediary factor 1-beta,Isoform 2 of Transcription intermediary factor 1-beta \| Endolysin,Transcription intermediary factor 1-beta,Isoform 2 of Transcription intermediary factor 1-beta; pdb \| 147L\_A \| 1.0 \| 0.0003051 \| 153 \| 0.197 \| 172 \| 98 \| 10 \| 6 \| 150 \| 2 \| 160 \| T4 LYSOZYME \| T4 LYSOZYME; pdb \| 2HUM\_B \| 1.0 \| 0.0003577 \| 153 \| 0.215 \| 172 \| 95 \| 10 \| 6 \| 150 \| 2 \| 160 \| Lysozyme \| Lysozyme; pdb \| 3OE8\_A \| 1.0 \| 0.0003392 \| 152 \| 0.205 \| 180 \| 105 \| 11 \| 4 \| 157 \| 198 \| 365 \| C-X-C chemokine receptor type 4, Lysozyme Chimera \| C-X-C chemokine receptor type 4, Lysozyme Chimera; pdb \| 3EML\_A \| 1.0 \| 0.0004915 \| 152 \| 0.201 \| 179 \| 103 \| 10 \| 4 \| 155 \| 198 \| 363 \| Human Adenosine A2A receptor/T4 lysozyme chimera \| Human Adenosine A2A receptor/T4 lysozyme chimera; pdb \| 216L\_B \| 1.0 \| 0.0003392 \| 150 \| 0.204 \| 166 \| 104 \| 9 \| 6 \| 146 \| 2 \| 164 \| T4 LYSOZYME \| T4 LYSOZYME; pdb \| 168L\_A \| 1.0 \| 0.0004915 \| 150 \| 0.205 \| 170 \| 100 \| 8 \| 6 \| 150 \| 2 \| 161 \| T4 LYSOZYME \| T4 LYSOZYME; pdb \| 3SBA\_D \| 1.0 \| 0.00121 \| 149 \| 0.196 \| 173 \| 104 \| 10 \| 4 \| 152 \| 1 \| 162 \| Lysozyme \| Lysozyme; pdb \| 5JGR\_A \| 1.0 \| 0.0004193 \| 149 \| 0.194 \| 170 \| 101 \| 8 \| 6 \| 150 \| 2 \| 160 \| Endolysin \| Endolysin; pdb \| 5VBA\_A \| 1.0 \| 0.0007122 \| 149 \| 0.174 \| 178 \| 105 \| 8 \| 1 \| 150 \| 3 \| 166 \| Lysozyme, ESX-1 secretion-associated protein EspG1 chimera \| Lysozyme, ESX-1 secretion-associated protein EspG1 chimera; pdb \| 5KI1\_A \| 1.0 \| 0.0004915 \| 148 \| 0.191 \| 172 \| 99 \| 9 \| 6 \| 150 \| 2 \| 160 \| Endolysin \| Endolysin; pdb \| 201L\_A \| 1.0 \| 0.0003976 \| 148 \| 0.201 \| 174 \| 97 \| 10 \| 6 \| 150 \| 2 \| 162 \| T4 LYSOZYME \| T4 LYSOZYME; pdb \| 8F11\_A \| 1.0 \| 0.0004421 \| 148 \| 0.194 \| 170 \| 101 \| 8 \| 6 \| 150 \| 2 \| 160 \| Endolysin \| Endolysin; pdb \| 3FAD\_A \| 1.0 \| 0.0006075 \| 148 \| 0.208 \| 173 \| 100 \| 10 \| 6 \| 153 \| 2 \| 162 \| Lysozyme \| Lysozyme; pdb \| 5JGN\_A \| 1.0 \| 0.0004915 \| 148 \| 0.194 \| 170 \| 101 \| 8 \| 6 \| 150 \| 2 \| 160 \| Endolysin \| Endolysin; pdb \| 6U0C\_A \| 1.0 \| 0.0004661 \| 148 \| 0.203 \| 172 \| 97 \| 10 \| 6 \| 150 \| 2 \| 160 \| Endolysin \| Endolysin; pdb \| 6V51\_A \| 1.0 \| 0.0005183 \| 148 \| 0.203 \| 172 \| 97 \| 9 \| 6 \| 150 \| 2 \| 160 \| Endolysin \| Endolysin; pdb \| 4PJZ\_A \| 1.0 \| 0.0007918 \| 148 \| 0.209 \| 177 \| 99 \| 11 \| 6 \| 155 \| 2 \| 164 \| Lysozyme \| Lysozyme; pdb \| 5VBA\_B \| 1.0 \| 0.0007122 \| 148 \| 0.191 \| 178 \| 104 \| 10 \| 1 \| 151 \| 6 \| 170 \| Lysozyme, ESX-1 secretion-associated protein EspG1 chimera \| Lysozyme, ESX-1 secretion-associated protein EspG1 chimera; pdb \| 1QTB\_A \| 1.0 \| 0.00121 \| 147 \| 0.207 \| 169 \| 100 \| 10 \| 6 \| 150 \| 2 \| 160 \| LYSOZYME \| LYSOZYME; pdb \| 169L\_B \| 1.0 \| 0.0007918 \| 146 \| 0.205 \| 170 \| 99 \| 9 \| 7 \| 151 \| 3 \| 161 \| T4 LYSOZYME \| T4 LYSOZYME; pdb \| 1B6I\_A \| 1.0 \| 0.001088 \| 146 \| 0.191 \| 167 \| 105 \| 8 \| 6 \| 150 \| 2 \| 160 \| PROTEIN (LYSOZYME) \| PROTEIN (LYSOZYME); pdb \| 1L67\_A \| 1.0 \| 0.001032 \| 146 \| 0.203 \| 162 \| 95 \| 9 \| 6 \| 143 \| 2 \| 153 \| LYSOZYME \| LYSOZYME; pdb \| 7XEA\_A \| 1.0 \| 0.0005762 \| 146 \| 0.2 \| 170 \| 100 \| 9 \| 6 \| 150 \| 2 \| 160 \| Endolysin \| Endolysin; pdb \| 5JGU\_A \| 1.0 \| 0.0006075 \| 146 \| 0.194 \| 170 \| 101 \| 8 \| 6 \| 150 \| 2 \| 160 \| Endolysin \| Endolysin; pdb \| 5JGX\_A \| 1.0 \| 0.0007122 \| 146 \| 0.196 \| 173 \| 102 \| 9 \| 6 \| 153 \| 2 \| 162 \| Endolysin \| Endolysin; pdb \| 2NTG\_A \| 1.0 \| 0.0006075 \| 146 \| 0.194 \| 170 \| 101 \| 8 \| 6 \| 150 \| 2 \| 160 \| Lysozyme \| Lysozyme; pdb \| 3SB6\_A \| 1.0 \| 0.00121 \| 146 \| 0.191 \| 167 \| 105 \| 8 \| 6 \| 150 \| 4 \| 162 \| Lysozyme \| Lysozyme; pdb \| 197L\_A \| 1.0 \| 0.001147 \| 145 \| 0.167 \| 167 \| 109 \| 8 \| 6 \| 150 \| 2 \| 160 \| LYSOZYME \| LYSOZYME; pdb \| 3F9L\_A \| 1.0 \| 0.0005465 \| 145 \| 0.205 \| 170 \| 99 \| 9 \| 6 \| 150 \| 2 \| 160 \| Lysozyme \| Lysozyme; pdb \| 3CDO\_C \| 1.0 \| 0.0006075 \| 145 \| 0.2 \| 170 \| 100 \| 9 \| 6 \| 150 \| 2 \| 160 \| Lysozyme \| Lysozyme; pdb \| 3F8V\_A \| 1.0 \| 0.0007918 \| 145 \| 0.202 \| 173 \| 101 \| 10 \| 6 \| 153 \| 2 \| 162 \| Lysozyme \| Lysozyme; pdb \| 1G0P\_A \| 1.0 \| 0.001495 \| 145 \| 0.195 \| 169 \| 102 \| 9 \| 6 \| 150 \| 2 \| 160 \| PROTEIN (LYSOZYME) \| PROTEIN (LYSOZYME); pdb \| 1L18\_A \| 1.0 \| 0.000751 \| 145 \| 0.2 \| 175 \| 99 \| 11 \| 6 \| 153 \| 2 \| 162 \| T4 LYSOZYME \| T4 LYSOZYME; pdb \| 1L21\_A \| 1.0 \| 0.001663 \| 145 \| 0.205 \| 170 \| 104 \| 10 \| 6 \| 153 \| 2 \| 162 \| T4 LYSOZYME \| T4 LYSOZYME; pdb \| 1P36\_A \| 1.0 \| 0.001495 \| 145 \| 0.191 \| 167 \| 105 \| 7 \| 6 \| 150 \| 2 \| 160 \| LYSOZYME \| LYSOZYME; pdb \| 4WTV\_A \| 1.0 \| 0.0007122 \| 145 \| 0.201 \| 174 \| 99 \| 10 \| 4 \| 150 \| 56 \| 216 \| Phosphatidylinositol 4-kinase type 2-beta,Endolysin,Phosphatidylinositol 4-kinase type 2-beta \| Phosphatidylinositol 4-kinase type 2-beta,Endolysin,Phosphatidylinositol 4-kinase type 2-beta; pdb \| 260L\_A \| 1.0 \| 0.001088 \| 144 \| 0.196 \| 163 \| 102 \| 8 \| 6 \| 146 \| 2 \| 157 \| PROTEIN (LYSOZYME) \| PROTEIN (LYSOZYME); pdb \| 1EPY\_A \| 1.0 \| 0.0008349 \| 144 \| 0.201 \| 159 \| 97 \| 9 \| 6 \| 140 \| 2 \| 154 \| LYSOZYME \| LYSOZYME; pdb \| 1T8F\_A \| 1.0 \| 0.0006075 \| 144 \| 0.205 \| 170 \| 99 \| 8 \| 6 \| 150 \| 2 \| 160 \| Lysozyme \| Lysozyme; pdb \| 3SBB\_C \| 1.0 \| 0.0007918 \| 144 \| 0.195 \| 174 \| 102 \| 10 \| 4 \| 151 \| 1 \| 162 \| Lysozyme \| Lysozyme; pdb \| 2A4T\_A \| 1.0 \| 0.0006755 \| 144 \| 0.194 \| 170 \| 101 \| 8 \| 6 \| 150 \| 2 \| 160 \| Lysozyme \| Lysozyme; pdb \| 5KGR\_A \| 1.0 \| 0.0007918 \| 144 \| 0.196 \| 173 \| 102 \| 9 \| 6 \| 153 \| 2 \| 162 \| Endolysin \| Endolysin; pdb \| 2Q9D\_A \| 1.0 \| 0.0006406 \| 144 \| 0.2 \| 170 \| 100 \| 9 \| 6 \| 150 \| 2 \| 160 \| Lysozyme \| Lysozyme; pdb \| 3SB8\_A \| 1.0 \| 0.001577 \| 144 \| 0.192 \| 171 \| 104 \| 9 \| 4 \| 150 \| 3 \| 163 \| Lysozyme \| Lysozyme; pdb \| 2QB0\_D \| 1.0 \| 0.0005183 \| 144 \| 0.187 \| 165 \| 98 \| 7 \| 4 \| 143 \| 79 \| 232 \| TELSAM domain - Lysozyme chimera \| TELSAM domain - Lysozyme chimera; pdb \| 6CM4\_A \| 1.0 \| 0.0004421 \| 144 \| 0.222 \| 162 \| 90 \| 10 \| 6 \| 140 \| 185 \| 337 \| D(2) dopamine receptor, endolysin chimera \| D(2) dopamine receptor, endolysin chimera; pdb \| 1C65\_A \| 1.0 \| 0.001147 \| 143 \| 0.19 \| 173 \| 98 \| 10 \| 6 \| 150 \| 2 \| 160 \| PROTEIN (LYSOZYME) \| PROTEIN (LYSOZYME); pdb \| 3G3W\_A \| 1.0 \| 0.000751 \| 143 \| 0.203 \| 172 \| 97 \| 9 \| 6 \| 150 \| 2 \| 160 \| Lysozyme \| Lysozyme; pdb \| 3K2R\_A \| 1.0 \| 0.0007918 \| 143 \| 0.197 \| 172 \| 98 \| 9 \| 6 \| 150 \| 2 \| 160 \| Lysozyme \| Lysozyme; pdb \| 1L73\_A \| 1.0 \| 0.001577 \| 143 \| 0.19 \| 168 \| 102 \| 9 \| 7 \| 150 \| 3 \| 160 \| LYSOZYME \| LYSOZYME; pdb \| 3CDT\_A \| 1.0 \| 0.001663 \| 143 \| 0.195 \| 169 \| 102 \| 9 \| 6 \| 150 \| 2 \| 160 \| Lysozyme \| Lysozyme; pdb \| 1JQU\_C \| 1.0 \| 0.0002603 \| 143 \| 0.209 \| 172 \| 96 \| 12 \| 6 \| 146 \| 2 \| 164 \| Lysozyme \| Lysozyme; pdb \| 111L\_A \| 1.0 \| 0.002055 \| 142 \| 0.201 \| 169 \| 101 \| 10 \| 6 \| 150 \| 2 \| 160 \| T4 LYSOZYME \| T4 LYSOZYME; pdb \| 1CUP\_A \| 1.0 \| 0.001848 \| 142 \| 0.195 \| 169 \| 102 \| 9 \| 6 \| 150 \| 2 \| 160 \| LYSOZYME \| LYSOZYME; pdb \| 5JGV\_A \| 1.0 \| 0.0008349 \| 142 \| 0.197 \| 172 \| 98 \| 9 \| 6 \| 150 \| 2 \| 160 \| Endolysin \| Endolysin; pdb \| 4Z9G\_A \| 1.0 \| 0.0003771 \| 142 \| 0.198 \| 186 \| 107 \| 10 \| 5 \| 157 \| 110 \| 286 \| Corticotropin-releasing factor receptor 1,Lysozyme,Corticotropin-releasing factor receptor 1 \| Corticotropin-releasing factor receptor 1,Lysozyme,Corticotropin-releasing factor receptor 1; pdb \| 115L\_A \| 1.0 \| 0.0007122 \| 141 \| 0.2 \| 170 \| 100 \| 9 \| 6 \| 150 \| 2 \| 160 \| T4 LYSOZYME \| T4 LYSOZYME; pdb \| 122L\_A \| 1.0 \| 0.0007918 \| 141 \| 0.205 \| 170 \| 99 \| 9 \| 6 \| 150 \| 2 \| 160 \| T4 LYSOZYME \| T4 LYSOZYME; pdb \| 5KIM\_A \| 1.0 \| 0.0008349 \| 141 \| 0.2 \| 170 \| 100 \| 9 \| 6 \| 150 \| 2 \| 160 \| Endolysin \| Endolysin; pdb \| 2O79\_A \| 1.0 \| 0.0007918 \| 141 \| 0.194 \| 175 \| 101 \| 10 \| 1 \| 150 \| 1 \| 160 \| Lysozyme \| Lysozyme; pdb \| 1P6Y\_A \| 1.0 \| 0.0008803 \| 141 \| 0.188 \| 170 \| 102 \| 8 \| 6 \| 150 \| 2 \| 160 \| LYSOZYME \| LYSOZYME; pdb \| 5XPF\_A \| 1.0 \| 0.0008803 \| 141 \| 0.194 \| 170 \| 101 \| 9 \| 6 \| 150 \| 2 \| 160 \| Endolysin \| Endolysin; pdb \| 3PBL\_B \| 1.0 \| 0.0008803 \| 141 \| 0.209 \| 172 \| 96 \| 10 \| 4 \| 148 \| 180 \| 338 \| D(3) dopamine receptor, Lysozyme chimera \| D(3) dopamine receptor, Lysozyme chimera; pdb \| 4QKX\_A \| 1.0 \| 0.0007122 \| 141 \| 0.203 \| 177 \| 101 \| 11 \| 1 \| 150 \| 5 \| 168 \| Beta-2 adrenergic receptor \| Beta-2 adrenergic receptor; pdb \| 114L\_A \| 1.0 \| 0.0009282 \| 140 \| 0.194 \| 170 \| 101 \| 8 \| 6 \| 150 \| 2 \| 160 \| T4 LYSOZYME \| T4 LYSOZYME; pdb \| 232L\_A \| 1.0 \| 0.002055 \| 140 \| 0.185 \| 167 \| 106 \| 8 \| 6 \| 150 \| 2 \| 160 \| T4 LYSOZYME \| T4 LYSOZYME; pdb \| 162L\_A \| 1.0 \| 0.0007918 \| 140 \| 0.209 \| 172 \| 96 \| 11 \| 6 \| 150 \| 2 \| 160 \| T4 LYSOZYME \| T4 LYSOZYME; pdb \| 3C8R\_A \| 1.0 \| 0.0009282 \| 140 \| 0.203 \| 172 \| 97 \| 10 \| 6 \| 150 \| 2 \| 160 \| Lysozyme \| Lysozyme; pdb \| 1ZYT\_A \| 1.0 \| 0.0009282 \| 140 \| 0.2 \| 170 \| 100 \| 9 \| 6 \| 150 \| 2 \| 160 \| Lysozyme \| Lysozyme; pdb \| 4W58\_A \| 1.0 \| 0.001276 \| 140 \| 0.2 \| 175 \| 99 \| 11 \| 6 \| 153 \| 2 \| 162 \| Endolysin \| Endolysin; pdb \| 6P5V\_A \| 1.0 \| 0.001345 \| 140 \| 0.224 \| 178 \| 95 \| 12 \| 4 \| 153 \| 13 \| 175 \| Lysozyme,DCN1-like protein 1 fusion \| Lysozyme,DCN1-like protein 1 fusion; pdb \| 217L\_A \| 1.0 \| 0.0009282 \| 139 \| 0.194 \| 170 \| 101 \| 8 \| 6 \| 150 \| 2 \| 160 \| T4 LYSOZYME \| T4 LYSOZYME; pdb \| 161L\_A \| 1.0 \| 0.0008803 \| 139 \| 0.2 \| 170 \| 100 \| 9 \| 6 \| 150 \| 2 \| 160 \| T4 LYSOZYME \| T4 LYSOZYME; pdb \| 1L88\_A \| 1.0 \| 0.001088 \| 139 \| 0.191 \| 172 \| 99 \| 9 \| 6 \| 150 \| 2 \| 160 \| T4 LYSOZYME \| T4 LYSOZYME; pdb \| 3L64\_A \| 1.0 \| 0.00254 \| 139 \| 0.189 \| 169 \| 103 \| 9 \| 6 \| 150 \| 2 \| 160 \| Lysozyme \| Lysozyme; pdb \| 1P46\_A \| 1.0 \| 0.001418 \| 139 \| 0.196 \| 173 \| 102 \| 9 \| 6 \| 153 \| 2 \| 162 \| LYSOZYME \| LYSOZYME; pdb \| 1PQK\_C \| 1.0 \| 0.001418 \| 139 \| 0.208 \| 173 \| 97 \| 10 \| 6 \| 151 \| 2 \| 161 \| Lysozyme \| Lysozyme; pdb \| 5VNQ\_A \| 1.0 \| 0.001032 \| 139 \| 0.197 \| 172 \| 98 \| 9 \| 6 \| 150 \| 2 \| 160 \| Endolysin \| Endolysin; pdb \| 5V88\_A \| 1.0 \| 0.001663 \| 139 \| 0.207 \| 178 \| 104 \| 10 \| 1 \| 153 \| 9 \| 174 \| Lysozyme,DCN1-like protein 1 \| Lysozyme,DCN1-like protein 1; pdb \| 4DJH\_B \| 1.0 \| 0.00121 \| 139 \| 0.209 \| 181 \| 99 \| 12 \| 5 \| 155 \| 207 \| 373 \| Kappa-type opioid receptor, Lysozyme \| Kappa-type opioid receptor, Lysozyme; pdb \| 5KII\_A \| 1.0 \| 0.001032 \| 138 \| 0.201 \| 169 \| 100 \| 9 \| 6 \| 150 \| 2 \| 159 \| Endolysin \| Endolysin; pdb \| 175L\_A \| 1.0 \| 0.001418 \| 138 \| 0.203 \| 172 \| 97 \| 10 \| 6 \| 150 \| 2 \| 160 \| T4 LYSOZYME \| T4 LYSOZYME; pdb \| 192L\_A \| 1.0 \| 0.0006075 \| 138 \| 0.218 \| 165 \| 89 \| 10 \| 6 \| 143 \| 2 \| 153 \| LYSOZYME \| LYSOZYME; pdb \| 1L74\_A \| 1.0 \| 0.0006755 \| 138 \| 0.189 \| 169 \| 101 \| 9 \| 7 \| 150 \| 3 \| 160 \| LYSOZYME \| LYSOZYME; pdb \| 237L\_A \| 1.0 \| 0.0009788 \| 138 \| 0.203 \| 172 \| 97 \| 10 \| 6 \| 150 \| 2 \| 160 \| T4 LYSOZYME \| T4 LYSOZYME; pdb \| 1L26\_A \| 1.0 \| 0.0009788 \| 138 \| 0.195 \| 169 \| 100 \| 9 \| 7 \| 150 \| 3 \| 160 \| T4 LYSOZYME \| T4 LYSOZYME; pdb \| 1L53\_A \| 1.0 \| 0.0004915 \| 138 \| 0.216 \| 162 \| 91 \| 11 \| 6 \| 140 \| 2 \| 154 \| T4 LYSOZYME \| T4 LYSOZYME; pdb \| 1P2R\_A \| 1.0 \| 0.00121 \| 138 \| 0.197 \| 172 \| 98 \| 9 \| 6 \| 150 \| 2 \| 160 \| LYSOZYME \| LYSOZYME; pdb \| 2F47\_A \| 1.0 \| 0.0008803 \| 138 \| 0.176 \| 181 \| 102 \| 7 \| 6 \| 150 \| 2 \| 171 \| Lysozyme \| Lysozyme; pdb \| 4ARJ\_B \| 1.0 \| 0.0007918 \| 138 \| 0.209 \| 162 \| 92 \| 10 \| 6 \| 140 \| 152 \| 304 \| PESTICIN, LYSOZYME \| PESTICIN, LYSOZYME; pdb \| 8A5X\_A \| 1.0 \| 0.0005762 \| 138 \| 0.201 \| 164 \| 94 \| 11 \| 4 \| 140 \| 70 \| 223 \| Phosphatidylinositol 4-kinase type 2-beta,Endolysin \| Phosphatidylinositol 4-kinase type 2-beta,Endolysin; pdb \| 4EVX\_B \| 1.0 \| 0.008593 \| 137 \| 0.25 \| 92 \| 66 \| 2 \| 2 \| 93 \| 1 \| 89 \| Putative phage endolysin \| Putative phage endolysin; pdb \| 7LX8\_A \| 1.0 \| 0.001032 \| 137 \| 0.195 \| 169 \| 101 \| 8 \| 6 \| 150 \| 2 \| 159 \| Lysozyme \| Lysozyme; pdb \| 139L\_A \| 1.0 \| 0.001276 \| 137 \| 0.203 \| 172 \| 97 \| 10 \| 6 \| 150 \| 2 \| 160 \| T4 LYSOZYME \| T4 LYSOZYME; pdb \| 1L66\_A \| 1.0 \| 0.001276 \| 137 \| 0.203 \| 172 \| 97 \| 10 \| 6 \| 150 \| 2 \| 160 \| LYSOZYME \| LYSOZYME; pdb \| 2RAY\_X \| 1.0 \| 0.001276 \| 137 \| 0.194 \| 170 \| 101 \| 9 \| 6 \| 150 \| 2 \| 160 \| Lysozyme \| Lysozyme; pdb \| 3CDV\_A \| 1.0 \| 0.00121 \| 137 \| 0.203 \| 172 \| 97 \| 10 \| 6 \| 150 \| 2 \| 160 \| Lysozyme \| Lysozyme; pdb \| 1L22\_A \| 1.0 \| 0.001418 \| 137 \| 0.208 \| 173 \| 100 \| 10 \| 6 \| 153 \| 2 \| 162 \| T4 LYSOZYME \| T4 LYSOZYME; pdb \| 6A73\_B \| 1.0 \| 0.000751 \| 137 \| 0.222 \| 162 \| 90 \| 10 \| 6 \| 140 \| 131 \| 283 \| COP9 signalosome complex subunit 2,Endolysin \| COP9 signalosome complex subunit 2,Endolysin; pdb \| 1XJU\_B \| 1.0 \| 0.001753 \| 136 \| 0.184 \| 163 \| 98 \| 7 \| 2 \| 144 \| 1 \| 148 \| Lysozyme \| Lysozyme; pdb \| 180L\_B \| 1.0 \| 0.002978 \| 136 \| 0.183 \| 169 \| 104 \| 9 \| 6 \| 150 \| 2 \| 160 \| LYSOZYME \| LYSOZYME; pdb \| 3C7Y\_A \| 1.0 \| 0.001418 \| 136 \| 0.194 \| 170 \| 101 \| 8 \| 6 \| 150 \| 2 \| 160 \| lysozyme \| lysozyme; pdb \| 3LZM\_A \| 1.0 \| 0.001577 \| 136 \| 0.217 \| 175 \| 96 \| 13 \| 6 \| 153 \| 2 \| 162 \| T4 LYSOZYME \| T4 LYSOZYME; pdb \| 2OU9\_A \| 1.0 \| 0.001418 \| 136 \| 0.2 \| 170 \| 100 \| 9 \| 6 \| 150 \| 2 \| 160 \| Lysozyme \| Lysozyme; pdb \| 159L\_A \| 1.0 \| 0.001663 \| 135 \| 0.2 \| 170 \| 100 \| 8 \| 6 \| 150 \| 2 \| 160 \| T4 LYSOZYME \| T4 LYSOZYME; pdb \| 7LOC\_A \| 1.0 \| 0.001577 \| 135 \| 0.203 \| 172 \| 101 \| 9 \| 6 \| 153 \| 2 \| 161 \| Lysozyme \| Lysozyme; pdb \| 4W56\_A \| 1.0 \| 0.001495 \| 135 \| 0.194 \| 170 \| 101 \| 9 \| 6 \| 150 \| 2 \| 160 \| Endolysin \| Endolysin; pdb \| 6XC0\_B \| 1.0 \| 0.00314 \| 135 \| 0.184 \| 173 \| 102 \| 7 \| 7 \| 150 \| 2 \| 164 \| Lysozyme \| Lysozyme; pdb \| 4K5Y\_A \| 1.0 \| 0.002285 \| 135 \| 0.195 \| 174 \| 104 \| 9 \| 5 \| 153 \| 106 \| 268 \| Corticotropin-releasing factor receptor 1, T4-Lysozyme chimeric construct \| Corticotropin-releasing factor receptor 1, T4-Lysozyme chimeric construct; pdb \| 158L\_A \| 1.0 \| 0.001495 \| 134 \| 0.203 \| 172 \| 97 \| 10 \| 6 \| 150 \| 2 \| 160 \| T4 LYSOZYME \| T4 LYSOZYME; pdb \| 173L\_A \| 1.0 \| 0.002824 \| 134 \| 0.19 \| 173 \| 98 \| 9 \| 7 \| 151 \| 3 \| 161 \| T4 LYSOZYME \| T4 LYSOZYME; pdb \| 1TLA\_A \| 1.0 \| 0.001495 \| 134 \| 0.205 \| 170 \| 99 \| 10 \| 6 \| 150 \| 2 \| 160 \| T4 LYSOZYME \| T4 LYSOZYME; pdb \| 213L\_A \| 1.0 \| 0.0009282 \| 134 \| 0.218 \| 160 \| 94 \| 8 \| 6 \| 140 \| 2 \| 155 \| T4 LYSOZYME \| T4 LYSOZYME; pdb \| 5NX0\_A \| 1.0 \| 0.001577 \| 134 \| 0.203 \| 172 \| 97 \| 10 \| 6 \| 150 \| 2 \| 160 \| Endolysin \| Endolysin; pdb \| 2Q9E\_B \| 1.0 \| 0.0007918 \| 134 \| 0.212 \| 160 \| 94 \| 9 \| 6 \| 140 \| 2 \| 154 \| Lysozyme \| Lysozyme; pdb \| 1JTN\_B \| 1.0 \| 0.001753 \| 134 \| 0.192 \| 177 \| 102 \| 11 \| 6 \| 155 \| 2 \| 164 \| LYSOZYME \| LYSOZYME; pdb \| 6D9M\_A \| 1.0 \| 0.002285 \| 134 \| 0.202 \| 173 \| 98 \| 10 \| 6 \| 151 \| 2 \| 161 \| Fusion protein of Endolysin,Response receiver sensor diguanylate cyclase, GAF domain-containing \| Fusion protein of Endolysin,Response receiver sensor diguanylate cyclase, GAF domain-containing; pdb \| 3QAK\_A \| 1.0 \| 0.001949 \| 134 \| 0.198 \| 181 \| 106 \| 10 \| 4 \| 156 \| 196 \| 365 \| Adenosine receptor A2a,lysozyme chimera \| Adenosine receptor A2a,lysozyme chimera; pdb \| 7BU7\_A \| 1.0 \| 0.001147 \| 134 \| 0.183 \| 185 \| 108 \| 11 \| 1 \| 157 \| 1 \| 170 \| Endolysin,Endolysin,Beta-1 adrenergic receptor chimera \| Endolysin,Endolysin,Beta-1 adrenergic receptor chimera; pdb \| 1KW5\_A \| 1.0 \| 0.001753 \| 133 \| 0.188 \| 170 \| 102 \| 9 \| 6 \| 150 \| 2 \| 160 \| LYSOZYME \| LYSOZYME; pdb \| 230L\_A \| 1.0 \| 0.00254 \| 133 \| 0.196 \| 163 \| 102 \| 8 \| 6 \| 146 \| 2 \| 157 \| T4 LYSOZYME \| T4 LYSOZYME; pdb \| 3C7Z\_A \| 1.0 \| 0.00121 \| 133 \| 0.204 \| 166 \| 97 \| 9 \| 6 \| 146 \| 2 \| 157 \| Lysozyme \| Lysozyme; pdb \| 1L97\_B \| 1.0 \| 0.0009282 \| 133 \| 0.223 \| 161 \| 89 \| 11 \| 7 \| 140 \| 3 \| 154 \| T4 LYSOZYME \| T4 LYSOZYME; pdb \| 1JTM\_A \| 1.0 \| 0.001147 \| 133 \| 0.221 \| 167 \| 102 \| 9 \| 6 \| 147 \| 2 \| 165 \| LYSOZYME \| LYSOZYME; pdb \| 3VW7\_A \| 1.0 \| 0.00254 \| 133 \| 0.204 \| 176 \| 104 \| 9 \| 4 \| 154 \| 205 \| 369 \| Proteinase-activated receptor 1, Lysozyme \| Proteinase-activated receptor 1, Lysozyme; pdb \| 6WJC\_A \| 1.0 \| 0.001032 \| 133 \| 0.192 \| 177 \| 97 \| 11 \| 6 \| 154 \| 203 \| 361 \| Muscarinic acetylcholine receptor M1,Endolysin fusion \| Muscarinic acetylcholine receptor M1,Endolysin fusion; pdb \| 5T04\_A \| 1.0 \| 0.0007122 \| 133 \| 0.22 \| 186 \| 101 \| 13 \| 6 \| 157 \| 219 \| 394 \| Neurotensin receptor type 1,Endolysin,Neurotensin receptor type 1 \| Neurotensin receptor type 1,Endolysin,Neurotensin receptor type 1; pdb \| 137L\_A \| 1.0 \| 0.004092 \| 132 \| 0.194 \| 170 \| 101 \| 9 \| 6 \| 150 \| 2 \| 160 \| T4 LYSOZYME \| T4 LYSOZYME; pdb \| 180L\_A \| 1.0 \| 0.001848 \| 132 \| 0.182 \| 170 \| 103 \| 8 \| 6 \| 150 \| 2 \| 160 \| LYSOZYME \| LYSOZYME; pdb \| 1L87\_A \| 1.0 \| 0.001276 \| 132 \| 0.216 \| 166 \| 95 \| 10 \| 6 \| 146 \| 2 \| 157 \| T4 LYSOZYME \| T4 LYSOZYME; pdb \| 1L95\_A \| 1.0 \| 0.001949 \| 132 \| 0.197 \| 172 \| 98 \| 10 \| 6 \| 150 \| 2 \| 160 \| T4 LYSOZYME \| T4 LYSOZYME; pdb \| 102L\_A \| 1.0 \| 0.001949 \| 132 \| 0.208 \| 173 \| 96 \| 11 \| 6 \| 150 \| 2 \| 161 \| T4 LYSOZYME \| T4 LYSOZYME; pdb \| 1PQO\_A \| 1.0 \| 0.002285 \| 132 \| 0.19 \| 173 \| 103 \| 10 \| 6 \| 153 \| 2 \| 162 \| Lysozyme \| Lysozyme; pdb \| 262L\_B \| 1.0 \| 0.002055 \| 132 \| 0.176 \| 181 \| 102 \| 7 \| 6 \| 150 \| 2 \| 171 \| LYSOZYME \| LYSOZYME; pdb \| 206L\_A \| 1.0 \| 0.002055 \| 131 \| 0.203 \| 172 \| 97 \| 10 \| 6 \| 150 \| 2 \| 160 \| LYSOZYME \| LYSOZYME; pdb \| 109L\_A \| 1.0 \| 0.001753 \| 131 \| 0.209 \| 172 \| 96 \| 11 \| 6 \| 150 \| 2 \| 160 \| T4 LYSOZYME \| T4 LYSOZYME; pdb \| 174L\_B \| 1.0 \| 0.0008349 \| 131 \| 0.202 \| 163 \| 92 \| 10 \| 6 \| 140 \| 2 \| 154 \| T4 LYSOZYME \| T4 LYSOZYME; pdb \| 178L\_A \| 1.0 \| 0.003491 \| 131 \| 0.209 \| 172 \| 96 \| 10 \| 6 \| 150 \| 2 \| 160 \| T4 LYSOZYME \| T4 LYSOZYME; pdb \| 1CV3\_A \| 1.0 \| 0.002055 \| 131 \| 0.203 \| 172 \| 97 \| 10 \| 6 \| 150 \| 2 \| 160 \| LYSOZYME \| LYSOZYME; pdb \| 6PGZ\_B \| 1.0 \| 0.001848 \| 131 \| 0.19 \| 173 \| 98 \| 9 \| 6 \| 150 \| 2 \| 160 \| Endolysin \| Endolysin; pdb \| 1L00\_A \| 1.0 \| 0.002978 \| 131 \| 0.195 \| 164 \| 99 \| 9 \| 7 \| 146 \| 3 \| 157 \| T4 LYSOZYME \| T4 LYSOZYME; pdb \| 1P64\_A \| 1.0 \| 0.002055 \| 131 \| 0.205 \| 170 \| 99 \| 9 \| 6 \| 150 \| 2 \| 160 \| LYSOZYME \| LYSOZYME; pdb \| 7CN7\_A \| 1.0 \| 0.002167 \| 131 \| 0.194 \| 175 \| 92 \| 9 \| 7 \| 148 \| 12 \| 170 \| Baseplate central spike complex protein gp5 \| Baseplate central spike complex protein gp5; pdb \| 1D9W\_A \| 1.0 \| 0.001848 \| 130 \| 0.21 \| 166 \| 96 \| 10 \| 6 \| 146 \| 2 \| 157 \| PROTEIN (LYSOZYME) \| PROTEIN (LYSOZYME); pdb \| 1KW7\_A \| 1.0 \| 0.002285 \| 130 \| 0.191 \| 172 \| 99 \| 10 \| 6 \| 150 \| 2 \| 160 \| LYSOZYME \| LYSOZYME; pdb \| 2RBO\_A \| 1.0 \| 0.002285 \| 130 \| 0.194 \| 170 \| 101 \| 9 \| 6 \| 150 \| 2 \| 160 \| Lysozyme \| Lysozyme; pdb \| 7DWS\_A \| 1.0 \| 0.001577 \| 130 \| 0.205 \| 170 \| 96 \| 9 \| 4 \| 146 \| 2 \| 159 \| Endolysin \| Endolysin; pdb \| 1G0J\_A \| 1.0 \| 0.002285 \| 130 \| 0.203 \| 172 \| 97 \| 10 \| 6 \| 150 \| 2 \| 160 \| PROTEIN (LYSOZYME) \| PROTEIN (LYSOZYME); pdb \| 1L19\_A \| 1.0 \| 0.00314 \| 130 \| 0.206 \| 165 \| 98 \| 10 \| 6 \| 146 \| 2 \| 157 \| T4 LYSOZYME \| T4 LYSOZYME; pdb \| 1QT5\_A \| 1.0 \| 0.002285 \| 130 \| 0.194 \| 170 \| 101 \| 9 \| 6 \| 150 \| 2 \| 160 \| PROTEIN (T4 LYSOZYME) \| PROTEIN (T4 LYSOZYME); pdb \| 1SWZ\_A \| 1.0 \| 0.003311 \| 130 \| 0.208 \| 173 \| 100 \| 10 \| 6 \| 153 \| 2 \| 162 \| Lysozyme \| Lysozyme; pdb \| 4YXA\_F \| 1.0 \| 0.002285 \| 130 \| 0.182 \| 170 \| 101 \| 8 \| 7 \| 151 \| 36 \| 192 \| Oxygen-regulated invasion protein OrgB,Endolysin \| Oxygen-regulated invasion protein OrgB,Endolysin; pdb \| 5YQR\_A \| 1.0 \| 0.001418 \| 130 \| 0.198 \| 161 \| 96 \| 7 \| 5 \| 143 \| 1 \| 150 \| Endolysin/Membrane-anchored lipid-binding protein LAM6 fusion protein \| Endolysin/Membrane-anchored lipid-binding protein LAM6 fusion protein; pdb \| 7MWY\_A \| 1.0 \| 0.003881 \| 130 \| 0.194 \| 175 \| 101 \| 10 \| 4 \| 151 \| 1 \| 162 \| STING \| STING; pdb \| 1T8G\_A \| 1.0 \| 0.001848 \| 129 \| 0.208 \| 158 \| 93 \| 9 \| 6 \| 140 \| 2 \| 150 \| Lysozyme \| Lysozyme; pdb \| 195L\_A \| 1.0 \| 0.002285 \| 129 \| 0.197 \| 172 \| 98 \| 10 \| 6 \| 150 \| 2 \| 160 \| LYSOZYME \| LYSOZYME; pdb \| 1C64\_A \| 1.0 \| 0.002678 \| 129 \| 0.203 \| 172 \| 97 \| 10 \| 6 \| 150 \| 2 \| 160 \| PROTEIN (LYSOZYME) \| PROTEIN (LYSOZYME); pdb \| 149L\_A \| 1.0 \| 0.001949 \| 129 \| 0.215 \| 167 \| 92 \| 11 \| 7 \| 146 \| 3 \| 157 \| T4 LYSOZYME \| T4 LYSOZYME; pdb \| 1JQU\_A \| 1.0 \| 0.0004661 \| 129 \| 0.195 \| 169 \| 100 \| 10 \| 7 \| 146 \| 3 \| 164 \| Lysozyme \| Lysozyme; pdb \| 1L35\_A \| 1.0 \| 0.00254 \| 129 \| 0.203 \| 172 \| 97 \| 10 \| 6 \| 150 \| 2 \| 160 \| T4 LYSOZYME \| T4 LYSOZYME; pdb \| 6XMV\_A \| 1.0 \| 0.000751 \| 129 \| 0.197 \| 167 \| 100 \| 8 \| 4 \| 142 \| 145 \| 305 \| Hemolysin,Endolysin \| Hemolysin,Endolysin; pdb \| 5KI2\_A \| 1.0 \| 0.002285 \| 128 \| 0.191 \| 172 \| 98 \| 10 \| 6 \| 150 \| 2 \| 159 \| Endolysin \| Endolysin; pdb \| 118L\_A \| 1.0 \| 0.002678 \| 128 \| 0.194 \| 170 \| 101 \| 9 \| 6 \| 150 \| 2 \| 160 \| T4 LYSOZYME \| T4 LYSOZYME; pdb \| 199L\_A \| 1.0 \| 0.002978 \| 128 \| 0.191 \| 172 \| 99 \| 9 \| 6 \| 150 \| 2 \| 160 \| LYSOZYME \| LYSOZYME; pdb \| 1LYE\_A \| 1.0 \| 0.00254 \| 128 \| 0.203 \| 172 \| 97 \| 11 \| 6 \| 150 \| 2 \| 160 \| T4 LYSOZYME \| T4 LYSOZYME; pdb \| 175L\_B \| 1.0 \| 0.0007918 \| 128 \| 0.207 \| 169 \| 100 \| 9 \| 6 \| 146 \| 2 \| 164 \| T4 LYSOZYME \| T4 LYSOZYME; pdb \| 1JQU\_B \| 1.0 \| 0.0006406 \| 128 \| 0.213 \| 169 \| 97 \| 10 \| 7 \| 146 \| 3 \| 164 \| Lysozyme \| Lysozyme; pdb \| 1L11\_A \| 1.0 \| 0.003491 \| 128 \| 0.2 \| 175 \| 99 \| 10 \| 6 \| 153 \| 2 \| 162 \| T4 LYSOZYME \| T4 LYSOZYME; pdb \| 1P3N\_A \| 1.0 \| 0.00314 \| 128 \| 0.203 \| 172 \| 97 \| 9 \| 6 \| 150 \| 2 \| 160 \| LYSOZYME \| LYSOZYME; pdb \| 3JR6\_D \| 1.0 \| 0.00314 \| 128 \| 0.194 \| 175 \| 99 \| 9 \| 7 \| 150 \| 3 \| 166 \| Lysozyme \| Lysozyme; pdb \| 6XON\_A \| 1.0 \| 0.004092 \| 128 \| 0.2 \| 170 \| 99 \| 9 \| 7 \| 153 \| 2 \| 157 \| Lysozyme DCN1-like protein 1 chimera \| Lysozyme DCN1-like protein 1 chimera; pdb \| 3V2Y\_A \| 1.0 \| 0.005334 \| 128 \| 0.196 \| 163 \| 97 \| 7 \| 6 \| 144 \| 210 \| 362 \| Sphingosine 1-phosphate receptor 1, Lysozyme chimera (E.C.3.2.1.17) \| Sphingosine 1-phosphate receptor 1, Lysozyme chimera (E.C.3.2.1.17); pdb \| 3JR6\_B \| 1.0 \| 0.003681 \| 127 \| 0.205 \| 151 \| 97 \| 9 \| 7 \| 140 \| 3 \| 147 \| Lysozyme \| Lysozyme; pdb \| 5JWU\_A \| 1.0 \| 0.002978 \| 127 \| 0.197 \| 172 \| 98 \| 10 \| 6 \| 150 \| 2 \| 160 \| Endolysin \| Endolysin; pdb \| 1L89\_A \| 1.0 \| 0.002055 \| 127 \| 0.18 \| 172 \| 101 \| 9 \| 6 \| 150 \| 2 \| 160 \| T4 LYSOZYME \| T4 LYSOZYME; pdb \| 1L05\_A \| 1.0 \| 0.003681 \| 127 \| 0.2 \| 175 \| 99 \| 10 \| 6 \| 153 \| 2 \| 162 \| T4 LYSOZYME \| T4 LYSOZYME; pdb \| 262L\_A \| 1.0 \| 0.002167 \| 127 \| 0.191 \| 178 \| 96 \| 9 \| 6 \| 146 \| 2 \| 168 \| LYSOZYME \| LYSOZYME; pdb \| 6WSK\_A \| 1.0 \| 0.002824 \| 127 \| 0.184 \| 157 \| 95 \| 8 \| 4 \| 150 \| 1 \| 134 \| Endolysin,CB1 cannabinoid receptor-interacting protein 1 fusion \| Endolysin,CB1 cannabinoid receptor-interacting protein 1 fusion; pdb \| 120L\_A \| 1.0 \| 0.001949 \| 126 \| 0.213 \| 169 \| 92 \| 11 \| 6 \| 146 \| 2 \| 157 \| T4 LYSOZYME \| T4 LYSOZYME; pdb \| 234L\_A \| 1.0 \| 0.002167 \| 126 \| 0.21 \| 166 \| 96 \| 9 \| 6 \| 146 \| 2 \| 157 \| T4 LYSOZYME \| T4 LYSOZYME; pdb \| 160L\_A \| 1.0 \| 0.003311 \| 126 \| 0.203 \| 172 \| 97 \| 11 \| 6 \| 150 \| 2 \| 160 \| T4 LYSOZYME \| T4 LYSOZYME; pdb \| 4S0W\_B \| 1.0 \| 0.001577 \| 126 \| 0.222 \| 162 \| 90 \| 10 \| 6 \| 140 \| 2 \| 154 \| Lysozyme \| Lysozyme; pdb \| 3SB8\_C \| 1.0 \| 0.005059 \| 126 \| 0.187 \| 165 \| 105 \| 8 \| 4 \| 146 \| 1 \| 158 \| Lysozyme \| Lysozyme; pdb \| 189L\_A \| 1.0 \| 0.001949 \| 126 \| 0.2 \| 170 \| 96 \| 11 \| 4 \| 146 \| 1 \| 157 \| T4 LYSOZYME \| T4 LYSOZYME; pdb \| 1L32\_A \| 1.0 \| 0.002409 \| 126 \| 0.21 \| 166 \| 96 \| 10 \| 6 \| 146 \| 2 \| 157 \| T4 LYSOZYME \| T4 LYSOZYME; pdb \| 1L34\_A \| 1.0 \| 0.002409 \| 126 \| 0.22 \| 168 \| 92 \| 12 \| 6 \| 146 \| 2 \| 157 \| T4 LYSOZYME \| T4 LYSOZYME; pdb \| 1QTZ\_A \| 1.0 \| 0.006952 \| 126 \| 0.189 \| 169 \| 103 \| 9 \| 6 \| 150 \| 2 \| 160 \| PROTEIN (T4 LYSOZYME) \| PROTEIN (T4 LYSOZYME); pdb \| 1T97\_B \| 1.0 \| 0.001949 \| 126 \| 0.206 \| 179 \| 92 \| 11 \| 6 \| 146 \| 2 \| 168 \| Lysozyme \| Lysozyme; pdb \| 7F8X\_A \| 1.0 \| 0.002824 \| 126 \| 0.194 \| 180 \| 108 \| 11 \| 4 \| 158 \| 203 \| 370 \| Cholecystokinin receptor type A,Endolysin \| Cholecystokinin receptor type A,Endolysin; pdb \| 6E67\_A \| 1.0 \| 0.004092 \| 126 \| 0.202 \| 178 \| 100 \| 11 \| 6 \| 154 \| 205 \| 369 \| Beta-2 adrenergic receptor,Endolysin,Guanine nucleotide-binding protein G(s) subunit alpha isoforms short,Beta-2 adrenergic receptor chimera \| Beta-2 adrenergic receptor,Endolysin,Guanine nucleotide-binding protein G(s) subunit alpha isoforms short,Beta-2 adrenergic receptor chimera; pdb \| 5XF1\_A \| 1.0 \| 0.001345 \| 126 \| 0.169 \| 177 \| 116 \| 8 \| 5 \| 157 \| 230 \| 399 \| Glucagon receptor,Endolysin,Glucagon receptor \| Glucagon receptor,Endolysin,Glucagon receptor; pdb \| 236L\_A \| 1.0 \| 0.00314 \| 125 \| 0.2 \| 170 \| 100 \| 9 \| 6 \| 150 \| 2 \| 160 \| T4 LYSOZYME \| T4 LYSOZYME; pdb \| 177L\_A \| 1.0 \| 0.004797 \| 125 \| 0.176 \| 170 \| 97 \| 8 \| 6 \| 146 \| 2 \| 157 \| T4 LYSOZYME \| T4 LYSOZYME; pdb \| 1CX7\_A \| 1.0 \| 0.003491 \| 125 \| 0.176 \| 170 \| 104 \| 9 \| 6 \| 150 \| 2 \| 160 \| LYSOZYME \| LYSOZYME; pdb \| 1KY0\_A \| 1.0 \| 0.001848 \| 125 \| 0.2 \| 160 \| 96 \| 9 \| 6 \| 140 \| 2 \| 154 \| LYSOZYME \| LYSOZYME; pdb \| 2L78\_A \| 1.0 \| 0.002285 \| 125 \| 0.22 \| 168 \| 92 \| 11 \| 6 \| 146 \| 2 \| 157 \| T4 LYSOZYME \| T4 LYSOZYME; pdb \| 3CDR\_A \| 1.0 \| 0.003681 \| 125 \| 0.203 \| 172 \| 97 \| 10 \| 6 \| 150 \| 2 \| 160 \| Lysozyme \| Lysozyme; pdb \| 1L20\_A \| 1.0 \| 0.005624 \| 125 \| 0.212 \| 165 \| 97 \| 11 \| 6 \| 146 \| 2 \| 157 \| T4 LYSOZYME \| T4 LYSOZYME; pdb \| 7MWZ\_A \| 1.0 \| 0.002055 \| 125 \| 0.207 \| 164 \| 94 \| 10 \| 4 \| 140 \| 1 \| 155 \| STING \| STING; pdb \| 7MI1\_A \| 1.0 \| 0.003491 \| 125 \| 0.206 \| 165 \| 98 \| 11 \| 6 \| 146 \| 1663 \| 1818 \| Chimera protein of Dynein and Endolysin \| Chimera protein of Dynein and Endolysin; pdb \| 5JWW\_A \| 1.0 \| 0.003311 \| 124 \| 0.2 \| 170 \| 100 \| 9 \| 6 \| 150 \| 2 \| 160 \| Endolysin \| Endolysin; pdb \| 156L\_A \| 1.0 \| 0.003881 \| 124 \| 0.197 \| 172 \| 98 \| 9 \| 6 \| 150 \| 2 \| 160 \| T4 LYSOZYME \| T4 LYSOZYME; pdb \| 1L77\_A \| 1.0 \| 0.002055 \| 124 \| 0.206 \| 165 \| 91 \| 10 \| 6 \| 143 \| 2 \| 153 \| T4 LYSOZYME \| T4 LYSOZYME; pdb \| 1L09\_A \| 1.0 \| 0.00593 \| 124 \| 0.196 \| 163 \| 102 \| 8 \| 6 \| 146 \| 2 \| 157 \| T4 LYSOZYME \| T4 LYSOZYME; pdb \| 1L31\_A \| 1.0 \| 0.00314 \| 124 \| 0.198 \| 166 \| 98 \| 8 \| 6 \| 146 \| 2 \| 157 \| T4 LYSOZYME \| T4 LYSOZYME; pdb \| 1SSY\_B \| 1.0 \| 0.00455 \| 124 \| 0.182 \| 159 \| 100 \| 9 \| 6 \| 140 \| 2 \| 154 \| Lysozyme \| Lysozyme; pdb \| 1T97\_A \| 1.0 \| 0.003311 \| 124 \| 0.193 \| 176 \| 97 \| 10 \| 6 \| 150 \| 2 \| 163 \| Lysozyme \| Lysozyme; pdb \| 7XKA\_A \| 1.0 \| 0.002285 \| 124 \| 0.196 \| 168 \| 99 \| 9 \| 1 \| 143 \| 5 \| 161 \| Endolysin,Beta-2 adrenergic receptor \| Endolysin,Beta-2 adrenergic receptor; pdb \| 231L\_A \| 1.0 \| 0.004315 \| 123 \| 0.203 \| 172 \| 97 \| 10 \| 6 \| 150 \| 2 \| 160 \| T4 LYSOZYME \| T4 LYSOZYME; pdb \| 233L\_A \| 1.0 \| 0.004092 \| 123 \| 0.197 \| 172 \| 98 \| 10 \| 6 \| 150 \| 2 \| 160 \| T4 LYSOZYME \| T4 LYSOZYME; pdb \| 1C6N\_A \| 1.0 \| 0.004315 \| 123 \| 0.188 \| 170 \| 102 \| 9 \| 6 \| 150 \| 2 \| 160 \| PROTEIN (LYSOZYME) \| PROTEIN (LYSOZYME); pdb \| 1LLH\_A \| 1.0 \| 0.00314 \| 123 \| 0.204 \| 166 \| 97 \| 9 \| 6 \| 146 \| 2 \| 157 \| Lysozyme \| Lysozyme; pdb \| 4LZM\_A \| 1.0 \| 0.002978 \| 123 \| 0.197 \| 172 \| 98 \| 10 \| 6 \| 150 \| 2 \| 160 \| T4 LYSOZYME \| T4 LYSOZYME; pdb \| 176L\_B \| 1.0 \| 0.00314 \| 123 \| 0.21 \| 166 \| 96 \| 7 \| 6 \| 146 \| 2 \| 157 \| T4 LYSOZYME \| T4 LYSOZYME; pdb \| 1L14\_A \| 1.0 \| 0.005059 \| 123 \| 0.196 \| 173 \| 102 \| 9 \| 6 \| 153 \| 2 \| 162 \| T4 LYSOZYME \| T4 LYSOZYME; pdb \| 4EXM\_C \| 1.0 \| 0.002824 \| 123 \| 0.191 \| 172 \| 97 \| 11 \| 5 \| 146 \| 166 \| 325 \| Pesticin, Lysozyme Chimera \| Pesticin, Lysozyme Chimera; pdb \| 5DGY\_A \| 1.0 \| 0.002409 \| 123 \| 0.217 \| 161 \| 90 \| 10 \| 7 \| 140 \| 2 \| 153 \| Endolysin,Rhodopsin,S-arrestin \| Endolysin,Rhodopsin,S-arrestin; pdb \| 210L\_A \| 1.0 \| 0.004315 \| 122 \| 0.192 \| 166 \| 98 \| 9 \| 6 \| 146 \| 2 \| 156 \| T4 LYSOZYME \| T4 LYSOZYME; pdb \| 198L\_A \| 1.0 \| 0.00455 \| 122 \| 0.197 \| 172 \| 98 \| 10 \| 6 \| 150 \| 2 \| 160 \| LYSOZYME \| LYSOZYME; pdb \| 1D3N\_A \| 1.0 \| 0.00455 \| 122 \| 0.176 \| 170 \| 104 \| 8 \| 6 \| 150 \| 2 \| 160 \| LYSOZYME \| LYSOZYME; pdb \| 168L\_D \| 1.0 \| 0.00314 \| 122 \| 0.207 \| 169 \| 93 \| 11 \| 6 \| 146 \| 2 \| 157 \| T4 LYSOZYME \| T4 LYSOZYME; pdb \| 1P7S\_A \| 1.0 \| 0.005059 \| 122 \| 0.201 \| 159 \| 97 \| 10 \| 6 \| 140 \| 2 \| 154 \| LYSOZYME \| LYSOZYME; pdb \| 1QTV\_A \| 1.0 \| 0.003311 \| 122 \| 0.198 \| 166 \| 98 \| 9 \| 6 \| 146 \| 2 \| 157 \| PROTEIN (T4 LYSOZYME) \| PROTEIN (T4 LYSOZYME); pdb \| 1SSW\_A \| 1.0 \| 0.005624 \| 122 \| 0.194 \| 175 \| 100 \| 11 \| 6 \| 153 \| 2 \| 162 \| Lysozyme \| Lysozyme; pdb \| 6K69\_B \| 1.0 \| 0.005334 \| 122 \| 0.188 \| 180 \| 100 \| 8 \| 6 \| 150 \| 2 \| 170 \| Engineered T4 lysozyme \| Engineered T4 lysozyme; pdb \| 1JTN\_A \| 1.0 \| 0.002678 \| 122 \| 0.198 \| 176 \| 97 \| 12 \| 10 \| 155 \| 3 \| 164 \| LYSOZYME \| LYSOZYME; pdb \| 6QAJ\_A \| 1.0 \| 0.002678 \| 122 \| 0.206 \| 179 \| 105 \| 11 \| 1 \| 153 \| 2 \| 169 \| Endolysin,Transcription intermediary factor 1-beta \| Endolysin,Transcription intermediary factor 1-beta; pdb \| 235L\_A \| 1.0 \| 0.00455 \| 121 \| 0.203 \| 172 \| 97 \| 10 \| 6 \| 150 \| 2 \| 160 \| T4 LYSOZYME \| T4 LYSOZYME; pdb \| 172L\_A \| 1.0 \| 0.00733 \| 121 \| 0.2 \| 170 \| 100 \| 9 \| 6 \| 150 \| 2 \| 160 \| T4 LYSOZYME \| T4 LYSOZYME; pdb \| 2HUK\_A \| 1.0 \| 0.002824 \| 121 \| 0.201 \| 174 \| 98 \| 12 \| 4 \| 146 \| 1 \| 164 \| Lysozyme \| Lysozyme; pdb \| 2Q9E\_C \| 1.0 \| 0.00314 \| 121 \| 0.191 \| 172 \| 99 \| 11 \| 6 \| 146 \| 2 \| 164 \| Lysozyme \| Lysozyme; pdb \| 6XYR\_A \| 1.0 \| 0.006952 \| 121 \| 0.203 \| 177 \| 101 \| 10 \| 6 \| 155 \| 34 \| 197 \| T4Lnano,Endolysin,Calmodulin,Endolysin,Calmodulin-1 \| T4Lnano,Endolysin,Calmodulin,Endolysin,Calmodulin-1; pdb \| 6LB8\_C \| 1.0 \| 0.00906 \| 121 \| 0.201 \| 174 \| 104 \| 11 \| 6 \| 155 \| 2 \| 164 \| Endolysin,Calcium uptake protein 1, mitochondrial \| Endolysin,Calcium uptake protein 1, mitochondrial; pdb \| 141L\_A \| 1.0 \| 0.005624 \| 120 \| 0.191 \| 172 \| 99 \| 9 \| 6 \| 150 \| 2 \| 160 \| T4 LYSOZYME \| T4 LYSOZYME; pdb \| 1OVK\_A \| 1.0 \| 0.006952 \| 120 \| 0.203 \| 172 \| 97 \| 11 \| 6 \| 150 \| 2 \| 160 \| Lysozyme \| Lysozyme; pdb \| 1L07\_A \| 1.0 \| 0.004315 \| 120 \| 0.198 \| 166 \| 98 \| 8 \| 6 \| 146 \| 2 \| 157 \| T4 LYSOZYME \| T4 LYSOZYME; pdb \| 1L15\_A \| 1.0 \| 0.003881 \| 120 \| 0.204 \| 166 \| 97 \| 9 \| 6 \| 146 \| 2 \| 157 \| T4 LYSOZYME \| T4 LYSOZYME; pdb \| 1K28\_A \| 1.0 \| 0.007729 \| 120 \| 0.193 \| 176 \| 102 \| 9 \| 4 \| 150 \| 169 \| 333 \| TAIL-ASSOCIATED LYSOZYME \| TAIL-ASSOCIATED LYSOZYME; pdb \| 244L\_A \| 1.0 \| 0.006593 \| 119 \| 0.203 \| 172 \| 97 \| 10 \| 6 \| 150 \| 2 \| 160 \| T4 LYSOZYME \| T4 LYSOZYME; pdb \| 157L\_A \| 1.0 \| 0.006253 \| 119 \| 0.197 \| 172 \| 98 \| 10 \| 6 \| 150 \| 2 \| 160 \| T4 LYSOZYME \| T4 LYSOZYME; pdb \| 1L01\_A \| 1.0 \| 0.001848 \| 119 \| 0.207 \| 169 \| 100 \| 8 \| 6 \| 146 \| 2 \| 164 \| T4 LYSOZYME \| T4 LYSOZYME; pdb \| 1L33\_A \| 1.0 \| 0.00455 \| 119 \| 0.214 \| 168 \| 93 \| 11 \| 6 \| 146 \| 2 \| 157 \| T4 LYSOZYME \| T4 LYSOZYME; pdb \| 1L50\_A \| 1.0 \| 0.005059 \| 119 \| 0.198 \| 166 \| 105 \| 9 \| 6 \| 146 \| 2 \| 164 \| T4 LYSOZYME \| T4 LYSOZYME; pdb \| 1KNI\_A \| 1.0 \| 0.001753 \| 118 \| 0.21 \| 166 \| 97 \| 9 \| 6 \| 143 \| 2 \| 161 \| LYSOZYME \| LYSOZYME; pdb \| 1L02\_A \| 1.0 \| 0.005624 \| 118 \| 0.198 \| 166 \| 98 \| 8 \| 6 \| 146 \| 2 \| 157 \| T4 LYSOZYME \| T4 LYSOZYME; pdb \| 2HUL\_A \| 1.0 \| 0.002678 \| 117 \| 0.216 \| 162 \| 91 \| 10 \| 6 \| 140 \| 2 \| 154 \| Lysozyme \| Lysozyme; pdb \| 1L12\_A \| 1.0 \| 0.00906 \| 117 \| 0.2 \| 175 \| 99 \| 10 \| 6 \| 153 \| 2 \| 162 \| T4 LYSOZYME \| T4 LYSOZYME; pdb \| 1L16\_A \| 1.0 \| 0.005334 \| 117 \| 0.198 \| 166 \| 105 \| 8 \| 6 \| 146 \| 2 \| 164 \| T4 LYSOZYME \| T4 LYSOZYME; pdb \| 5B2G\_G \| 1.0 \| 0.003881 \| 117 \| 0.207 \| 169 \| 106 \| 6 \| 4 \| 147 \| 6 \| 171 \| Endolysin,Claudin-4 \| Endolysin,Claudin-4; pdb \| 3OE8\_C \| 1.0 \| 0.001577 \| 117 \| 0.23 \| 182 \| 104 \| 11 \| 5 \| 156 \| 190 \| 365 \| C-X-C chemokine receptor type 4, Lysozyme Chimera \| C-X-C chemokine receptor type 4, Lysozyme Chimera; pdb \| 3OE0\_A \| 1.0 \| 0.004315 \| 117 \| 0.208 \| 163 \| 91 \| 10 \| 6 \| 140 \| 201 \| 353 \| C-X-C chemokine receptor type 4, Lysozyme Chimera \| C-X-C chemokine receptor type 4, Lysozyme Chimera; pdb \| 112L\_A \| 1.0 \| 0.008149 \| 116 \| 0.205 \| 170 \| 99 \| 10 \| 6 \| 150 \| 2 \| 160 \| T4 LYSOZYME \| T4 LYSOZYME; pdb \| 191L\_A \| 1.0 \| 0.003311 \| 116 \| 0.192 \| 166 \| 99 \| 9 \| 6 \| 146 \| 2 \| 157 \| LYSOZYME \| LYSOZYME; pdb \| 3GUK\_A \| 1.0 \| 0.005624 \| 116 \| 0.194 \| 170 \| 101 \| 9 \| 6 \| 150 \| 2 \| 160 \| Lysozyme \| Lysozyme; pdb \| 253L\_A \| 1.0 \| 0.002409 \| 116 \| 0.213 \| 169 \| 99 \| 10 \| 6 \| 146 \| 2 \| 164 \| LYSOZYME \| LYSOZYME; pdb \| 1L13\_A \| 1.0 \| 0.002678 \| 116 \| 0.201 \| 169 \| 101 \| 8 \| 6 \| 146 \| 2 \| 164 \| T4 LYSOZYME \| T4 LYSOZYME; pdb \| 4DAJ\_D \| 1.0 \| 0.003681 \| 116 \| 0.208 \| 168 \| 94 \| 11 \| 6 \| 146 \| 197 \| 352 \| Muscarinic acetylcholine receptor M3, Lysozyme \| Muscarinic acetylcholine receptor M3, Lysozyme; pdb \| 250L\_A \| 1.0 \| 0.00593 \| 114 \| 0.214 \| 168 \| 93 \| 11 \| 6 \| 146 \| 2 \| 157 \| T4 LYSOZYME \| T4 LYSOZYME; pdb \| 1L57\_A \| 1.0 \| 0.006952 \| 114 \| 0.204 \| 166 \| 97 \| 9 \| 6 \| 146 \| 2 \| 157 \| LYSOZYME \| LYSOZYME; pdb \| 1L06\_A \| 1.0 \| 0.003681 \| 114 \| 0.213 \| 169 \| 99 \| 9 \| 6 \| 146 \| 2 \| 164 \| T4 LYSOZYME \| T4 LYSOZYME; pdb \| 1L10\_A \| 1.0 \| 0.008149 \| 113 \| 0.198 \| 166 \| 98 \| 8 \| 6 \| 146 \| 2 \| 157 \| T4 LYSOZYME \| T4 LYSOZYME; pdb \| 4YXA\_C \| 1.0 \| 0.00593 \| 113 \| 0.18 \| 155 \| 91 \| 7 \| 7 \| 150 \| 36 \| 165 \| Oxygen-regulated invasion protein OrgB,Endolysin \| Oxygen-regulated invasion protein OrgB,Endolysin; pdb \| 5BZ6\_A \| 1.0 \| 0.008593 \| 113 \| 0.212 \| 165 \| 90 \| 11 \| 6 \| 143 \| 2 \| 153 \| Lysozyme,Calcium uniporter protein, mitochondrial \| Lysozyme,Calcium uniporter protein, mitochondrial; pdb \| 4XSJ\_A \| 1.0 \| 0.005334 \| 112 \| 0.196 \| 168 \| 96 \| 11 \| 6 \| 146 \| 2 \| 157 \| Lysozyme,Calcium uniporter protein, mitochondrial \| Lysozyme,Calcium uniporter protein, mitochondrial; pdb \| 6A9J\_B \| 1.0 \| 0.008593 \| 112 \| 0.201 \| 174 \| 100 \| 10 \| 6 \| 151 \| 2 \| 164 \| Endolysin,Autophagy-related protein 2 \| Endolysin,Autophagy-related protein 2; pdb \| 6KK7\_A \| 1.0 \| 0.004092 \| 112 \| 0.153 \| 195 \| 111 \| 12 \| 6 \| 156 \| 98 \| 282 \| Glucagon-like peptide 1 receptor,Endolysin,Glucagon-like peptide 1 receptor \| Glucagon-like peptide 1 receptor,Endolysin,Glucagon-like peptide 1 receptor; pdb \| 3OE8\_B \| 1.0 \| 0.00455 \| 112 \| 0.198 \| 186 \| 107 \| 11 \| 5 \| 157 \| 194 \| 370 \| C-X-C chemokine receptor type 4, Lysozyme Chimera \| C-X-C chemokine receptor type 4, Lysozyme Chimera; pdb \| 5I0N\_A \| 1.0 \| 0.001753 \| 112 \| 0.198 \| 176 \| 102 \| 11 \| 4 \| 146 \| 70 \| 239 \| Phosphatidylinositol 4-kinase type 2-alpha,Lysozyme,Phosphatidylinositol 4-kinase type 2-alpha \| Phosphatidylinositol 4-kinase type 2-alpha,Lysozyme,Phosphatidylinositol 4-kinase type 2-alpha; pdb \| 6KK1\_A \| 1.0 \| 0.004797 \| 111 \| 0.173 \| 196 \| 106 \| 10 \| 6 \| 156 \| 98 \| 282 \| Glucagon-like peptide 1 receptor,Endolysin,Glucagon-like peptide 1 receptor \| Glucagon-like peptide 1 receptor,Endolysin,Glucagon-like peptide 1 receptor; pdb \| 5T1A\_A \| 1.0 \| 0.00593 \| 110 \| 0.197 \| 167 \| 95 \| 11 \| 4 \| 140 \| 193 \| 350 \| Chimera protein of CC chemokine receptor type 2 isoform B and T4-lysozyme,Lysozyme \| Chimera protein of CC chemokine receptor type 2 isoform B and T4-lysozyme,Lysozyme; pdb \| 150L\_C \| 1.0 \| 0.006593 \| 109 \| 0.222 \| 157 \| 88 \| 9 \| 8 \| 136 \| 4 \| 154 \| T4 LYSOZYME \| T4 LYSOZYME; pdb \| 3GUJ\_A \| 1.0 \| 0.00593 \| 108 \| 0.194 \| 170 \| 94 \| 11 \| 6 \| 146 \| 2 \| 157 \| Lysozyme \| Lysozyme; pdb \| 6KVX\_A \| 1.0 \| 0.005059 \| 108 \| 0.201 \| 159 \| 92 \| 10 \| 10 \| 140 \| 3 \| 154 \| Endolysin,Calcium uniporter protein, mitochondrial \| Endolysin,Calcium uniporter protein, mitochondrial; pdb \| 6KK7\_B \| 1.0 \| 0.004315 \| 108 \| 0.158 \| 196 \| 109 \| 10 \| 6 \| 156 \| 97 \| 281 \| Glucagon-like peptide 1 receptor,Endolysin,Glucagon-like peptide 1 receptor \| Glucagon-like peptide 1 receptor,Endolysin,Glucagon-like peptide 1 receptor; pdb \| 6KJV\_A \| 1.0 \| 0.00455 \| 108 \| 0.175 \| 194 \| 108 \| 12 \| 6 \| 156 \| 98 \| 282 \| Glucagon-like peptide 1 receptor,Endolysin,Glucagon-like peptide 1 receptor \| Glucagon-like peptide 1 receptor,Endolysin,Glucagon-like peptide 1 receptor; pdb \| 6KK1\_B \| 1.0 \| 0.004092 \| 107 \| 0.182 \| 203 \| 106 \| 11 \| 6 \| 157 \| 100 \| 293 \| Glucagon-like peptide 1 receptor,Endolysin,Glucagon-like peptide 1 receptor \| Glucagon-like peptide 1 receptor,Endolysin,Glucagon-like peptide 1 receptor; pdb \| 5GLH\_A \| 1.0 \| 0.002978 \| 107 \| 0.209 \| 186 \| 94 \| 9 \| 6 \| 144 \| 202 \| 381 \| Endothelin Receptor Subtype-B \| Endothelin Receptor Subtype-B; pdb \| 4Z9G\_B \| 1.0 \| 0.004092 \| 106 \| 0.213 \| 192 \| 101 \| 12 \| 6 \| 156 \| 101 \| 283 \| Corticotropin-releasing factor receptor 1,Lysozyme,Corticotropin-releasing factor receptor 1 \| Corticotropin-releasing factor receptor 1,Lysozyme,Corticotropin-releasing factor receptor 1; pdb \| 5VEX\_A \| 1.0 \| 0.00733 \| 106 \| 0.166 \| 198 \| 105 \| 11 \| 6 \| 156 \| 98 \| 282 \| Glucagon-like peptide 1 receptor, Endolysin chimera \| Glucagon-like peptide 1 receptor, Endolysin chimera; pdb \| 6KJV\_B \| 1.0 \| 0.004315 \| 105 \| 0.17 \| 194 \| 109 \| 12 \| 6 \| 156 \| 100 \| 284 \| Glucagon-like peptide 1 receptor,Endolysin,Glucagon-like peptide 1 receptor \| Glucagon-like peptide 1 receptor,Endolysin,Glucagon-like peptide 1 receptor; pdb \| 7JOZ\_R \| 1.0 \| 0.008149 \| 101 \| 0.203 \| 157 \| 91 \| 7 \| 8 \| 136 \| 2 \| 152 \| Endolysin,D(1A) dopamine receptor \| Endolysin,D(1A) dopamine receptor; pdb \| 6LB8\_A \| 0.999 \| 0.009553 \| 97 \| 0.186 \| 172 \| 102 \| 9 \| 6 \| 153 \| 2 \| 159 \| Endolysin,Calcium uptake protein 1, mitochondrial \| Endolysin,Calcium uptake protein 1, mitochondrial; afdb-proteome \| AF-Q8ZQ99-F1-MODEL\_V4 \| 1.0 \| 2.577e-15 \| 693 \| 0.567 \| 148 \| 60 \| 1 \| 1 \| 144 \| 2 \| 149 \| Lysozyme \| Lysozyme; afdb-proteome \| AF-Q8ZMY9-F1-MODEL\_V4 \| 1.0 \| 1.164e-15 \| 676 \| 0.554 \| 148 \| 62 \| 2 \| 1 \| 144 \| 2 \| 149 \| Lysozyme \| Lysozyme; afdb-proteome \| AF-P78285-F1-MODEL\_V4 \| 1.0 \| 1.901e-09 \| 404 \| 0.319 \| 144 \| 90 \| 3 \| 6 \| 144 \| 21 \| 161 \| Lysozyme RrrD \| Lysozyme RrrD; afdb-proteome \| AF-A0A0H3GKK1-F1-MODEL\_V4 \| 1.0 \| 6.1e-09 \| 399 \| 0.34 \| 144 \| 91 \| 3 \| 1 \| 144 \| 27 \| 166 \| Lysozyme \| Lysozyme; afdb-proteome \| AF-Q8ZMT7-F1-MODEL\_V4 \| 1.0 \| 7.54e-09 \| 395 \| 0.368 \| 144 \| 87 \| 3 \| 1 \| 144 \| 27 \| 166 \| Lysozyme \| Lysozyme; afdb-proteome \| AF-A0A175VZF5-F1-MODEL\_V4 \| 1.0 \| 1.281e-08 \| 359 \| 0.317 \| 151 \| 91 \| 5 \| 1 \| 142 \| 107 \| 254 \| Lysozyme \| Lysozyme; afdb-proteome \| AF-C0NHY3-F1-MODEL\_V4 \| 1.0 \| 6.979e-08 \| 331 \| 0.331 \| 151 \| 87 \| 6 \| 1 \| 142 \| 31 \| 176 \| Lysozyme \| Lysozyme; afdb-proteome \| AF-Q32GL6-F1-MODEL\_V4 \| 1.0 \| 2.918e-07 \| 303 \| 0.283 \| 155 \| 95 \| 6 \| 4 \| 145 \| 22 \| 173 \| Lysozyme \| Lysozyme; afdb-proteome \| AF-A0A0H3GP93-F1-MODEL\_V4 \| 1.0 \| 4.701e-07 \| 294 \| 0.301 \| 156 \| 92 \| 6 \| 4 \| 145 \| 20 \| 172 \| Lysozyme \| Lysozyme; afdb-proteome \| AF-A0A0H3GWD4-F1-MODEL\_V4 \| 1.0 \| 6.461e-07 \| 290 \| 0.307 \| 156 \| 91 \| 6 \| 4 \| 145 \| 20 \| 172 \| Lysozyme \| Lysozyme; afdb-proteome \| AF-P76159-F1-MODEL\_V4 \| 1.0 \| 9.872e-07 \| 283 \| 0.294 \| 153 \| 92 \| 6 \| 4 \| 143 \| 22 \| 171 \| Probable prophage lysozyme \| Probable prophage lysozyme; afdb-proteome \| AF-Q8ZLC6-F1-MODEL\_V4 \| 1.0 \| 0.001828 \| 150 \| 0.24 \| 108 \| 74 \| 3 \| 3 \| 110 \| 8 \| 107 \| Putative phage endolysin \| Putative phage endolysin; afdb-uniprot50 \| AF-A0A1S2UDK3-F1-MODEL\_V4 \| 1.0 \| 1.808e-21 \| 1036 \| 0.679 \| 159 \| 51 \| 0 \| 1 \| 159 \| 1 \| 159 \| Lysozyme \| Lysozyme; afdb-uniprot50 \| AF-A0A090V1M7-F1-MODEL\_V4 \| 1.0 \| 1.394e-19 \| 934 \| 0.554 \| 184 \| 57 \| 2 \| 1 \| 159 \| 1 \| 184 \| Lysozyme \| Lysozyme; afdb-uniprot50 \| AF-A0A857JAQ9-F1-MODEL\_V4 \| 1.0 \| 1.441e-16 \| 785 \| 0.519 \| 156 \| 74 \| 1 \| 1 \| 155 \| 1 \| 156 \| Glycoside hydrolase family protein \| Glycoside hydrolase family protein; afdb-uniprot50 \| AF-A0A6G6JNR3-F1-MODEL\_V4 \| 1.0 \| 7.236e-17 \| 783 \| 0.543 \| 171 \| 66 \| 1 \| 1 \| 159 \| 1 \| 171 \| Lysozyme \| Lysozyme; afdb-uniprot50 \| AF-A0A3G6HI33-F1-MODEL\_V4 \| 1.0 \| 4.384e-16 \| 758 \| 0.509 \| 159 \| 75 \| 2 \| 2 \| 159 \| 36 \| 192 \| Lysozyme \| Lysozyme; afdb-uniprot50 \| AF-A0A844U3X0-F1-MODEL\_V4 \| 1.0 \| 1.079e-15 \| 742 \| 0.493 \| 160 \| 77 \| 3 \| 1 \| 159 \| 1 \| 157 \| Glycoside hydrolase family protein \| Glycoside hydrolase family protein; afdb-uniprot50 \| AF-A0A1W0CID5-F1-MODEL\_V4 \| 1.0 \| 3.849e-15 \| 708 \| 0.595 \| 146 \| 56 \| 2 \| 1 \| 146 \| 1 \| 143 \| Lysozyme \| Lysozyme; afdb-uniprot50 \| AF-A0A286B0J8-F1-MODEL\_V4 \| 1.0 \| 6.2e-15 \| 696 \| 0.451 \| 184 \| 74 \| 3 \| 1 \| 157 \| 5 \| 188 \| Lysozyme \| Lysozyme; afdb-uniprot50 \| AF-A0A2G1LY75-F1-MODEL\_V4 \| 1.0 \| 1.111e-14 \| 686 \| 0.565 \| 145 \| 62 \| 1 \| 2 \| 145 \| 38 \| 182 \| Lysozyme \| Lysozyme; afdb-uniprot50 \| AF-A0A7T2GKL9-F1-MODEL\_V4 \| 1.0 \| 6.893e-15 \| 681 \| 0.627 \| 145 \| 53 \| 1 \| 1 \| 145 \| 42 \| 185 \| Lysozyme \| Lysozyme; afdb-uniprot50 \| AF-A0A8B1MMR9-F1-MODEL\_V4 \| 1.0 \| 2.211e-14 \| 676 \| 0.54 \| 148 \| 64 \| 1 \| 1 \| 144 \| 1 \| 148 \| Lysozyme RrrD \| Lysozyme RrrD; afdb-uniprot50 \| AF-A0A2W7M8R8-F1-MODEL\_V4 \| 1.0 \| 1.886e-14 \| 675 \| 0.53 \| 147 \| 66 \| 2 \| 1 \| 147 \| 1 \| 144 \| Lysozyme \| Lysozyme; afdb-uniprot50 \| AF-A0A2W4VX53-F1-MODEL\_V4 \| 1.0 \| 1.789e-14 \| 666 \| 0.49 \| 153 \| 74 \| 1 \| 3 \| 151 \| 1 \| 153 \| Lysozyme \| Lysozyme; afdb-uniprot50 \| AF-A0A4R3ZEW6-F1-MODEL\_V4 \| 1.0 \| 5.443e-14 \| 661 \| 0.533 \| 148 \| 65 \| 1 \| 1 \| 144 \| 1 \| 148 \| Lysozyme \| Lysozyme; afdb-uniprot50 \| AF-A0A7Z0MUC7-F1-MODEL\_V4 \| 1.0 \| 7.481e-14 \| 661 \| 0.452 \| 159 \| 84 \| 2 \| 2 \| 159 \| 34 \| 190 \| Lysozyme \| Lysozyme; afdb-uniprot50 \| AF-A0A841WQ48-F1-MODEL\_V4 \| 1.0 \| 2.332e-14 \| 656 \| 0.456 \| 151 \| 78 \| 3 \| 1 \| 151 \| 4 \| 150 \| Glycoside hydrolase family protein \| Glycoside hydrolase family protein; afdb-uniprot50 \| AF-A0A4Q0YFE7-F1-MODEL\_V4 \| 1.0 \| 4.404e-14 \| 653 \| 0.469 \| 147 \| 72 \| 3 \| 1 \| 144 \| 3 \| 146 \| Lysozyme \| Lysozyme; afdb-uniprot50 \| AF-A0A747EZ57-F1-MODEL\_V4 \| 1.0 \| 1.989e-14 \| 648 \| 0.543 \| 151 \| 65 \| 2 \| 1 \| 147 \| 2 \| 152 \| Lysozyme \| Lysozyme; afdb-uniprot50 \| AF-W1INZ3-F1-MODEL\_V4 \| 1.0 \| 8.317e-14 \| 642 \| 0.5 \| 144 \| 71 \| 1 \| 1 \| 144 \| 1 \| 143 \| Lysozyme \| Lysozyme; afdb-uniprot50 \| AF-A0A0J1IN41-F1-MODEL\_V4 \| 1.0 \| 9.247e-14 \| 642 \| 0.537 \| 147 \| 67 \| 1 \| 1 \| 147 \| 1 \| 146 \| Lysozyme \| Lysozyme; afdb-uniprot50 \| AF-A0A2D8XAB4-F1-MODEL\_V4 \| 1.0 \| 7.481e-14 \| 634 \| 0.486 \| 150 \| 73 \| 3 \| 1 \| 150 \| 1 \| 146 \| Lysozyme \| Lysozyme; afdb-uniprot50 \| AF-A0A4P5ZKT6-F1-MODEL\_V4 \| 1.0 \| 1.028e-13 \| 629 \| 0.506 \| 150 \| 70 \| 2 \| 1 \| 150 \| 67 \| 212 \| Lysozyme \| Lysozyme; afdb-uniprot50 \| AF-A0A2X0S8V7-F1-MODEL\_V4 \| 1.0 \| 1.656e-13 \| 625 \| 0.516 \| 149 \| 71 \| 1 \| 1 \| 149 \| 1 \| 148 \| Lysozyme \| Lysozyme; afdb-uniprot50 \| AF-A0A3P1XHR8-F1-MODEL\_V4 \| 1.0 \| 1.34e-13 \| 624 \| 0.523 \| 147 \| 67 \| 2 \| 1 \| 147 \| 350 \| 493 \| Lysozyme \| Lysozyme; afdb-uniprot50 \| AF-A0A846A290-F1-MODEL\_V4 \| 1.0 \| 1.656e-13 \| 622 \| 0.473 \| 150 \| 75 \| 3 \| 2 \| 151 \| 81 \| 226 \| Lysozyme \| Lysozyme; afdb-uniprot50 \| AF-A0A7U2UZ55-F1-MODEL\_V4 \| 1.0 \| 1.34e-13 \| 619 \| 0.496 \| 147 \| 70 \| 1 \| 1 \| 143 \| 1 \| 147 \| Lysozyme \| Lysozyme; afdb-uniprot50 \| AF-A0A4Q7YUD0-F1-MODEL\_V4 \| 1.0 \| 1.571e-13 \| 616 \| 0.51 \| 143 \| 67 \| 2 \| 1 \| 143 \| 4 \| 143 \| Lysozyme \| Lysozyme; afdb-uniprot50 \| AF-A0A3M9Z9E8-F1-MODEL\_V4 \| 1.0 \| 4.298e-13 \| 615 \| 0.496 \| 145 \| 70 \| 2 \| 2 \| 146 \| 29 \| 170 \| Lysozyme \| Lysozyme; afdb-uniprot50 \| AF-A0A5J6N4Q1-F1-MODEL\_V4 \| 1.0 \| 3.667e-13 \| 614 \| 0.479 \| 144 \| 72 \| 2 \| 1 \| 144 \| 1 \| 141 \| Lysozyme \| Lysozyme; afdb-uniprot50 \| AF-A0A6I5P031-F1-MODEL\_V4 \| 1.0 \| 3.298e-13 \| 612 \| 0.483 \| 151 \| 74 \| 3 \| 1 \| 151 \| 30 \| 176 \| Lysozyme \| Lysozyme; afdb-uniprot50 \| AF-A0A2V9UU03-F1-MODEL\_V4 \| 1.0 \| 4.298e-13 \| 605 \| 0.486 \| 146 \| 72 \| 2 \| 1 \| 146 \| 7 \| 149 \| Lysozyme \| Lysozyme; afdb-uniprot50 \| AF-A0A1S1NZ32-F1-MODEL\_V4 \| 1.0 \| 7.302e-13 \| 605 \| 0.503 \| 143 \| 67 \| 3 \| 1 \| 143 \| 1 \| 139 \| Lysozyme \| Lysozyme; afdb-uniprot50 \| AF-A0A3M9ZBC4-F1-MODEL\_V4 \| 1.0 \| 3.866e-13 \| 605 \| 0.526 \| 150 \| 67 \| 3 \| 2 \| 151 \| 112 \| 257 \| Lysozyme \| Lysozyme; afdb-uniprot50 \| AF-A0A7D0IRK1-F1-MODEL\_V4 \| 1.0 \| 5.602e-13 \| 603 \| 0.479 \| 144 \| 75 \| 0 \| 1 \| 144 \| 1 \| 144 \| Lysozyme \| Lysozyme; afdb-uniprot50 \| AF-A0A7T9EKQ7-F1-MODEL\_V4 \| 1.0 \| 8.56e-13 \| 603 \| 0.51 \| 145 \| 68 \| 2 \| 1 \| 145 \| 8 \| 149 \| Lysozyme \| Lysozyme; afdb-uniprot50 \| AF-A0A2T1DR14-F1-MODEL\_V4 \| 1.0 \| 4.298e-13 \| 602 \| 0.486 \| 150 \| 73 \| 3 \| 2 \| 151 \| 8 \| 153 \| Lysozyme \| Lysozyme; afdb-uniprot50 \| AF-A0A4S3LZ40-F1-MODEL\_V4 \| 1.0 \| 5.313e-13 \| 601 \| 0.486 \| 146 \| 71 \| 3 \| 1 \| 146 \| 1 \| 142 \| Lysozyme \| Lysozyme; afdb-uniprot50 \| AF-A0A0Q2RDV9-F1-MODEL\_V4 \| 1.0 \| 1.841e-13 \| 601 \| 0.446 \| 159 \| 83 \| 4 \| 2 \| 158 \| 3 \| 158 \| Lysozyme \| Lysozyme; afdb-uniprot50 \| AF-A0A5E8KST5-F1-MODEL\_V4 \| 1.0 \| 7.302e-13 \| 600 \| 0.466 \| 148 \| 78 \| 1 \| 1 \| 148 \| 20 \| 166 \| Lysozyme \| Lysozyme; afdb-uniprot50 \| AF-A0A239PTC6-F1-MODEL\_V4 \| 1.0 \| 3.667e-13 \| 600 \| 0.52 \| 148 \| 64 \| 4 \| 1 \| 144 \| 1 \| 145 \| Lysozyme \| Lysozyme; afdb-uniprot50 \| AF-A0A348HCF7-F1-MODEL\_V4 \| 1.0 \| 9.517e-13 \| 600 \| 0.468 \| 145 \| 76 \| 1 \| 1 \| 144 \| 2 \| 146 \| Lysozyme \| Lysozyme; afdb-uniprot50 \| AF-A0A7Y6TTM5-F1-MODEL\_V4 \| 1.0 \| 2.4e-13 \| 599 \| 0.44 \| 175 \| 69 \| 2 \| 1 \| 146 \| 1 \| 175 \| Lysozyme \| Lysozyme; afdb-uniprot50 \| AF-S3JZ44-F1-MODEL\_V4 \| 1.0 \| 1.058e-12 \| 598 \| 0.468 \| 145 \| 73 \| 3 \| 1 \| 145 \| 1 \| 141 \| Lysozyme \| Lysozyme; afdb-uniprot50 \| AF-A0A5E4NVC9-F1-MODEL\_V4 \| 1.0 \| 6.925e-13 \| 594 \| 0.459 \| 148 \| 76 \| 3 \| 1 \| 148 \| 3 \| 146 \| Lysozyme \| Lysozyme; afdb-uniprot50 \| AF-A0A2E4ZEW9-F1-MODEL\_V4 \| 1.0 \| 3.866e-13 \| 593 \| 0.454 \| 154 \| 79 \| 4 \| 1 \| 153 \| 1 \| 150 \| Lysozyme \| Lysozyme; afdb-uniprot50 \| AF-A0A381SG74-F1-MODEL\_V4 \| 1.0 \| 5.602e-13 \| 593 \| 0.457 \| 153 \| 79 \| 3 \| 1 \| 153 \| 22 \| 170 \| Lysozyme \| Lysozyme; afdb-uniprot50 \| AF-A0A7H0LF26-F1-MODEL\_V4 \| 1.0 \| 6.229e-13 \| 592 \| 0.472 \| 146 \| 74 \| 2 \| 2 \| 146 \| 49 \| 192 \| Lysozyme \| Lysozyme; afdb-uniprot50 \| AF-A0A1C4GBY3-F1-MODEL\_V4 \| 1.0 \| 1.797e-12 \| 590 \| 0.513 \| 148 \| 65 \| 3 \| 1 \| 144 \| 1 \| 145 \| Lysozyme \| Lysozyme; afdb-uniprot50 \| AF-A0A3N8QQG0-F1-MODEL\_V4 \| 1.0 \| 4.532e-13 \| 587 \| 0.421 \| 173 \| 71 \| 1 \| 1 \| 144 \| 3 \| 175 \| Lysozyme \| Lysozyme; afdb-uniprot50 \| AF-A0A158DN86-F1-MODEL\_V4 \| 1.0 \| 1.24e-12 \| 585 \| 0.444 \| 144 \| 80 \| 0 \| 1 \| 144 \| 1 \| 144 \| Lysozyme \| Lysozyme; afdb-uniprot50 \| AF-J2HCF9-F1-MODEL\_V4 \| 1.0 \| 7.302e-13 \| 584 \| 0.486 \| 148 \| 71 \| 3 \| 1 \| 145 \| 1 \| 146 \| Lysozyme \| Lysozyme; afdb-uniprot50 \| AF-A0A846FGE4-F1-MODEL\_V4 \| 1.0 \| 1.454e-12 \| 583 \| 0.49 \| 151 \| 73 \| 3 \| 1 \| 151 \| 53 \| 199 \| Lysozyme \| Lysozyme; afdb-uniprot50 \| AF-A0A6I2IUJ3-F1-MODEL\_V4 \| 1.0 \| 6.568e-13 \| 581 \| 0.455 \| 156 \| 80 \| 4 \| 2 \| 155 \| 3 \| 155 \| Lysozyme \| Lysozyme; afdb-uniprot50 \| AF-A0A494X2V0-F1-MODEL\_V4 \| 1.0 \| 4.779e-13 \| 581 \| 0.434 \| 182 \| 72 \| 2 \| 2 \| 153 \| 6 \| 186 \| Lysozyme \| Lysozyme; afdb-uniprot50 \| AF-A0A3B9WX79-F1-MODEL\_V4 \| 1.0 \| 8.118e-13 \| 580 \| 0.464 \| 153 \| 75 \| 3 \| 1 \| 150 \| 1 \| 149 \| Lysozyme \| Lysozyme; afdb-uniprot50 \| AF-A0A7W7K3I1-F1-MODEL\_V4 \| 1.0 \| 1.176e-12 \| 580 \| 0.49 \| 151 \| 69 \| 2 \| 1 \| 143 \| 1 \| 151 \| Lysozyme \| Lysozyme; afdb-uniprot50 \| AF-A0A4S1WM04-F1-MODEL\_V4 \| 1.0 \| 1.705e-12 \| 579 \| 0.526 \| 152 \| 64 \| 2 \| 1 \| 144 \| 24 \| 175 \| Lysozyme \| Lysozyme; afdb-uniprot50 \| AF-A0A2G2MWX1-F1-MODEL\_V4 \| 1.0 \| 4.532e-13 \| 579 \| 0.489 \| 147 \| 68 \| 4 \| 1 \| 143 \| 1 \| 144 \| Lysozyme \| Lysozyme; afdb-uniprot50 \| AF-A0A2A2JXU9-F1-MODEL\_V4 \| 1.0 \| 8.56e-13 \| 578 \| 0.506 \| 144 \| 67 \| 3 \| 1 \| 144 \| 1 \| 140 \| Lysozyme \| Lysozyme; afdb-uniprot50 \| AF-A0A5C7QC42-F1-MODEL\_V4 \| 1.0 \| 1.454e-12 \| 575 \| 0.493 \| 146 \| 70 \| 3 \| 1 \| 145 \| 1 \| 143 \| Lysozyme \| Lysozyme; afdb-uniprot50 \| AF-A0A3S0BYI5-F1-MODEL\_V4 \| 1.0 \| 2.107e-12 \| 575 \| 0.482 \| 143 \| 70 \| 2 \| 1 \| 143 \| 4 \| 142 \| Lysozyme \| Lysozyme; afdb-uniprot50 \| AF-A0A1M6MED4-F1-MODEL\_V4 \| 1.0 \| 6.41e-12 \| 575 \| 0.486 \| 144 \| 73 \| 1 \| 1 \| 144 \| 3 \| 145 \| Lysozyme \| Lysozyme; afdb-uniprot50 \| AF-A0A0K1NGA9-F1-MODEL\_V4 \| 1.0 \| 3.219e-12 \| 573 \| 0.468 \| 143 \| 73 \| 2 \| 1 \| 143 \| 1 \| 140 \| Lysozyme \| Lysozyme; afdb-uniprot50 \| AF-A0A0C1VC21-F1-MODEL\_V4 \| 1.0 \| 2.342e-12 \| 571 \| 0.441 \| 154 \| 79 \| 3 \| 2 \| 151 \| 11 \| 161 \| Lysozyme \| Lysozyme; afdb-uniprot50 \| AF-A0A4Q7EGY7-F1-MODEL\_V4 \| 1.0 \| 2.47e-12 \| 569 \| 0.441 \| 154 \| 79 \| 3 \| 2 \| 151 \| 80 \| 230 \| Lysozyme \| Lysozyme; afdb-uniprot50 \| AF-N8WAZ3-F1-MODEL\_V4 \| 1.0 \| 3.579e-12 \| 568 \| 0.493 \| 148 \| 68 \| 2 \| 1 \| 144 \| 63 \| 207 \| Lysozyme \| Lysozyme; afdb-uniprot50 \| AF-Q9P9P8-F1-MODEL\_V4 \| 1.0 \| 1.176e-12 \| 567 \| 0.465 \| 159 \| 80 \| 4 \| 2 \| 158 \| 41 \| 196 \| Lysozyme \| Lysozyme; afdb-uniprot50 \| AF-A0A349GTM3-F1-MODEL\_V4 \| 1.0 \| 2.342e-12 \| 566 \| 0.442 \| 149 \| 76 \| 4 \| 1 \| 145 \| 1 \| 146 \| Lysozyme \| Lysozyme; afdb-uniprot50 \| AF-A0A6P0MVH8-F1-MODEL\_V4 \| 1.0 \| 1.003e-12 \| 565 \| 0.449 \| 158 \| 73 \| 2 \| 1 \| 147 \| 1 \| 155 \| Lysozyme \| Lysozyme; afdb-uniprot50 \| AF-A0A6B9YSE3-F1-MODEL\_V4 \| 1.0 \| 3.579e-12 \| 565 \| 0.503 \| 143 \| 68 \| 2 \| 1 \| 143 \| 4 \| 143 \| Lysozyme \| Lysozyme; afdb-uniprot50 \| AF-A0A3B8XMC2-F1-MODEL\_V4 \| 1.0 \| 2.895e-12 \| 565 \| 0.413 \| 145 \| 81 \| 2 \| 1 \| 145 \| 3 \| 143 \| Lysozyme \| Lysozyme; afdb-uniprot50 \| AF-A0A653CSV6-F1-MODEL\_V4 \| 1.0 \| 5.766e-12 \| 564 \| 0.5 \| 144 \| 68 \| 2 \| 1 \| 144 \| 112 \| 251 \| Lysozyme \| Lysozyme; afdb-uniprot50 \| AF-B6VNM2-F1-MODEL\_V4 \| 1.0 \| 5.766e-12 \| 563 \| 0.496 \| 143 \| 68 \| 2 \| 1 \| 143 \| 25 \| 163 \| Lysozyme \| Lysozyme; afdb-uniprot50 \| AF-A0A1I0J1Q9-F1-MODEL\_V4 \| 1.0 \| 1.454e-12 \| 563 \| 0.516 \| 149 \| 63 \| 4 \| 1 \| 143 \| 1 \| 146 \| Lysozyme \| Lysozyme; afdb-uniprot50 \| AF-I3YEG4-F1-MODEL\_V4 \| 1.0 \| 5.186e-12 \| 561 \| 0.475 \| 143 \| 71 \| 3 \| 1 \| 143 \| 10 \| 148 \| Lysozyme \| Lysozyme; afdb-uniprot50 \| AF-A0A5A9FA75-F1-MODEL\_V4 \| 1.0 \| 8.118e-13 \| 561 \| 0.44 \| 161 \| 73 \| 1 \| 1 \| 144 \| 2 \| 162 \| Lysozyme \| Lysozyme; afdb-uniprot50 \| AF-A0A1H8EK82-F1-MODEL\_V4 \| 1.0 \| 6.08e-12 \| 559 \| 0.469 \| 147 \| 76 \| 2 \| 1 \| 146 \| 3 \| 148 \| Lysozyme \| Lysozyme; afdb-uniprot50 \| AF-A0A349JKR9-F1-MODEL\_V4 \| 1.0 \| 3.774e-12 \| 558 \| 0.47 \| 151 \| 75 \| 3 \| 3 \| 151 \| 98 \| 245 \| Lysozyme \| Lysozyme; afdb-uniprot50 \| AF-A0A840YK85-F1-MODEL\_V4 \| 1.0 \| 3.979e-12 \| 557 \| 0.476 \| 147 \| 72 \| 4 \| 1 \| 147 \| 46 \| 187 \| Lysozyme \| Lysozyme; afdb-uniprot50 \| AF-A0A4Q6Y9K8-F1-MODEL\_V4 \| 1.0 \| 4.424e-12 \| 555 \| 0.517 \| 141 \| 67 \| 1 \| 4 \| 144 \| 1 \| 140 \| Lysozyme \| Lysozyme; afdb-uniprot50 \| AF-A0A2X1ULF4-F1-MODEL\_V4 \| 1.0 \| 9.517e-13 \| 554 \| 0.477 \| 157 \| 67 \| 5 \| 1 \| 146 \| 5 \| 157 \| Lysozyme \| Lysozyme; afdb-uniprot50 \| AF-A0A840I6B0-F1-MODEL\_V4 \| 1.0 \| 5.186e-12 \| 551 \| 0.466 \| 150 \| 73 \| 4 \| 1 \| 146 \| 1 \| 147 \| Lysozyme \| Lysozyme; afdb-uniprot50 \| AF-A0A1I6KZC5-F1-MODEL\_V4 \| 1.0 \| 5.766e-12 \| 551 \| 0.472 \| 148 \| 75 \| 2 \| 1 \| 148 \| 15 \| 159 \| Lysozyme \| Lysozyme; afdb-uniprot50 \| AF-A0A442TIX8-F1-MODEL\_V4 \| 1.0 \| 3.774e-12 \| 550 \| 0.493 \| 150 \| 69 \| 3 \| 1 \| 146 \| 1 \| 147 \| Lysozyme \| Lysozyme; afdb-uniprot50 \| AF-A0A2U1WSM1-F1-MODEL\_V4 \| 1.0 \| 2.47e-12 \| 549 \| 0.424 \| 158 \| 81 \| 6 \| 1 \| 154 \| 3 \| 154 \| Lysozyme \| Lysozyme; afdb-uniprot50 \| AF-A0A858RDI0-F1-MODEL\_V4 \| 1.0 \| 6.08e-12 \| 549 \| 0.466 \| 150 \| 70 \| 4 \| 1 \| 145 \| 43 \| 187 \| Lysozyme \| Lysozyme; afdb-uniprot50 \| AF-A0A6C8N2Y9-F1-MODEL\_V4 \| 1.0 \| 2.746e-12 \| 549 \| 0.439 \| 148 \| 81 \| 2 \| 1 \| 147 \| 1 \| 147 \| Lysozyme \| Lysozyme; afdb-uniprot50 \| AF-A0A7J6JRX3-F1-MODEL\_V4 \| 1.0 \| 1.211e-11 \| 549 \| 0.514 \| 142 \| 66 \| 1 \| 2 \| 143 \| 6 \| 144 \| Lysozyme \| Lysozyme; afdb-uniprot50 \| AF-T2JMX1-F1-MODEL\_V4 \| 1.0 \| 6.759e-12 \| 549 \| 0.391 \| 148 \| 87 \| 2 \| 1 \| 148 \| 1 \| 145 \| Lysozyme \| Lysozyme; afdb-uniprot50 \| AF-A0A844YLN9-F1-MODEL\_V4 \| 1.0 \| 4.919e-12 \| 548 \| 0.496 \| 149 \| 71 \| 2 \| 3 \| 147 \| 46 \| 194 \| Glycoside hydrolase family protein \| Glycoside hydrolase family protein; afdb-uniprot50 \| AF-A0A7Y3W4Z2-F1-MODEL\_V4 \| 1.0 \| 7.515e-12 \| 548 \| 0.472 \| 148 \| 71 \| 3 \| 1 \| 144 \| 1 \| 145 \| Lysozyme \| Lysozyme; afdb-uniprot50 \| AF-A0A2N3AFJ7-F1-MODEL\_V4 \| 1.0 \| 6.41e-12 \| 546 \| 0.466 \| 148 \| 72 \| 2 \| 2 \| 145 \| 25 \| 169 \| Lysozyme \| Lysozyme; afdb-uniprot50 \| AF-A0A537UHK2-F1-MODEL\_V4 \| 1.0 \| 2.746e-12 \| 546 \| 0.478 \| 165 \| 72 \| 4 \| 1 \| 156 \| 81 \| 240 \| Lysozyme \| Lysozyme; afdb-uniprot50 \| AF-A0A2V2RWC3-F1-MODEL\_V4 \| 1.0 \| 1.617e-12 \| 545 \| 0.475 \| 166 \| 70 \| 5 \| 1 \| 155 \| 5 \| 164 \| Lysozyme \| Lysozyme; afdb-uniprot50 \| AF-A0A7W1RQP8-F1-MODEL\_V4 \| 1.0 \| 6.759e-12 \| 544 \| 0.423 \| 151 \| 77 \| 3 \| 1 \| 146 \| 2 \| 147 \| Lysozyme \| Lysozyme; afdb-uniprot50 \| AF-A0A546YGV8-F1-MODEL\_V4 \| 1.0 \| 7.515e-12 \| 544 \| 0.462 \| 147 \| 71 \| 4 \| 1 \| 143 \| 1 \| 143 \| Lysozyme \| Lysozyme; afdb-uniprot50 \| AF-A3Z6C9-F1-MODEL\_V4 \| 1.0 \| 8.355e-12 \| 544 \| 0.406 \| 150 \| 84 \| 4 \| 1 \| 147 \| 3 \| 150 \| Lysozyme \| Lysozyme; afdb-uniprot50 \| AF-A0A1I6PPK5-F1-MODEL\_V4 \| 1.0 \| 9.289e-12 \| 543 \| 0.496 \| 149 \| 70 \| 2 \| 1 \| 144 \| 43 \| 191 \| Lysozyme \| Lysozyme; afdb-uniprot50 \| AF-L8M8V2-F1-MODEL\_V4 \| 1.0 \| 1.277e-11 \| 543 \| 0.443 \| 142 \| 75 \| 2 \| 3 \| 144 \| 59 \| 196 \| Lysozyme \| Lysozyme; afdb-uniprot50 \| AF-A0A2D9TFC8-F1-MODEL\_V4 \| 1.0 \| 7.515e-12 \| 543 \| 0.469 \| 147 \| 71 \| 4 \| 1 \| 146 \| 1 \| 141 \| Lysozyme \| Lysozyme; afdb-uniprot50 \| AF-A0A0J7Y672-F1-MODEL\_V4 \| 1.0 \| 5.186e-12 \| 543 \| 0.459 \| 148 \| 76 \| 3 \| 2 \| 147 \| 5 \| 150 \| Lysozyme \| Lysozyme; afdb-uniprot50 \| AF-V4R5W7-F1-MODEL\_V4 \| 1.0 \| 6.568e-13 \| 542 \| 0.384 \| 177 \| 84 \| 5 \| 1 \| 155 \| 1 \| 174 \| Lysozyme \| Lysozyme; afdb-uniprot50 \| AF-N9YWK1-F1-MODEL\_V4 \| 1.0 \| 2.286e-11 \| 541 \| 0.513 \| 144 \| 66 \| 3 \| 1 \| 144 \| 340 \| 479 \| Lysozyme \| Lysozyme; afdb-uniprot50 \| AF-A0A504F0U0-F1-MODEL\_V4 \| 1.0 \| 4.424e-12 \| 539 \| 0.452 \| 148 \| 73 \| 5 \| 1 \| 144 \| 3 \| 146 \| Lysozyme \| Lysozyme; afdb-uniprot50 \| AF-A0A1K1QDJ6-F1-MODEL\_V4 \| 1.0 \| 1.754e-11 \| 539 \| 0.46 \| 139 \| 75 \| 0 \| 6 \| 144 \| 6 \| 144 \| Lysozyme \| Lysozyme; afdb-uniprot50 \| AF-A0A4Y9EQV7-F1-MODEL\_V4 \| 1.0 \| 9.794e-12 \| 539 \| 0.465 \| 146 \| 71 \| 3 \| 3 \| 143 \| 2 \| 145 \| Lysozyme \| Lysozyme; afdb-uniprot50 \| AF-M1N4J0-F1-MODEL\_V4 \| 1.0 \| 1.089e-11 \| 539 \| 0.462 \| 147 \| 75 \| 3 \| 1 \| 147 \| 2 \| 144 \| Lysozyme \| Lysozyme; afdb-uniprot50 \| AF-A0A1M3DRS2-F1-MODEL\_V4 \| 1.0 \| 6.759e-12 \| 537 \| 0.391 \| 143 \| 84 \| 2 \| 2 \| 144 \| 3 \| 142 \| Lysozyme \| Lysozyme; afdb-uniprot50 \| AF-A0A4P6L9W1-F1-MODEL\_V4 \| 1.0 \| 4.919e-12 \| 537 \| 0.464 \| 153 \| 79 \| 2 \| 1 \| 150 \| 1 \| 153 \| Lysozyme \| Lysozyme; afdb-uniprot50 \| AF-A0A2E0ELT0-F1-MODEL\_V4 \| 1.0 \| 1.211e-11 \| 535 \| 0.413 \| 145 \| 81 \| 3 \| 1 \| 144 \| 2 \| 143 \| Lysozyme \| Lysozyme; afdb-uniprot50 \| AF-A0A225DZS3-F1-MODEL\_V4 \| 1.0 \| 5.766e-12 \| 535 \| 0.412 \| 143 \| 81 \| 2 \| 1 \| 143 \| 1 \| 140 \| Lysozyme \| Lysozyme; afdb-uniprot50 \| AF-A0A4Z1QPU2-F1-MODEL\_V4 \| 1.0 \| 1.346e-11 \| 534 \| 0.463 \| 151 \| 72 \| 4 \| 1 \| 146 \| 3 \| 149 \| Lysozyme \| Lysozyme; afdb-uniprot50 \| AF-A0A2W7HI45-F1-MODEL\_V4 \| 1.0 \| 6.08e-12 \| 534 \| 0.393 \| 173 \| 77 \| 1 \| 1 \| 145 \| 75 \| 247 \| Lysozyme \| Lysozyme; afdb-uniprot50 \| AF-A0A371R7Q5-F1-MODEL\_V4 \| 1.0 \| 2.542e-11 \| 534 \| 0.465 \| 144 \| 74 \| 2 \| 1 \| 144 \| 1 \| 141 \| Lysozyme \| Lysozyme; afdb-uniprot50 \| AF-A0A756FUB3-F1-MODEL\_V4 \| 1.0 \| 5.766e-12 \| 533 \| 0.35 \| 160 \| 102 \| 2 \| 1 \| 159 \| 2 \| 160 \| Lysozyme \| Lysozyme; afdb-uniprot50 \| AF-R7PL86-F1-MODEL\_V4 \| 1.0 \| 2.411e-11 \| 532 \| 0.451 \| 144 \| 75 \| 3 \| 1 \| 144 \| 3 \| 142 \| Lysozyme \| Lysozyme; afdb-uniprot50 \| AF-A0A560B879-F1-MODEL\_V4 \| 1.0 \| 7.924e-12 \| 531 \| 0.448 \| 147 \| 73 \| 5 \| 1 \| 143 \| 28 \| 170 \| Lysozyme \| Lysozyme; afdb-uniprot50 \| AF-Q4FSX3-F1-MODEL\_V4 \| 1.0 \| 1.277e-11 \| 531 \| 0.493 \| 150 \| 68 \| 3 \| 3 \| 147 \| 57 \| 203 \| Lysozyme \| Lysozyme; afdb-uniprot50 \| AF-A0A318N2U1-F1-MODEL\_V4 \| 1.0 \| 3.493e-11 \| 530 \| 0.37 \| 143 \| 86 \| 3 \| 1 \| 143 \| 1 \| 139 \| Lysozyme \| Lysozyme; afdb-uniprot50 \| AF-A0A4D8QPL9-F1-MODEL\_V4 \| 1.0 \| 7.924e-12 \| 530 \| 0.432 \| 148 \| 76 \| 5 \| 3 \| 146 \| 5 \| 148 \| Lysozyme \| Lysozyme; afdb-uniprot50 \| AF-A0A4D8QAE7-F1-MODEL\_V4 \| 1.0 \| 4.424e-12 \| 530 \| 0.429 \| 163 \| 85 \| 5 \| 1 \| 159 \| 5 \| 163 \| Lysozyme \| Lysozyme; afdb-uniprot50 \| AF-A0A0A2W020-F1-MODEL\_V4 \| 1.0 \| 6.759e-12 \| 530 \| 0.458 \| 144 \| 74 \| 3 \| 4 \| 146 \| 421 \| 561 \| Lysozyme \| Lysozyme; afdb-uniprot50 \| AF-A0A6M3XRR4-F1-MODEL\_V4 \| 1.0 \| 1.419e-11 \| 529 \| 0.445 \| 146 \| 77 \| 3 \| 1 \| 145 \| 1 \| 143 \| Lysozyme \| Lysozyme; afdb-uniprot50 \| AF-A0A495FZX1-F1-MODEL\_V4 \| 1.0 \| 6.41e-12 \| 529 \| 0.406 \| 172 \| 74 \| 2 \| 1 \| 144 \| 33 \| 204 \| Lysozyme \| Lysozyme; afdb-uniprot50 \| AF-A0A0F9HUD0-F1-MODEL\_V4 \| 1.0 \| 6.759e-12 \| 528 \| 0.414 \| 152 \| 81 \| 3 \| 1 \| 147 \| 1 \| 149 \| Lysozyme \| Lysozyme; afdb-uniprot50 \| AF-A0A662Z686-F1-MODEL\_V4 \| 1.0 \| 2.542e-11 \| 527 \| 0.442 \| 147 \| 78 \| 2 \| 1 \| 144 \| 1 \| 146 \| Lysozyme \| Lysozyme; afdb-uniprot50 \| AF-A0A561M7L1-F1-MODEL\_V4 \| 1.0 \| 1.277e-11 \| 527 \| 0.465 \| 144 \| 73 \| 3 \| 2 \| 144 \| 7 \| 147 \| Lysozyme \| Lysozyme; afdb-uniprot50 \| AF-A0A366I5P1-F1-MODEL\_V4 \| 1.0 \| 1.346e-11 \| 527 \| 0.458 \| 144 \| 74 \| 3 \| 2 \| 144 \| 7 \| 147 \| Lysozyme \| Lysozyme; afdb-uniprot50 \| AF-A0A2E7GSA7-F1-MODEL\_V4 \| 1.0 \| 2.826e-11 \| 527 \| 0.436 \| 142 \| 76 \| 3 \| 2 \| 143 \| 43 \| 180 \| Lysozyme \| Lysozyme; afdb-uniprot50 \| AF-A0A2W5V772-F1-MODEL\_V4 \| 1.0 \| 2.68e-11 \| 527 \| 0.521 \| 142 \| 65 \| 2 \| 2 \| 143 \| 73 \| 211 \| Lysozyme \| Lysozyme; afdb-uniprot50 \| AF-A0A522A0K7-F1-MODEL\_V4 \| 1.0 \| 2.98e-11 \| 526 \| 0.462 \| 147 \| 75 \| 3 \| 1 \| 147 \| 1 \| 143 \| Lysozyme \| Lysozyme; afdb-uniprot50 \| AF-A0A6P0VW89-F1-MODEL\_V4 \| 1.0 \| 9.794e-12 \| 526 \| 0.433 \| 157 \| 75 \| 3 \| 1 \| 146 \| 1 \| 154 \| Lysozyme \| Lysozyme; afdb-uniprot50 \| AF-A0A2E2D638-F1-MODEL\_V4 \| 1.0 \| 2.057e-11 \| 525 \| 0.413 \| 145 \| 81 \| 3 \| 1 \| 145 \| 21 \| 161 \| Lysozyme \| Lysozyme; afdb-uniprot50 \| AF-A0A4V3SSF5-F1-MODEL\_V4 \| 1.0 \| 5.186e-12 \| 522 \| 0.419 \| 155 \| 77 \| 4 \| 1 \| 145 \| 1 \| 152 \| Lysozyme \| Lysozyme; afdb-uniprot50 \| AF-L7U523-F1-MODEL\_V4 \| 1.0 \| 5.934e-11 \| 521 \| 0.406 \| 150 \| 82 \| 4 \| 1 \| 147 \| 1 \| 146 \| Lysozyme \| Lysozyme; afdb-uniprot50 \| AF-A0A2A4Z0X3-F1-MODEL\_V4 \| 1.0 \| 1.496e-11 \| 521 \| 0.463 \| 151 \| 72 \| 4 \| 1 \| 146 \| 2 \| 148 \| Lysozyme \| Lysozyme; afdb-uniprot50 \| AF-J1K1J4-F1-MODEL\_V4 \| 1.0 \| 3.142e-11 \| 521 \| 0.412 \| 148 \| 80 \| 3 \| 1 \| 144 \| 2 \| 146 \| Lysozyme \| Lysozyme; afdb-uniprot50 \| AF-A0A433J7K8-F1-MODEL\_V4 \| 1.0 \| 1.95e-11 \| 521 \| 0.44 \| 150 \| 76 \| 5 \| 1 \| 146 \| 12 \| 157 \| Lysozyme \| Lysozyme; afdb-uniprot50 \| AF-A0A858RGZ4-F1-MODEL\_V4 \| 1.0 \| 2.542e-11 \| 521 \| 0.439 \| 157 \| 76 \| 5 \| 1 \| 152 \| 129 \| 278 \| Lysozyme \| Lysozyme; afdb-uniprot50 \| AF-A0A840E055-F1-MODEL\_V4 \| 1.0 \| 3.884e-11 \| 520 \| 0.452 \| 148 \| 74 \| 3 \| 1 \| 144 \| 2 \| 146 \| Lysozyme \| Lysozyme; afdb-uniprot50 \| AF-A0A7T9C8N5-F1-MODEL\_V4 \| 1.0 \| 3.313e-11 \| 519 \| 0.421 \| 140 \| 78 \| 2 \| 4 \| 143 \| 9 \| 145 \| Lysozyme \| Lysozyme; afdb-uniprot50 \| AF-A0A3A4WDP1-F1-MODEL\_V4 \| 1.0 \| 4.424e-12 \| 519 \| 0.426 \| 157 \| 80 \| 5 \| 3 \| 153 \| 2 \| 154 \| Lysozyme \| Lysozyme; afdb-uniprot50 \| AF-S7Y3M8-F1-MODEL\_V4 \| 1.0 \| 3.142e-11 \| 519 \| 0.506 \| 148 \| 66 \| 3 \| 1 \| 144 \| 33 \| 177 \| Lysozyme \| Lysozyme; afdb-uniprot50 \| AF-A0A3S3SML5-F1-MODEL\_V4 \| 1.0 \| 3.313e-11 \| 519 \| 0.438 \| 146 \| 79 \| 2 \| 1 \| 146 \| 4 \| 146 \| Lysozyme \| Lysozyme; afdb-uniprot50 \| AF-A0A8A7U198-F1-MODEL\_V4 \| 1.0 \| 8.355e-12 \| 519 \| 0.5 \| 154 \| 66 \| 5 \| 1 \| 146 \| 1 \| 151 \| Glycoside hydrolase family protein \| Glycoside hydrolase family protein; afdb-uniprot50 \| AF-A9IN99-F1-MODEL\_V4 \| 1.0 \| 3.493e-11 \| 519 \| 0.418 \| 148 \| 79 \| 3 \| 1 \| 144 \| 2 \| 146 \| Lysozyme \| Lysozyme; afdb-uniprot50 \| AF-A0A355T917-F1-MODEL\_V4 \| 1.0 \| 3.884e-11 \| 518 \| 0.44 \| 143 \| 76 \| 2 \| 1 \| 143 \| 18 \| 156 \| Lysozyme \| Lysozyme; afdb-uniprot50 \| AF-A0A2Z6C8X5-F1-MODEL\_V4 \| 1.0 \| 1.277e-11 \| 518 \| 0.416 \| 161 \| 80 \| 3 \| 2 \| 152 \| 69 \| 225 \| Lysozyme \| Lysozyme; afdb-uniprot50 \| AF-A0A3E0CLR5-F1-MODEL\_V4 \| 1.0 \| 1.754e-11 \| 516 \| 0.401 \| 172 \| 75 \| 1 \| 1 \| 144 \| 22 \| 193 \| Lysozyme \| Lysozyme; afdb-uniprot50 \| AF-A0A3B0S578-F1-MODEL\_V4 \| 1.0 \| 3.142e-11 \| 516 \| 0.439 \| 148 \| 76 \| 3 \| 1 \| 144 \| 1 \| 145 \| Lysozyme \| Lysozyme; afdb-uniprot50 \| AF-A0A5J4X7D3-F1-MODEL\_V4 \| 1.0 \| 1.664e-11 \| 515 \| 0.479 \| 148 \| 69 \| 5 \| 1 \| 144 \| 31 \| 174 \| Lysozyme \| Lysozyme; afdb-uniprot50 \| AF-A0A2U1YTM3-F1-MODEL\_V4 \| 1.0 \| 1.211e-11 \| 514 \| 0.496 \| 133 \| 66 \| 1 \| 15 \| 146 \| 2 \| 134 \| Lysozyme \| Lysozyme; afdb-uniprot50 \| AF-A0A7M5WWG7-F1-MODEL\_V4 \| 1.0 \| 6.598e-11 \| 514 \| 0.444 \| 144 \| 76 \| 3 \| 1 \| 144 \| 21 \| 160 \| Lysozyme \| Lysozyme; afdb-uniprot50 \| AF-A0A371YID5-F1-MODEL\_V4 \| 1.0 \| 2.286e-11 \| 514 \| 0.506 \| 146 \| 65 \| 4 \| 1 \| 143 \| 39 \| 180 \| Lysozyme \| Lysozyme; afdb-uniprot50 \| AF-A0A3A2IVA4-F1-MODEL\_V4 \| 1.0 \| 2.542e-11 \| 514 \| 0.401 \| 152 \| 82 \| 4 \| 1 \| 146 \| 3 \| 151 \| Lysozyme \| Lysozyme; afdb-uniprot50 \| AF-A0A2X0QI68-F1-MODEL\_V4 \| 1.0 \| 2.168e-11 \| 514 \| 0.465 \| 144 \| 73 \| 2 \| 1 \| 144 \| 1 \| 140 \| Lysozyme \| Lysozyme; afdb-uniprot50 \| AF-A0A839W1G4-F1-MODEL\_V4 \| 1.0 \| 2.168e-11 \| 513 \| 0.433 \| 150 \| 77 \| 5 \| 1 \| 146 \| 12 \| 157 \| GH24 family phage-related lysozyme (Muramidase) \| GH24 family phage-related lysozyme (Muramidase); afdb-uniprot50 \| AF-A0A844IYI4-F1-MODEL\_V4 \| 1.0 \| 9.56e-11 \| 513 \| 0.486 \| 144 \| 70 \| 3 \| 1 \| 144 \| 3 \| 142 \| Glycoside hydrolase family protein \| Glycoside hydrolase family protein; afdb-uniprot50 \| AF-A0A261R4U0-F1-MODEL\_V4 \| 1.0 \| 3.884e-11 \| 512 \| 0.397 \| 146 \| 87 \| 1 \| 1 \| 145 \| 12 \| 157 \| Lysozyme \| Lysozyme; afdb-uniprot50 \| AF-A0A7S6VXX3-F1-MODEL\_V4 \| 1.0 \| 1.85e-11 \| 512 \| 0.506 \| 146 \| 65 \| 4 \| 1 \| 143 \| 36 \| 177 \| Lysozyme \| Lysozyme; afdb-uniprot50 \| AF-A0A850T1I2-F1-MODEL\_V4 \| 1.0 \| 4.553e-11 \| 512 \| 0.48 \| 150 \| 70 \| 5 \| 1 \| 146 \| 1 \| 146 \| Lysozyme \| Lysozyme; afdb-uniprot50 \| AF-N6UDE9-F1-MODEL\_V4 \| 1.0 \| 5.338e-11 \| 511 \| 0.391 \| 148 \| 83 \| 2 \| 1 \| 144 \| 2 \| 146 \| Lysozyme \| Lysozyme; afdb-uniprot50 \| AF-A0A4P6LA78-F1-MODEL\_V4 \| 1.0 \| 6.257e-11 \| 511 \| 0.41 \| 146 \| 83 \| 2 \| 2 \| 145 \| 12 \| 156 \| Lysozyme \| Lysozyme; afdb-uniprot50 \| AF-A0A2H5SCH4-F1-MODEL\_V4 \| 1.0 \| 3.142e-11 \| 510 \| 0.397 \| 146 \| 82 \| 3 \| 1 \| 143 \| 20 \| 162 \| Lysozyme \| Lysozyme; afdb-uniprot50 \| AF-A0A2N7VBE1-F1-MODEL\_V4 \| 1.0 \| 4.095e-11 \| 509 \| 0.465 \| 144 \| 73 \| 3 \| 2 \| 144 \| 7 \| 147 \| Lysozyme \| Lysozyme; afdb-uniprot50 \| AF-A0A5C7LRY7-F1-MODEL\_V4 \| 1.0 \| 4.318e-11 \| 508 \| 0.43 \| 151 \| 79 \| 4 \| 1 \| 147 \| 59 \| 206 \| Lysozyme \| Lysozyme; afdb-uniprot50 \| AF-A0A2C9BRJ3-F1-MODEL\_V4 \| 1.0 \| 5.062e-11 \| 507 \| 0.451 \| 144 \| 75 \| 3 \| 2 \| 144 \| 7 \| 147 \| Lysozyme \| Lysozyme; afdb-uniprot50 \| AF-A0A522QL01-F1-MODEL\_V4 \| 1.0 \| 6.598e-11 \| 505 \| 0.351 \| 168 \| 95 \| 5 \| 2 \| 159 \| 6 \| 169 \| Lysozyme \| Lysozyme; afdb-uniprot50 \| AF-A0A7H9ARJ5-F1-MODEL\_V4 \| 1.0 \| 3.313e-11 \| 504 \| 0.401 \| 152 \| 83 \| 4 \| 1 \| 146 \| 1 \| 150 \| Lysozyme \| Lysozyme; afdb-uniprot50 \| AF-C9XUR2-F1-MODEL\_V4 \| 1.0 \| 6.957e-11 \| 504 \| 0.462 \| 145 \| 74 \| 3 \| 2 \| 145 \| 25 \| 166 \| Lysozyme \| Lysozyme; afdb-uniprot50 \| AF-A0A2U3AP30-F1-MODEL\_V4 \| 1.0 \| 1.85e-11 \| 504 \| 0.405 \| 143 \| 78 \| 3 \| 1 \| 143 \| 1 \| 136 \| Lysozyme \| Lysozyme; afdb-uniprot50 \| AF-A0A5C7RF17-F1-MODEL\_V4 \| 1.0 \| 6.957e-11 \| 503 \| 0.39 \| 146 \| 86 \| 2 \| 1 \| 146 \| 6 \| 148 \| Lysozyme \| Lysozyme; afdb-uniprot50 \| AF-A0A7X0CNU8-F1-MODEL\_V4 \| 1.0 \| 2.826e-11 \| 503 \| 0.503 \| 155 \| 63 \| 6 \| 1 \| 144 \| 1 \| 152 \| Lysozyme \| Lysozyme; afdb-uniprot50 \| AF-B3T3K9-F1-MODEL\_V4 \| 1.0 \| 2.98e-11 \| 503 \| 0.457 \| 153 \| 73 \| 3 \| 1 \| 144 \| 21 \| 172 \| Lysozyme \| Lysozyme; afdb-uniprot50 \| AF-A0A4Y8P1F7-F1-MODEL\_V4 \| 1.0 \| 4.553e-11 \| 503 \| 0.442 \| 147 \| 78 \| 2 \| 1 \| 143 \| 1 \| 147 \| Lysozyme \| Lysozyme; afdb-uniprot50 \| AF-V4Q636-F1-MODEL\_V4 \| 1.0 \| 4.318e-11 \| 502 \| 0.456 \| 149 \| 74 \| 3 \| 1 \| 144 \| 1 \| 147 \| Lysozyme \| Lysozyme; afdb-uniprot50 \| AF-A0A4Q4GWB6-F1-MODEL\_V4 \| 1.0 \| 4.318e-11 \| 501 \| 0.486 \| 146 \| 68 \| 4 \| 1 \| 143 \| 71 \| 212 \| Lysozyme \| Lysozyme; afdb-uniprot50 \| AF-A0A6P0MMN1-F1-MODEL\_V4 \| 1.0 \| 3.313e-11 \| 501 \| 0.411 \| 158 \| 79 \| 3 \| 1 \| 147 \| 5 \| 159 \| Lysozyme \| Lysozyme; afdb-uniprot50 \| AF-A0A060DQX0-F1-MODEL\_V4 \| 1.0 \| 3.884e-11 \| 500 \| 0.413 \| 150 \| 80 \| 5 \| 1 \| 146 \| 3 \| 148 \| Lysozyme \| Lysozyme; afdb-uniprot50 \| AF-A0A2R4F7Z5-F1-MODEL\_V4 \| 1.0 \| 9.067e-11 \| 500 \| 0.434 \| 145 \| 78 \| 3 \| 1 \| 145 \| 3 \| 143 \| Lysozyme \| Lysozyme; afdb-uniprot50 \| AF-A0A7Y0FFV2-F1-MODEL\_V4 \| 1.0 \| 9.794e-12 \| 500 \| 0.372 \| 185 \| 71 \| 2 \| 6 \| 145 \| 63 \| 247 \| Lysozyme \| Lysozyme; afdb-uniprot50 \| AF-A0A6P0KSA3-F1-MODEL\_V4 \| 1.0 \| 6.257e-11 \| 499 \| 0.433 \| 150 \| 74 \| 4 \| 2 \| 144 \| 118 \| 263 \| Lysozyme \| Lysozyme; afdb-uniprot50 \| AF-A0A6P0MU79-F1-MODEL\_V4 \| 1.0 \| 2.286e-11 \| 499 \| 0.427 \| 159 \| 76 \| 3 \| 1 \| 147 \| 5 \| 160 \| Lysozyme \| Lysozyme; afdb-uniprot50 \| AF-K6ZSA0-F1-MODEL\_V4 \| 1.0 \| 1.806e-10 \| 498 \| 0.419 \| 143 \| 79 \| 3 \| 2 \| 144 \| 35 \| 173 \| Lysozyme \| Lysozyme; afdb-uniprot50 \| AF-A0A5K7ZI99-F1-MODEL\_V4 \| 1.0 \| 3.493e-11 \| 498 \| 0.407 \| 152 \| 80 \| 5 \| 1 \| 146 \| 3 \| 150 \| Lysozyme \| Lysozyme; afdb-uniprot50 \| AF-Q0B7R2-F1-MODEL\_V4 \| 1.0 \| 7.335e-11 \| 497 \| 0.451 \| 144 \| 75 \| 3 \| 2 \| 144 \| 7 \| 147 \| Lysozyme \| Lysozyme; afdb-uniprot50 \| AF-A0A5C0UIT8-F1-MODEL\_V4 \| 1.0 \| 3.684e-11 \| 497 \| 0.379 \| 145 \| 87 \| 2 \| 1 \| 145 \| 1 \| 142 \| Lysozyme \| Lysozyme; afdb-uniprot50 \| AF-A0A7H4K350-F1-MODEL\_V4 \| 1.0 \| 4.095e-11 \| 497 \| 0.486 \| 148 \| 69 \| 4 \| 1 \| 144 \| 10 \| 154 \| Lysozyme \| Lysozyme; afdb-uniprot50 \| AF-A0A2Z5ZIU1-F1-MODEL\_V4 \| 1.0 \| 2.168e-11 \| 497 \| 0.442 \| 156 \| 72 \| 3 \| 2 \| 145 \| 7 \| 159 \| Lysozyme \| Lysozyme; afdb-uniprot50 \| AF-B3T4S1-F1-MODEL\_V4 \| 1.0 \| 3.493e-11 \| 497 \| 0.457 \| 153 \| 73 \| 2 \| 1 \| 144 \| 59 \| 210 \| Lysozyme \| Lysozyme; afdb-uniprot50 \| AF-A0A7R8WLX0-F1-MODEL\_V4 \| 1.0 \| 6.257e-11 \| 497 \| 0.386 \| 145 \| 86 \| 2 \| 2 \| 146 \| 84 \| 225 \| Lysozyme \| Lysozyme; afdb-uniprot50 \| AF-A0A5P2QSJ9-F1-MODEL\_V4 \| 1.0 \| 9.067e-11 \| 496 \| 0.414 \| 147 \| 77 \| 3 \| 1 \| 143 \| 1 \| 142 \| Lysozyme \| Lysozyme; afdb-uniprot50 \| AF-A0A2T3N1M3-F1-MODEL\_V4 \| 1.0 \| 1.121e-10 \| 495 \| 0.416 \| 144 \| 80 \| 2 \| 1 \| 144 \| 26 \| 165 \| Lysozyme \| Lysozyme; afdb-uniprot50 \| AF-E8X0S4-F1-MODEL\_V4 \| 1.0 \| 5.062e-11 \| 494 \| 0.408 \| 159 \| 81 \| 4 \| 1 \| 158 \| 4 \| 150 \| Lysozyme \| Lysozyme; afdb-uniprot50 \| AF-A0A6N1BS85-F1-MODEL\_V4 \| 1.0 \| 9.067e-11 \| 494 \| 0.41 \| 139 \| 78 \| 3 \| 7 \| 145 \| 17 \| 151 \| Lysozyme \| Lysozyme; afdb-uniprot50 \| AF-F8WVD9-F1-MODEL\_V4 \| 1.0 \| 1.385e-10 \| 494 \| 0.409 \| 149 \| 80 \| 4 \| 1 \| 144 \| 37 \| 182 \| Lysozyme \| Lysozyme; afdb-uniprot50 \| AF-A0A8B6S7C1-F1-MODEL\_V4 \| 1.0 \| 5.628e-11 \| 494 \| 0.416 \| 149 \| 78 \| 5 \| 1 \| 145 \| 1 \| 144 \| Lysozyme \| Lysozyme; afdb-uniprot50 \| AF-A0A4P6GWD7-F1-MODEL\_V4 \| 1.0 \| 1.385e-10 \| 493 \| 0.394 \| 142 \| 82 \| 3 \| 3 \| 144 \| 5 \| 142 \| Lysozyme \| Lysozyme; afdb-uniprot50 \| AF-S5MZ25-F1-MODEL\_V4 \| 1.0 \| 1.385e-10 \| 493 \| 0.386 \| 150 \| 85 \| 3 \| 1 \| 146 \| 1 \| 147 \| Lysozyme \| Lysozyme; afdb-uniprot50 \| AF-A0A2B8BD98-F1-MODEL\_V4 \| 1.0 \| 7.335e-11 \| 493 \| 0.442 \| 149 \| 75 \| 5 \| 1 \| 145 \| 4 \| 148 \| Lysozyme \| Lysozyme; afdb-uniprot50 \| AF-A0A1M3KZA5-F1-MODEL\_V4 \| 1.0 \| 2.232e-10 \| 492 \| 0.445 \| 146 \| 76 \| 4 \| 1 \| 145 \| 2 \| 143 \| Lysozyme \| Lysozyme; afdb-uniprot50 \| AF-K6Z298-F1-MODEL\_V4 \| 1.0 \| 1.806e-10 \| 492 \| 0.421 \| 140 \| 77 \| 3 \| 4 \| 143 \| 49 \| 184 \| Lysozyme \| Lysozyme; afdb-uniprot50 \| AF-A0A5M6I5G4-F1-MODEL\_V4 \| 1.0 \| 4.553e-11 \| 492 \| 0.418 \| 153 \| 79 \| 5 \| 2 \| 148 \| 4 \| 152 \| Lysozyme \| Lysozyme; afdb-uniprot50 \| AF-A0A397TI77-F1-MODEL\_V4 \| 1.0 \| 6.257e-11 \| 491 \| 0.416 \| 149 \| 77 \| 4 \| 1 \| 144 \| 1 \| 144 \| Lysozyme \| Lysozyme; afdb-uniprot50 \| AF-A0A316JPI6-F1-MODEL\_V4 \| 1.0 \| 5.628e-11 \| 491 \| 0.472 \| 146 \| 70 \| 4 \| 1 \| 143 \| 1 \| 142 \| Lysozyme \| Lysozyme; afdb-uniprot50 \| AF-A0A2N1MZ41-F1-MODEL\_V4 \| 1.0 \| 1.121e-10 \| 490 \| 0.356 \| 146 \| 87 \| 2 \| 1 \| 143 \| 5 \| 146 \| Lysozyme \| Lysozyme; afdb-uniprot50 \| AF-A0A271JAU0-F1-MODEL\_V4 \| 1.0 \| 1.008e-10 \| 489 \| 0.439 \| 148 \| 77 \| 4 \| 1 \| 146 \| 9 \| 152 \| Lysozyme \| Lysozyme; afdb-uniprot50 \| AF-A0A5F0MKV9-F1-MODEL\_V4 \| 1.0 \| 1.063e-10 \| 489 \| 0.363 \| 146 \| 89 \| 3 \| 2 \| 147 \| 7 \| 148 \| Lysozyme \| Lysozyme; afdb-uniprot50 \| AF-A0A1M6VYX9-F1-MODEL\_V4 \| 1.0 \| 6.598e-11 \| 488 \| 0.409 \| 149 \| 79 \| 5 \| 1 \| 143 \| 1 \| 146 \| Lysozyme \| Lysozyme; afdb-uniprot50 \| AF-A0A286GNA9-F1-MODEL\_V4 \| 1.0 \| 5.062e-11 \| 488 \| 0.452 \| 148 \| 72 \| 4 \| 6 \| 147 \| 12 \| 156 \| Lysozyme \| Lysozyme; afdb-uniprot50 \| AF-A0A151QE01-F1-MODEL\_V4 \| 1.0 \| 1.121e-10 \| 488 \| 0.379 \| 145 \| 87 \| 2 \| 2 \| 146 \| 17 \| 158 \| Lysozyme \| Lysozyme; afdb-uniprot50 \| AF-A0A848YF79-F1-MODEL\_V4 \| 1.0 \| 3.884e-11 \| 488 \| 0.422 \| 161 \| 85 \| 4 \| 1 \| 157 \| 1 \| 157 \| Lysozyme \| Lysozyme; afdb-uniprot50 \| AF-A0A149TN60-F1-MODEL\_V4 \| 1.0 \| 5.338e-11 \| 486 \| 0.409 \| 154 \| 77 \| 3 \| 1 \| 143 \| 1 \| 151 \| Lysozyme \| Lysozyme; afdb-uniprot50 \| AF-A0A225M2C7-F1-MODEL\_V4 \| 1.0 \| 2.117e-10 \| 486 \| 0.371 \| 140 \| 85 \| 2 \| 5 \| 144 \| 22 \| 158 \| Lysozyme \| Lysozyme; afdb-uniprot50 \| AF-A0A239Q0C8-F1-MODEL\_V4 \| 1.0 \| 4.318e-11 \| 486 \| 0.424 \| 172 \| 74 \| 6 \| 1 \| 150 \| 7 \| 175 \| Lysozyme \| Lysozyme; afdb-uniprot50 \| AF-A0A3Q8ZIN2-F1-MODEL\_V4 \| 1.0 \| 3.313e-11 \| 486 \| 0.445 \| 155 \| 75 \| 5 \| 1 \| 147 \| 1 \| 152 \| Lysozyme \| Lysozyme; afdb-uniprot50 \| AF-A0A241W0V3-F1-MODEL\_V4 \| 1.0 \| 1.385e-10 \| 486 \| 0.476 \| 149 \| 70 \| 3 \| 1 \| 145 \| 87 \| 231 \| Lysozyme \| Lysozyme; afdb-uniprot50 \| AF-A0A428KE41-F1-MODEL\_V4 \| 1.0 \| 7.335e-11 \| 485 \| 0.423 \| 151 \| 79 \| 3 \| 1 \| 146 \| 5 \| 152 \| Lysozyme \| Lysozyme; afdb-uniprot50 \| AF-A0A402TH11-F1-MODEL\_V4 \| 1.0 \| 8.599e-11 \| 485 \| 0.394 \| 142 \| 81 \| 3 \| 6 \| 145 \| 23 \| 161 \| Lysozyme \| Lysozyme; afdb-uniprot50 \| AF-A0A4Y3ZNK5-F1-MODEL\_V4 \| 1.0 \| 4.801e-11 \| 485 \| 0.433 \| 159 \| 80 \| 5 \| 1 \| 155 \| 1 \| 153 \| Lysozyme \| Lysozyme; afdb-uniprot50 \| AF-A0A839Y491-F1-MODEL\_V4 \| 1.0 \| 1.121e-10 \| 485 \| 0.457 \| 166 \| 80 \| 6 \| 1 \| 159 \| 94 \| 256 \| GH24 family phage-related lysozyme (Muramidase) \| GH24 family phage-related lysozyme (Muramidase); afdb-uniprot50 \| AF-Q28PX4-F1-MODEL\_V4 \| 1.0 \| 9.56e-11 \| 485 \| 0.401 \| 147 \| 81 \| 4 \| 1 \| 143 \| 1 \| 144 \| Lysozyme \| Lysozyme; afdb-uniprot50 \| AF-A0A3S0C8Q3-F1-MODEL\_V4 \| 1.0 \| 1.461e-10 \| 484 \| 0.4 \| 145 \| 84 \| 2 \| 1 \| 145 \| 1 \| 142 \| Lysozyme \| Lysozyme; afdb-uniprot50 \| AF-A0A2M8AHY6-F1-MODEL\_V4 \| 1.0 \| 1.806e-10 \| 484 \| 0.449 \| 149 \| 76 \| 3 \| 1 \| 146 \| 1 \| 146 \| Lysozyme \| Lysozyme; afdb-uniprot50 \| AF-A0A3B9NR42-F1-MODEL\_V4 \| 1.0 \| 1.121e-10 \| 484 \| 0.408 \| 142 \| 77 \| 4 \| 6 \| 144 \| 63 \| 200 \| Lysozyme \| Lysozyme; afdb-uniprot50 \| AF-A0A3P1XK73-F1-MODEL\_V4 \| 1.0 \| 3.595e-10 \| 484 \| 0.454 \| 143 \| 75 \| 2 \| 1 \| 143 \| 99 \| 238 \| Lysozyme \| Lysozyme; afdb-uniprot50 \| AF-A0A167HMV7-F1-MODEL\_V4 \| 1.0 \| 3.067e-10 \| 483 \| 0.45 \| 151 \| 75 \| 3 \| 1 \| 146 \| 1 \| 148 \| Lysozyme \| Lysozyme; afdb-uniprot50 \| AF-A0A3D1CPK1-F1-MODEL\_V4 \| 1.0 \| 1.182e-10 \| 483 \| 0.348 \| 152 \| 91 \| 3 \| 1 \| 147 \| 1 \| 149 \| Lysozyme \| Lysozyme; afdb-uniprot50 \| AF-A0A2E4PLW6-F1-MODEL\_V4 \| 1.0 \| 1.246e-10 \| 483 \| 0.363 \| 154 \| 88 \| 2 \| 1 \| 145 \| 1 \| 153 \| Lysozyme \| Lysozyme; afdb-uniprot50 \| AF-A0A258A958-F1-MODEL\_V4 \| 1.0 \| 4.553e-11 \| 483 \| 0.398 \| 168 \| 84 \| 7 \| 2 \| 157 \| 5 \| 167 \| Lysozyme \| Lysozyme; afdb-uniprot50 \| AF-A0A552BAQ5-F1-MODEL\_V4 \| 1.0 \| 4.095e-11 \| 483 \| 0.456 \| 151 \| 71 \| 6 \| 1 \| 143 \| 1 \| 148 \| Lysozyme \| Lysozyme; afdb-uniprot50 \| AF-A0A3N0V5X1-F1-MODEL\_V4 \| 1.0 \| 2.007e-10 \| 482 \| 0.409 \| 144 \| 81 \| 3 \| 1 \| 144 \| 5 \| 144 \| Lysozyme \| Lysozyme; afdb-uniprot50 \| AF-A0A709N1J6-F1-MODEL\_V4 \| 1.0 \| 5.062e-11 \| 482 \| 0.401 \| 152 \| 81 \| 5 \| 1 \| 145 \| 1 \| 149 \| Lysozyme \| Lysozyme; afdb-uniprot50 \| AF-A0A6P0UJS3-F1-MODEL\_V4 \| 1.0 \| 9.067e-11 \| 482 \| 0.428 \| 147 \| 76 \| 5 \| 1 \| 143 \| 1 \| 143 \| Lysozyme \| Lysozyme; afdb-uniprot50 \| AF-A0A1G6JCW9-F1-MODEL\_V4 \| 1.0 \| 2.353e-10 \| 482 \| 0.432 \| 148 \| 77 \| 3 \| 1 \| 144 \| 60 \| 204 \| Lysozyme \| Lysozyme; afdb-uniprot50 \| AF-A0A6M0G4D1-F1-MODEL\_V4 \| 1.0 \| 7.734e-11 \| 482 \| 0.398 \| 158 \| 81 \| 2 \| 1 \| 147 \| 5 \| 159 \| Lysozyme \| Lysozyme; afdb-uniprot50 \| AF-A0A2E2VXS1-F1-MODEL\_V4 \| 1.0 \| 8.599e-11 \| 481 \| 0.344 \| 148 \| 89 \| 3 \| 1 \| 143 \| 1 \| 145 \| Lysozyme \| Lysozyme; afdb-uniprot50 \| AF-A0A5U3EPL2-F1-MODEL\_V4 \| 1.0 \| 9.56e-11 \| 481 \| 0.427 \| 152 \| 77 \| 5 \| 1 \| 147 \| 1 \| 147 \| Lysozyme \| Lysozyme; afdb-uniprot50 \| AF-A0A1E8CGD7-F1-MODEL\_V4 \| 1.0 \| 6.957e-11 \| 481 \| 0.384 \| 151 \| 85 \| 4 \| 1 \| 147 \| 1 \| 147 \| Lysozyme \| Lysozyme; afdb-uniprot50 \| AF-A0A3S4Z1L8-F1-MODEL\_V4 \| 1.0 \| 1.712e-10 \| 481 \| 0.384 \| 151 \| 83 \| 4 \| 1 \| 144 \| 2 \| 149 \| Lysozyme \| Lysozyme; afdb-uniprot50 \| AF-A0A317H2W4-F1-MODEL\_V4 \| 1.0 \| 8.155e-11 \| 481 \| 0.425 \| 155 \| 75 \| 4 \| 1 \| 144 \| 10 \| 161 \| Lysozyme \| Lysozyme; afdb-uniprot50 \| AF-A0A3B0SY71-F1-MODEL\_V4 \| 1.0 \| 3.791e-10 \| 481 \| 0.409 \| 144 \| 80 \| 2 \| 1 \| 144 \| 94 \| 232 \| Lysozyme \| Lysozyme; afdb-uniprot50 \| AF-A0A357LA02-F1-MODEL\_V4 \| 1.0 \| 1.182e-10 \| 481 \| 0.371 \| 148 \| 85 \| 4 \| 1 \| 144 \| 1 \| 144 \| Lysozyme \| Lysozyme; afdb-uniprot50 \| AF-A0A4Q5XWR4-F1-MODEL\_V4 \| 1.0 \| 1.54e-10 \| 481 \| 0.398 \| 148 \| 81 \| 5 \| 1 \| 144 \| 23 \| 166 \| Lysozyme \| Lysozyme; afdb-uniprot50 \| AF-A0A1F6ZTS5-F1-MODEL\_V4 \| 1.0 \| 2.232e-10 \| 480 \| 0.379 \| 145 \| 85 \| 2 \| 3 \| 146 \| 1 \| 141 \| Lysozyme \| Lysozyme; afdb-uniprot50 \| AF-A0A2P1P7G1-F1-MODEL\_V4 \| 1.0 \| 1.712e-10 \| 480 \| 0.353 \| 147 \| 92 \| 2 \| 1 \| 147 \| 1 \| 144 \| Lysozyme \| Lysozyme; afdb-uniprot50 \| AF-A0A3D5RRE8-F1-MODEL\_V4 \| 1.0 \| 1.904e-10 \| 480 \| 0.326 \| 150 \| 93 \| 2 \| 2 \| 146 \| 19 \| 165 \| Lysozyme \| Lysozyme; afdb-uniprot50 \| AF-A0A5C7PF01-F1-MODEL\_V4 \| 1.0 \| 2.98e-11 \| 480 \| 0.364 \| 162 \| 86 \| 4 \| 3 \| 150 \| 2 \| 160 \| Lysozyme \| Lysozyme; afdb-uniprot50 \| AF-R3W5G4-F1-MODEL\_V4 \| 1.0 \| 2.232e-10 \| 480 \| 0.439 \| 148 \| 78 \| 4 \| 1 \| 147 \| 6 \| 149 \| Lysozyme \| Lysozyme; afdb-uniprot50 \| AF-A0A4Q3DIP3-F1-MODEL\_V4 \| 1.0 \| 2.616e-10 \| 479 \| 0.381 \| 144 \| 86 \| 2 \| 1 \| 144 \| 3 \| 143 \| Lysozyme \| Lysozyme; afdb-uniprot50 \| AF-A0A1U7NBN1-F1-MODEL\_V4 \| 1.0 \| 2.353e-10 \| 479 \| 0.44 \| 150 \| 73 \| 4 \| 2 \| 144 \| 8 \| 153 \| Lysozyme \| Lysozyme; afdb-uniprot50 \| AF-A0A0G3I3X2-F1-MODEL\_V4 \| 1.0 \| 2.007e-10 \| 479 \| 0.335 \| 146 \| 93 \| 2 \| 3 \| 146 \| 20 \| 163 \| Lysozyme \| Lysozyme; afdb-uniprot50 \| AF-A0A845N7Q8-F1-MODEL\_V4 \| 1.0 \| 2.007e-10 \| 479 \| 0.391 \| 148 \| 84 \| 3 \| 1 \| 145 \| 2 \| 146 \| Glycoside hydrolase family protein \| Glycoside hydrolase family protein; afdb-uniprot50 \| AF-A0A7W5C0N3-F1-MODEL\_V4 \| 1.0 \| 3.234e-10 \| 479 \| 0.397 \| 141 \| 81 \| 3 \| 4 \| 144 \| 1 \| 137 \| Lysozyme \| Lysozyme; afdb-uniprot50 \| AF-A0A0M3A474-F1-MODEL\_V4 \| 1.0 \| 8.155e-11 \| 479 \| 0.469 \| 149 \| 72 \| 4 \| 2 \| 146 \| 52 \| 197 \| Lysozyme \| Lysozyme; afdb-uniprot50 \| AF-A0A239Q0D5-F1-MODEL\_V4 \| 1.0 \| 4.941e-10 \| 479 \| 0.388 \| 144 \| 83 \| 3 \| 1 \| 144 \| 76 \| 214 \| Lysozyme \| Lysozyme; afdb-uniprot50 \| AF-A0A7G8FMI4-F1-MODEL\_V4 \| 1.0 \| 1.314e-10 \| 478 \| 0.424 \| 146 \| 78 \| 5 \| 2 \| 144 \| 3 \| 145 \| Lysozyme \| Lysozyme; afdb-uniprot50 \| AF-A0A1F9BXF9-F1-MODEL\_V4 \| 1.0 \| 1.008e-10 \| 477 \| 0.44 \| 143 \| 77 \| 2 \| 1 \| 143 \| 89 \| 228 \| Lysozyme \| Lysozyme; afdb-uniprot50 \| AF-A0A1H9MQ78-F1-MODEL\_V4 \| 1.0 \| 2.616e-10 \| 476 \| 0.391 \| 143 \| 82 \| 4 \| 5 \| 146 \| 17 \| 155 \| Lysozyme \| Lysozyme; afdb-uniprot50 \| AF-A0A496C4E2-F1-MODEL\_V4 \| 1.0 \| 2.117e-10 \| 476 \| 0.434 \| 145 \| 74 \| 6 \| 2 \| 144 \| 17 \| 155 \| Lysozyme \| Lysozyme; afdb-uniprot50 \| AF-A0A522GMT4-F1-MODEL\_V4 \| 1.0 \| 5.338e-11 \| 476 \| 0.407 \| 162 \| 84 \| 4 \| 1 \| 153 \| 1 \| 159 \| Lysozyme \| Lysozyme; afdb-uniprot50 \| AF-A0A4P6YSL8-F1-MODEL\_V4 \| 1.0 \| 1.712e-10 \| 476 \| 0.452 \| 146 \| 76 \| 3 \| 1 \| 146 \| 5 \| 146 \| Lysozyme \| Lysozyme; afdb-uniprot50 \| AF-A0A6H9GT10-F1-MODEL\_V4 \| 1.0 \| 8.599e-11 \| 476 \| 0.438 \| 155 \| 76 \| 6 \| 1 \| 147 \| 1 \| 152 \| Lysozyme \| Lysozyme; afdb-uniprot50 \| AF-A0A0N1L9V2-F1-MODEL\_V4 \| 1.0 \| 3.067e-10 \| 475 \| 0.387 \| 147 \| 85 \| 3 \| 1 \| 147 \| 1 \| 142 \| Lysozyme \| Lysozyme; afdb-uniprot50 \| AF-A0A2N0R9A1-F1-MODEL\_V4 \| 1.0 \| 2.616e-10 \| 475 \| 0.394 \| 152 \| 80 \| 5 \| 1 \| 146 \| 3 \| 148 \| Lysozyme \| Lysozyme; afdb-uniprot50 \| AF-A0A3M9MZF2-F1-MODEL\_V4 \| 1.0 \| 1.182e-10 \| 475 \| 0.409 \| 149 \| 82 \| 3 \| 1 \| 146 \| 3 \| 148 \| Lysozyme \| Lysozyme; afdb-uniprot50 \| AF-A0A1T4RPF3-F1-MODEL\_V4 \| 1.0 \| 5.934e-11 \| 475 \| 0.448 \| 156 \| 71 \| 2 \| 1 \| 144 \| 1 \| 153 \| Lysozyme \| Lysozyme; afdb-uniprot50 \| AF-A0A0F9NJU4-F1-MODEL\_V4 \| 1.0 \| 9.067e-11 \| 475 \| 0.407 \| 152 \| 78 \| 3 \| 1 \| 143 \| 1 \| 149 \| Lysozyme \| Lysozyme; afdb-uniprot50 \| AF-A0A2D7ZXW1-F1-MODEL\_V4 \| 1.0 \| 2.481e-10 \| 475 \| 0.414 \| 152 \| 81 \| 3 \| 1 \| 147 \| 2 \| 150 \| Lysozyme \| Lysozyme; afdb-uniprot50 \| AF-A0A6D2B246-F1-MODEL\_V4 \| 1.0 \| 3.067e-10 \| 474 \| 0.534 \| 116 \| 50 \| 1 \| 1 \| 112 \| 2 \| 117 \| Lysozyme \| Lysozyme; afdb-uniprot50 \| AF-A0A2E1XDB6-F1-MODEL\_V4 \| 1.0 \| 2.481e-10 \| 474 \| 0.364 \| 151 \| 87 \| 4 \| 1 \| 147 \| 1 \| 146 \| Lysozyme \| Lysozyme; afdb-uniprot50 \| AF-A0A109BL19-F1-MODEL\_V4 \| 1.0 \| 1.624e-10 \| 474 \| 0.418 \| 153 \| 78 \| 4 \| 1 \| 146 \| 15 \| 163 \| Lysozyme \| Lysozyme; afdb-uniprot50 \| AF-A0A7G8J609-F1-MODEL\_V4 \| 1.0 \| 3.41e-10 \| 474 \| 0.393 \| 150 \| 87 \| 3 \| 1 \| 147 \| 3 \| 151 \| Lysozyme \| Lysozyme; afdb-uniprot50 \| AF-A0A496C4S7-F1-MODEL\_V4 \| 1.0 \| 4.215e-10 \| 473 \| 0.427 \| 145 \| 75 \| 6 \| 1 \| 143 \| 1 \| 139 \| Lysozyme \| Lysozyme; afdb-uniprot50 \| AF-A0A7S7C260-F1-MODEL\_V4 \| 1.0 \| 3.884e-11 \| 473 \| 0.401 \| 152 \| 80 \| 6 \| 4 \| 147 \| 1 \| 149 \| Lysozyme \| Lysozyme; afdb-uniprot50 \| AF-A0A2S0V4Q7-F1-MODEL\_V4 \| 1.0 \| 9.331e-10 \| 473 \| 0.376 \| 146 \| 87 \| 3 \| 1 \| 146 \| 76 \| 217 \| Lysozyme \| Lysozyme; afdb-uniprot50 \| AF-A0A2S6HCU7-F1-MODEL\_V4 \| 1.0 \| 1.246e-10 \| 473 \| 0.394 \| 152 \| 82 \| 5 \| 1 \| 146 \| 3 \| 150 \| Lysozyme \| Lysozyme; afdb-uniprot50 \| AF-A0A1M3AIX6-F1-MODEL\_V4 \| 1.0 \| 9.56e-11 \| 473 \| 0.403 \| 156 \| 82 \| 6 \| 1 \| 148 \| 1 \| 153 \| Lysozyme \| Lysozyme; afdb-uniprot50 \| AF-A0A608BLC2-F1-MODEL\_V4 \| 1.0 \| 3.067e-10 \| 472 \| 0.414 \| 152 \| 79 \| 5 \| 1 \| 147 \| 1 \| 147 \| Lysozyme \| Lysozyme; afdb-uniprot50 \| AF-K9ZAF2-F1-MODEL\_V4 \| 1.0 \| 5.792e-10 \| 472 \| 0.387 \| 147 \| 83 \| 3 \| 2 \| 144 \| 80 \| 223 \| Lysozyme \| Lysozyme; afdb-uniprot50 \| AF-A0A7Z1UR98-F1-MODEL\_V4 \| 1.0 \| 4.444e-10 \| 472 \| 0.426 \| 150 \| 79 \| 3 \| 1 \| 146 \| 81 \| 227 \| Lysozyme \| Lysozyme; afdb-uniprot50 \| AF-A0A3Q9BKE1-F1-MODEL\_V4 \| 1.0 \| 2.759e-10 \| 472 \| 0.451 \| 144 \| 75 \| 3 \| 1 \| 144 \| 6 \| 145 \| Lysozyme \| Lysozyme; afdb-uniprot50 \| AF-A0A1J5F4W9-F1-MODEL\_V4 \| 1.0 \| 4.941e-10 \| 470 \| 0.364 \| 151 \| 88 \| 4 \| 1 \| 146 \| 1 \| 148 \| Lysozyme \| Lysozyme; afdb-uniprot50 \| AF-A0A1F2Z7P5-F1-MODEL\_V4 \| 1.0 \| 2.481e-10 \| 470 \| 0.352 \| 142 \| 89 \| 2 \| 5 \| 146 \| 15 \| 153 \| Lysozyme \| Lysozyme; afdb-uniprot50 \| AF-A0A258GYK9-F1-MODEL\_V4 \| 1.0 \| 2.909e-10 \| 470 \| 0.391 \| 148 \| 82 \| 5 \| 1 \| 144 \| 11 \| 154 \| Lysozyme \| Lysozyme; afdb-uniprot50 \| AF-A0A2M7K8A1-F1-MODEL\_V4 \| 1.0 \| 5.21e-10 \| 469 \| 0.393 \| 145 \| 85 \| 2 \| 1 \| 145 \| 3 \| 144 \| Lysozyme \| Lysozyme; afdb-uniprot50 \| AF-A0A4Q7BRX3-F1-MODEL\_V4 \| 1.0 \| 2.759e-10 \| 469 \| 0.489 \| 147 \| 67 \| 5 \| 1 \| 143 \| 22 \| 164 \| Lysozyme \| Lysozyme; afdb-uniprot50 \| AF-A0A7Y3RIZ1-F1-MODEL\_V4 \| 1.0 \| 9.331e-10 \| 469 \| 0.435 \| 140 \| 76 \| 2 \| 6 \| 145 \| 435 \| 571 \| Lysozyme \| Lysozyme; afdb-uniprot50 \| AF-A0A250LRH8-F1-MODEL\_V4 \| 1.0 \| 4.686e-10 \| 468 \| 0.349 \| 146 \| 91 \| 3 \| 1 \| 145 \| 3 \| 145 \| Lysozyme \| Lysozyme; afdb-uniprot50 \| AF-A0A2W6VVK0-F1-MODEL\_V4 \| 1.0 \| 8.155e-11 \| 468 \| 0.402 \| 149 \| 81 \| 5 \| 1 \| 145 \| 1 \| 145 \| Lysozyme \| Lysozyme; afdb-uniprot50 \| AF-A0A5C7QE19-F1-MODEL\_V4 \| 1.0 \| 2.117e-10 \| 468 \| 0.396 \| 154 \| 82 \| 3 \| 1 \| 147 \| 1 \| 150 \| Lysozyme \| Lysozyme; afdb-uniprot50 \| AF-A0A1I5CM08-F1-MODEL\_V4 \| 1.0 \| 2.353e-10 \| 467 \| 0.528 \| 125 \| 58 \| 1 \| 20 \| 144 \| 2 \| 125 \| Lysozyme \| Lysozyme; afdb-uniprot50 \| AF-Q2NTN7-F1-MODEL\_V4 \| 1.0 \| 3.791e-10 \| 467 \| 0.414 \| 147 \| 78 \| 4 \| 1 \| 143 \| 2 \| 144 \| Lysozyme \| Lysozyme; afdb-uniprot50 \| AF-A0A619AGL7-F1-MODEL\_V4 \| 1.0 \| 2.759e-10 \| 467 \| 0.414 \| 152 \| 79 \| 5 \| 1 \| 147 \| 1 \| 147 \| Lysozyme \| Lysozyme; afdb-uniprot50 \| AF-A0A5E4SJE8-F1-MODEL\_V4 \| 1.0 \| 7.549e-10 \| 467 \| 0.407 \| 135 \| 76 \| 3 \| 10 \| 144 \| 22 \| 152 \| Lysozyme \| Lysozyme; afdb-uniprot50 \| AF-A0A4Q0D5W1-F1-MODEL\_V4 \| 1.0 \| 4.215e-10 \| 467 \| 0.417 \| 146 \| 81 \| 3 \| 1 \| 146 \| 6 \| 147 \| Lysozyme \| Lysozyme; afdb-uniprot50 \| AF-A0A6P1NBT1-F1-MODEL\_V4 \| 1.0 \| 2.117e-10 \| 466 \| 0.397 \| 146 \| 80 \| 4 \| 5 \| 145 \| 6 \| 148 \| Lysozyme \| Lysozyme; afdb-uniprot50 \| AF-S2K1B0-F1-MODEL\_V4 \| 1.0 \| 4.444e-10 \| 466 \| 0.398 \| 143 \| 81 \| 4 \| 5 \| 146 \| 32 \| 170 \| Lysozyme \| Lysozyme; afdb-uniprot50 \| AF-A0A7C5WIK9-F1-MODEL\_V4 \| 1.0 \| 1.461e-10 \| 466 \| 0.387 \| 160 \| 91 \| 4 \| 2 \| 157 \| 55 \| 211 \| Lysozyme \| Lysozyme; afdb-uniprot50 \| AF-A0A551Z3X8-F1-MODEL\_V4 \| 1.0 \| 2.232e-10 \| 466 \| 0.448 \| 156 \| 75 \| 5 \| 1 \| 148 \| 1 \| 153 \| Lysozyme \| Lysozyme; afdb-uniprot50 \| AF-A0A521YLX8-F1-MODEL\_V4 \| 1.0 \| 7.549e-10 \| 465 \| 0.401 \| 147 \| 82 \| 5 \| 1 \| 145 \| 4 \| 146 \| Lysozyme \| Lysozyme; afdb-uniprot50 \| AF-A0A8A8CIQ3-F1-MODEL\_V4 \| 1.0 \| 5.792e-10 \| 465 \| 0.402 \| 139 \| 80 \| 2 \| 7 \| 145 \| 19 \| 154 \| Lysozyme \| Lysozyme; afdb-uniprot50 \| AF-A0A1V0DDE0-F1-MODEL\_V4 \| 1.0 \| 3.234e-10 \| 465 \| 0.406 \| 150 \| 79 \| 4 \| 2 \| 144 \| 12 \| 158 \| Lysozyme \| Lysozyme; afdb-uniprot50 \| AF-A0A192H186-F1-MODEL\_V4 \| 1.0 \| 2.481e-10 \| 465 \| 0.407 \| 152 \| 79 \| 5 \| 1 \| 144 \| 538 \| 686 \| Lysozyme \| Lysozyme; afdb-uniprot50 \| AF-A0A0S8L002-F1-MODEL\_V4 \| 1.0 \| 2.007e-10 \| 464 \| 0.445 \| 148 \| 75 \| 2 \| 12 \| 155 \| 2 \| 146 \| Lysozyme \| Lysozyme; afdb-uniprot50 \| AF-A0A4Z0DE45-F1-MODEL\_V4 \| 1.0 \| 9.56e-11 \| 464 \| 0.387 \| 147 \| 83 \| 3 \| 2 \| 144 \| 5 \| 148 \| Lysozyme \| Lysozyme; afdb-uniprot50 \| AF-A0A2G2HRY1-F1-MODEL\_V4 \| 1.0 \| 3.41e-10 \| 464 \| 0.358 \| 148 \| 87 \| 4 \| 3 \| 146 \| 7 \| 150 \| Lysozyme \| Lysozyme; afdb-uniprot50 \| AF-A0A1V2N9M7-F1-MODEL\_V4 \| 1.0 \| 4.686e-10 \| 463 \| 0.378 \| 148 \| 87 \| 3 \| 1 \| 146 \| 2 \| 146 \| Lysozyme \| Lysozyme; afdb-uniprot50 \| AF-A0A5V7UAR2-F1-MODEL\_V4 \| 1.0 \| 4.215e-10 \| 463 \| 0.42 \| 150 \| 77 \| 5 \| 1 \| 145 \| 1 \| 145 \| Lysozyme \| Lysozyme; afdb-uniprot50 \| AF-A0A1G9W3A4-F1-MODEL\_V4 \| 1.0 \| 3.791e-10 \| 462 \| 0.366 \| 150 \| 89 \| 3 \| 1 \| 147 \| 2 \| 148 \| Lysozyme \| Lysozyme; afdb-uniprot50 \| AF-A0A7S9LKT4-F1-MODEL\_V4 \| 1.0 \| 6.107e-10 \| 462 \| 0.377 \| 143 \| 84 \| 4 \| 5 \| 146 \| 16 \| 154 \| Lysozyme \| Lysozyme; afdb-uniprot50 \| AF-W6M7Y1-F1-MODEL\_V4 \| 1.0 \| 9.56e-11 \| 462 \| 0.43 \| 158 \| 75 \| 6 \| 1 \| 146 \| 8 \| 162 \| Lysozyme \| Lysozyme; afdb-uniprot50 \| AF-A0A6J4RNY5-F1-MODEL\_V4 \| 1.0 \| 9.56e-11 \| 462 \| 0.415 \| 154 \| 77 \| 2 \| 1 \| 144 \| 622 \| 772 \| Lysozyme \| Lysozyme; afdb-uniprot50 \| AF-Z9JHW0-F1-MODEL\_V4 \| 1.0 \| 4.686e-10 \| 461 \| 0.34 \| 141 \| 90 \| 2 \| 5 \| 145 \| 4 \| 141 \| Lysozyme \| Lysozyme; afdb-uniprot50 \| AF-L1M7X0-F1-MODEL\_V4 \| 1.0 \| 6.79e-10 \| 461 \| 0.361 \| 144 \| 88 \| 3 \| 1 \| 144 \| 1 \| 140 \| Lysozyme \| Lysozyme; afdb-uniprot50 \| AF-A0A629H5A7-F1-MODEL\_V4 \| 1.0 \| 4.444e-10 \| 461 \| 0.42 \| 150 \| 77 \| 5 \| 1 \| 145 \| 1 \| 145 \| Lysozyme \| Lysozyme; afdb-uniprot50 \| AF-A0A7W5WZ35-F1-MODEL\_V4 \| 1.0 \| 3.234e-10 \| 461 \| 0.335 \| 146 \| 93 \| 2 \| 1 \| 143 \| 5 \| 149 \| Lysozyme \| Lysozyme; afdb-uniprot50 \| AF-A0A1G4TH47-F1-MODEL\_V4 \| 1.0 \| 1.904e-10 \| 461 \| 0.422 \| 154 \| 81 \| 5 \| 1 \| 150 \| 5 \| 154 \| Lysozyme \| Lysozyme; afdb-uniprot50 \| AF-A0A4Y8S5U9-F1-MODEL\_V4 \| 1.0 \| 3.41e-10 \| 460 \| 0.423 \| 151 \| 79 \| 4 \| 1 \| 146 \| 1 \| 148 \| Lysozyme \| Lysozyme; afdb-uniprot50 \| AF-A0A3A1YD39-F1-MODEL\_V4 \| 1.0 \| 3.067e-10 \| 460 \| 0.412 \| 143 \| 79 \| 3 \| 6 \| 146 \| 10 \| 149 \| Lysozyme \| Lysozyme; afdb-uniprot50 \| AF-A0A494QR79-F1-MODEL\_V4 \| 1.0 \| 5.493e-10 \| 460 \| 0.435 \| 147 \| 79 \| 3 \| 1 \| 147 \| 6 \| 148 \| Lysozyme \| Lysozyme; afdb-uniprot50 \| AF-C6XGM6-F1-MODEL\_V4 \| 1.0 \| 6.107e-10 \| 459 \| 0.371 \| 148 \| 89 \| 3 \| 1 \| 146 \| 26 \| 171 \| Lysozyme \| Lysozyme; afdb-uniprot50 \| AF-A0A257RPK0-F1-MODEL\_V4 \| 1.0 \| 1.246e-10 \| 458 \| 0.405 \| 143 \| 77 \| 4 \| 6 \| 143 \| 7 \| 146 \| Lysozyme \| Lysozyme; afdb-uniprot50 \| AF-A0A7U1H6U5-F1-MODEL\_V4 \| 1.0 \| 8.393e-10 \| 458 \| 0.388 \| 139 \| 82 \| 2 \| 7 \| 145 \| 19 \| 154 \| Lysozyme \| Lysozyme; afdb-uniprot50 \| AF-A0A3P3QN20-F1-MODEL\_V4 \| 1.0 \| 9.839e-10 \| 458 \| 0.366 \| 139 \| 85 \| 1 \| 6 \| 144 \| 20 \| 155 \| Lysozyme \| Lysozyme; afdb-uniprot50 \| AF-A0A344WGG6-F1-MODEL\_V4 \| 1.0 \| 5.493e-10 \| 458 \| 0.375 \| 149 \| 84 \| 6 \| 1 \| 144 \| 5 \| 149 \| Lysozyme \| Lysozyme; afdb-uniprot50 \| AF-A0A328B6S8-F1-MODEL\_V4 \| 1.0 \| 3.595e-10 \| 458 \| 0.375 \| 152 \| 86 \| 6 \| 2 \| 148 \| 6 \| 153 \| Lysozyme \| Lysozyme; afdb-uniprot50 \| AF-A0A7T8NV96-F1-MODEL\_V4 \| 1.0 \| 3.791e-10 \| 458 \| 0.339 \| 159 \| 94 \| 5 \| 1 \| 154 \| 1 \| 153 \| Lysozyme \| Lysozyme; afdb-uniprot50 \| AF-A0A4Q2JA85-F1-MODEL\_V4 \| 1.0 \| 7.549e-10 \| 457 \| 0.392 \| 140 \| 81 \| 3 \| 7 \| 146 \| 21 \| 156 \| Lysozyme \| Lysozyme; afdb-uniprot50 \| AF-A0A494Y8X4-F1-MODEL\_V4 \| 1.0 \| 3.997e-10 \| 457 \| 0.387 \| 147 \| 84 \| 4 \| 1 \| 144 \| 1 \| 144 \| Lysozyme \| Lysozyme; afdb-uniprot50 \| AF-A0A6N7D1K0-F1-MODEL\_V4 \| 1.0 \| 5.21e-10 \| 457 \| 0.405 \| 143 \| 84 \| 1 \| 5 \| 147 \| 23 \| 164 \| Lysozyme \| Lysozyme; afdb-uniprot50 \| AF-A0A7K1SLB8-F1-MODEL\_V4 \| 1.0 \| 2.353e-10 \| 456 \| 0.435 \| 147 \| 75 \| 5 \| 2 \| 143 \| 5 \| 148 \| Lysozyme \| Lysozyme; afdb-uniprot50 \| AF-A0A5P1R9I0-F1-MODEL\_V4 \| 1.0 \| 5.493e-10 \| 456 \| 0.349 \| 146 \| 90 \| 4 \| 5 \| 150 \| 7 \| 147 \| Lysozyme \| Lysozyme; afdb-uniprot50 \| AF-A0A1I5WI32-F1-MODEL\_V4 \| 1.0 \| 6.79e-10 \| 456 \| 0.395 \| 144 \| 83 \| 2 \| 1 \| 144 \| 63 \| 202 \| Lysozyme \| Lysozyme; afdb-uniprot50 \| AF-A0A3D3Z792-F1-MODEL\_V4 \| 1.0 \| 5.21e-10 \| 456 \| 0.36 \| 150 \| 88 \| 5 \| 1 \| 146 \| 4 \| 149 \| Lysozyme \| Lysozyme; afdb-uniprot50 \| AF-A0A2D9RBB7-F1-MODEL\_V4 \| 1.0 \| 5.21e-10 \| 456 \| 0.4 \| 150 \| 82 \| 5 \| 1 \| 146 \| 5 \| 150 \| Lysozyme \| Lysozyme; afdb-uniprot50 \| AF-A0A7C9TZG6-F1-MODEL\_V4 \| 1.0 \| 1.008e-10 \| 456 \| 0.39 \| 151 \| 79 \| 4 \| 2 \| 143 \| 79 \| 225 \| Lysozyme \| Lysozyme; afdb-uniprot50 \| AF-A0A443JYE9-F1-MODEL\_V4 \| 1.0 \| 1.094e-09 \| 454 \| 0.333 \| 141 \| 91 \| 2 \| 4 \| 144 \| 1 \| 138 \| Lysozyme \| Lysozyme; afdb-uniprot50 \| AF-A0A397U2F2-F1-MODEL\_V4 \| 1.0 \| 2.909e-10 \| 454 \| 0.376 \| 154 \| 86 \| 4 \| 2 \| 148 \| 9 \| 159 \| Lysozyme \| Lysozyme; afdb-uniprot50 \| AF-A0A3R7C6X1-F1-MODEL\_V4 \| 1.0 \| 1.712e-10 \| 454 \| 0.339 \| 153 \| 89 \| 3 \| 1 \| 146 \| 20 \| 167 \| Lysozyme \| Lysozyme; afdb-uniprot50 \| AF-A0A1A9WZ64-F1-MODEL\_V4 \| 1.0 \| 6.44e-10 \| 454 \| 0.359 \| 139 \| 86 \| 2 \| 7 \| 145 \| 141 \| 276 \| Lysozyme \| Lysozyme; afdb-uniprot50 \| AF-A0A3B0MJC2-F1-MODEL\_V4 \| 1.0 \| 8.85e-10 \| 453 \| 0.34 \| 141 \| 90 \| 2 \| 6 \| 146 \| 76 \| 213 \| Lysozyme \| Lysozyme; afdb-uniprot50 \| AF-L0EV97-F1-MODEL\_V4 \| 1.0 \| 2.232e-10 \| 453 \| 0.429 \| 149 \| 78 \| 4 \| 1 \| 145 \| 9 \| 154 \| Lysozyme \| Lysozyme; afdb-uniprot50 \| AF-A0A828NXT7-F1-MODEL\_V4 \| 1.0 \| 7.549e-10 \| 452 \| 0.362 \| 138 \| 84 \| 3 \| 6 \| 143 \| 2 \| 135 \| Lysozyme \| Lysozyme; afdb-uniprot50 \| AF-C1D7V6-F1-MODEL\_V4 \| 1.0 \| 1.585e-09 \| 452 \| 0.365 \| 145 \| 89 \| 2 \| 1 \| 145 \| 10 \| 151 \| Lysozyme \| Lysozyme; afdb-uniprot50 \| AF-A0A1I7J3V8-F1-MODEL\_V4 \| 1.0 \| 4.444e-10 \| 452 \| 0.398 \| 143 \| 76 \| 5 \| 6 \| 144 \| 14 \| 150 \| Lysozyme \| Lysozyme; afdb-uniprot50 \| AF-A0A850SM26-F1-MODEL\_V4 \| 1.0 \| 1.216e-09 \| 452 \| 0.384 \| 151 \| 84 \| 5 \| 1 \| 147 \| 67 \| 212 \| Lysozyme \| Lysozyme; afdb-uniprot50 \| AF-A0A6B3UKC5-F1-MODEL\_V4 \| 1.0 \| 1.153e-09 \| 452 \| 0.418 \| 148 \| 78 \| 5 \| 1 \| 144 \| 5 \| 148 \| Lysozyme \| Lysozyme; afdb-uniprot50 \| AF-A0A2M9PCB6-F1-MODEL\_V4 \| 1.0 \| 1.037e-09 \| 451 \| 0.39 \| 151 \| 84 \| 3 \| 3 \| 148 \| 1 \| 148 \| Lysozyme \| Lysozyme; afdb-uniprot50 \| AF-A0A7Z2B7B3-F1-MODEL\_V4 \| 1.0 \| 9.839e-10 \| 451 \| 0.397 \| 146 \| 84 \| 3 \| 1 \| 146 \| 1 \| 142 \| Lysozyme \| Lysozyme; afdb-uniprot50 \| AF-A0A318AUS2-F1-MODEL\_V4 \| 1.0 \| 8.85e-10 \| 451 \| 0.374 \| 147 \| 84 \| 5 \| 2 \| 144 \| 6 \| 148 \| Lysozyme \| Lysozyme; afdb-uniprot50 \| AF-A0A7J7BR61-F1-MODEL\_V4 \| 1.0 \| 4.444e-10 \| 451 \| 0.482 \| 145 \| 68 \| 4 \| 2 \| 142 \| 171 \| 312 \| Lysozyme \| Lysozyme; afdb-uniprot50 \| AF-A0A062U607-F1-MODEL\_V4 \| 1.0 \| 5.21e-10 \| 451 \| 0.35 \| 151 \| 89 \| 6 \| 1 \| 146 \| 1 \| 147 \| Lysozyme \| Lysozyme; afdb-uniprot50 \| AF-A0A2E7F2D9-F1-MODEL\_V4 \| 1.0 \| 7.549e-10 \| 451 \| 0.382 \| 149 \| 84 \| 5 \| 2 \| 146 \| 6 \| 150 \| Lysozyme \| Lysozyme; afdb-uniprot50 \| AF-A0A3D3IVC7-F1-MODEL\_V4 \| 1.0 \| 6.44e-10 \| 451 \| 0.376 \| 154 \| 82 \| 5 \| 1 \| 145 \| 11 \| 159 \| Lysozyme \| Lysozyme; afdb-uniprot50 \| AF-V4PU64-F1-MODEL\_V4 \| 1.0 \| 3.41e-10 \| 451 \| 0.415 \| 154 \| 82 \| 5 \| 1 \| 150 \| 5 \| 154 \| Lysozyme \| Lysozyme; afdb-uniprot50 \| AF-A0A1I3ZI12-F1-MODEL\_V4 \| 1.0 \| 1.624e-10 \| 450 \| 0.343 \| 160 \| 99 \| 3 \| 2 \| 155 \| 10 \| 169 \| Lysozyme \| Lysozyme; afdb-uniprot50 \| AF-A0A2D6VSD9-F1-MODEL\_V4 \| 1.0 \| 6.107e-10 \| 450 \| 0.367 \| 147 \| 85 \| 5 \| 3 \| 146 \| 5 \| 146 \| Lysozyme \| Lysozyme; afdb-uniprot50 \| AF-A0A0B1MTD0-F1-MODEL\_V4 \| 1.0 \| 1.352e-09 \| 449 \| 0.382 \| 149 \| 82 \| 4 \| 1 \| 144 \| 1 \| 144 \| Lysozyme \| Lysozyme; afdb-uniprot50 \| AF-A0A0J1CTD0-F1-MODEL\_V4 \| 1.0 \| 1.806e-10 \| 449 \| 0.411 \| 170 \| 74 \| 4 \| 1 \| 147 \| 4 \| 170 \| Lysozyme \| Lysozyme; afdb-uniprot50 \| AF-A0A7V8DKX2-F1-MODEL\_V4 \| 1.0 \| 6.79e-10 \| 449 \| 0.385 \| 148 \| 83 \| 5 \| 3 \| 146 \| 6 \| 149 \| Lysozyme \| Lysozyme; afdb-uniprot50 \| AF-A0A1G4WUT2-F1-MODEL\_V4 \| 1.0 \| 1.959e-09 \| 449 \| 0.34 \| 144 \| 92 \| 2 \| 1 \| 143 \| 201 \| 342 \| Lysozyme \| Lysozyme; afdb-uniprot50 \| AF-A0A2P2E800-F1-MODEL\_V4 \| 1.0 \| 7.549e-10 \| 449 \| 0.366 \| 150 \| 87 \| 5 \| 1 \| 146 \| 4 \| 149 \| Lysozyme \| Lysozyme; afdb-uniprot50 \| AF-K1KHD9-F1-MODEL\_V4 \| 1.0 \| 5.21e-10 \| 448 \| 0.443 \| 142 \| 74 \| 3 \| 6 \| 145 \| 14 \| 152 \| Lysozyme \| Lysozyme; afdb-uniprot50 \| AF-A0A7V4U4L1-F1-MODEL\_V4 \| 1.0 \| 8.85e-10 \| 448 \| 0.305 \| 144 \| 96 \| 3 \| 2 \| 144 \| 15 \| 155 \| Lysozyme \| Lysozyme; afdb-uniprot50 \| AF-A0A645BK91-F1-MODEL\_V4 \| 1.0 \| 8.393e-10 \| 447 \| 0.37 \| 151 \| 82 \| 6 \| 1 \| 143 \| 1 \| 146 \| Lysozyme \| Lysozyme; afdb-uniprot50 \| AF-A0A7C9RJG9-F1-MODEL\_V4 \| 1.0 \| 1.153e-09 \| 447 \| 0.307 \| 140 \| 95 \| 2 \| 5 \| 144 \| 15 \| 152 \| Lysozyme \| Lysozyme; afdb-uniprot50 \| AF-A0A1S1NR31-F1-MODEL\_V4 \| 1.0 \| 1.503e-09 \| 447 \| 0.38 \| 142 \| 81 \| 3 \| 7 \| 144 \| 19 \| 157 \| Lysozyme \| Lysozyme; afdb-uniprot50 \| AF-A0A512NCQ8-F1-MODEL\_V4 \| 1.0 \| 1.426e-09 \| 447 \| 0.417 \| 151 \| 79 \| 6 \| 1 \| 146 \| 3 \| 149 \| Lysozyme \| Lysozyme; afdb-uniprot50 \| AF-A0A354U3U6-F1-MODEL\_V4 \| 1.0 \| 1.216e-09 \| 447 \| 0.38 \| 150 \| 85 \| 5 \| 1 \| 146 \| 5 \| 150 \| Lysozyme \| Lysozyme; afdb-uniprot50 \| AF-A0A849N0U3-F1-MODEL\_V4 \| 1.0 \| 7.549e-10 \| 446 \| 0.335 \| 149 \| 94 \| 3 \| 1 \| 145 \| 1 \| 148 \| Lysozyme \| Lysozyme; afdb-uniprot50 \| AF-A0A833N5J4-F1-MODEL\_V4 \| 1.0 \| 1.585e-09 \| 445 \| 0.365 \| 145 \| 89 \| 2 \| 1 \| 145 \| 2 \| 143 \| Lysozyme \| Lysozyme; afdb-uniprot50 \| AF-U3TZ86-F1-MODEL\_V4 \| 1.0 \| 6.44e-10 \| 445 \| 0.35 \| 140 \| 86 \| 3 \| 6 \| 143 \| 21 \| 157 \| Lysozyme \| Lysozyme; afdb-uniprot50 \| AF-C5A7Z2-F1-MODEL\_V4 \| 1.0 \| 9.067e-11 \| 445 \| 0.357 \| 193 \| 73 \| 6 \| 4 \| 145 \| 58 \| 250 \| Lysozyme \| Lysozyme; afdb-uniprot50 \| AF-A0A1B1ALJ1-F1-MODEL\_V4 \| 1.0 \| 5.21e-10 \| 445 \| 0.342 \| 146 \| 88 \| 4 \| 3 \| 144 \| 5 \| 146 \| Lysozyme \| Lysozyme; afdb-uniprot50 \| AF-A0A096BIF3-F1-MODEL\_V4 \| 1.0 \| 6.79e-10 \| 444 \| 0.41 \| 146 \| 79 \| 4 \| 1 \| 143 \| 3 \| 144 \| Lysozyme \| Lysozyme; afdb-uniprot50 \| AF-A0A2S4E741-F1-MODEL\_V4 \| 1.0 \| 3.791e-10 \| 444 \| 0.418 \| 153 \| 75 \| 6 \| 2 \| 144 \| 3 \| 151 \| Lysozyme \| Lysozyme; afdb-uniprot50 \| AF-A0A7V5SPV2-F1-MODEL\_V4 \| 1.0 \| 3.595e-10 \| 444 \| 0.36 \| 147 \| 85 \| 5 \| 1 \| 143 \| 22 \| 163 \| Lysozyme \| Lysozyme; afdb-uniprot50 \| AF-A0A7V8A8R5-F1-MODEL\_V4 \| 1.0 \| 8.393e-10 \| 444 \| 0.385 \| 148 \| 83 \| 5 \| 3 \| 146 \| 6 \| 149 \| Lysozyme \| Lysozyme; afdb-uniprot50 \| AF-R9LWU8-F1-MODEL\_V4 \| 1.0 \| 3.234e-10 \| 444 \| 0.408 \| 147 \| 77 \| 4 \| 1 \| 143 \| 3 \| 143 \| Lysozyme \| Lysozyme; afdb-uniprot50 \| AF-A0A7C5PMR5-F1-MODEL\_V4 \| 1.0 \| 7.549e-10 \| 444 \| 0.433 \| 157 \| 76 \| 5 \| 1 \| 148 \| 1 \| 153 \| Lysozyme \| Lysozyme; afdb-uniprot50 \| AF-A0A4R2VIK8-F1-MODEL\_V4 \| 1.0 \| 6.44e-10 \| 444 \| 0.428 \| 154 \| 77 \| 5 \| 1 \| 146 \| 68 \| 218 \| Lysozyme \| Lysozyme; afdb-uniprot50 \| AF-F5RAK7-F1-MODEL\_V4 \| 1.0 \| 3.234e-10 \| 444 \| 0.395 \| 149 \| 82 \| 5 \| 1 \| 145 \| 96 \| 240 \| Lysozyme \| Lysozyme; afdb-uniprot50 \| AF-R6FSL7-F1-MODEL\_V4 \| 1.0 \| 7.16e-10 \| 444 \| 0.406 \| 145 \| 80 \| 4 \| 1 \| 144 \| 1 \| 140 \| Lysozyme \| Lysozyme; afdb-uniprot50 \| AF-A0A059DT11-F1-MODEL\_V4 \| 1.0 \| 1.216e-09 \| 444 \| 0.355 \| 149 \| 87 \| 6 \| 1 \| 144 \| 5 \| 149 \| Lysozyme \| Lysozyme; afdb-uniprot50 \| AF-A0A2D7XZM4-F1-MODEL\_V4 \| 1.0 \| 1.216e-09 \| 444 \| 0.393 \| 150 \| 83 \| 5 \| 1 \| 146 \| 5 \| 150 \| Lysozyme \| Lysozyme; afdb-uniprot50 \| AF-A0A6P0NKT7-F1-MODEL\_V4 \| 1.0 \| 5.493e-10 \| 443 \| 0.363 \| 154 \| 87 \| 5 \| 1 \| 146 \| 5 \| 155 \| Lysozyme \| Lysozyme; afdb-uniprot50 \| AF-I4FUS8-F1-MODEL\_V4 \| 1.0 \| 2.481e-10 \| 443 \| 0.389 \| 167 \| 76 \| 5 \| 1 \| 144 \| 25 \| 188 \| Lysozyme \| Lysozyme; afdb-uniprot50 \| AF-A0A2G2IYC1-F1-MODEL\_V4 \| 1.0 \| 5.493e-10 \| 443 \| 0.396 \| 154 \| 78 \| 6 \| 1 \| 143 \| 4 \| 153 \| Lysozyme \| Lysozyme; afdb-uniprot50 \| AF-A0A2E2QM79-F1-MODEL\_V4 \| 1.0 \| 5.21e-10 \| 442 \| 0.404 \| 163 \| 75 \| 4 \| 1 \| 145 \| 3 \| 161 \| Lysozyme \| Lysozyme; afdb-uniprot50 \| AF-A0A5C7NLG2-F1-MODEL\_V4 \| 1.0 \| 1.153e-09 \| 442 \| 0.371 \| 159 \| 83 \| 4 \| 2 \| 146 \| 20 \| 175 \| Lysozyme \| Lysozyme; afdb-uniprot50 \| AF-A0A2R4NSE3-F1-MODEL\_V4 \| 1.0 \| 7.16e-10 \| 441 \| 0.412 \| 148 \| 79 \| 4 \| 1 \| 144 \| 1 \| 144 \| Lysozyme \| Lysozyme; afdb-uniprot50 \| AF-A0A7V8JUE6-F1-MODEL\_V4 \| 1.0 \| 1.153e-09 \| 441 \| 0.349 \| 143 \| 89 \| 2 \| 5 \| 147 \| 4 \| 142 \| Lysozyme \| Lysozyme; afdb-uniprot50 \| AF-A0A6P2LU80-F1-MODEL\_V4 \| 1.0 \| 3.41e-10 \| 440 \| 0.371 \| 159 \| 86 \| 5 \| 1 \| 148 \| 3 \| 158 \| Lysozyme \| Lysozyme; afdb-uniprot50 \| AF-A0A348HDV6-F1-MODEL\_V4 \| 1.0 \| 1.858e-09 \| 440 \| 0.344 \| 145 \| 88 \| 3 \| 5 \| 145 \| 15 \| 156 \| Lysozyme \| Lysozyme; afdb-uniprot50 \| AF-A0A3S1C1X9-F1-MODEL\_V4 \| 1.0 \| 4.215e-10 \| 440 \| 0.385 \| 153 \| 83 \| 3 \| 1 \| 145 \| 61 \| 210 \| Lysozyme \| Lysozyme; afdb-uniprot50 \| AF-A0A850BAY3-F1-MODEL\_V4 \| 1.0 \| 3.791e-10 \| 440 \| 0.398 \| 158 \| 77 \| 5 \| 1 \| 143 \| 738 \| 892 \| Type VI secretion system tip protein VgrG \| Type VI secretion system tip protein VgrG; afdb-uniprot50 \| AF-A0A0F3Q9P1-F1-MODEL\_V4 \| 1.0 \| 1.094e-09 \| 439 \| 0.337 \| 145 \| 90 \| 4 \| 6 \| 147 \| 6 \| 147 \| Lysozyme \| Lysozyme; afdb-uniprot50 \| AF-A0A1M3KMN9-F1-MODEL\_V4 \| 1.0 \| 1.352e-09 \| 439 \| 0.393 \| 150 \| 84 \| 3 \| 1 \| 143 \| 1 \| 150 \| Lysozyme \| Lysozyme; afdb-uniprot50 \| AF-A0A839LRE8-F1-MODEL\_V4 \| 1.0 \| 3.41e-10 \| 438 \| 0.325 \| 160 \| 102 \| 3 \| 2 \| 155 \| 24 \| 183 \| Lysozyme \| Lysozyme; afdb-uniprot50 \| AF-A0A2E4QSB6-F1-MODEL\_V4 \| 1.0 \| 1.426e-09 \| 438 \| 0.331 \| 148 \| 91 \| 5 \| 1 \| 144 \| 5 \| 148 \| Lysozyme \| Lysozyme; afdb-uniprot50 \| AF-A0A4Q3UUN7-F1-MODEL\_V4 \| 1.0 \| 1.282e-09 \| 437 \| 0.422 \| 142 \| 74 \| 5 \| 1 \| 138 \| 5 \| 142 \| Lysozyme \| Lysozyme; afdb-uniprot50 \| AF-A0A4Q0QES9-F1-MODEL\_V4 \| 1.0 \| 7.549e-10 \| 437 \| 0.522 \| 132 \| 57 \| 2 \| 18 \| 146 \| 7 \| 135 \| Lysozyme \| Lysozyme; afdb-uniprot50 \| AF-Q2NSF6-F1-MODEL\_V4 \| 1.0 \| 1.426e-09 \| 437 \| 0.384 \| 130 \| 77 \| 2 \| 7 \| 136 \| 21 \| 147 \| Lysozyme \| Lysozyme; afdb-uniprot50 \| AF-A0A1W1V408-F1-MODEL\_V4 \| 1.0 \| 1.216e-09 \| 437 \| 0.339 \| 153 \| 88 \| 4 \| 1 \| 143 \| 2 \| 151 \| Lysozyme \| Lysozyme; afdb-uniprot50 \| AF-A0A7Y0GSW2-F1-MODEL\_V4 \| 1.0 \| 3.7e-09 \| 437 \| 0.363 \| 146 \| 92 \| 1 \| 1 \| 146 \| 13 \| 157 \| Lysozyme \| Lysozyme; afdb-uniprot50 \| AF-A0A4P7L828-F1-MODEL\_V4 \| 1.0 \| 2.297e-09 \| 437 \| 0.361 \| 141 \| 87 \| 2 \| 6 \| 146 \| 32 \| 169 \| Lysozyme \| Lysozyme; afdb-uniprot50 \| AF-A0A829TCL8-F1-MODEL\_V4 \| 1.0 \| 3.157e-09 \| 436 \| 0.364 \| 137 \| 84 \| 2 \| 11 \| 147 \| 2 \| 135 \| Putative phage lysozyme \| Putative phage lysozyme; afdb-uniprot50 \| AF-B1M176-F1-MODEL\_V4 \| 1.0 \| 2.554e-09 \| 436 \| 0.328 \| 140 \| 93 \| 1 \| 5 \| 144 \| 55 \| 193 \| Lysozyme \| Lysozyme; afdb-uniprot50 \| AF-A0A0N0JEL9-F1-MODEL\_V4 \| 1.0 \| 4.215e-10 \| 436 \| 0.372 \| 180 \| 83 \| 7 \| 1 \| 153 \| 10 \| 186 \| Lysozyme \| Lysozyme; afdb-uniprot50 \| AF-A0A7W9CFD1-F1-MODEL\_V4 \| 1.0 \| 1.426e-09 \| 436 \| 0.314 \| 159 \| 97 \| 4 \| 1 \| 154 \| 9 \| 160 \| Lysozyme \| Lysozyme; afdb-uniprot50 \| AF-A0A7W6JCA7-F1-MODEL\_V4 \| 1.0 \| 1.503e-09 \| 436 \| 0.333 \| 159 \| 95 \| 5 \| 1 \| 154 \| 9 \| 161 \| Lysozyme \| Lysozyme; afdb-uniprot50 \| AF-A0A829EU62-F1-MODEL\_V4 \| 1.0 \| 1.503e-09 \| 436 \| 0.308 \| 149 \| 96 \| 4 \| 1 \| 145 \| 1 \| 146 \| Uncharacterized protein \| Uncharacterized protein; afdb-uniprot50 \| AF-A0A1M7CNB5-F1-MODEL\_V4 \| 1.0 \| 1.426e-09 \| 435 \| 0.387 \| 147 \| 84 \| 5 \| 1 \| 146 \| 1 \| 142 \| Lysozyme \| Lysozyme; afdb-uniprot50 \| AF-A0A840G3M7-F1-MODEL\_V4 \| 1.0 \| 1.585e-09 \| 435 \| 0.414 \| 147 \| 81 \| 3 \| 1 \| 146 \| 4 \| 146 \| Lysozyme \| Lysozyme; afdb-uniprot50 \| AF-K1JIE7-F1-MODEL\_V4 \| 1.0 \| 4.215e-10 \| 435 \| 0.381 \| 152 \| 84 \| 5 \| 1 \| 145 \| 1 \| 149 \| Lysozyme \| Lysozyme; afdb-uniprot50 \| AF-A0A847VHY1-F1-MODEL\_V4 \| 1.0 \| 1.426e-09 \| 435 \| 0.323 \| 142 \| 92 \| 3 \| 5 \| 145 \| 22 \| 160 \| Lysozyme \| Lysozyme; afdb-uniprot50 \| AF-A0A4Q3SIK8-F1-MODEL\_V4 \| 1.0 \| 1.094e-09 \| 435 \| 0.348 \| 152 \| 87 \| 6 \| 1 \| 144 \| 1 \| 148 \| Lysozyme \| Lysozyme; afdb-uniprot50 \| AF-A0A420WVM4-F1-MODEL\_V4 \| 1.0 \| 1.503e-09 \| 435 \| 0.385 \| 148 \| 82 \| 4 \| 4 \| 145 \| 16 \| 160 \| Lysozyme \| Lysozyme; afdb-uniprot50 \| AF-A0A514ETB6-F1-MODEL\_V4 \| 1.0 \| 1.153e-09 \| 435 \| 0.317 \| 145 \| 91 \| 3 \| 6 \| 145 \| 21 \| 162 \| Lysozyme \| Lysozyme; afdb-uniprot50 \| AF-A0A257J4H1-F1-MODEL\_V4 \| 1.0 \| 1.503e-09 \| 435 \| 0.346 \| 147 \| 88 \| 5 \| 2 \| 144 \| 6 \| 148 \| Lysozyme \| Lysozyme; afdb-uniprot50 \| AF-A0A2N6KSA2-F1-MODEL\_V4 \| 1.0 \| 5.493e-10 \| 435 \| 0.369 \| 149 \| 84 \| 4 \| 3 \| 147 \| 131 \| 273 \| Lysozyme \| Lysozyme; afdb-uniprot50 \| AF-A0A601HWC6-F1-MODEL\_V4 \| 1.0 \| 9.839e-10 \| 434 \| 0.554 \| 128 \| 53 \| 2 \| 21 \| 144 \| 1 \| 128 \| Lysozyme \| Lysozyme; afdb-uniprot50 \| AF-A0A4Q3S555-F1-MODEL\_V4 \| 1.0 \| 2.297e-09 \| 434 \| 0.374 \| 147 \| 84 \| 5 \| 2 \| 144 \| 6 \| 148 \| Lysozyme \| Lysozyme; afdb-uniprot50 \| AF-A0A7R9QUH5-F1-MODEL\_V4 \| 1.0 \| 5.493e-10 \| 434 \| 0.405 \| 158 \| 84 \| 6 \| 1 \| 153 \| 18 \| 170 \| Lysozyme \| Lysozyme; afdb-uniprot50 \| AF-E0TGL6-F1-MODEL\_V4 \| 1.0 \| 6.44e-10 \| 434 \| 0.402 \| 159 \| 77 \| 5 \| 1 \| 144 \| 1 \| 156 \| Lysozyme \| Lysozyme; afdb-uniprot50 \| AF-A0A5B2VH29-F1-MODEL\_V4 \| 1.0 \| 8.85e-10 \| 432 \| 0.378 \| 148 \| 86 \| 4 \| 1 \| 144 \| 6 \| 151 \| Lysozyme \| Lysozyme; afdb-uniprot50 \| AF-A0A5C4PLN9-F1-MODEL\_V4 \| 1.0 \| 3.41e-10 \| 432 \| 0.37 \| 154 \| 85 \| 7 \| 1 \| 145 \| 1 \| 151 \| Lysozyme \| Lysozyme; afdb-uniprot50 \| AF-A0A1S9AW89-F1-MODEL\_V4 \| 1.0 \| 5.21e-10 \| 432 \| 0.362 \| 174 \| 80 \| 5 \| 1 \| 146 \| 1 \| 171 \| Lysozyme \| Lysozyme; afdb-uniprot50 \| AF-A0A2G2MTD8-F1-MODEL\_V4 \| 1.0 \| 9.839e-10 \| 432 \| 0.364 \| 151 \| 88 \| 5 \| 1 \| 147 \| 5 \| 151 \| Lysozyme \| Lysozyme; afdb-uniprot50 \| AF-A0A1V5WW45-F1-MODEL\_V4 \| 1.0 \| 3.7e-09 \| 431 \| 0.363 \| 146 \| 88 \| 4 \| 2 \| 146 \| 5 \| 146 \| Lysozyme \| Lysozyme; afdb-uniprot50 \| AF-A0A2H0QTW7-F1-MODEL\_V4 \| 1.0 \| 7.96e-10 \| 431 \| 0.331 \| 169 \| 87 \| 5 \| 1 \| 146 \| 2 \| 167 \| Lysozyme \| Lysozyme; afdb-uniprot50 \| AF-Q87BX4-F1-MODEL\_V4 \| 1.0 \| 1.153e-09 \| 431 \| 0.472 \| 129 \| 63 \| 4 \| 2 \| 128 \| 19 \| 144 \| Lysozyme \| Lysozyme; afdb-uniprot50 \| AF-A0A6M0HX53-F1-MODEL\_V4 \| 1.0 \| 7.549e-10 \| 431 \| 0.4 \| 150 \| 78 \| 5 \| 1 \| 146 \| 1 \| 142 \| Lysozyme \| Lysozyme; afdb-uniprot50 \| AF-A0A6G7CH63-F1-MODEL\_V4 \| 1.0 \| 4.338e-09 \| 430 \| 0.306 \| 137 \| 92 \| 2 \| 7 \| 143 \| 22 \| 155 \| Lysozyme \| Lysozyme; afdb-uniprot50 \| AF-A0A846BN22-F1-MODEL\_V4 \| 1.0 \| 3.791e-10 \| 430 \| 0.375 \| 152 \| 82 \| 4 \| 2 \| 143 \| 27 \| 175 \| Lysozyme \| Lysozyme; afdb-uniprot50 \| AF-A0A7R9M991-F1-MODEL\_V4 \| 1.0 \| 1.858e-09 \| 429 \| 0.373 \| 150 \| 82 \| 5 \| 1 \| 144 \| 19 \| 162 \| Lysozyme \| Lysozyme; afdb-uniprot50 \| AF-A0A7X9HPJ2-F1-MODEL\_V4 \| 1.0 \| 6.107e-10 \| 429 \| 0.364 \| 159 \| 82 \| 4 \| 2 \| 145 \| 59 \| 213 \| Lysozyme \| Lysozyme; afdb-uniprot50 \| AF-A3XEZ7-F1-MODEL\_V4 \| 1.0 \| 2.297e-09 \| 429 \| 0.358 \| 145 \| 88 \| 4 \| 1 \| 144 \| 94 \| 234 \| Lysozyme \| Lysozyme; afdb-uniprot50 \| AF-I0HR64-F1-MODEL\_V4 \| 1.0 \| 4.444e-10 \| 429 \| 0.381 \| 181 \| 81 \| 6 \| 1 \| 153 \| 6 \| 183 \| Lysozyme \| Lysozyme; afdb-uniprot50 \| AF-A0A1Z4LWR5-F1-MODEL\_V4 \| 1.0 \| 6.79e-10 \| 429 \| 0.379 \| 153 \| 82 \| 4 \| 1 \| 144 \| 105 \| 253 \| Lysozyme \| Lysozyme; afdb-uniprot50 \| AF-A0A3P2RCD3-F1-MODEL\_V4 \| 1.0 \| 6.44e-10 \| 429 \| 0.395 \| 149 \| 84 \| 5 \| 1 \| 147 \| 1 \| 145 \| Lysozyme \| Lysozyme; afdb-uniprot50 \| AF-A0A2A7S155-F1-MODEL\_V4 \| 1.0 \| 2.232e-10 \| 429 \| 0.336 \| 196 \| 78 \| 5 \| 1 \| 145 \| 77 \| 271 \| Lysozyme \| Lysozyme; afdb-uniprot50 \| AF-A0A7C4STG1-F1-MODEL\_V4 \| 1.0 \| 6.44e-10 \| 429 \| 0.351 \| 165 \| 86 \| 5 \| 1 \| 150 \| 204 \| 362 \| Lysozyme \| Lysozyme; afdb-uniprot50 \| AF-A0A399R703-F1-MODEL\_V4 \| 1.0 \| 2.839e-09 \| 429 \| 0.35 \| 151 \| 89 \| 6 \| 1 \| 146 \| 5 \| 151 \| Lysozyme \| Lysozyme; afdb-uniprot50 \| AF-R0CYV1-F1-MODEL\_V4 \| 1.0 \| 2.994e-09 \| 429 \| 0.367 \| 147 \| 85 \| 5 \| 2 \| 144 \| 6 \| 148 \| Lysozyme \| Lysozyme; afdb-uniprot50 \| AF-A0A7X7AHC1-F1-MODEL\_V4 \| 1.0 \| 2.839e-09 \| 428 \| 0.354 \| 141 \| 85 \| 4 \| 6 \| 144 \| 10 \| 146 \| Lysozyme \| Lysozyme; afdb-uniprot50 \| AF-A0A8B2QY50-F1-MODEL\_V4 \| 1.0 \| 9.331e-10 \| 428 \| 0.304 \| 148 \| 90 \| 3 \| 7 \| 144 \| 2 \| 146 \| Lysozyme \| Lysozyme; afdb-uniprot50 \| AF-A0A259NPH4-F1-MODEL\_V4 \| 1.0 \| 3.234e-10 \| 428 \| 0.392 \| 158 \| 76 \| 4 \| 4 \| 143 \| 1 \| 156 \| Lysozyme \| Lysozyme; afdb-uniprot50 \| AF-A0A258GXH3-F1-MODEL\_V4 \| 1.0 \| 2.994e-09 \| 428 \| 0.347 \| 144 \| 89 \| 2 \| 3 \| 146 \| 48 \| 186 \| Lysozyme \| Lysozyme; afdb-uniprot50 \| AF-A0A1M3KKZ9-F1-MODEL\_V4 \| 1.0 \| 3.997e-10 \| 428 \| 0.383 \| 159 \| 83 \| 6 \| 1 \| 148 \| 6 \| 160 \| Lysozyme \| Lysozyme; afdb-uniprot50 \| AF-A0A7I8ZS27-F1-MODEL\_V4 \| 1.0 \| 1.503e-09 \| 428 \| 0.317 \| 145 \| 91 \| 3 \| 6 \| 145 \| 21 \| 162 \| Lysozyme \| Lysozyme; afdb-uniprot50 \| AF-A0A7W1LGJ0-F1-MODEL\_V4 \| 1.0 \| 5.21e-10 \| 428 \| 0.409 \| 154 \| 78 \| 4 \| 2 \| 146 \| 238 \| 387 \| Lysozyme \| Lysozyme; afdb-uniprot50 \| AF-A0A7W0G9B2-F1-MODEL\_V4 \| 1.0 \| 4.941e-10 \| 428 \| 0.385 \| 153 \| 83 \| 3 \| 2 \| 146 \| 321 \| 470 \| Lysozyme \| Lysozyme; afdb-uniprot50 \| AF-A0A421BJ35-F1-MODEL\_V4 \| 1.0 \| 4.574e-09 \| 427 \| 0.364 \| 140 \| 85 \| 3 \| 5 \| 144 \| 9 \| 144 \| Lysozyme \| Lysozyme; afdb-uniprot50 \| AF-G4QCV9-F1-MODEL\_V4 \| 1.0 \| 1.762e-09 \| 427 \| 0.356 \| 146 \| 86 \| 5 \| 4 \| 146 \| 12 \| 152 \| Lysozyme \| Lysozyme; afdb-uniprot50 \| AF-A0A060H052-F1-MODEL\_V4 \| 1.0 \| 5.21e-10 \| 427 \| 0.331 \| 148 \| 89 \| 3 \| 5 \| 145 \| 32 \| 176 \| Lysozyme \| Lysozyme; afdb-uniprot50 \| AF-A0A1I2RRC9-F1-MODEL\_V4 \| 1.0 \| 8.85e-10 \| 427 \| 0.421 \| 147 \| 73 \| 6 \| 1 \| 143 \| 1 \| 139 \| Lysozyme \| Lysozyme; afdb-uniprot50 \| AF-A0A226ESI2-F1-MODEL\_V4 \| 1.0 \| 9.839e-10 \| 427 \| 0.371 \| 159 \| 86 \| 7 \| 1 \| 148 \| 23 \| 178 \| Lysozyme \| Lysozyme; afdb-uniprot50 \| AF-A0A2D7IXU0-F1-MODEL\_V4 \| 1.0 \| 3.902e-09 \| 427 \| 0.386 \| 150 \| 84 \| 5 \| 1 \| 146 \| 5 \| 150 \| Lysozyme \| Lysozyme; afdb-uniprot50 \| AF-D9Y906-F1-MODEL\_V4 \| 1.0 \| 1.503e-09 \| 426 \| 0.376 \| 146 \| 82 \| 5 \| 6 \| 146 \| 2 \| 143 \| Lysozyme \| Lysozyme; afdb-uniprot50 \| AF-A0A4Y5YHV6-F1-MODEL\_V4 \| 1.0 \| 3.328e-09 \| 426 \| 0.352 \| 142 \| 87 \| 4 \| 6 \| 146 \| 18 \| 155 \| Lysozyme \| Lysozyme; afdb-uniprot50 \| AF-A0A7R9PWQ4-F1-MODEL\_V4 \| 1.0 \| 2.297e-09 \| 426 \| 0.377 \| 151 \| 81 \| 6 \| 1 \| 144 \| 18 \| 162 \| Lysozyme \| Lysozyme; afdb-uniprot50 \| AF-A0A4R2MG20-F1-MODEL\_V4 \| 1.0 \| 2.481e-10 \| 426 \| 0.37 \| 181 \| 81 \| 7 \| 1 \| 152 \| 16 \| 192 \| Lysozyme \| Lysozyme; afdb-uniprot50 \| AF-A0A1D2MI83-F1-MODEL\_V4 \| 1.0 \| 1.671e-09 \| 426 \| 0.364 \| 159 \| 86 \| 6 \| 1 \| 147 \| 70 \| 225 \| Lysozyme \| Lysozyme; afdb-uniprot50 \| AF-A0A1M7K5Y8-F1-MODEL\_V4 \| 1.0 \| 7.369e-09 \| 425 \| 0.328 \| 140 \| 91 \| 2 \| 7 \| 146 \| 21 \| 157 \| Lysozyme \| Lysozyme; afdb-uniprot50 \| AF-A0A7Y3N265-F1-MODEL\_V4 \| 1.0 \| 5.085e-09 \| 425 \| 0.352 \| 139 \| 87 \| 2 \| 5 \| 143 \| 39 \| 174 \| Lysozyme \| Lysozyme; afdb-uniprot50 \| AF-A0A7W0G4G5-F1-MODEL\_V4 \| 1.0 \| 6.79e-10 \| 425 \| 0.284 \| 169 \| 103 \| 4 \| 3 \| 155 \| 16 \| 182 \| Lysozyme \| Lysozyme; afdb-uniprot50 \| AF-A0A857C5S3-F1-MODEL\_V4 \| 1.0 \| 2.297e-09 \| 425 \| 0.333 \| 141 \| 88 \| 4 \| 6 \| 144 \| 17 \| 153 \| Glycoside hydrolase family protein \| Glycoside hydrolase family protein; afdb-uniprot50 \| AF-A0A125SM95-F1-MODEL\_V4 \| 1.0 \| 9.839e-10 \| 425 \| 0.384 \| 169 \| 78 \| 4 \| 1 \| 146 \| 49 \| 214 \| Lysozyme \| Lysozyme; afdb-uniprot50 \| AF-S3JF93-F1-MODEL\_V4 \| 1.0 \| 9.839e-10 \| 425 \| 0.389 \| 167 \| 76 \| 5 \| 1 \| 144 \| 114 \| 277 \| Lysozyme \| Lysozyme; afdb-uniprot50 \| AF-A0A7Z1R6F5-F1-MODEL\_V4 \| 1.0 \| 3.7e-09 \| 425 \| 0.377 \| 143 \| 79 \| 5 \| 6 \| 144 \| 177 \| 313 \| Lysozyme \| Lysozyme; afdb-uniprot50 \| AF-A0A0A2V9H7-F1-MODEL\_V4 \| 1.0 \| 2.839e-09 \| 425 \| 0.302 \| 149 \| 98 \| 3 \| 1 \| 146 \| 1 \| 146 \| Lysozyme \| Lysozyme; afdb-uniprot50 \| AF-E2YNY1-F1-MODEL\_V4 \| 1.0 \| 1.858e-09 \| 425 \| 0.313 \| 153 \| 97 \| 5 \| 1 \| 148 \| 1 \| 150 \| Lysozyme \| Lysozyme; afdb-uniprot50 \| AF-A0A525KSI6-F1-MODEL\_V4 \| 1.0 \| 4.114e-09 \| 425 \| 0.34 \| 147 \| 89 \| 5 \| 2 \| 144 \| 6 \| 148 \| Lysozyme \| Lysozyme; afdb-uniprot50 \| AF-A0A357L761-F1-MODEL\_V4 \| 1.0 \| 4.338e-09 \| 424 \| 0.353 \| 150 \| 88 \| 5 \| 1 \| 146 \| 38 \| 182 \| Lysozyme \| Lysozyme; afdb-uniprot50 \| AF-A0A327X3T2-F1-MODEL\_V4 \| 1.0 \| 4.823e-09 \| 424 \| 0.357 \| 140 \| 86 \| 3 \| 5 \| 144 \| 19 \| 154 \| Lysozyme \| Lysozyme; afdb-uniprot50 \| AF-A0A443L9N3-F1-MODEL\_V4 \| 1.0 \| 4.574e-09 \| 424 \| 0.364 \| 140 \| 85 \| 3 \| 5 \| 144 \| 104 \| 239 \| Lysozyme \| Lysozyme; afdb-uniprot50 \| AF-A0A7V8YU39-F1-MODEL\_V4 \| 1.0 \| 5.21e-10 \| 424 \| 0.405 \| 153 \| 78 \| 4 \| 2 \| 145 \| 112 \| 260 \| Lysozyme \| Lysozyme; afdb-uniprot50 \| AF-A0A7W9E8M6-F1-MODEL\_V4 \| 1.0 \| 1.762e-09 \| 424 \| 0.3 \| 163 \| 99 \| 6 \| 1 \| 154 \| 1 \| 157 \| Lysozyme \| Lysozyme; afdb-uniprot50 \| AF-A0A525JRD6-F1-MODEL\_V4 \| 1.0 \| 3.328e-09 \| 424 \| 0.364 \| 151 \| 88 \| 5 \| 2 \| 148 \| 6 \| 152 \| Lysozyme \| Lysozyme; afdb-uniprot50 \| AF-A0A7G9RLL0-F1-MODEL\_V4 \| 1.0 \| 2.693e-09 \| 423 \| 0.372 \| 145 \| 83 \| 4 \| 3 \| 143 \| 11 \| 151 \| Lysozyme \| Lysozyme; afdb-uniprot50 \| AF-A0A0F9RCT0-F1-MODEL\_V4 \| 1.0 \| 1.585e-09 \| 423 \| 0.342 \| 152 \| 88 \| 3 \| 1 \| 143 \| 1 \| 149 \| Lysozyme \| Lysozyme; afdb-uniprot50 \| AF-A0A397UIN9-F1-MODEL\_V4 \| 1.0 \| 1.585e-09 \| 423 \| 0.374 \| 155 \| 86 \| 4 \| 2 \| 148 \| 177 \| 328 \| Lysozyme \| Lysozyme; afdb-uniprot50 \| AF-A0A2G2LIQ0-F1-MODEL\_V4 \| 1.0 \| 2.422e-09 \| 423 \| 0.358 \| 148 \| 87 \| 5 \| 3 \| 146 \| 6 \| 149 \| Lysozyme \| Lysozyme; afdb-uniprot50 \| AF-A0A7M5XGT8-F1-MODEL\_V4 \| 1.0 \| 2.839e-09 \| 422 \| 0.46 \| 128 \| 66 \| 2 \| 2 \| 129 \| 5 \| 129 \| Lysozyme \| Lysozyme; afdb-uniprot50 \| AF-A0A009FLU0-F1-MODEL\_V4 \| 1.0 \| 1.037e-09 \| 422 \| 0.518 \| 133 \| 57 \| 4 \| 18 \| 146 \| 1 \| 130 \| Lysozyme \| Lysozyme; afdb-uniprot50 \| AF-A0A2N9LSD0-F1-MODEL\_V4 \| 1.0 \| 5.085e-09 \| 422 \| 0.391 \| 148 \| 82 \| 5 \| 1 \| 144 \| 1 \| 144 \| Lysozyme \| Lysozyme; afdb-uniprot50 \| AF-C7RLP6-F1-MODEL\_V4 \| 1.0 \| 7.16e-10 \| 422 \| 0.378 \| 169 \| 75 \| 5 \| 1 \| 144 \| 4 \| 167 \| Lysozyme \| Lysozyme; afdb-uniprot50 \| AF-A0A845WUP1-F1-MODEL\_V4 \| 1.0 \| 1.503e-09 \| 422 \| 0.33 \| 142 \| 87 \| 4 \| 5 \| 144 \| 49 \| 184 \| Glycoside hydrolase family protein \| Glycoside hydrolase family protein; afdb-uniprot50 \| AF-S4FZ78-F1-MODEL\_V4 \| 1.0 \| 3.7e-09 \| 422 \| 0.31 \| 148 \| 95 \| 4 \| 1 \| 144 \| 1 \| 145 \| Lysozyme \| Lysozyme; afdb-uniprot50 \| AF-A0A0S9NGX8-F1-MODEL\_V4 \| 1.0 \| 1.671e-09 \| 422 \| 0.312 \| 157 \| 97 \| 5 \| 3 \| 154 \| 1 \| 151 \| Lysozyme \| Lysozyme; afdb-uniprot50 \| AF-A0A6I5R3J3-F1-MODEL\_V4 \| 1.0 \| 4.444e-10 \| 422 \| 0.368 \| 152 \| 88 \| 5 \| 4 \| 151 \| 219 \| 366 \| Lysozyme \| Lysozyme; afdb-uniprot50 \| AF-A0A329A2Y0-F1-MODEL\_V4 \| 1.0 \| 1.426e-09 \| 421 \| 0.36 \| 158 \| 84 \| 4 \| 1 \| 144 \| 1 \| 155 \| Lysozyme \| Lysozyme; afdb-uniprot50 \| AF-A0A1V5YWE3-F1-MODEL\_V4 \| 1.0 \| 1.585e-09 \| 421 \| 0.385 \| 148 \| 85 \| 4 \| 1 \| 147 \| 1 \| 143 \| Lysozyme \| Lysozyme; afdb-uniprot50 \| AF-A0A1Y1YA73-F1-MODEL\_V4 \| 1.0 \| 1.959e-09 \| 421 \| 0.375 \| 149 \| 81 \| 5 \| 2 \| 143 \| 84 \| 227 \| Lysozyme \| Lysozyme; afdb-uniprot50 \| AF-A0A847VEY3-F1-MODEL\_V4 \| 1.0 \| 4.338e-09 \| 421 \| 0.412 \| 148 \| 79 \| 5 \| 3 \| 146 \| 11 \| 154 \| Glycoside hydrolase family protein \| Glycoside hydrolase family protein; afdb-uniprot50 \| AF-A0A136H867-F1-MODEL\_V4 \| 1.0 \| 7.77e-09 \| 420 \| 0.342 \| 140 \| 88 \| 3 \| 5 \| 144 \| 10 \| 145 \| Lysozyme \| Lysozyme; afdb-uniprot50 \| AF-A0A6B3LIZ4-F1-MODEL\_V4 \| 1.0 \| 2.178e-09 \| 420 \| 0.375 \| 149 \| 82 \| 3 \| 4 \| 145 \| 29 \| 173 \| Lysozyme \| Lysozyme; afdb-uniprot50 \| AF-A0A520R102-F1-MODEL\_V4 \| 1.0 \| 9.108e-09 \| 420 \| 0.342 \| 146 \| 88 \| 4 \| 1 \| 144 \| 69 \| 208 \| Lysozyme \| Lysozyme; afdb-uniprot50 \| AF-A0A7W6U5N3-F1-MODEL\_V4 \| 1.0 \| 1.858e-09 \| 420 \| 0.358 \| 170 \| 94 \| 4 \| 1 \| 159 \| 52 \| 217 \| Lysozyme \| Lysozyme; afdb-uniprot50 \| AF-A0A7T9XKC1-F1-MODEL\_V4 \| 1.0 \| 1.013e-08 \| 419 \| 0.328 \| 140 \| 91 \| 2 \| 7 \| 146 \| 21 \| 157 \| Lysozyme \| Lysozyme; afdb-uniprot50 \| AF-A0A2D8XMB5-F1-MODEL\_V4 \| 1.0 \| 2.178e-09 \| 419 \| 0.35 \| 157 \| 86 \| 5 \| 3 \| 146 \| 6 \| 159 \| Lysozyme \| Lysozyme; afdb-uniprot50 \| AF-A0A3B9L692-F1-MODEL\_V4 \| 1.0 \| 1.282e-09 \| 419 \| 0.456 \| 151 \| 71 \| 5 \| 1 \| 143 \| 9 \| 156 \| Lysozyme \| Lysozyme; afdb-uniprot50 \| AF-A0A376MWV7-F1-MODEL\_V4 \| 1.0 \| 3.157e-09 \| 419 \| 0.319 \| 144 \| 90 \| 3 \| 6 \| 144 \| 21 \| 161 \| Lysozyme \| Lysozyme; afdb-uniprot50 \| AF-A0A350KLD8-F1-MODEL\_V4 \| 1.0 \| 2.422e-09 \| 419 \| 0.364 \| 151 \| 92 \| 3 \| 1 \| 150 \| 29 \| 176 \| Lysozyme \| Lysozyme; afdb-uniprot50 \| AF-E0WUE6-F1-MODEL\_V4 \| 1.0 \| 3.902e-09 \| 419 \| 0.337 \| 145 \| 90 \| 3 \| 4 \| 145 \| 61 \| 202 \| Lysozyme \| Lysozyme; afdb-uniprot50 \| AF-A0A7U7FZ79-F1-MODEL\_V4 \| 1.0 \| 6.628e-09 \| 418 \| 0.407 \| 130 \| 74 \| 2 \| 14 \| 143 \| 31 \| 157 \| Lysozyme \| Lysozyme; afdb-uniprot50 \| AF-V7ZPU5-F1-MODEL\_V4 \| 1.0 \| 5.362e-09 \| 418 \| 0.331 \| 148 \| 92 \| 4 \| 1 \| 144 \| 1 \| 145 \| Lysozyme \| Lysozyme; afdb-uniprot50 \| AF-A0A7R9M799-F1-MODEL\_V4 \| 1.0 \| 5.362e-09 \| 418 \| 0.405 \| 148 \| 79 \| 5 \| 1 \| 144 \| 100 \| 242 \| Lysozyme \| Lysozyme; afdb-uniprot50 \| AF-A0A328I7V7-F1-MODEL\_V4 \| 1.0 \| 1.762e-09 \| 418 \| 0.369 \| 149 \| 84 \| 4 \| 3 \| 147 \| 131 \| 273 \| Lysozyme \| Lysozyme; afdb-uniprot50 \| AF-A0A7X2HR52-F1-MODEL\_V4 \| 1.0 \| 4.114e-09 \| 417 \| 0.388 \| 144 \| 84 \| 2 \| 1 \| 144 \| 39 \| 178 \| Lysozyme \| Lysozyme; afdb-uniprot50 \| AF-E6EY07-F1-MODEL\_V4 \| 1.0 \| 5.085e-09 \| 417 \| 0.31 \| 148 \| 95 \| 4 \| 1 \| 144 \| 1 \| 145 \| Lysozyme \| Lysozyme; afdb-uniprot50 \| AF-E8RMG8-F1-MODEL\_V4 \| 1.0 \| 5.085e-09 \| 417 \| 0.393 \| 150 \| 83 \| 5 \| 1 \| 146 \| 5 \| 150 \| Lysozyme \| Lysozyme; afdb-uniprot50 \| AF-A0A4Z0P380-F1-MODEL\_V4 \| 1.0 \| 3.7e-09 \| 415 \| 0.425 \| 155 \| 76 \| 6 \| 1 \| 146 \| 3 \| 153 \| Lysozyme \| Lysozyme; afdb-uniprot50 \| AF-A0A257UY12-F1-MODEL\_V4 \| 1.0 \| 2.759e-10 \| 415 \| 0.394 \| 157 \| 83 \| 5 \| 6 \| 154 \| 23 \| 175 \| Lysozyme \| Lysozyme; afdb-uniprot50 \| AF-A0A2G2G4E8-F1-MODEL\_V4 \| 1.0 \| 3.157e-09 \| 415 \| 0.38 \| 150 \| 85 \| 5 \| 1 \| 146 \| 5 \| 150 \| Lysozyme \| Lysozyme; afdb-uniprot50 \| AF-A0A3B0R8Q5-F1-MODEL\_V4 \| 1.0 \| 6.628e-09 \| 415 \| 0.337 \| 148 \| 90 \| 5 \| 3 \| 146 \| 7 \| 150 \| Lysozyme \| Lysozyme; afdb-uniprot50 \| AF-A0A2C5THG2-F1-MODEL\_V4 \| 1.0 \| 1.126e-08 \| 414 \| 0.34 \| 135 \| 86 \| 2 \| 11 \| 145 \| 2 \| 133 \| Lysozyme \| Lysozyme; afdb-uniprot50 \| AF-A0A1W1XTA9-F1-MODEL\_V4 \| 1.0 \| 2.178e-09 \| 414 \| 0.32 \| 150 \| 94 \| 4 \| 5 \| 150 \| 14 \| 159 \| Lysozyme \| Lysozyme; afdb-uniprot50 \| AF-A0A7C5SDY7-F1-MODEL\_V4 \| 1.0 \| 1.068e-08 \| 414 \| 0.379 \| 153 \| 85 \| 5 \| 1 \| 146 \| 37 \| 186 \| Lysozyme \| Lysozyme; afdb-uniprot50 \| AF-L8M4C1-F1-MODEL\_V4 \| 1.0 \| 2.839e-09 \| 414 \| 0.373 \| 150 \| 84 \| 5 \| 2 \| 144 \| 107 \| 253 \| Lysozyme \| Lysozyme; afdb-uniprot50 \| AF-A0A843K231-F1-MODEL\_V4 \| 1.0 \| 1.216e-09 \| 414 \| 0.361 \| 155 \| 86 \| 5 \| 1 \| 145 \| 27 \| 178 \| Lysozyme \| Lysozyme; afdb-uniprot50 \| AF-A0A3T0EDP8-F1-MODEL\_V4 \| 1.0 \| 1.585e-09 \| 414 \| 0.378 \| 148 \| 82 \| 6 \| 5 \| 146 \| 332 \| 475 \| Lysozyme \| Lysozyme; afdb-uniprot50 \| AF-A0A158G2K7-F1-MODEL\_V4 \| 1.0 \| 5.654e-09 \| 413 \| 0.557 \| 104 \| 46 \| 0 \| 44 \| 147 \| 1 \| 104 \| Lysozyme \| Lysozyme; afdb-uniprot50 \| AF-A0A1F4JWI6-F1-MODEL\_V4 \| 1.0 \| 5.085e-09 \| 413 \| 0.307 \| 143 \| 93 \| 3 \| 3 \| 145 \| 16 \| 152 \| Lysozyme \| Lysozyme; afdb-uniprot50 \| AF-D7N2W0-F1-MODEL\_V4 \| 1.0 \| 6.286e-09 \| 413 \| 0.29 \| 155 \| 99 \| 2 \| 1 \| 147 \| 5 \| 156 \| Lysozyme \| Lysozyme; afdb-uniprot50 \| AF-A0A401IDR8-F1-MODEL\_V4 \| 1.0 \| 1.426e-09 \| 413 \| 0.398 \| 148 \| 78 \| 5 \| 1 \| 140 \| 1 \| 145 \| Lysozyme \| Lysozyme; afdb-uniprot50 \| AF-A0A7I6QAH2-F1-MODEL\_V4 \| 1.0 \| 2.554e-09 \| 413 \| 0.358 \| 145 \| 89 \| 3 \| 7 \| 151 \| 26 \| 166 \| Lysozyme \| Lysozyme; afdb-uniprot50 \| AF-A0A2W4UPU2-F1-MODEL\_V4 \| 1.0 \| 3.509e-09 \| 413 \| 0.383 \| 154 \| 80 \| 3 \| 1 \| 145 \| 1 \| 148 \| Lysozyme \| Lysozyme; afdb-uniprot50 \| AF-K9XTT4-F1-MODEL\_V4 \| 1.0 \| 2.066e-09 \| 413 \| 0.385 \| 153 \| 84 \| 5 \| 1 \| 146 \| 71 \| 220 \| Lysozyme \| Lysozyme; afdb-uniprot50 \| AF-A0A6L3STT2-F1-MODEL\_V4 \| 1.0 \| 2.994e-09 \| 413 \| 0.373 \| 150 \| 82 \| 5 \| 1 \| 146 \| 1 \| 142 \| Lysozyme \| Lysozyme; afdb-uniprot50 \| AF-A0A843LAG6-F1-MODEL\_V4 \| 1.0 \| 9.839e-10 \| 413 \| 0.385 \| 153 \| 81 \| 5 \| 1 \| 143 \| 27 \| 176 \| Glycoside hydrolase family protein \| Glycoside hydrolase family protein; afdb-uniprot50 \| AF-A0A0R2JFT7-F1-MODEL\_V4 \| 1.0 \| 4.338e-09 \| 412 \| 0.364 \| 148 \| 88 \| 5 \| 1 \| 146 \| 1 \| 144 \| Lysozyme \| Lysozyme; afdb-uniprot50 \| AF-A0A3N0WCV2-F1-MODEL\_V4 \| 1.0 \| 8.192e-09 \| 412 \| 0.366 \| 150 \| 85 \| 5 \| 1 \| 144 \| 25 \| 170 \| Lysozyme \| Lysozyme; afdb-uniprot50 \| AF-A0A822UE20-F1-MODEL\_V4 \| 1.0 \| 2.693e-09 \| 411 \| 0.313 \| 150 \| 90 \| 3 \| 5 \| 144 \| 17 \| 163 \| Phage lysozome \| Phage lysozome; afdb-uniprot50 \| AF-A0A366C290-F1-MODEL\_V4 \| 1.0 \| 7.77e-09 \| 411 \| 0.328 \| 140 \| 90 \| 3 \| 5 \| 144 \| 30 \| 165 \| Lysozyme \| Lysozyme; afdb-uniprot50 \| AF-A5FWW2-F1-MODEL\_V4 \| 1.0 \| 1.094e-09 \| 411 \| 0.367 \| 155 \| 86 \| 4 \| 1 \| 146 \| 21 \| 172 \| Lysozyme \| Lysozyme; afdb-uniprot50 \| AF-F9RLU7-F1-MODEL\_V4 \| 1.0 \| 3.902e-09 \| 411 \| 0.355 \| 166 \| 92 \| 5 \| 1 \| 157 \| 29 \| 188 \| Lysozyme \| Lysozyme; afdb-uniprot50 \| AF-A0A257R9J7-F1-MODEL\_V4 \| 1.0 \| 6.79e-10 \| 411 \| 0.384 \| 156 \| 80 \| 5 \| 4 \| 146 \| 37 \| 189 \| Lysozyme \| Lysozyme; afdb-uniprot50 \| AF-A0A258BAK0-F1-MODEL\_V4 \| 1.0 \| 3.902e-09 \| 411 \| 0.374 \| 155 \| 89 \| 5 \| 1 \| 151 \| 5 \| 155 \| Lysozyme \| Lysozyme; afdb-uniprot50 \| AF-A0A7C5QZP7-F1-MODEL\_V4 \| 1.0 \| 3.902e-09 \| 411 \| 0.355 \| 152 \| 90 \| 5 \| 1 \| 148 \| 5 \| 152 \| Lysozyme \| Lysozyme; afdb-uniprot50 \| AF-A0A514C2T5-F1-MODEL\_V4 \| 1.0 \| 8.192e-09 \| 411 \| 0.331 \| 160 \| 94 \| 6 \| 1 \| 154 \| 27 \| 179 \| Lysozyme \| Lysozyme; afdb-uniprot50 \| AF-A0A6P6XNS9-F1-MODEL\_V4 \| 1.0 \| 3.7e-09 \| 410 \| 0.396 \| 154 \| 81 \| 5 \| 1 \| 148 \| 33 \| 180 \| Lysozyme \| Lysozyme; afdb-uniprot50 \| AF-A0A7T2CTG2-F1-MODEL\_V4 \| 1.0 \| 3.7e-09 \| 410 \| 0.333 \| 150 \| 93 \| 3 \| 1 \| 146 \| 1 \| 147 \| Lysozyme \| Lysozyme; afdb-uniprot50 \| AF-A0A6M3LU93-F1-MODEL\_V4 \| 1.0 \| 3.902e-09 \| 409 \| 0.31 \| 145 \| 93 \| 4 \| 1 \| 143 \| 1 \| 140 \| Lysozyme \| Lysozyme; afdb-uniprot50 \| AF-A0A0R0CI69-F1-MODEL\_V4 \| 1.0 \| 3.328e-09 \| 409 \| 0.286 \| 150 \| 98 \| 4 \| 1 \| 144 \| 13 \| 159 \| Lysozyme \| Lysozyme; afdb-uniprot50 \| AF-A0A558IEL7-F1-MODEL\_V4 \| 1.0 \| 9.108e-09 \| 409 \| 0.338 \| 139 \| 87 \| 4 \| 6 \| 144 \| 18 \| 151 \| Lysozyme \| Lysozyme; afdb-uniprot50 \| AF-A0A6S7CW88-F1-MODEL\_V4 \| 1.0 \| 1.037e-09 \| 409 \| 0.374 \| 147 \| 82 \| 6 \| 6 \| 146 \| 22 \| 164 \| Lysozyme \| Lysozyme; afdb-uniprot50 \| AF-A0A1Q3Q7Z8-F1-MODEL\_V4 \| 1.0 \| 8.393e-10 \| 409 \| 0.361 \| 166 \| 79 \| 4 \| 4 \| 145 \| 1 \| 163 \| Lysozyme \| Lysozyme; afdb-uniprot50 \| AF-A0A5B0FQN8-F1-MODEL\_V4 \| 1.0 \| 2.066e-09 \| 409 \| 0.337 \| 154 \| 89 \| 5 \| 1 \| 146 \| 4 \| 152 \| Lysozyme \| Lysozyme; afdb-uniprot50 \| AF-A0A2G2GMC1-F1-MODEL\_V4 \| 1.0 \| 1.392e-08 \| 409 \| 0.335 \| 149 \| 91 \| 4 \| 2 \| 146 \| 6 \| 150 \| Lysozyme \| Lysozyme; afdb-uniprot50 \| AF-A0A6G9YUS1-F1-MODEL\_V4 \| 1.0 \| 2.994e-09 \| 409 \| 0.436 \| 149 \| 74 \| 4 \| 2 \| 143 \| 593 \| 738 \| Lysozyme \| Lysozyme; afdb-uniprot50 \| AF-A0A350KNR5-F1-MODEL\_V4 \| 1.0 \| 1.187e-08 \| 408 \| 0.349 \| 146 \| 90 \| 4 \| 3 \| 146 \| 2 \| 144 \| Lysozyme \| Lysozyme; afdb-uniprot50 \| AF-A0A397VV26-F1-MODEL\_V4 \| 1.0 \| 6.628e-09 \| 408 \| 0.337 \| 154 \| 90 \| 5 \| 2 \| 148 \| 116 \| 264 \| Lysozyme \| Lysozyme; afdb-uniprot50 \| AF-A0A2D7Y4X6-F1-MODEL\_V4 \| 1.0 \| 2.554e-09 \| 408 \| 0.377 \| 151 \| 82 \| 5 \| 3 \| 144 \| 632 \| 779 \| Lysozyme \| Lysozyme; afdb-uniprot50 \| AF-A0A3N5EFT6-F1-MODEL\_V4 \| 1.0 \| 1.762e-09 \| 407 \| 0.343 \| 157 \| 90 \| 6 \| 2 \| 145 \| 7 \| 163 \| Lysozyme \| Lysozyme; afdb-uniprot50 \| AF-A0A345P973-F1-MODEL\_V4 \| 1.0 \| 7.369e-09 \| 407 \| 0.3 \| 143 \| 97 \| 2 \| 4 \| 146 \| 30 \| 169 \| Lysozyme \| Lysozyme; afdb-uniprot50 \| AF-A0A2G2J894-F1-MODEL\_V4 \| 1.0 \| 7.77e-09 \| 407 \| 0.326 \| 150 \| 93 \| 5 \| 1 \| 146 \| 5 \| 150 \| Lysozyme \| Lysozyme; afdb-uniprot50 \| AF-A0A850B733-F1-MODEL\_V4 \| 1.0 \| 2.994e-09 \| 407 \| 0.427 \| 152 \| 75 \| 6 \| 1 \| 143 \| 748 \| 896 \| Type VI secretion system tip protein VgrG \| Type VI secretion system tip protein VgrG; afdb-uniprot50 \| AF-A0A060C089-F1-MODEL\_V4 \| 1.0 \| 7.77e-09 \| 406 \| 0.396 \| 131 \| 76 \| 2 \| 15 \| 145 \| 1 \| 128 \| Lysozyme \| Lysozyme; afdb-uniprot50 \| AF-A0A839EPA8-F1-MODEL\_V4 \| 1.0 \| 4.114e-09 \| 406 \| 0.325 \| 163 \| 92 \| 6 \| 1 \| 148 \| 3 \| 162 \| Lysozyme \| Lysozyme; afdb-uniprot50 \| AF-A0A0Q6RTE5-F1-MODEL\_V4 \| 1.0 \| 2.554e-09 \| 406 \| 0.343 \| 166 \| 90 \| 5 \| 1 \| 147 \| 3 \| 168 \| Lysozyme \| Lysozyme; afdb-uniprot50 \| AF-A0A443L6C9-F1-MODEL\_V4 \| 1.0 \| 1.631e-08 \| 405 \| 0.357 \| 140 \| 86 \| 3 \| 5 \| 144 \| 9 \| 144 \| Lysozyme \| Lysozyme; afdb-uniprot50 \| AF-A0A1Y0YBN9-F1-MODEL\_V4 \| 1.0 \| 3.157e-09 \| 405 \| 0.367 \| 147 \| 84 \| 4 \| 6 \| 146 \| 7 \| 150 \| Lysozyme \| Lysozyme; afdb-uniprot50 \| AF-A0A5R9KB34-F1-MODEL\_V4 \| 1.0 \| 1.216e-09 \| 405 \| 0.391 \| 156 \| 81 \| 6 \| 1 \| 145 \| 3 \| 155 \| Lysozyme \| Lysozyme; afdb-uniprot50 \| AF-A0A330M3W1-F1-MODEL\_V4 \| 1.0 \| 5.362e-09 \| 405 \| 0.326 \| 147 \| 91 \| 4 \| 3 \| 144 \| 12 \| 155 \| Lysozyme \| Lysozyme; afdb-uniprot50 \| AF-A0A838L2V2-F1-MODEL\_V4 \| 1.0 \| 3.7e-09 \| 405 \| 0.306 \| 160 \| 92 \| 4 \| 2 \| 146 \| 184 \| 339 \| Lysozyme \| Lysozyme; afdb-uniprot50 \| AF-A0A1X7KQT0-F1-MODEL\_V4 \| 1.0 \| 1.094e-09 \| 404 \| 0.481 \| 137 \| 56 \| 4 \| 20 \| 144 \| 2 \| 135 \| Lysozyme \| Lysozyme; afdb-uniprot50 \| AF-A0A839A1U5-F1-MODEL\_V4 \| 1.0 \| 9.108e-09 \| 404 \| 0.345 \| 142 \| 86 \| 4 \| 5 \| 143 \| 15 \| 152 \| Lysozyme \| Lysozyme; afdb-uniprot50 \| AF-A0A6N6MEH8-F1-MODEL\_V4 \| 1.0 \| 4.823e-09 \| 404 \| 0.38 \| 150 \| 81 \| 5 \| 1 \| 146 \| 1 \| 142 \| Lysozyme \| Lysozyme; afdb-uniprot50 \| AF-A0A2D9KP11-F1-MODEL\_V4 \| 1.0 \| 5.362e-09 \| 403 \| 0.326 \| 144 \| 91 \| 4 \| 4 \| 144 \| 16 \| 156 \| Lysozyme \| Lysozyme; afdb-uniprot50 \| AF-A0A5C8S7S7-F1-MODEL\_V4 \| 1.0 \| 3.7e-09 \| 403 \| 0.401 \| 147 \| 76 \| 6 \| 1 \| 143 \| 1 \| 139 \| Lysozyme \| Lysozyme; afdb-uniprot50 \| AF-A0A644VLX9-F1-MODEL\_V4 \| 1.0 \| 1.72e-08 \| 403 \| 0.35 \| 140 \| 87 \| 3 \| 5 \| 144 \| 133 \| 268 \| Lysozyme \| Lysozyme; afdb-uniprot50 \| AF-A0A4Y9RDX4-F1-MODEL\_V4 \| 1.0 \| 5.085e-09 \| 403 \| 0.333 \| 159 \| 95 \| 5 \| 1 \| 154 \| 16 \| 168 \| Lysozyme \| Lysozyme; afdb-uniprot50 \| AF-A0A522A8D2-F1-MODEL\_V4 \| 1.0 \| 5.085e-09 \| 403 \| 0.372 \| 145 \| 88 \| 2 \| 1 \| 145 \| 179 \| 320 \| Lysozyme \| Lysozyme; afdb-uniprot50 \| AF-A0A1S8P180-F1-MODEL\_V4 \| 1.0 \| 3.7e-09 \| 402 \| 0.361 \| 144 \| 88 \| 3 \| 2 \| 143 \| 131 \| 272 \| Lysozyme \| Lysozyme; afdb-uniprot50 \| AF-A0A059E111-F1-MODEL\_V4 \| 1.0 \| 1.126e-08 \| 402 \| 0.344 \| 151 \| 90 \| 6 \| 1 \| 146 \| 10 \| 156 \| Lysozyme \| Lysozyme; afdb-uniprot50 \| AF-C6XLA1-F1-MODEL\_V4 \| 1.0 \| 1.068e-08 \| 402 \| 0.344 \| 148 \| 89 \| 5 \| 1 \| 144 \| 5 \| 148 \| Lysozyme \| Lysozyme; afdb-uniprot50 \| AF-A0A2E5TFG5-F1-MODEL\_V4 \| 1.0 \| 2.126e-08 \| 401 \| 0.354 \| 141 \| 86 \| 4 \| 5 \| 144 \| 15 \| 151 \| Lysozyme \| Lysozyme; afdb-uniprot50 \| AF-U2N1X6-F1-MODEL\_V4 \| 1.0 \| 2.017e-08 \| 401 \| 0.384 \| 143 \| 84 \| 3 \| 1 \| 143 \| 26 \| 164 \| Lysozyme \| Lysozyme; afdb-uniprot50 \| AF-A0A1V2GWA1-F1-MODEL\_V4 \| 1.0 \| 5.085e-09 \| 401 \| 0.318 \| 160 \| 99 \| 5 \| 2 \| 155 \| 22 \| 177 \| Uncharacterized protein \| Uncharacterized protein; afdb-uniprot50 \| AF-A0A4U0YH08-F1-MODEL\_V4 \| 1.0 \| 1.547e-08 \| 401 \| 0.384 \| 138 \| 80 \| 4 \| 7 \| 144 \| 2 \| 134 \| Lysozyme \| Lysozyme; afdb-uniprot50 \| AF-A0A7Y0NU04-F1-MODEL\_V4 \| 1.0 \| 3.328e-09 \| 401 \| 0.361 \| 152 \| 87 \| 5 \| 2 \| 146 \| 88 \| 236 \| Lysozyme \| Lysozyme; afdb-uniprot50 \| AF-A0A2E6Y4S6-F1-MODEL\_V4 \| 1.0 \| 6.988e-09 \| 400 \| 0.349 \| 146 \| 86 \| 5 \| 4 \| 144 \| 12 \| 153 \| Lysozyme \| Lysozyme; afdb-uniprot50 \| AF-A0A8A3KUB0-F1-MODEL\_V4 \| 1.0 \| 3.902e-09 \| 400 \| 0.361 \| 163 \| 85 \| 5 \| 1 \| 144 \| 3 \| 165 \| Lysozyme \| Lysozyme; afdb-uniprot50 \| AF-A0A377A078-F1-MODEL\_V4 \| 1.0 \| 5.654e-09 \| 399 \| 0.335 \| 146 \| 92 \| 4 \| 7 \| 152 \| 8 \| 148 \| Lysozyme \| Lysozyme; afdb-uniprot50 \| AF-A0A827LRS5-F1-MODEL\_V4 \| 1.0 \| 1.912e-08 \| 399 \| 0.361 \| 144 \| 88 \| 3 \| 1 \| 144 \| 27 \| 166 \| Lysozyme \| Lysozyme; afdb-uniprot50 \| AF-A0A3P1W4Z0-F1-MODEL\_V4 \| 1.0 \| 2.994e-09 \| 399 \| 0.318 \| 160 \| 99 \| 2 \| 2 \| 153 \| 14 \| 171 \| Lysozyme \| Lysozyme; afdb-uniprot50 \| AF-A0A069IM11-F1-MODEL\_V4 \| 1.0 \| 5.961e-09 \| 399 \| 0.348 \| 158 \| 91 \| 6 \| 2 \| 154 \| 9 \| 159 \| Lysozyme \| Lysozyme; afdb-uniprot50 \| AF-A0A6P0MRG7-F1-MODEL\_V4 \| 1.0 \| 6.628e-09 \| 399 \| 0.32 \| 156 \| 92 \| 2 \| 2 \| 146 \| 6 \| 158 \| Lysozyme \| Lysozyme; afdb-uniprot50 \| AF-A0A7X0AWL3-F1-MODEL\_V4 \| 1.0 \| 2.554e-09 \| 398 \| 0.406 \| 150 \| 76 \| 7 \| 1 \| 143 \| 3 \| 146 \| Lysozyme \| Lysozyme; afdb-uniprot50 \| AF-A0A2T1CPP5-F1-MODEL\_V4 \| 1.0 \| 8.638e-09 \| 398 \| 0.389 \| 149 \| 81 \| 5 \| 3 \| 144 \| 63 \| 208 \| Lysozyme \| Lysozyme; afdb-uniprot50 \| AF-A0A2G1YWA0-F1-MODEL\_V4 \| 1.0 \| 9.604e-09 \| 398 \| 0.353 \| 147 \| 87 \| 5 \| 2 \| 144 \| 6 \| 148 \| Lysozyme \| Lysozyme; afdb-uniprot50 \| AF-A0A6N6L8L1-F1-MODEL\_V4 \| 1.0 \| 1.547e-08 \| 397 \| 0.356 \| 146 \| 84 \| 5 \| 3 \| 143 \| 2 \| 142 \| Lysozyme \| Lysozyme; afdb-uniprot50 \| AF-A0A764SMJ2-F1-MODEL\_V4 \| 1.0 \| 9.108e-09 \| 397 \| 0.316 \| 142 \| 89 \| 3 \| 6 \| 142 \| 21 \| 159 \| Lysozyme \| Lysozyme; afdb-uniprot50 \| AF-A0A389MSU7-F1-MODEL\_V4 \| 1.0 \| 5.362e-09 \| 397 \| 0.408 \| 147 \| 75 \| 5 \| 1 \| 143 \| 1 \| 139 \| Lysozyme \| Lysozyme; afdb-uniprot50 \| AF-C3L4H1-F1-MODEL\_V4 \| 1.0 \| 2.066e-09 \| 397 \| 0.397 \| 151 \| 81 \| 6 \| 2 \| 145 \| 431 \| 578 \| Lysozyme \| Lysozyme; afdb-uniprot50 \| AF-A0A2S5TTS2-F1-MODEL\_V4 \| 1.0 \| 1.912e-08 \| 396 \| 0.404 \| 146 \| 75 \| 5 \| 3 \| 144 \| 1 \| 138 \| Lysozyme \| Lysozyme; afdb-uniprot50 \| AF-A0A6L6G496-F1-MODEL\_V4 \| 1.0 \| 4.574e-09 \| 396 \| 0.307 \| 156 \| 93 \| 3 \| 2 \| 145 \| 3 \| 155 \| Lysozyme \| Lysozyme; afdb-uniprot50 \| AF-A0A5C8W707-F1-MODEL\_V4 \| 1.0 \| 6.988e-09 \| 396 \| 0.36 \| 150 \| 84 \| 5 \| 1 \| 146 \| 1 \| 142 \| Lysozyme \| Lysozyme; afdb-uniprot50 \| AF-A0A2V3TSZ6-F1-MODEL\_V4 \| 1.0 \| 5.085e-09 \| 396 \| 0.36 \| 147 \| 81 \| 5 \| 1 \| 143 \| 1 \| 138 \| Lysozyme \| Lysozyme; afdb-uniprot50 \| AF-A0A563VMR6-F1-MODEL\_V4 \| 1.0 \| 6.628e-09 \| 396 \| 0.337 \| 151 \| 90 \| 5 \| 3 \| 146 \| 51 \| 198 \| Lysozyme \| Lysozyme; afdb-uniprot50 \| AF-A0A519IYM2-F1-MODEL\_V4 \| 1.0 \| 1.72e-08 \| 396 \| 0.32 \| 159 \| 97 \| 5 \| 1 \| 154 \| 1 \| 153 \| Lysozyme \| Lysozyme; afdb-uniprot50 \| AF-A0A8A7UZQ7-F1-MODEL\_V4 \| 1.0 \| 8.192e-09 \| 396 \| 0.316 \| 161 \| 95 \| 6 \| 1 \| 154 \| 1 \| 153 \| Glycoside hydrolase family protein \| Glycoside hydrolase family protein; afdb-uniprot50 \| AF-A0A711C0N5-F1-MODEL\_V4 \| 1.0 \| 1.631e-08 \| 395 \| 0.542 \| 118 \| 50 \| 2 \| 31 \| 144 \| 2 \| 119 \| Lysozyme \| Lysozyme; afdb-uniprot50 \| AF-A0A4R8F7V4-F1-MODEL\_V4 \| 1.0 \| 2.017e-08 \| 395 \| 0.304 \| 141 \| 94 \| 3 \| 4 \| 144 \| 1 \| 137 \| Lysozyme \| Lysozyme; afdb-uniprot50 \| AF-A0A7V3LJU9-F1-MODEL\_V4 \| 1.0 \| 1.392e-08 \| 395 \| 0.319 \| 147 \| 93 \| 5 \| 1 \| 144 \| 1 \| 143 \| Lysozyme \| Lysozyme; afdb-uniprot50 \| AF-A0A233HE82-F1-MODEL\_V4 \| 1.0 \| 4.823e-09 \| 395 \| 0.297 \| 158 \| 93 \| 4 \| 1 \| 143 \| 1 \| 155 \| Lysozyme \| Lysozyme; afdb-uniprot50 \| AF-A0A1X0X4Q9-F1-MODEL\_V4 \| 1.0 \| 1.631e-08 \| 395 \| 0.33 \| 142 \| 90 \| 4 \| 4 \| 144 \| 32 \| 169 \| Lysozyme \| Lysozyme; afdb-uniprot50 \| AF-A0A345CWD3-F1-MODEL\_V4 \| 1.0 \| 6.988e-09 \| 395 \| 0.363 \| 143 \| 86 \| 4 \| 10 \| 152 \| 62 \| 199 \| Lysozyme \| Lysozyme; afdb-uniprot50 \| AF-A0A1H8ACR7-F1-MODEL\_V4 \| 1.0 \| 3.7e-09 \| 395 \| 0.37 \| 151 \| 82 \| 7 \| 1 \| 146 \| 71 \| 213 \| Lysozyme \| Lysozyme; afdb-uniprot50 \| AF-A0A258FEZ4-F1-MODEL\_V4 \| 1.0 \| 1.068e-08 \| 395 \| 0.339 \| 159 \| 94 \| 5 \| 1 \| 154 \| 9 \| 161 \| Lysozyme \| Lysozyme; afdb-uniprot50 \| AF-A0A6B8KGM4-F1-MODEL\_V4 \| 1.0 \| 1.216e-09 \| 394 \| 0.403 \| 156 \| 75 \| 8 \| 1 \| 146 \| 2 \| 149 \| Lysozyme \| Lysozyme; afdb-uniprot50 \| AF-A0A2U1SSP9-F1-MODEL\_V4 \| 1.0 \| 1.912e-08 \| 394 \| 0.391 \| 148 \| 78 \| 4 \| 1 \| 144 \| 3 \| 142 \| Lysozyme \| Lysozyme; afdb-uniprot50 \| AF-A0A2K9CTZ1-F1-MODEL\_V4 \| 1.0 \| 2.364e-08 \| 393 \| 0.581 \| 98 \| 41 \| 0 \| 47 \| 144 \| 1 \| 98 \| Lysozyme \| Lysozyme; afdb-uniprot50 \| AF-A0A2P7SF92-F1-MODEL\_V4 \| 1.0 \| 1.72e-08 \| 393 \| 0.286 \| 143 \| 96 \| 4 \| 4 \| 144 \| 1 \| 139 \| Lysozyme \| Lysozyme; afdb-uniprot50 \| AF-A0A7L7IWX3-F1-MODEL\_V4 \| 1.0 \| 2.839e-09 \| 393 \| 0.363 \| 157 \| 83 \| 4 \| 1 \| 143 \| 1 \| 154 \| Lysozyme \| Lysozyme; afdb-uniprot50 \| AF-A0A1G1BDY3-F1-MODEL\_V4 \| 1.0 \| 4.823e-09 \| 393 \| 0.314 \| 159 \| 100 \| 5 \| 1 \| 155 \| 5 \| 158 \| Lysozyme \| Lysozyme; afdb-uniprot50 \| AF-A0A3D0HRH7-F1-MODEL\_V4 \| 1.0 \| 1.126e-08 \| 393 \| 0.419 \| 155 \| 75 \| 7 \| 2 \| 145 \| 7 \| 157 \| Lysozyme \| Lysozyme; afdb-uniprot50 \| AF-H1KV31-F1-MODEL\_V4 \| 1.0 \| 5.085e-09 \| 393 \| 0.387 \| 147 \| 78 \| 6 \| 1 \| 143 \| 1 \| 139 \| Lysozyme \| Lysozyme; afdb-uniprot50 \| AF-A0A252DI96-F1-MODEL\_V4 \| 1.0 \| 3.249e-08 \| 392 \| 0.447 \| 105 \| 58 \| 0 \| 47 \| 151 \| 1 \| 105 \| Lysozyme \| Lysozyme; afdb-uniprot50 \| AF-A0A2W2BWY0-F1-MODEL\_V4 \| 1.0 \| 3.509e-09 \| 392 \| 0.343 \| 157 \| 91 \| 4 \| 1 \| 146 \| 2 \| 157 \| Lysozyme \| Lysozyme; afdb-uniprot50 \| AF-A0A6L3SVH8-F1-MODEL\_V4 \| 1.0 \| 8.192e-09 \| 392 \| 0.373 \| 150 \| 82 \| 6 \| 1 \| 146 \| 1 \| 142 \| Lysozyme \| Lysozyme; afdb-uniprot50 \| AF-A0A455RUP7-F1-MODEL\_V4 \| 1.0 \| 6.821e-08 \| 391 \| 0.551 \| 98 \| 44 \| 0 \| 47 \| 144 \| 1 \| 98 \| Lysozyme \| Lysozyme; afdb-uniprot50 \| AF-A0A291E4Z2-F1-MODEL\_V4 \| 1.0 \| 2.771e-08 \| 391 \| 0.321 \| 146 \| 92 \| 4 \| 5 \| 146 \| 4 \| 146 \| Lysozyme \| Lysozyme; afdb-uniprot50 \| AF-A0A0L0AVI2-F1-MODEL\_V4 \| 1.0 \| 8.638e-09 \| 391 \| 0.317 \| 145 \| 95 \| 3 \| 1 \| 144 \| 6 \| 147 \| Lysozyme \| Lysozyme; afdb-uniprot50 \| AF-A0A1W2A982-F1-MODEL\_V4 \| 1.0 \| 2.017e-08 \| 391 \| 0.314 \| 143 \| 95 \| 2 \| 5 \| 145 \| 40 \| 181 \| Lysozyme \| Lysozyme; afdb-uniprot50 \| AF-A0A5C8UIV5-F1-MODEL\_V4 \| 1.0 \| 1.013e-08 \| 391 \| 0.375 \| 144 \| 81 \| 5 \| 3 \| 146 \| 7 \| 141 \| Lysozyme \| Lysozyme; afdb-uniprot50 \| AF-A0A1X2GML7-F1-MODEL\_V4 \| 1.0 \| 3.902e-09 \| 391 \| 0.387 \| 147 \| 82 \| 4 \| 2 \| 143 \| 122 \| 265 \| Lysozyme \| Lysozyme; afdb-uniprot50 \| AF-B4WDB1-F1-MODEL\_V4 \| 1.0 \| 1.126e-08 \| 391 \| 0.341 \| 158 \| 92 \| 6 \| 2 \| 154 \| 9 \| 159 \| Lysozyme \| Lysozyme; afdb-uniprot50 \| AF-A0A5M8SLB4-F1-MODEL\_V4 \| 1.0 \| 9.108e-09 \| 391 \| 0.344 \| 145 \| 91 \| 3 \| 5 \| 149 \| 381 \| 521 \| Lysozyme \| Lysozyme; afdb-uniprot50 \| AF-A0A741PPE7-F1-MODEL\_V4 \| 1.0 \| 7.192e-08 \| 390 \| 0.561 \| 98 \| 43 \| 0 \| 47 \| 144 \| 1 \| 98 \| Lysozyme \| Lysozyme; afdb-uniprot50 \| AF-A0A704ZET1-F1-MODEL\_V4 \| 1.0 \| 2.242e-08 \| 390 \| 0.538 \| 117 \| 50 \| 2 \| 32 \| 144 \| 1 \| 117 \| Lysozyme \| Lysozyme; afdb-uniprot50 \| AF-A0A7X5L1T0-F1-MODEL\_V4 \| 1.0 \| 5.085e-09 \| 390 \| 0.366 \| 150 \| 80 \| 6 \| 1 \| 144 \| 1 \| 141 \| Lysozyme \| Lysozyme; afdb-uniprot50 \| AF-A0A7R9PXF8-F1-MODEL\_V4 \| 1.0 \| 1.187e-08 \| 390 \| 0.375 \| 152 \| 82 \| 7 \| 1 \| 144 \| 18 \| 164 \| Lysozyme \| Lysozyme; afdb-uniprot50 \| AF-A0A7U0N3R0-F1-MODEL\_V4 \| 1.0 \| 2.628e-08 \| 390 \| 0.319 \| 147 \| 95 \| 3 \| 1 \| 146 \| 26 \| 168 \| Lysozyme \| Lysozyme; afdb-uniprot50 \| AF-A0A2U8HQZ1-F1-MODEL\_V4 \| 1.0 \| 8.638e-09 \| 390 \| 0.391 \| 148 \| 78 \| 6 \| 1 \| 144 \| 1 \| 140 \| Lysozyme \| Lysozyme; afdb-uniprot50 \| AF-A0A4Q7ICI5-F1-MODEL\_V4 \| 1.0 \| 5.085e-09 \| 390 \| 0.358 \| 156 \| 83 \| 4 \| 2 \| 143 \| 3 \| 155 \| Lysozyme \| Lysozyme; afdb-uniprot50 \| AF-A0A4P6YRS8-F1-MODEL\_V4 \| 1.0 \| 1.72e-08 \| 390 \| 0.404 \| 146 \| 81 \| 5 \| 1 \| 145 \| 5 \| 145 \| Lysozyme \| Lysozyme; afdb-uniprot50 \| AF-A0A7R9KVP1-F1-MODEL\_V4 \| 1.0 \| 5.961e-09 \| 390 \| 0.364 \| 159 \| 83 \| 4 \| 2 \| 143 \| 140 \| 297 \| Lysozyme \| Lysozyme; afdb-uniprot50 \| AF-A0A539DDV1-F1-MODEL\_V4 \| 1.0 \| 2.126e-08 \| 390 \| 0.32 \| 150 \| 94 \| 4 \| 1 \| 146 \| 25 \| 170 \| Lysozyme \| Lysozyme; afdb-uniprot50 \| AF-A0A450YWU4-F1-MODEL\_V4 \| 1.0 \| 1.187e-08 \| 390 \| 0.348 \| 155 \| 87 \| 4 \| 2 \| 146 \| 620 \| 770 \| Lysozyme \| Lysozyme; afdb-uniprot50 \| AF-A0A7C1EUW6-F1-MODEL\_V4 \| 1.0 \| 3.808e-08 \| 389 \| 0.376 \| 146 \| 84 \| 5 \| 3 \| 144 \| 1 \| 143 \| Lysozyme \| Lysozyme; afdb-uniprot50 \| AF-A0A7X9JNE0-F1-MODEL\_V4 \| 1.0 \| 4.114e-09 \| 389 \| 0.282 \| 163 \| 101 \| 5 \| 1 \| 154 \| 1 \| 156 \| Lysozyme \| Lysozyme; afdb-uniprot50 \| AF-A0A6I1MR44-F1-MODEL\_V4 \| 1.0 \| 1.671e-09 \| 389 \| 0.395 \| 167 \| 75 \| 4 \| 1 \| 144 \| 4 \| 167 \| Lysozyme \| Lysozyme; afdb-uniprot50 \| AF-A0A2G6EIY6-F1-MODEL\_V4 \| 1.0 \| 7.369e-09 \| 389 \| 0.36 \| 158 \| 82 \| 4 \| 1 \| 143 \| 17 \| 170 \| Lysozyme \| Lysozyme; afdb-uniprot50 \| AF-A0A4R5DVM8-F1-MODEL\_V4 \| 1.0 \| 3.509e-09 \| 389 \| 0.372 \| 153 \| 83 \| 6 \| 1 \| 143 \| 4 \| 153 \| Lysozyme \| Lysozyme; afdb-uniprot50 \| AF-A0A8B5V1R1-F1-MODEL\_V4 \| 1.0 \| 2.017e-08 \| 389 \| 0.339 \| 153 \| 90 \| 3 \| 1 \| 145 \| 49 \| 198 \| Lysozyme \| Lysozyme; afdb-uniprot50 \| AF-A0A326KZ45-F1-MODEL\_V4 \| 1.0 \| 1.467e-08 \| 389 \| 0.339 \| 159 \| 92 \| 7 \| 2 \| 154 \| 18 \| 169 \| Lysozyme \| Lysozyme; afdb-uniprot50 \| AF-A0A316HJ74-F1-MODEL\_V4 \| 1.0 \| 1.013e-08 \| 388 \| 0.353 \| 150 \| 87 \| 5 \| 2 \| 144 \| 5 \| 151 \| Lysozyme \| Lysozyme; afdb-uniprot50 \| AF-A0A0A2YID5-F1-MODEL\_V4 \| 1.0 \| 5.961e-09 \| 388 \| 0.357 \| 154 \| 87 \| 7 \| 1 \| 145 \| 1 \| 151 \| Lysozyme \| Lysozyme; afdb-uniprot50 \| AF-A0A501QET1-F1-MODEL\_V4 \| 1.0 \| 2.242e-08 \| 388 \| 0.372 \| 153 \| 81 \| 3 \| 1 \| 144 \| 32 \| 178 \| Lysozyme \| Lysozyme; afdb-uniprot50 \| AF-A0A679KB21-F1-MODEL\_V4 \| 1.0 \| 6.286e-09 \| 388 \| 0.355 \| 169 \| 89 \| 6 \| 1 \| 150 \| 3 \| 170 \| Lysozyme \| Lysozyme; afdb-uniprot50 \| AF-A0A5C1Q659-F1-MODEL\_V4 \| 1.0 \| 6.286e-09 \| 388 \| 0.375 \| 157 \| 84 \| 5 \| 1 \| 146 \| 303 \| 456 \| Lysozyme \| Lysozyme; afdb-uniprot50 \| AF-A0A349NKU6-F1-MODEL\_V4 \| 1.0 \| 3.425e-08 \| 387 \| 0.437 \| 112 \| 59 \| 2 \| 1 \| 112 \| 1 \| 108 \| Lysozyme \| Lysozyme; afdb-uniprot50 \| AF-A0A1H9KD71-F1-MODEL\_V4 \| 1.0 \| 1.72e-08 \| 387 \| 0.348 \| 155 \| 90 \| 4 \| 1 \| 147 \| 56 \| 207 \| Lysozyme \| Lysozyme; afdb-uniprot50 \| AF-A0A348HI68-F1-MODEL\_V4 \| 1.0 \| 5.961e-09 \| 386 \| 0.333 \| 153 \| 88 \| 6 \| 4 \| 146 \| 10 \| 158 \| Lysozyme \| Lysozyme; afdb-uniprot50 \| AF-A0A081NGH9-F1-MODEL\_V4 \| 1.0 \| 5.085e-09 \| 386 \| 0.373 \| 158 \| 84 \| 6 \| 1 \| 143 \| 1 \| 158 \| Lysozyme \| Lysozyme; afdb-uniprot50 \| AF-A0A380TT26-F1-MODEL\_V4 \| 1.0 \| 3.081e-08 \| 386 \| 0.344 \| 148 \| 90 \| 4 \| 1 \| 144 \| 26 \| 170 \| Lysozyme \| Lysozyme; afdb-uniprot50 \| AF-A0A255YT31-F1-MODEL\_V4 \| 1.0 \| 4.338e-09 \| 386 \| 0.348 \| 178 \| 80 \| 8 \| 1 \| 146 \| 1 \| 174 \| Lysozyme \| Lysozyme; afdb-uniprot50 \| AF-A0A2W6XAR0-F1-MODEL\_V4 \| 1.0 \| 5.085e-09 \| 386 \| 0.352 \| 159 \| 96 \| 5 \| 2 \| 159 \| 10 \| 162 \| Lysozyme \| Lysozyme; afdb-uniprot50 \| AF-A0A2G2HXP0-F1-MODEL\_V4 \| 1.0 \| 4.234e-08 \| 386 \| 0.337 \| 151 \| 91 \| 6 \| 1 \| 146 \| 4 \| 150 \| Lysozyme \| Lysozyme; afdb-uniprot50 \| AF-A0A661CVB8-F1-MODEL\_V4 \| 1.0 \| 4.823e-09 \| 386 \| 0.296 \| 162 \| 90 \| 3 \| 2 \| 143 \| 243 \| 400 \| Lysozyme \| Lysozyme; afdb-uniprot50 \| AF-A0A1Z9AVU0-F1-MODEL\_V4 \| 1.0 \| 2.364e-08 \| 385 \| 0.388 \| 134 \| 76 \| 3 \| 2 \| 133 \| 6 \| 135 \| Lysozyme \| Lysozyme; afdb-uniprot50 \| AF-A0A7V3K1A6-F1-MODEL\_V4 \| 1.0 \| 1.126e-08 \| 385 \| 0.341 \| 164 \| 87 \| 3 \| 1 \| 145 \| 1 \| 162 \| Lysozyme \| Lysozyme; afdb-uniprot50 \| AF-A0A358J110-F1-MODEL\_V4 \| 1.0 \| 2.493e-08 \| 385 \| 0.314 \| 143 \| 94 \| 3 \| 1 \| 143 \| 26 \| 164 \| Lysozyme \| Lysozyme; afdb-uniprot50 \| AF-A0A0Q6SQZ2-F1-MODEL\_V4 \| 1.0 \| 2.628e-08 \| 385 \| 0.318 \| 154 \| 92 \| 4 \| 1 \| 145 \| 50 \| 199 \| Lysozyme \| Lysozyme; afdb-uniprot50 \| AF-A0A2W4T8V5-F1-MODEL\_V4 \| 1.0 \| 1.187e-08 \| 385 \| 0.329 \| 158 \| 92 \| 5 \| 1 \| 152 \| 54 \| 203 \| Lysozyme \| Lysozyme; afdb-uniprot50 \| AF-A0A2U9U7G2-F1-MODEL\_V4 \| 1.0 \| 2.126e-08 \| 384 \| 0.302 \| 149 \| 96 \| 5 \| 1 \| 145 \| 1 \| 145 \| Lysozyme \| Lysozyme; afdb-uniprot50 \| AF-A0A6L6YI06-F1-MODEL\_V4 \| 1.0 \| 4.464e-08 \| 384 \| 0.345 \| 139 \| 87 \| 3 \| 6 \| 144 \| 13 \| 147 \| Lysozyme \| Lysozyme; afdb-uniprot50 \| AF-A0A1H8IPR6-F1-MODEL\_V4 \| 1.0 \| 2.628e-08 \| 384 \| 0.356 \| 146 \| 90 \| 4 \| 1 \| 145 \| 10 \| 152 \| Lysozyme \| Lysozyme; afdb-uniprot50 \| AF-A0A7Y8MEE6-F1-MODEL\_V4 \| 1.0 \| 1.72e-08 \| 384 \| 0.344 \| 151 \| 91 \| 4 \| 1 \| 147 \| 4 \| 150 \| Lysozyme \| Lysozyme; afdb-uniprot50 \| AF-A0A420WMK7-F1-MODEL\_V4 \| 1.0 \| 3.612e-08 \| 384 \| 0.333 \| 150 \| 92 \| 5 \| 1 \| 146 \| 10 \| 155 \| Lysozyme \| Lysozyme; afdb-uniprot50 \| AF-V5TVX0-F1-MODEL\_V4 \| 1.0 \| 4.016e-08 \| 383 \| 0.337 \| 148 \| 93 \| 4 \| 1 \| 148 \| 17 \| 159 \| Lysozyme \| Lysozyme; afdb-uniprot50 \| AF-A0A8B3HN41-F1-MODEL\_V4 \| 1.0 \| 8.89e-08 \| 383 \| 0.244 \| 139 \| 105 \| 0 \| 6 \| 144 \| 24 \| 162 \| Lysozyme \| Lysozyme; afdb-uniprot50 \| AF-A0A4Q2RBR2-F1-MODEL\_V4 \| 1.0 \| 3.081e-08 \| 383 \| 0.391 \| 148 \| 78 \| 4 \| 1 \| 144 \| 3 \| 142 \| Lysozyme \| Lysozyme; afdb-uniprot50 \| AF-A0A4Q5WCJ2-F1-MODEL\_V4 \| 1.0 \| 2.493e-08 \| 383 \| 0.348 \| 135 \| 83 \| 2 \| 3 \| 137 \| 86 \| 215 \| Lysozyme \| Lysozyme; afdb-uniprot50 \| AF-A0A1M6L8G3-F1-MODEL\_V4 \| 1.0 \| 6.628e-09 \| 383 \| 0.321 \| 165 \| 100 \| 6 \| 2 \| 157 \| 74 \| 235 \| Lysozyme \| Lysozyme; afdb-uniprot50 \| AF-A0A847VFE4-F1-MODEL\_V4 \| 1.0 \| 6.988e-09 \| 383 \| 0.361 \| 180 \| 83 \| 7 \| 1 \| 152 \| 6 \| 181 \| Lysozyme \| Lysozyme; afdb-uniprot50 \| AF-A0A5P9KRI5-F1-MODEL\_V4 \| 1.0 \| 1.467e-08 \| 383 \| 0.333 \| 162 \| 93 \| 7 \| 1 \| 154 \| 1 \| 155 \| Lysozyme \| Lysozyme; afdb-uniprot50 \| AF-A0A0F9UX49-F1-MODEL\_V4 \| 1.0 \| 1.72e-08 \| 383 \| 0.366 \| 153 \| 84 \| 4 \| 1 \| 143 \| 559 \| 708 \| Lysozyme \| Lysozyme; afdb-uniprot50 \| AF-A0A6V8I6G4-F1-MODEL\_V4 \| 1.0 \| 1.187e-08 \| 382 \| 0.346 \| 150 \| 86 \| 5 \| 6 \| 146 \| 7 \| 153 \| Lysozyme \| Lysozyme; afdb-uniprot50 \| AF-A0A4R3HYS8-F1-MODEL\_V4 \| 1.0 \| 1.013e-08 \| 382 \| 0.347 \| 164 \| 86 \| 7 \| 1 \| 146 \| 5 \| 165 \| Lysozyme \| Lysozyme; afdb-uniprot50 \| AF-A0A832PMR6-F1-MODEL\_V4 \| 1.0 \| 6.988e-09 \| 382 \| 0.357 \| 151 \| 84 \| 4 \| 4 \| 144 \| 26 \| 173 \| Lysozyme \| Lysozyme; afdb-uniprot50 \| AF-A0A7W0JH81-F1-MODEL\_V4 \| 1.0 \| 1.068e-08 \| 382 \| 0.367 \| 166 \| 90 \| 4 \| 1 \| 158 \| 2 \| 160 \| Lysozyme \| Lysozyme; afdb-uniprot50 \| AF-A0A3S5DFC5-F1-MODEL\_V4 \| 1.0 \| 4.707e-08 \| 381 \| 0.353 \| 147 \| 89 \| 3 \| 4 \| 150 \| 96 \| 236 \| Lysozyme \| Lysozyme; afdb-uniprot50 \| AF-A0A6C8Z019-F1-MODEL\_V4 \| 1.0 \| 6.135e-08 \| 380 \| 0.564 \| 101 \| 44 \| 0 \| 44 \| 144 \| 1 \| 101 \| Lysozyme \| Lysozyme; afdb-uniprot50 \| AF-H8L2I5-F1-MODEL\_V4 \| 1.0 \| 3.425e-08 \| 380 \| 0.298 \| 144 \| 94 \| 3 \| 6 \| 145 \| 16 \| 156 \| Lysozyme \| Lysozyme; afdb-uniprot50 \| AF-A0A3P3DCL8-F1-MODEL\_V4 \| 1.0 \| 2.364e-08 \| 380 \| 0.388 \| 144 \| 80 \| 4 \| 5 \| 144 \| 33 \| 172 \| Lysozyme \| Lysozyme; afdb-uniprot50 \| AF-A0A4S2CSZ9-F1-MODEL\_V4 \| 1.0 \| 6.628e-09 \| 380 \| 0.346 \| 156 \| 88 \| 5 \| 1 \| 143 \| 26 \| 180 \| Lysozyme \| Lysozyme; afdb-uniprot50 \| AF-A0A147I3J2-F1-MODEL\_V4 \| 1.0 \| 2.628e-08 \| 379 \| 0.393 \| 150 \| 80 \| 6 \| 13 \| 156 \| 3 \| 147 \| Lysozyme \| Lysozyme; afdb-uniprot50 \| AF-A0A238JZZ0-F1-MODEL\_V4 \| 1.0 \| 3.081e-08 \| 379 \| 0.277 \| 144 \| 96 \| 4 \| 6 \| 145 \| 15 \| 154 \| Lysozyme \| Lysozyme; afdb-uniprot50 \| AF-A0A715LSR0-F1-MODEL\_V4 \| 1.0 \| 2.922e-08 \| 378 \| 0.389 \| 131 \| 76 \| 3 \| 14 \| 144 \| 2 \| 128 \| Lysozyme \| Lysozyme; afdb-uniprot50 \| AF-A0A0Q5LBE4-F1-MODEL\_V4 \| 1.0 \| 1.013e-08 \| 378 \| 0.352 \| 153 \| 85 \| 5 \| 1 \| 145 \| 4 \| 150 \| Lysozyme \| Lysozyme; afdb-uniprot50 \| AF-A0A1Q3HI04-F1-MODEL\_V4 \| 1.0 \| 5.961e-09 \| 378 \| 0.393 \| 155 \| 75 \| 5 \| 1 \| 143 \| 9 \| 156 \| Lysozyme \| Lysozyme; afdb-uniprot50 \| AF-A0A2X9ZYS3-F1-MODEL\_V4 \| 1.0 \| 1.392e-08 \| 378 \| 0.342 \| 149 \| 92 \| 4 \| 7 \| 153 \| 29 \| 173 \| Lysozyme \| Lysozyme; afdb-uniprot50 \| AF-A0A0A2X3S0-F1-MODEL\_V4 \| 1.0 \| 5.819e-08 \| 377 \| 0.242 \| 140 \| 102 \| 3 \| 6 \| 145 \| 4 \| 139 \| Lysozyme \| Lysozyme; afdb-uniprot50 \| AF-A0A2D5RX06-F1-MODEL\_V4 \| 1.0 \| 2.493e-08 \| 377 \| 0.335 \| 149 \| 93 \| 5 \| 5 \| 153 \| 17 \| 159 \| Lysozyme \| Lysozyme; afdb-uniprot50 \| AF-A0A519PZJ4-F1-MODEL\_V4 \| 1.0 \| 3.612e-08 \| 377 \| 0.337 \| 148 \| 88 \| 5 \| 1 \| 146 \| 27 \| 166 \| Lysozyme \| Lysozyme; afdb-uniprot50 \| AF-A0A1X3I4E1-F1-MODEL\_V4 \| 1.0 \| 4.707e-08 \| 377 \| 0.317 \| 145 \| 93 \| 5 \| 1 \| 144 \| 49 \| 188 \| Lysozyme \| Lysozyme; afdb-uniprot50 \| AF-A0A2T8WJ16-F1-MODEL\_V4 \| 1.0 \| 5.234e-08 \| 376 \| 0.425 \| 120 \| 62 \| 3 \| 1 \| 116 \| 1 \| 117 \| Lysozyme \| Lysozyme; afdb-uniprot50 \| AF-A0A327SGY1-F1-MODEL\_V4 \| 1.0 \| 3.612e-08 \| 376 \| 0.34 \| 150 \| 88 \| 5 \| 1 \| 144 \| 1 \| 145 \| Lysozyme \| Lysozyme; afdb-uniprot50 \| AF-A0A356FYC9-F1-MODEL\_V4 \| 1.0 \| 7.369e-09 \| 376 \| 0.36 \| 136 \| 83 \| 3 \| 11 \| 145 \| 2 \| 134 \| Lysozyme \| Lysozyme; afdb-uniprot50 \| AF-A0A1L7AKL0-F1-MODEL\_V4 \| 1.0 \| 7.77e-09 \| 376 \| 0.337 \| 160 \| 89 \| 5 \| 9 \| 155 \| 2 \| 157 \| Lysozyme \| Lysozyme; afdb-uniprot50 \| AF-A0A2S2CGK8-F1-MODEL\_V4 \| 1.0 \| 1.32e-08 \| 376 \| 0.301 \| 166 \| 99 \| 7 \| 2 \| 155 \| 5 \| 165 \| Lysozyme \| Lysozyme; afdb-uniprot50 \| AF-A0A3D5P936-F1-MODEL\_V4 \| 1.0 \| 1.631e-08 \| 376 \| 0.342 \| 146 \| 90 \| 5 \| 3 \| 145 \| 157 \| 299 \| Lysozyme \| Lysozyme; afdb-uniprot50 \| AF-J7QGR0-F1-MODEL\_V4 \| 1.0 \| 1.32e-08 \| 376 \| 0.374 \| 171 \| 83 \| 6 \| 1 \| 155 \| 4 \| 166 \| Lysozyme \| Lysozyme; afdb-uniprot50 \| AF-A0A0Q5UY70-F1-MODEL\_V4 \| 1.0 \| 1.72e-08 \| 376 \| 0.301 \| 159 \| 100 \| 5 \| 1 \| 154 \| 9 \| 161 \| Lysozyme \| Lysozyme; afdb-uniprot50 \| AF-A0A198H0T0-F1-MODEL\_V4 \| 1.0 \| 1.72e-08 \| 375 \| 0.319 \| 147 \| 96 \| 3 \| 1 \| 146 \| 6 \| 149 \| Lysozyme \| Lysozyme; afdb-uniprot50 \| AF-A0A366F692-F1-MODEL\_V4 \| 1.0 \| 2.364e-08 \| 375 \| 0.287 \| 146 \| 100 \| 2 \| 1 \| 145 \| 6 \| 148 \| Lysozyme \| Lysozyme; afdb-uniprot50 \| AF-A0A7S6S8H3-F1-MODEL\_V4 \| 1.0 \| 7.369e-09 \| 375 \| 0.333 \| 171 \| 86 \| 5 \| 1 \| 146 \| 1 \| 168 \| Lysozyme \| Lysozyme; afdb-uniprot50 \| AF-A0A3P6JLZ0-F1-MODEL\_V4 \| 1.0 \| 1.013e-08 \| 375 \| 0.35 \| 151 \| 87 \| 6 \| 1 \| 144 \| 144 \| 290 \| Lysozyme \| Lysozyme; afdb-uniprot50 \| AF-A0A1D9HAL1-F1-MODEL\_V4 \| 1.0 \| 4.964e-08 \| 374 \| 0.324 \| 145 \| 91 \| 4 \| 4 \| 145 \| 21 \| 161 \| Lysozyme \| Lysozyme; afdb-uniprot50 \| AF-A0A1W9JSH2-F1-MODEL\_V4 \| 1.0 \| 1.126e-08 \| 374 \| 0.289 \| 176 \| 88 \| 4 \| 2 \| 143 \| 8 \| 180 \| Lysozyme \| Lysozyme; afdb-uniprot50 \| AF-A0A0J1D667-F1-MODEL\_V4 \| 1.0 \| 4.574e-09 \| 373 \| 0.452 \| 137 \| 71 \| 2 \| 20 \| 154 \| 4 \| 138 \| Lysozyme \| Lysozyme; afdb-uniprot50 \| AF-A0A844D768-F1-MODEL\_V4 \| 1.0 \| 1.547e-08 \| 373 \| 0.277 \| 180 \| 91 \| 6 \| 3 \| 146 \| 1 \| 177 \| Glycoside hydrolase family protein \| Glycoside hydrolase family protein; afdb-uniprot50 \| AF-A0A2A6JMP3-F1-MODEL\_V4 \| 1.0 \| 4.707e-08 \| 373 \| 0.326 \| 150 \| 91 \| 4 \| 1 \| 144 \| 63 \| 208 \| Lysozyme \| Lysozyme; afdb-uniprot50 \| AF-A0A2W4UDA1-F1-MODEL\_V4 \| 1.0 \| 4.016e-08 \| 373 \| 0.301 \| 159 \| 100 \| 5 \| 1 \| 154 \| 9 \| 161 \| Lysozyme \| Lysozyme; afdb-uniprot50 \| AF-A0A6H1ZIA5-F1-MODEL\_V4 \| 1.0 \| 2.493e-08 \| 372 \| 0.307 \| 140 \| 89 \| 5 \| 6 \| 143 \| 11 \| 144 \| Lysozyme \| Lysozyme; afdb-uniprot50 \| AF-A0A4V1EHC6-F1-MODEL\_V4 \| 1.0 \| 2.126e-08 \| 372 \| 0.325 \| 160 \| 89 \| 4 \| 1 \| 145 \| 3 \| 158 \| Lysozyme \| Lysozyme; afdb-uniprot50 \| AF-A0A4P8HTQ2-F1-MODEL\_V4 \| 1.0 \| 5.234e-08 \| 372 \| 0.294 \| 153 \| 95 \| 3 \| 1 \| 144 \| 60 \| 208 \| Lysozyme \| Lysozyme; afdb-uniprot50 \| AF-A0A177NPS6-F1-MODEL\_V4 \| 1.0 \| 1.547e-08 \| 372 \| 0.343 \| 157 \| 89 \| 6 \| 1 \| 147 \| 90 \| 242 \| Lysozyme \| Lysozyme; afdb-uniprot50 \| AF-A0A7G8Q4E4-F1-MODEL\_V4 \| 1.0 \| 5.362e-09 \| 372 \| 0.343 \| 163 \| 84 \| 5 \| 1 \| 143 \| 4 \| 163 \| Lysozyme \| Lysozyme; afdb-uniprot50 \| AF-A0A7R9LHS2-F1-MODEL\_V4 \| 1.0 \| 4.964e-08 \| 372 \| 0.304 \| 148 \| 95 \| 5 \| 1 \| 144 \| 241 \| 384 \| Lysozyme \| Lysozyme; afdb-uniprot50 \| AF-A0A709FKY7-F1-MODEL\_V4 \| 1.0 \| 2.852e-07 \| 371 \| 0.595 \| 89 \| 36 \| 0 \| 56 \| 144 \| 2 \| 90 \| Lysozyme \| Lysozyme; afdb-uniprot50 \| AF-A0A316KGZ2-F1-MODEL\_V4 \| 1.0 \| 1.126e-08 \| 371 \| 0.459 \| 135 \| 62 \| 4 \| 1 \| 126 \| 1 \| 133 \| Lysozyme \| Lysozyme; afdb-uniprot50 \| AF-A0A5C7ZEJ8-F1-MODEL\_V4 \| 1.0 \| 3.425e-08 \| 371 \| 0.363 \| 143 \| 83 \| 6 \| 6 \| 145 \| 11 \| 148 \| Lysozyme \| Lysozyme; afdb-uniprot50 \| AF-A0A840DW28-F1-MODEL\_V4 \| 1.0 \| 4.464e-08 \| 370 \| 0.426 \| 129 \| 67 \| 2 \| 1 \| 125 \| 2 \| 127 \| GH24 family phage-related lysozyme (Muramidase) \| GH24 family phage-related lysozyme (Muramidase); afdb-uniprot50 \| AF-A0A7R9KMC6-F1-MODEL\_V4 \| 1.0 \| 9.374e-08 \| 370 \| 0.308 \| 149 \| 96 \| 4 \| 1 \| 146 \| 3 \| 147 \| Lysozyme \| Lysozyme; afdb-uniprot50 \| AF-A0A7X6IQC2-F1-MODEL\_V4 \| 1.0 \| 8.89e-08 \| 370 \| 0.342 \| 146 \| 85 \| 4 \| 7 \| 145 \| 19 \| 160 \| Lysozyme \| Lysozyme; afdb-uniprot50 \| AF-A0A2F0P4T0-F1-MODEL\_V4 \| 1.0 \| 3.249e-08 \| 369 \| 0.315 \| 149 \| 93 \| 4 \| 1 \| 146 \| 13 \| 155 \| Lysozyme \| Lysozyme; afdb-uniprot50 \| AF-A0A011RLS9-F1-MODEL\_V4 \| 1.0 \| 5.234e-08 \| 369 \| 0.308 \| 146 \| 94 \| 4 \| 1 \| 145 \| 12 \| 151 \| Lysozyme \| Lysozyme; afdb-uniprot50 \| AF-A0A379FFZ8-F1-MODEL\_V4 \| 1.0 \| 1.912e-08 \| 369 \| 0.349 \| 146 \| 85 \| 4 \| 5 \| 143 \| 15 \| 157 \| Lysozyme \| Lysozyme; afdb-uniprot50 \| AF-E2CPH6-F1-MODEL\_V4 \| 1.0 \| 4.234e-08 \| 369 \| 0.335 \| 161 \| 91 \| 6 \| 1 \| 149 \| 1 \| 157 \| Lysozyme \| Lysozyme; afdb-uniprot50 \| AF-A0A6C2CEI5-F1-MODEL\_V4 \| 1.0 \| 8.192e-09 \| 369 \| 0.386 \| 158 \| 78 \| 4 \| 1 \| 143 \| 463 \| 616 \| Lysozyme \| Lysozyme; afdb-uniprot50 \| AF-A0A2N0Q7R0-F1-MODEL\_V4 \| 1.0 \| 1.51e-07 \| 368 \| 0.46 \| 100 \| 54 \| 0 \| 49 \| 148 \| 17 \| 116 \| Lysozyme \| Lysozyme; afdb-uniprot50 \| AF-A0A6P4KAZ4-F1-MODEL\_V4 \| 1.0 \| 1.814e-08 \| 368 \| 0.375 \| 152 \| 81 \| 6 \| 6 \| 147 \| 7 \| 154 \| Lysozyme \| Lysozyme; afdb-uniprot50 \| AF-A0A0G3BMQ4-F1-MODEL\_V4 \| 1.0 \| 1.392e-08 \| 367 \| 0.314 \| 156 \| 90 \| 5 \| 1 \| 144 \| 6 \| 156 \| Lysozyme \| Lysozyme; afdb-uniprot50 \| AF-A0A5J4VHC4-F1-MODEL\_V4 \| 1.0 \| 8.192e-09 \| 367 \| 0.333 \| 165 \| 96 \| 7 \| 3 \| 157 \| 1 \| 161 \| Lysozyme \| Lysozyme; afdb-uniprot50 \| AF-A0A2A5BNU6-F1-MODEL\_V4 \| 1.0 \| 2.628e-08 \| 366 \| 0.35 \| 160 \| 82 \| 3 \| 1 \| 146 \| 1 \| 152 \| Lysozyme \| Lysozyme; afdb-uniprot50 \| AF-A0A707IK04-F1-MODEL\_V4 \| 1.0 \| 5.386e-07 \| 365 \| 0.611 \| 85 \| 33 \| 0 \| 60 \| 144 \| 5 \| 89 \| Lysozyme \| Lysozyme; afdb-uniprot50 \| AF-A0A1Y6D0R4-F1-MODEL\_V4 \| 1.0 \| 5.518e-08 \| 365 \| 0.295 \| 149 \| 99 \| 4 \| 1 \| 146 \| 5 \| 150 \| Lysozyme \| Lysozyme; afdb-uniprot50 \| AF-A0A6P4HYW2-F1-MODEL\_V4 \| 1.0 \| 9.108e-09 \| 365 \| 0.342 \| 152 \| 85 \| 6 \| 5 \| 144 \| 13 \| 161 \| Lysozyme \| Lysozyme; afdb-uniprot50 \| AF-A0A1G5SUL8-F1-MODEL\_V4 \| 1.0 \| 7.584e-08 \| 365 \| 0.282 \| 145 \| 96 \| 5 \| 4 \| 144 \| 28 \| 168 \| Lysozyme \| Lysozyme; afdb-uniprot50 \| AF-A0A708EKG9-F1-MODEL\_V4 \| 1.0 \| 3.525e-07 \| 364 \| 0.582 \| 91 \| 38 \| 0 \| 54 \| 144 \| 4 \| 94 \| Lysozyme \| Lysozyme; afdb-uniprot50 \| AF-A0A060NVJ0-F1-MODEL\_V4 \| 1.0 \| 1.013e-08 \| 364 \| 0.326 \| 156 \| 85 \| 6 \| 1 \| 143 \| 2 \| 150 \| Lysozyme \| Lysozyme; afdb-uniprot50 \| AF-A0A4S3KDY4-F1-MODEL\_V4 \| 1.0 \| 8.89e-08 \| 363 \| 0.25 \| 144 \| 103 \| 4 \| 4 \| 146 \| 1 \| 140 \| Lysozyme \| Lysozyme; afdb-uniprot50 \| AF-A0A3B9NE05-F1-MODEL\_V4 \| 1.0 \| 3.425e-08 \| 363 \| 0.32 \| 156 \| 93 \| 5 \| 1 \| 146 \| 1 \| 153 \| Lysozyme \| Lysozyme; afdb-uniprot50 \| AF-A0A3G3GKL6-F1-MODEL\_V4 \| 1.0 \| 1.068e-08 \| 363 \| 0.327 \| 171 \| 90 \| 7 \| 1 \| 153 \| 2 \| 165 \| Lysozyme \| Lysozyme; afdb-uniprot50 \| AF-A0A317CJ11-F1-MODEL\_V4 \| 1.0 \| 2.242e-08 \| 363 \| 0.337 \| 163 \| 85 \| 6 \| 1 \| 143 \| 2 \| 161 \| Lysozyme \| Lysozyme; afdb-uniprot50 \| AF-A0A538JLK8-F1-MODEL\_V4 \| 1.0 \| 3.249e-08 \| 363 \| 0.293 \| 160 \| 93 \| 5 \| 3 \| 146 \| 149 \| 304 \| Lysozyme \| Lysozyme; afdb-uniprot50 \| AF-A0A1X9TLR2-F1-MODEL\_V4 \| 1.0 \| 7.996e-08 \| 362 \| 0.317 \| 145 \| 95 \| 3 \| 1 \| 145 \| 26 \| 166 \| Lysozyme \| Lysozyme; afdb-uniprot50 \| AF-A0A849CLB2-F1-MODEL\_V4 \| 1.0 \| 4.016e-08 \| 362 \| 0.346 \| 147 \| 84 \| 4 \| 4 \| 144 \| 30 \| 170 \| Lysozyme \| Lysozyme; afdb-uniprot50 \| AF-A0A7U3CRJ8-F1-MODEL\_V4 \| 1.0 \| 1.912e-08 \| 362 \| 0.315 \| 152 \| 88 \| 5 \| 5 \| 144 \| 37 \| 184 \| Lysozyme \| Lysozyme; afdb-uniprot50 \| AF-A0A7W5W268-F1-MODEL\_V4 \| 1.0 \| 9.374e-08 \| 362 \| 0.335 \| 158 \| 89 \| 3 \| 1 \| 146 \| 56 \| 209 \| Lysozyme \| Lysozyme; afdb-uniprot50 \| AF-A0A2A7MJ10-F1-MODEL\_V4 \| 1.0 \| 2.017e-08 \| 362 \| 0.335 \| 152 \| 88 \| 6 \| 2 \| 145 \| 103 \| 249 \| Lysozyme \| Lysozyme; afdb-uniprot50 \| AF-A0A081C6S4-F1-MODEL\_V4 \| 1.0 \| 1.912e-08 \| 362 \| 0.333 \| 159 \| 89 \| 4 \| 1 \| 145 \| 1 \| 156 \| Lysozyme \| Lysozyme; afdb-uniprot50 \| AF-A0A242N7A0-F1-MODEL\_V4 \| 1.0 \| 7.402e-07 \| 361 \| 0.548 \| 82 \| 37 \| 0 \| 64 \| 145 \| 2 \| 83 \| Lysozyme \| Lysozyme; afdb-uniprot50 \| AF-A0A3D4K8L3-F1-MODEL\_V4 \| 1.0 \| 7.192e-08 \| 361 \| 0.357 \| 126 \| 71 \| 4 \| 1 \| 121 \| 2 \| 122 \| Lysozyme \| Lysozyme; afdb-uniprot50 \| AF-A0A0N7KV59-F1-MODEL\_V4 \| 1.0 \| 3.081e-08 \| 361 \| 0.326 \| 153 \| 89 \| 6 \| 6 \| 147 \| 8 \| 157 \| Lysozyme \| Lysozyme; afdb-uniprot50 \| AF-A0A1F4MGX4-F1-MODEL\_V4 \| 1.0 \| 3.612e-08 \| 361 \| 0.301 \| 166 \| 96 \| 4 \| 1 \| 146 \| 12 \| 177 \| Lysozyme \| Lysozyme; afdb-uniprot50 \| AF-A0A523FRG2-F1-MODEL\_V4 \| 1.0 \| 1.432e-07 \| 361 \| 0.313 \| 150 \| 95 \| 5 \| 1 \| 146 \| 65 \| 210 \| Lysozyme \| Lysozyme; afdb-uniprot50 \| AF-A0A2G6ENY7-F1-MODEL\_V4 \| 1.0 \| 1.631e-08 \| 361 \| 0.358 \| 145 \| 80 \| 3 \| 6 \| 143 \| 6 \| 144 \| Lysozyme \| Lysozyme; afdb-uniprot50 \| AF-A0A399R7V2-F1-MODEL\_V4 \| 1.0 \| 6.135e-08 \| 361 \| 0.362 \| 138 \| 79 \| 6 \| 14 \| 146 \| 3 \| 136 \| Lysozyme \| Lysozyme; afdb-uniprot50 \| AF-A0A640WAD0-F1-MODEL\_V4 \| 1.0 \| 6.821e-08 \| 360 \| 0.432 \| 125 \| 67 \| 3 \| 20 \| 144 \| 2 \| 122 \| Lysozyme \| Lysozyme; afdb-uniprot50 \| AF-A0A853WB76-F1-MODEL\_V4 \| 1.0 \| 6.135e-08 \| 360 \| 0.359 \| 128 \| 77 \| 3 \| 4 \| 129 \| 21 \| 145 \| Endolysin \| Endolysin; afdb-uniprot50 \| AF-A0A6G3Z5Y0-F1-MODEL\_V4 \| 1.0 \| 9.374e-08 \| 360 \| 0.482 \| 112 \| 51 \| 3 \| 2 \| 109 \| 105 \| 213 \| Lysozyme \| Lysozyme; afdb-uniprot50 \| AF-A0A1V1PUV1-F1-MODEL\_V4 \| 1.0 \| 1.042e-07 \| 360 \| 0.315 \| 146 \| 92 \| 4 \| 3 \| 144 \| 6 \| 147 \| Lysozyme \| Lysozyme; afdb-uniprot50 \| AF-K9XRH0-F1-MODEL\_V4 \| 1.0 \| 2.307e-07 \| 360 \| 0.478 \| 119 \| 58 \| 3 \| 2 \| 120 \| 67 \| 181 \| Lysozyme \| Lysozyme; afdb-uniprot50 \| AF-A0A1U9JSH0-F1-MODEL\_V4 \| 1.0 \| 3.612e-08 \| 360 \| 0.31 \| 148 \| 96 \| 3 \| 2 \| 145 \| 334 \| 479 \| Lysozyme \| Lysozyme; afdb-uniprot50 \| AF-Q0ANV5-F1-MODEL\_V4 \| 1.0 \| 2.922e-08 \| 360 \| 0.283 \| 148 \| 91 \| 5 \| 1 \| 144 \| 1 \| 137 \| Glycoside hydrolase, family 24 \| Glycoside hydrolase, family 24; afdb-uniprot50 \| AF-A0A3D6A2G8-F1-MODEL\_V4 \| 1.0 \| 2.242e-08 \| 359 \| 0.301 \| 146 \| 93 \| 4 \| 3 \| 144 \| 7 \| 147 \| Lysozyme \| Lysozyme; afdb-uniprot50 \| AF-A0A7W4IK65-F1-MODEL\_V4 \| 1.0 \| 3.808e-08 \| 359 \| 0.333 \| 156 \| 90 \| 6 \| 1 \| 145 \| 1 \| 153 \| Lysozyme \| Lysozyme; afdb-uniprot50 \| AF-A0A1E4ZVU3-F1-MODEL\_V4 \| 1.0 \| 1.912e-08 \| 359 \| 0.318 \| 160 \| 94 \| 6 \| 1 \| 150 \| 13 \| 167 \| Lysozyme \| Lysozyme; afdb-uniprot50 \| AF-A0A852Q327-F1-MODEL\_V4 \| 1.0 \| 1.042e-07 \| 359 \| 0.34 \| 144 \| 89 \| 3 \| 4 \| 144 \| 30 \| 170 \| Lysozyme \| Lysozyme; afdb-uniprot50 \| AF-A0A0F4VEW5-F1-MODEL\_V4 \| 1.0 \| 2.628e-08 \| 358 \| 0.4 \| 130 \| 70 \| 2 \| 20 \| 144 \| 12 \| 138 \| Lysozyme \| Lysozyme; afdb-uniprot50 \| AF-A0A6M3XN87-F1-MODEL\_V4 \| 1.0 \| 3.249e-08 \| 358 \| 0.318 \| 154 \| 86 \| 4 \| 6 \| 144 \| 16 \| 165 \| Lysozyme \| Lysozyme; afdb-uniprot50 \| AF-A0A103RZ68-F1-MODEL\_V4 \| 1.0 \| 1.467e-08 \| 358 \| 0.295 \| 166 \| 88 \| 7 \| 5 \| 144 \| 15 \| 177 \| Lysozyme \| Lysozyme; afdb-uniprot50 \| AF-A0A511F0V7-F1-MODEL\_V4 \| 1.0 \| 1.77e-07 \| 358 \| 0.309 \| 139 \| 93 \| 3 \| 7 \| 144 \| 46 \| 182 \| Lysozyme \| Lysozyme; afdb-uniprot50 \| AF-A0A358B152-F1-MODEL\_V4 \| 1.0 \| 2.188e-07 \| 357 \| 0.271 \| 140 \| 97 \| 4 \| 5 \| 143 \| 3 \| 138 \| Lysozyme \| Lysozyme; afdb-uniprot50 \| AF-A0A1M7YJN3-F1-MODEL\_V4 \| 1.0 \| 1.099e-07 \| 357 \| 0.287 \| 146 \| 97 \| 4 \| 1 \| 144 \| 1 \| 141 \| Lysozyme \| Lysozyme; afdb-uniprot50 \| AF-A0A3D3NAZ3-F1-MODEL\_V4 \| 1.0 \| 4.707e-08 \| 357 \| 0.305 \| 144 \| 89 \| 3 \| 10 \| 145 \| 2 \| 142 \| Lysozyme \| Lysozyme; afdb-uniprot50 \| AF-A0A4Q2QRI9-F1-MODEL\_V4 \| 1.0 \| 1.631e-08 \| 357 \| 0.339 \| 159 \| 83 \| 6 \| 1 \| 145 \| 293 \| 443 \| Lysozyme \| Lysozyme; afdb-uniprot50 \| AF-A0A1A7PVK4-F1-MODEL\_V4 \| 1.0 \| 7.996e-08 \| 356 \| 0.328 \| 152 \| 90 \| 4 \| 1 \| 146 \| 27 \| 172 \| Lysozyme \| Lysozyme; afdb-uniprot50 \| AF-G1UXF8-F1-MODEL\_V4 \| 1.0 \| 2.017e-08 \| 356 \| 0.34 \| 147 \| 82 \| 7 \| 7 \| 145 \| 31 \| 170 \| Lysozyme \| Lysozyme; afdb-uniprot50 \| AF-A0A3D3KGU8-F1-MODEL\_V4 \| 1.0 \| 7.584e-08 \| 356 \| 0.335 \| 155 \| 89 \| 6 \| 1 \| 144 \| 53 \| 204 \| Lysozyme \| Lysozyme; afdb-uniprot50 \| AF-A0A0U5ETJ2-F1-MODEL\_V4 \| 1.0 \| 2.771e-08 \| 356 \| 0.329 \| 155 \| 88 \| 6 \| 5 \| 148 \| 42 \| 191 \| Lysozyme \| Lysozyme; afdb-uniprot50 \| AF-A0A386TCM0-F1-MODEL\_V4 \| 1.0 \| 1.159e-07 \| 356 \| 0.312 \| 144 \| 90 \| 4 \| 6 \| 144 \| 142 \| 281 \| Lysozyme \| Lysozyme; afdb-uniprot50 \| AF-A0A023PRG0-F1-MODEL\_V4 \| 1.0 \| 1.467e-08 \| 356 \| 0.341 \| 155 \| 86 \| 6 \| 3 \| 143 \| 816 \| 968 \| Lysozyme \| Lysozyme; afdb-uniprot50 \| AF-I3DCN5-F1-MODEL\_V4 \| 1.0 \| 1.432e-07 \| 355 \| 0.324 \| 148 \| 93 \| 4 \| 3 \| 146 \| 29 \| 173 \| Lysozyme \| Lysozyme; afdb-uniprot50 \| AF-A0A0V8BQ91-F1-MODEL\_V4 \| 1.0 \| 1.72e-08 \| 355 \| 0.367 \| 158 \| 77 \| 6 \| 1 \| 147 \| 1 \| 146 \| Lysozyme \| Lysozyme; afdb-uniprot50 \| AF-A0A1Q9QMU9-F1-MODEL\_V4 \| 1.0 \| 4.016e-08 \| 354 \| 0.263 \| 152 \| 100 \| 6 \| 1 \| 144 \| 5 \| 152 \| Lysozyme \| Lysozyme; afdb-uniprot50 \| AF-A0A7U2KR66-F1-MODEL\_V4 \| 1.0 \| 2.922e-08 \| 354 \| 0.283 \| 155 \| 96 \| 5 \| 2 \| 145 \| 21 \| 171 \| Lysozyme \| Lysozyme; afdb-uniprot50 \| AF-E6VL46-F1-MODEL\_V4 \| 1.0 \| 5.819e-08 \| 354 \| 0.34 \| 150 \| 90 \| 3 \| 2 \| 144 \| 8 \| 155 \| Lysozyme \| Lysozyme; afdb-uniprot50 \| AF-A0A357YYC3-F1-MODEL\_V4 \| 1.0 \| 1.679e-07 \| 354 \| 0.337 \| 151 \| 89 \| 4 \| 1 \| 143 \| 58 \| 205 \| Lysozyme \| Lysozyme; afdb-uniprot50 \| AF-A0A430BCR4-F1-MODEL\_V4 \| 1.0 \| 3.343e-07 \| 354 \| 0.27 \| 144 \| 97 \| 4 \| 5 \| 145 \| 111 \| 249 \| Lysozyme \| Lysozyme; afdb-uniprot50 \| AF-A0A6N8JD00-F1-MODEL\_V4 \| 1.0 \| 2.242e-08 \| 354 \| 0.331 \| 160 \| 91 \| 5 \| 1 \| 147 \| 225 \| 381 \| Lysozyme \| Lysozyme; afdb-uniprot50 \| AF-A0A009G8D4-F1-MODEL\_V4 \| 1.0 \| 7.584e-08 \| 353 \| 0.535 \| 112 \| 47 \| 1 \| 35 \| 146 \| 8 \| 114 \| Lysozyme \| Lysozyme; afdb-uniprot50 \| AF-A0A7J0ADD2-F1-MODEL\_V4 \| 1.0 \| 1.51e-07 \| 353 \| 0.314 \| 143 \| 88 \| 6 \| 7 \| 144 \| 2 \| 139 \| Lysozyme \| Lysozyme; afdb-uniprot50 \| AF-A0A5C7LX91-F1-MODEL\_V4 \| 1.0 \| 4.964e-08 \| 353 \| 0.342 \| 149 \| 87 \| 6 \| 3 \| 146 \| 10 \| 152 \| Lysozyme \| Lysozyme; afdb-uniprot50 \| AF-A0A401JF78-F1-MODEL\_V4 \| 1.0 \| 1.72e-08 \| 352 \| 0.341 \| 155 \| 87 \| 6 \| 1 \| 146 \| 2 \| 150 \| Lysozyme \| Lysozyme; afdb-uniprot50 \| AF-A0A4R3JY04-F1-MODEL\_V4 \| 1.0 \| 1.392e-08 \| 352 \| 0.32 \| 159 \| 90 \| 6 \| 1 \| 147 \| 6 \| 158 \| Lysozyme \| Lysozyme; afdb-uniprot50 \| AF-A3MZM3-F1-MODEL\_V4 \| 1.0 \| 1.592e-07 \| 352 \| 0.28 \| 146 \| 98 \| 3 \| 4 \| 145 \| 36 \| 178 \| Lysozyme \| Lysozyme; afdb-uniprot50 \| AF-A0A341EDR9-F1-MODEL\_V4 \| 1.0 \| 3.425e-08 \| 352 \| 0.259 \| 154 \| 99 \| 5 \| 2 \| 144 \| 133 \| 282 \| Lysozyme \| Lysozyme; afdb-uniprot50 \| AF-A0A7W0Z8H7-F1-MODEL\_V4 \| 1.0 \| 2.017e-08 \| 352 \| 0.357 \| 154 \| 83 \| 4 \| 1 \| 144 \| 69 \| 216 \| Lysozyme \| Lysozyme; afdb-uniprot50 \| AF-A0A0F8Z6L9-F1-MODEL\_V4 \| 1.0 \| 6.469e-08 \| 352 \| 0.352 \| 153 \| 86 \| 5 \| 1 \| 143 \| 179 \| 328 \| Lysozyme \| Lysozyme; afdb-uniprot50 \| AF-A0A0Q5GY79-F1-MODEL\_V4 \| 1.0 \| 7.192e-08 \| 352 \| 0.271 \| 173 \| 99 \| 7 \| 1 \| 155 \| 1 \| 164 \| PG\_binding\_1 domain-containing protein \| PG\_binding\_1 domain-containing protein; afdb-uniprot50 \| AF-A0A375GDD6-F1-MODEL\_V4 \| 1.0 \| 1.042e-07 \| 351 \| 0.321 \| 143 \| 88 \| 4 \| 6 \| 142 \| 13 \| 152 \| Lysozyme \| Lysozyme; afdb-uniprot50 \| AF-A0A022G329-F1-MODEL\_V4 \| 1.0 \| 1.592e-07 \| 351 \| 0.303 \| 145 \| 94 \| 4 \| 4 \| 145 \| 15 \| 155 \| Lysozyme \| Lysozyme; afdb-uniprot50 \| AF-A0A7W9RA22-F1-MODEL\_V4 \| 1.0 \| 1.288e-07 \| 351 \| 0.293 \| 143 \| 93 \| 5 \| 5 \| 145 \| 28 \| 164 \| Lysozyme \| Lysozyme; afdb-uniprot50 \| AF-A0A0F9TJ14-F1-MODEL\_V4 \| 1.0 \| 4.707e-08 \| 351 \| 0.3 \| 150 \| 89 \| 5 \| 5 \| 143 \| 20 \| 164 \| Lysozyme \| Lysozyme; afdb-uniprot50 \| AF-A0A7X5Z2P4-F1-MODEL\_V4 \| 1.0 \| 4.016e-08 \| 351 \| 0.287 \| 146 \| 97 \| 4 \| 3 \| 144 \| 13 \| 155 \| Lysozyme \| Lysozyme; afdb-uniprot50 \| AF-X6L4V5-F1-MODEL\_V4 \| 1.0 \| 3.425e-08 \| 350 \| 0.315 \| 152 \| 88 \| 5 \| 5 \| 144 \| 124 \| 271 \| Lysozyme \| Lysozyme; afdb-uniprot50 \| AF-A0A1I1VPY7-F1-MODEL\_V4 \| 1.0 \| 3.612e-08 \| 350 \| 0.326 \| 156 \| 89 \| 6 \| 1 \| 143 \| 407 \| 559 \| Lysozyme \| Lysozyme; afdb-uniprot50 \| AF-A0A381FGI3-F1-MODEL\_V4 \| 1.0 \| 4.234e-08 \| 349 \| 0.251 \| 151 \| 105 \| 4 \| 6 \| 151 \| 8 \| 155 \| Lysozyme \| Lysozyme; afdb-uniprot50 \| AF-A0A752A1Q8-F1-MODEL\_V4 \| 1.0 \| 1.51e-07 \| 348 \| 0.319 \| 144 \| 89 \| 4 \| 7 \| 144 \| 19 \| 159 \| Lysozyme \| Lysozyme; afdb-uniprot50 \| AF-A0A285J525-F1-MODEL\_V4 \| 1.0 \| 4.234e-08 \| 348 \| 0.328 \| 152 \| 87 \| 6 \| 4 \| 144 \| 96 \| 243 \| Lysozyme \| Lysozyme; afdb-uniprot50 \| AF-A0A4P7EAC4-F1-MODEL\_V4 \| 1.0 \| 5.234e-08 \| 348 \| 0.297 \| 148 \| 93 \| 6 \| 4 \| 144 \| 95 \| 238 \| Lysozyme \| Lysozyme; afdb-uniprot50 \| AF-A0A2S5N400-F1-MODEL\_V4 \| 1.0 \| 1.867e-07 \| 348 \| 0.395 \| 149 \| 76 \| 6 \| 1 \| 144 \| 4 \| 143 \| Lysozyme \| Lysozyme; afdb-uniprot50 \| AF-A0A0C1GCU4-F1-MODEL\_V4 \| 1.0 \| 7.192e-08 \| 347 \| 0.263 \| 148 \| 98 \| 5 \| 3 \| 143 \| 1 \| 144 \| Lysozyme \| Lysozyme; afdb-uniprot50 \| AF-A0A2P5N8Y5-F1-MODEL\_V4 \| 1.0 \| 4.016e-08 \| 347 \| 0.303 \| 158 \| 93 \| 5 \| 1 \| 143 \| 9 \| 164 \| Lysozyme \| Lysozyme; afdb-uniprot50 \| AF-A0A2U1UUN7-F1-MODEL\_V4 \| 1.0 \| 1.358e-07 \| 347 \| 0.284 \| 151 \| 97 \| 3 \| 2 \| 146 \| 36 \| 181 \| Lysozyme \| Lysozyme; afdb-uniprot50 \| AF-A0A431NTU0-F1-MODEL\_V4 \| 1.0 \| 6.135e-08 \| 347 \| 0.293 \| 167 \| 97 \| 6 \| 1 \| 150 \| 1 \| 163 \| Lysozyme \| Lysozyme; afdb-uniprot50 \| AF-G8AQH1-F1-MODEL\_V4 \| 1.0 \| 8.432e-08 \| 346 \| 0.474 \| 118 \| 57 \| 3 \| 33 \| 146 \| 3 \| 119 \| Lysozyme \| Lysozyme; afdb-uniprot50 \| AF-A0A1H1ART9-F1-MODEL\_V4 \| 1.0 \| 6.469e-08 \| 346 \| 0.326 \| 153 \| 89 \| 6 \| 6 \| 147 \| 14 \| 163 \| Lysozyme \| Lysozyme; afdb-uniprot50 \| AF-S3UE20-F1-MODEL\_V4 \| 1.0 \| 1.193e-06 \| 345 \| 0.563 \| 87 \| 38 \| 0 \| 60 \| 146 \| 6 \| 92 \| Lysozyme \| Lysozyme; afdb-uniprot50 \| AF-A0A366HBV8-F1-MODEL\_V4 \| 1.0 \| 6.469e-08 \| 345 \| 0.301 \| 146 \| 91 \| 6 \| 6 \| 144 \| 11 \| 152 \| Lysozyme \| Lysozyme; afdb-uniprot50 \| AF-A0A143DGD6-F1-MODEL\_V4 \| 1.0 \| 9.374e-08 \| 345 \| 0.311 \| 151 \| 90 \| 4 \| 4 \| 144 \| 17 \| 163 \| Lysozyme \| Lysozyme; afdb-uniprot50 \| AF-A0A564WD90-F1-MODEL\_V4 \| 1.0 \| 4.234e-08 \| 345 \| 0.317 \| 170 \| 86 \| 5 \| 1 \| 144 \| 1 \| 166 \| Lysozyme \| Lysozyme; afdb-uniprot50 \| AF-A0A345PAS8-F1-MODEL\_V4 \| 1.0 \| 2.075e-07 \| 345 \| 0.291 \| 144 \| 99 \| 2 \| 4 \| 147 \| 31 \| 171 \| Lysozyme \| Lysozyme; afdb-uniprot50 \| AF-A0A495HUG2-F1-MODEL\_V4 \| 1.0 \| 8.89e-08 \| 345 \| 0.272 \| 169 \| 102 \| 4 \| 1 \| 148 \| 11 \| 179 \| GH24 family phage-related lysozyme (Muramidase) \| GH24 family phage-related lysozyme (Muramidase); afdb-uniprot50 \| AF-A0A702E4W0-F1-MODEL\_V4 \| 1.0 \| 3.612e-08 \| 345 \| 0.346 \| 156 \| 82 \| 5 \| 1 \| 143 \| 121 \| 269 \| Lysozyme \| Lysozyme; afdb-uniprot50 \| AF-A3UJS2-F1-MODEL\_V4 \| 1.0 \| 4.707e-08 \| 345 \| 0.285 \| 161 \| 101 \| 5 \| 1 \| 157 \| 5 \| 155 \| Lysozyme \| Lysozyme; afdb-uniprot50 \| AF-A0A6L6YDR0-F1-MODEL\_V4 \| 1.0 \| 8.89e-08 \| 344 \| 0.321 \| 143 \| 91 \| 4 \| 4 \| 144 \| 9 \| 147 \| Lysozyme \| Lysozyme; afdb-uniprot50 \| AF-A0A5C7SP49-F1-MODEL\_V4 \| 1.0 \| 7.584e-08 \| 344 \| 0.317 \| 148 \| 90 \| 7 \| 5 \| 146 \| 18 \| 160 \| Lysozyme \| Lysozyme; afdb-uniprot50 \| AF-V7DA39-F1-MODEL\_V4 \| 1.0 \| 2.922e-08 \| 343 \| 0.283 \| 166 \| 94 \| 4 \| 1 \| 144 \| 1 \| 163 \| Lysozyme \| Lysozyme; afdb-uniprot50 \| AF-A0A4S3M508-F1-MODEL\_V4 \| 1.0 \| 1.77e-07 \| 343 \| 0.344 \| 151 \| 90 \| 5 \| 5 \| 151 \| 18 \| 163 \| Lysozyme \| Lysozyme; afdb-uniprot50 \| AF-A0A521TYU1-F1-MODEL\_V4 \| 1.0 \| 7.996e-08 \| 343 \| 0.343 \| 160 \| 85 \| 7 \| 1 \| 144 \| 10 \| 165 \| Lysozyme \| Lysozyme; afdb-uniprot50 \| AF-A0A6L8KFJ1-F1-MODEL\_V4 \| 1.0 \| 1.288e-07 \| 343 \| 0.293 \| 143 \| 92 \| 4 \| 6 \| 142 \| 43 \| 182 \| Lysozyme \| Lysozyme; afdb-uniprot50 \| AF-A0A442BB57-F1-MODEL\_V4 \| 1.0 \| 1.77e-07 \| 343 \| 0.3 \| 153 \| 94 \| 3 \| 1 \| 143 \| 41 \| 190 \| Lysozyme \| Lysozyme; afdb-uniprot50 \| AF-A0A1H2EW19-F1-MODEL\_V4 \| 1.0 \| 8.432e-08 \| 343 \| 0.306 \| 176 \| 98 \| 8 \| 1 \| 157 \| 1 \| 171 \| Lysozyme \| Lysozyme; afdb-uniprot50 \| AF-A0A2U2BR73-F1-MODEL\_V4 \| 1.0 \| 4.964e-08 \| 343 \| 0.31 \| 158 \| 98 \| 5 \| 1 \| 154 \| 5 \| 155 \| Lysozyme \| Lysozyme; afdb-uniprot50 \| AF-A0A521ZXH5-F1-MODEL\_V4 \| 1.0 \| 3.808e-08 \| 342 \| 0.298 \| 151 \| 91 \| 6 \| 3 \| 144 \| 2 \| 146 \| Lysozyme \| Lysozyme; afdb-uniprot50 \| AF-B1ZJD5-F1-MODEL\_V4 \| 1.0 \| 1.592e-07 \| 342 \| 0.331 \| 145 \| 86 \| 6 \| 7 \| 146 \| 41 \| 179 \| Lysozyme \| Lysozyme; afdb-uniprot50 \| AF-A0A845GU55-F1-MODEL\_V4 \| 1.0 \| 1.51e-07 \| 342 \| 0.32 \| 159 \| 92 \| 6 \| 1 \| 147 \| 57 \| 211 \| Glycoside hydrolase family protein \| Glycoside hydrolase family protein; afdb-uniprot50 \| AF-J1JG31-F1-MODEL\_V4 \| 1.0 \| 2.565e-07 \| 341 \| 0.447 \| 123 \| 64 \| 1 \| 32 \| 150 \| 10 \| 132 \| Lysozyme \| Lysozyme; afdb-uniprot50 \| AF-A0A377M831-F1-MODEL\_V4 \| 1.0 \| 1.592e-07 \| 340 \| 0.413 \| 133 \| 69 \| 4 \| 1 \| 127 \| 6 \| 135 \| Lysozyme \| Lysozyme; afdb-uniprot50 \| AF-A0A519QHC5-F1-MODEL\_V4 \| 1.0 \| 7.996e-08 \| 340 \| 0.346 \| 147 \| 82 \| 7 \| 7 \| 143 \| 6 \| 148 \| Lysozyme \| Lysozyme; afdb-uniprot50 \| AF-A0A501W9K6-F1-MODEL\_V4 \| 1.0 \| 6.821e-08 \| 340 \| 0.313 \| 153 \| 92 \| 7 \| 1 \| 144 \| 8 \| 156 \| Lysozyme \| Lysozyme; afdb-uniprot50 \| AF-A0A512JI05-F1-MODEL\_V4 \| 1.0 \| 3.007e-07 \| 340 \| 0.354 \| 144 \| 84 \| 5 \| 3 \| 146 \| 7 \| 141 \| Lysozyme \| Lysozyme; afdb-uniprot50 \| AF-A0A1L3I5D1-F1-MODEL\_V4 \| 1.0 \| 1.432e-07 \| 340 \| 0.291 \| 144 \| 90 \| 5 \| 5 \| 140 \| 94 \| 233 \| Lysozyme \| Lysozyme; afdb-uniprot50 \| AF-G0JRZ2-F1-MODEL\_V4 \| 1.0 \| 7.192e-08 \| 340 \| 0.32 \| 156 \| 92 \| 4 \| 1 \| 143 \| 156 \| 310 \| Lysozyme \| Lysozyme; afdb-uniprot50 \| AF-A0A5C7PGH8-F1-MODEL\_V4 \| 1.0 \| 1.159e-07 \| 339 \| 0.301 \| 156 \| 92 \| 8 \| 1 \| 143 \| 1 \| 152 \| Lysozyme \| Lysozyme; afdb-uniprot50 \| AF-A0A431M8G9-F1-MODEL\_V4 \| 1.0 \| 1.679e-07 \| 339 \| 0.313 \| 153 \| 92 \| 4 \| 3 \| 144 \| 11 \| 161 \| Lysozyme \| Lysozyme; afdb-uniprot50 \| AF-A0A442XXC8-F1-MODEL\_V4 \| 1.0 \| 1.77e-07 \| 339 \| 0.292 \| 147 \| 92 \| 6 \| 6 \| 144 \| 33 \| 175 \| Lysozyme \| Lysozyme; afdb-uniprot50 \| AF-A0A2A7MJ58-F1-MODEL\_V4 \| 1.0 \| 1.222e-07 \| 339 \| 0.322 \| 149 \| 92 \| 5 \| 2 \| 144 \| 134 \| 279 \| Lysozyme \| Lysozyme; afdb-uniprot50 \| AF-J0QIK5-F1-MODEL\_V4 \| 1.0 \| 1.679e-07 \| 339 \| 0.45 \| 120 \| 61 \| 2 \| 33 \| 147 \| 376 \| 495 \| Lysozyme \| Lysozyme; afdb-uniprot50 \| AF-A0A7L5Y684-F1-MODEL\_V4 \| 1.0 \| 7.584e-08 \| 338 \| 0.309 \| 155 \| 92 \| 5 \| 1 \| 144 \| 20 \| 170 \| Lysozyme \| Lysozyme; afdb-uniprot50 \| AF-A0A7H0VBA6-F1-MODEL\_V4 \| 1.0 \| 1.679e-07 \| 338 \| 0.337 \| 151 \| 88 \| 6 \| 1 \| 146 \| 38 \| 181 \| Lysozyme \| Lysozyme; afdb-uniprot50 \| AF-A0A450X4Q4-F1-MODEL\_V4 \| 1.0 \| 3.92e-07 \| 338 \| 0.296 \| 155 \| 96 \| 4 \| 1 \| 145 \| 63 \| 214 \| Lysozyme \| Lysozyme; afdb-uniprot50 \| AF-A0A2E0EKN3-F1-MODEL\_V4 \| 1.0 \| 7.192e-08 \| 337 \| 0.289 \| 169 \| 104 \| 7 \| 1 \| 157 \| 11 \| 175 \| Lysozyme \| Lysozyme; afdb-uniprot50 \| AF-A0A368HDJ6-F1-MODEL\_V4 \| 1.0 \| 1.068e-08 \| 336 \| 0.392 \| 158 \| 84 \| 5 \| 2 \| 159 \| 6 \| 151 \| Lysozyme \| Lysozyme; afdb-uniprot50 \| AF-M1WVJ0-F1-MODEL\_V4 \| 1.0 \| 6.135e-08 \| 336 \| 0.312 \| 160 \| 87 \| 5 \| 1 \| 143 \| 318 \| 471 \| Lysozyme \| Lysozyme; afdb-uniprot50 \| AF-A0A257SN27-F1-MODEL\_V4 \| 1.0 \| 1.554e-06 \| 335 \| 0.526 \| 93 \| 44 \| 0 \| 53 \| 145 \| 1 \| 93 \| Lysozyme \| Lysozyme; afdb-uniprot50 \| AF-A0A2G3JY01-F1-MODEL\_V4 \| 1.0 \| 3.525e-07 \| 335 \| 0.243 \| 152 \| 102 \| 3 \| 6 \| 145 \| 18 \| 168 \| Lysozyme \| Lysozyme; afdb-uniprot50 \| AF-A0A7V8IN83-F1-MODEL\_V4 \| 1.0 \| 1.592e-07 \| 335 \| 0.243 \| 148 \| 103 \| 4 \| 6 \| 144 \| 23 \| 170 \| Lysozyme \| Lysozyme; afdb-uniprot50 \| AF-A0A1E4GWM4-F1-MODEL\_V4 \| 1.0 \| 4.358e-07 \| 335 \| 0.268 \| 145 \| 95 \| 6 \| 5 \| 144 \| 9 \| 147 \| Lysozyme \| Lysozyme; afdb-uniprot50 \| AF-A0A5A8A1Q9-F1-MODEL\_V4 \| 1.0 \| 3.343e-07 \| 335 \| 0.355 \| 152 \| 86 \| 5 \| 1 \| 148 \| 1 \| 144 \| Lysozyme \| Lysozyme; afdb-uniprot50 \| AF-E5YB66-F1-MODEL\_V4 \| 1.0 \| 5.234e-08 \| 334 \| 0.333 \| 144 \| 81 \| 7 \| 10 \| 145 \| 2 \| 138 \| Lysozyme \| Lysozyme; afdb-uniprot50 \| AF-A0A5C8TXF5-F1-MODEL\_V4 \| 1.0 \| 4.595e-07 \| 334 \| 0.368 \| 141 \| 80 \| 5 \| 3 \| 143 \| 11 \| 142 \| Lysozyme \| Lysozyme; afdb-uniprot50 \| AF-A0A2N1Y888-F1-MODEL\_V4 \| 1.0 \| 1.159e-07 \| 334 \| 0.287 \| 167 \| 96 \| 8 \| 3 \| 150 \| 13 \| 175 \| Lysozyme \| Lysozyme; afdb-uniprot50 \| AF-A0A7Z6L4X9-F1-MODEL\_V4 \| 1.0 \| 1.679e-07 \| 334 \| 0.351 \| 162 \| 93 \| 7 \| 1 \| 156 \| 3 \| 158 \| Lysozyme \| Lysozyme; afdb-uniprot50 \| AF-A0A4S2H276-F1-MODEL\_V4 \| 1.0 \| 6.135e-08 \| 334 \| 0.29 \| 155 \| 99 \| 5 \| 3 \| 153 \| 7 \| 154 \| Lysozyme \| Lysozyme; afdb-uniprot50 \| AF-A0A1V2N8E9-F1-MODEL\_V4 \| 1.0 \| 3.717e-07 \| 333 \| 0.354 \| 127 \| 77 \| 2 \| 22 \| 146 \| 1 \| 124 \| Lysozyme \| Lysozyme; afdb-uniprot50 \| AF-A0A7V8QYJ7-F1-MODEL\_V4 \| 1.0 \| 1.099e-07 \| 333 \| 0.246 \| 158 \| 101 \| 6 \| 1 \| 144 \| 2 \| 155 \| Lysozyme \| Lysozyme; afdb-uniprot50 \| AF-A0A3C1GCQ9-F1-MODEL\_V4 \| 1.0 \| 6.821e-08 \| 333 \| 0.354 \| 158 \| 82 \| 9 \| 1 \| 145 \| 4 \| 154 \| Lysozyme \| Lysozyme; afdb-uniprot50 \| AF-A0A7L9RU02-F1-MODEL\_V4 \| 1.0 \| 2.075e-07 \| 333 \| 0.31 \| 145 \| 92 \| 5 \| 4 \| 144 \| 36 \| 176 \| Lysozyme RrrD \| Lysozyme RrrD; afdb-uniprot50 \| AF-A0A366C3Q8-F1-MODEL\_V4 \| 1.0 \| 3.343e-07 \| 333 \| 0.251 \| 147 \| 100 \| 5 \| 6 \| 147 \| 47 \| 188 \| Lysozyme \| Lysozyme; afdb-uniprot50 \| AF-A0A080LZI7-F1-MODEL\_V4 \| 1.0 \| 1.679e-07 \| 333 \| 0.352 \| 153 \| 83 \| 4 \| 1 \| 143 \| 251 \| 397 \| Lysozyme \| Lysozyme; afdb-uniprot50 \| AF-A0A562U8C4-F1-MODEL\_V4 \| 1.0 \| 1.968e-07 \| 332 \| 0.398 \| 138 \| 68 \| 5 \| 20 \| 146 \| 4 \| 137 \| Lysozyme \| Lysozyme; afdb-uniprot50 \| AF-A0A246JYB8-F1-MODEL\_V4 \| 1.0 \| 3.171e-07 \| 332 \| 0.32 \| 150 \| 92 \| 6 \| 3 \| 146 \| 7 \| 152 \| Lysozyme \| Lysozyme; afdb-uniprot50 \| AF-A0A0Q4R256-F1-MODEL\_V4 \| 1.0 \| 8.23e-07 \| 332 \| 0.276 \| 141 \| 95 \| 5 \| 5 \| 144 \| 40 \| 174 \| Lysozyme \| Lysozyme; afdb-uniprot50 \| AF-A0A074TFU6-F1-MODEL\_V4 \| 1.0 \| 5.989e-07 \| 332 \| 0.303 \| 145 \| 93 \| 3 \| 6 \| 145 \| 114 \| 255 \| Lysozyme \| Lysozyme; afdb-uniprot50 \| AF-A0A1F6QQW0-F1-MODEL\_V4 \| 1.0 \| 2.017e-08 \| 331 \| 0.333 \| 144 \| 85 \| 6 \| 10 \| 146 \| 4 \| 143 \| Lysozyme \| Lysozyme; afdb-uniprot50 \| AF-A0A4R5DTI4-F1-MODEL\_V4 \| 1.0 \| 1.432e-07 \| 331 \| 0.28 \| 157 \| 96 \| 6 \| 2 \| 146 \| 13 \| 164 \| Lysozyme \| Lysozyme; afdb-uniprot50 \| AF-A0A2T3HZF6-F1-MODEL\_V4 \| 1.0 \| 2.705e-07 \| 331 \| 0.342 \| 140 \| 80 \| 4 \| 3 \| 135 \| 31 \| 165 \| Lysozyme \| Lysozyme; afdb-uniprot50 \| AF-A0A1L9BSJ1-F1-MODEL\_V4 \| 1.0 \| 1.099e-07 \| 331 \| 0.261 \| 153 \| 102 \| 4 \| 1 \| 145 \| 22 \| 171 \| Lysozyme \| Lysozyme; afdb-uniprot50 \| AF-A0A0R3DQ56-F1-MODEL\_V4 \| 1.0 \| 6.135e-08 \| 331 \| 0.277 \| 162 \| 101 \| 4 \| 1 \| 146 \| 1 \| 162 \| Lysozyme \| Lysozyme; afdb-uniprot50 \| AF-A0A1Y5SFD3-F1-MODEL\_V4 \| 1.0 \| 1.592e-07 \| 331 \| 0.293 \| 150 \| 89 \| 6 \| 6 \| 144 \| 93 \| 236 \| Lysozyme \| Lysozyme; afdb-uniprot50 \| AF-A0A5C7USV4-F1-MODEL\_V4 \| 1.0 \| 8.432e-08 \| 331 \| 0.31 \| 158 \| 93 \| 6 \| 1 \| 145 \| 43 \| 197 \| Lysozyme \| Lysozyme; afdb-uniprot50 \| AF-A0A7X5QS31-F1-MODEL\_V4 \| 1.0 \| 1.592e-07 \| 330 \| 0.251 \| 151 \| 104 \| 3 \| 1 \| 145 \| 4 \| 151 \| Lysozyme \| Lysozyme; afdb-uniprot50 \| AF-A0A2A2S6I3-F1-MODEL\_V4 \| 1.0 \| 1.288e-07 \| 330 \| 0.264 \| 155 \| 103 \| 7 \| 1 \| 145 \| 1 \| 154 \| Lysozyme \| Lysozyme; afdb-uniprot50 \| AF-A0A530R7Q7-F1-MODEL\_V4 \| 1.0 \| 2.705e-07 \| 330 \| 0.285 \| 147 \| 93 \| 6 \| 6 \| 144 \| 48 \| 190 \| Lysozyme \| Lysozyme; afdb-uniprot50 \| AF-E2CJT6-F1-MODEL\_V4 \| 1.0 \| 1.222e-07 \| 330 \| 0.307 \| 153 \| 90 \| 6 \| 5 \| 145 \| 102 \| 250 \| Lysozyme \| Lysozyme; afdb-uniprot50 \| AF-A0A7W7AKQ6-F1-MODEL\_V4 \| 1.0 \| 1.77e-07 \| 329 \| 0.339 \| 153 \| 85 \| 7 \| 3 \| 145 \| 18 \| 164 \| Lysozyme \| Lysozyme; afdb-uniprot50 \| AF-A0A1V5ITR2-F1-MODEL\_V4 \| 1.0 \| 6.658e-07 \| 329 \| 0.427 \| 117 \| 66 \| 1 \| 27 \| 143 \| 2 \| 117 \| Lysozyme \| Lysozyme; afdb-uniprot50 \| AF-A0A1U6JV99-F1-MODEL\_V4 \| 1.0 \| 1.822e-06 \| 329 \| 0.4 \| 100 \| 60 \| 0 \| 44 \| 143 \| 2 \| 101 \| Lysozyme \| Lysozyme; afdb-uniprot50 \| AF-A0A1I7GQC5-F1-MODEL\_V4 \| 1.0 \| 5.68e-07 \| 329 \| 0.333 \| 141 \| 84 \| 5 \| 7 \| 143 \| 55 \| 189 \| Lysozyme \| Lysozyme; afdb-uniprot50 \| AF-A0A1G9VVQ5-F1-MODEL\_V4 \| 1.0 \| 2.188e-07 \| 329 \| 0.263 \| 144 \| 95 \| 4 \| 1 \| 144 \| 5 \| 137 \| Lysozyme \| Lysozyme; afdb-uniprot50 \| AF-A0A529XVD8-F1-MODEL\_V4 \| 1.0 \| 2.188e-07 \| 328 \| 0.241 \| 153 \| 102 \| 5 \| 2 \| 144 \| 3 \| 151 \| Lysozyme \| Lysozyme; afdb-uniprot50 \| AF-A0A7X0DG28-F1-MODEL\_V4 \| 1.0 \| 1.252e-08 \| 328 \| 0.378 \| 156 \| 84 \| 7 \| 13 \| 159 \| 3 \| 154 \| Lysozyme \| Lysozyme; afdb-uniprot50 \| AF-A0A2S9RDE4-F1-MODEL\_V4 \| 1.0 \| 3.007e-07 \| 328 \| 0.347 \| 144 \| 88 \| 3 \| 4 \| 144 \| 12 \| 152 \| Lysozyme \| Lysozyme; afdb-uniprot50 \| AF-A0A6P2W0B7-F1-MODEL\_V4 \| 1.0 \| 9.884e-08 \| 328 \| 0.301 \| 166 \| 87 \| 7 \| 5 \| 144 \| 15 \| 177 \| Lysozyme \| Lysozyme; afdb-uniprot50 \| AF-A0A074TMQ1-F1-MODEL\_V4 \| 1.0 \| 1.867e-07 \| 328 \| 0.277 \| 155 \| 99 \| 5 \| 1 \| 145 \| 41 \| 192 \| Lysozyme \| Lysozyme; afdb-uniprot50 \| AF-E2CNP7-F1-MODEL\_V4 \| 1.0 \| 2.705e-07 \| 328 \| 0.318 \| 160 \| 87 \| 7 \| 1 \| 143 \| 1 \| 155 \| Peptidoglycan binding domain-containing protein \| Peptidoglycan binding domain-containing protein; afdb-uniprot50 \| AF-A0A5M6ZAJ4-F1-MODEL\_V4 \| 1.0 \| 1.51e-07 \| 328 \| 0.294 \| 156 \| 96 \| 5 \| 2 \| 153 \| 6 \| 151 \| Lysozyme \| Lysozyme; afdb-uniprot50 \| AF-A0A708BZD6-F1-MODEL\_V4 \| 1.0 \| 4.986e-06 \| 327 \| 0.632 \| 79 \| 29 \| 0 \| 66 \| 144 \| 2 \| 80 \| Lysozyme \| Lysozyme; afdb-uniprot50 \| AF-A0A5U2FB15-F1-MODEL\_V4 \| 1.0 \| 8.23e-07 \| 327 \| 0.49 \| 102 \| 52 \| 0 \| 43 \| 144 \| 2 \| 103 \| Lysozyme \| Lysozyme; afdb-uniprot50 \| AF-Q5NXI8-F1-MODEL\_V4 \| 1.0 \| 4.358e-07 \| 327 \| 0.251 \| 167 \| 101 \| 4 \| 1 \| 145 \| 11 \| 175 \| Lysozyme \| Lysozyme; afdb-uniprot50 \| AF-A0A1V6EFU8-F1-MODEL\_V4 \| 1.0 \| 2.852e-07 \| 327 \| 0.38 \| 147 \| 77 \| 7 \| 1 \| 143 \| 1 \| 137 \| Lysozyme \| Lysozyme; afdb-uniprot50 \| AF-A0A1H7CRP1-F1-MODEL\_V4 \| 1.0 \| 1.77e-07 \| 327 \| 0.288 \| 156 \| 97 \| 4 \| 1 \| 144 \| 92 \| 245 \| Lysozyme \| Lysozyme; afdb-uniprot50 \| AF-A0A4Q5U1B8-F1-MODEL\_V4 \| 1.0 \| 1.592e-07 \| 327 \| 0.268 \| 160 \| 94 \| 6 \| 1 \| 143 \| 337 \| 490 \| Lysozyme \| Lysozyme; afdb-uniprot50 \| AF-A0A7T8PHM4-F1-MODEL\_V4 \| 1.0 \| 1.51e-07 \| 327 \| 0.322 \| 158 \| 89 \| 4 \| 1 \| 143 \| 832 \| 986 \| Lysozyme \| Lysozyme; afdb-uniprot50 \| AF-A0A0M3SVL5-F1-MODEL\_V4 \| 1.0 \| 6.314e-07 \| 326 \| 0.45 \| 102 \| 56 \| 0 \| 44 \| 145 \| 2 \| 103 \| Lysozyme \| Lysozyme; afdb-uniprot50 \| AF-A0A146H6X4-F1-MODEL\_V4 \| 1.0 \| 2.307e-07 \| 326 \| 0.375 \| 144 \| 79 \| 4 \| 2 \| 138 \| 90 \| 229 \| Lysozyme \| Lysozyme; afdb-uniprot50 \| AF-A0A2E5UZW7-F1-MODEL\_V4 \| 1.0 \| 2.565e-07 \| 326 \| 0.284 \| 144 \| 92 \| 5 \| 1 \| 144 \| 1 \| 133 \| Uncharacterized protein \| Uncharacterized protein; afdb-uniprot50 \| AF-A0A553DTV1-F1-MODEL\_V4 \| 1.0 \| 1.432e-07 \| 326 \| 0.339 \| 159 \| 89 \| 4 \| 1 \| 145 \| 765 \| 921 \| Lysozyme \| Lysozyme; afdb-uniprot50 \| AF-A0A848V5S5-F1-MODEL\_V4 \| 1.0 \| 1.017e-06 \| 325 \| 0.297 \| 148 \| 95 \| 5 \| 1 \| 144 \| 5 \| 147 \| Lysozyme \| Lysozyme; afdb-uniprot50 \| AF-A0A2M7GL79-F1-MODEL\_V4 \| 1.0 \| 4.595e-07 \| 325 \| 0.286 \| 143 \| 91 \| 5 \| 2 \| 144 \| 6 \| 137 \| Lysozyme \| Lysozyme; afdb-uniprot50 \| AF-A0A7L4TJR0-F1-MODEL\_V4 \| 1.0 \| 2.705e-07 \| 325 \| 0.296 \| 155 \| 94 \| 5 \| 2 \| 145 \| 576 \| 726 \| Lysozyme \| Lysozyme; afdb-uniprot50 \| AF-A0A7W5YB65-F1-MODEL\_V4 \| 1.0 \| 2.433e-07 \| 324 \| 0.291 \| 151 \| 92 \| 8 \| 6 \| 144 \| 12 \| 159 \| Lysozyme \| Lysozyme; afdb-uniprot50 \| AF-A0A1N7NPB8-F1-MODEL\_V4 \| 1.0 \| 6.314e-07 \| 323 \| 0.345 \| 133 \| 80 \| 3 \| 18 \| 146 \| 2 \| 131 \| Lysozyme \| Lysozyme; afdb-uniprot50 \| AF-K6ZFL4-F1-MODEL\_V4 \| 1.0 \| 1.193e-06 \| 323 \| 0.423 \| 104 \| 58 \| 2 \| 4 \| 107 \| 30 \| 131 \| Lysozyme \| Lysozyme; afdb-uniprot50 \| AF-A0A846MWR1-F1-MODEL\_V4 \| 1.0 \| 3.343e-07 \| 323 \| 0.286 \| 150 \| 95 \| 6 \| 6 \| 146 \| 13 \| 159 \| Lysozyme \| Lysozyme; afdb-uniprot50 \| AF-A0A811SYU9-F1-MODEL\_V4 \| 1.0 \| 2.565e-07 \| 323 \| 0.271 \| 166 \| 94 \| 7 \| 3 \| 144 \| 14 \| 176 \| Lysozyme RrrD \| Lysozyme RrrD; afdb-uniprot50 \| AF-A0A5X3P0C1-F1-MODEL\_V4 \| 1.0 \| 2.705e-07 \| 323 \| 0.285 \| 154 \| 97 \| 5 \| 1 \| 143 \| 87 \| 238 \| Lysozyme \| Lysozyme; afdb-uniprot50 \| AF-A0A060C188-F1-MODEL\_V4 \| 1.0 \| 2.026e-06 \| 322 \| 0.396 \| 111 \| 67 \| 0 \| 33 \| 143 \| 3 \| 113 \| Lysozyme \| Lysozyme; afdb-uniprot50 \| AF-A0A328A9J3-F1-MODEL\_V4 \| 1.0 \| 2.705e-07 \| 322 \| 0.266 \| 154 \| 98 \| 4 \| 5 \| 156 \| 18 \| 158 \| Lysozyme \| Lysozyme; afdb-uniprot50 \| AF-A0A839ZWC1-F1-MODEL\_V4 \| 1.0 \| 3.343e-07 \| 322 \| 0.277 \| 148 \| 94 \| 6 \| 5 \| 145 \| 20 \| 161 \| Lysozyme \| Lysozyme; afdb-uniprot50 \| AF-A0A1I4AZP2-F1-MODEL\_V4 \| 1.0 \| 9.374e-08 \| 322 \| 0.342 \| 184 \| 78 \| 5 \| 2 \| 143 \| 117 \| 299 \| Lysozyme \| Lysozyme; afdb-uniprot50 \| AF-A0A2D8BR65-F1-MODEL\_V4 \| 1.0 \| 6.314e-07 \| 322 \| 0.281 \| 142 \| 91 \| 5 \| 2 \| 143 \| 21 \| 151 \| Lysozyme \| Lysozyme; afdb-uniprot50 \| AF-A0A2D3TE31-F1-MODEL\_V4 \| 1.0 \| 4.595e-07 \| 321 \| 0.398 \| 108 \| 61 \| 2 \| 1 \| 108 \| 1 \| 104 \| Lysozyme \| Lysozyme; afdb-uniprot50 \| AF-A0A5E8H3J2-F1-MODEL\_V4 \| 1.0 \| 1.358e-07 \| 321 \| 0.32 \| 159 \| 87 \| 9 \| 1 \| 143 \| 31 \| 184 \| Phage-related lysozyme (Muraminidase) \| Phage-related lysozyme (Muraminidase); afdb-uniprot50 \| AF-A0A832GZF3-F1-MODEL\_V4 \| 1.0 \| 5.108e-07 \| 321 \| 0.446 \| 121 \| 63 \| 2 \| 29 \| 146 \| 1 \| 120 \| CHAP domain-containing protein \| CHAP domain-containing protein; afdb-uniprot50 \| AF-A0A0P7VAN4-F1-MODEL\_V4 \| 1.0 \| 3.007e-07 \| 321 \| 0.27 \| 148 \| 99 \| 4 \| 1 \| 144 \| 5 \| 147 \| Lysozyme \| Lysozyme; afdb-uniprot50 \| AF-A0A1M5MU42-F1-MODEL\_V4 \| 1.0 \| 2.852e-07 \| 321 \| 0.303 \| 155 \| 93 \| 5 \| 2 \| 145 \| 727 \| 877 \| Lysozyme \| Lysozyme; afdb-uniprot50 \| AF-A0A5C7QBV1-F1-MODEL\_V4 \| 1.0 \| 7.02e-07 \| 320 \| 0.273 \| 139 \| 92 \| 5 \| 7 \| 144 \| 19 \| 149 \| Lysozyme \| Lysozyme; afdb-uniprot50 \| AF-A0A1G5SFW3-F1-MODEL\_V4 \| 1.0 \| 6.658e-07 \| 320 \| 0.309 \| 155 \| 91 \| 5 \| 1 \| 144 \| 115 \| 264 \| Lysozyme \| Lysozyme; afdb-uniprot50 \| AF-A0A6I5RH79-F1-MODEL\_V4 \| 1.0 \| 3.263e-06 \| 319 \| 0.357 \| 98 \| 63 \| 0 \| 49 \| 146 \| 1 \| 98 \| Lysozyme \| Lysozyme; afdb-uniprot50 \| AF-A0A397TFN8-F1-MODEL\_V4 \| 1.0 \| 6.658e-07 \| 319 \| 0.344 \| 122 \| 74 \| 3 \| 3 \| 121 \| 1 \| 119 \| Lysozyme \| Lysozyme; afdb-uniprot50 \| AF-A0A2I1EL27-F1-MODEL\_V4 \| 1.0 \| 1.432e-07 \| 318 \| 0.387 \| 129 \| 74 \| 2 \| 23 \| 148 \| 10 \| 136 \| Lysozyme \| Lysozyme; afdb-uniprot50 \| AF-A0A3N2S834-F1-MODEL\_V4 \| 1.0 \| 4.595e-07 \| 318 \| 0.286 \| 143 \| 94 \| 4 \| 2 \| 143 \| 6 \| 141 \| Lysozyme \| Lysozyme; afdb-uniprot50 \| AF-A0A516SAT2-F1-MODEL\_V4 \| 1.0 \| 1.679e-07 \| 318 \| 0.26 \| 161 \| 98 \| 6 \| 1 \| 143 \| 8 \| 165 \| Lysozyme \| Lysozyme; afdb-uniprot50 \| AF-A0A1L9CZQ6-F1-MODEL\_V4 \| 1.0 \| 3.717e-07 \| 318 \| 0.32 \| 156 \| 95 \| 7 \| 2 \| 151 \| 4 \| 154 \| PG\_binding\_1 domain-containing protein \| PG\_binding\_1 domain-containing protein; afdb-uniprot50 \| AF-A0A549T0T9-F1-MODEL\_V4 \| 1.0 \| 4.358e-07 \| 318 \| 0.295 \| 159 \| 91 \| 6 \| 2 \| 143 \| 5 \| 159 \| Lysozyme \| Lysozyme; afdb-uniprot50 \| AF-A0A2X3C7T3-F1-MODEL\_V4 \| 1.0 \| 1.131e-06 \| 316 \| 0.388 \| 103 \| 60 \| 2 \| 15 \| 117 \| 33 \| 132 \| Lysozyme \| Lysozyme; afdb-uniprot50 \| AF-A0A8B3NKR9-F1-MODEL\_V4 \| 1.0 \| 4.845e-07 \| 316 \| 0.263 \| 152 \| 93 \| 5 \| 7 \| 144 \| 20 \| 166 \| Lysozyme \| Lysozyme; afdb-uniprot50 \| AF-A0A0K8MB38-F1-MODEL\_V4 \| 1.0 \| 4.595e-07 \| 316 \| 0.335 \| 134 \| 85 \| 3 \| 11 \| 143 \| 2 \| 132 \| Lysozyme \| Lysozyme; afdb-uniprot50 \| AF-A0A0S1B3T1-F1-MODEL\_V4 \| 1.0 \| 3.171e-07 \| 316 \| 0.247 \| 174 \| 110 \| 7 \| 2 \| 157 \| 17 \| 187 \| Lysozyme \| Lysozyme; afdb-uniprot50 \| AF-A0A3S0MS16-F1-MODEL\_V4 \| 1.0 \| 9.648e-07 \| 316 \| 0.234 \| 158 \| 108 \| 6 \| 6 \| 155 \| 17 \| 169 \| Lysozyme \| Lysozyme; afdb-uniprot50 \| AF-A0A378N5X9-F1-MODEL\_V4 \| 1.0 \| 1.326e-06 \| 314 \| 0.474 \| 99 \| 50 \| 2 \| 47 \| 145 \| 1 \| 97 \| Lysozyme \| Lysozyme; afdb-uniprot50 \| AF-A0A2I1F4H6-F1-MODEL\_V4 \| 1.0 \| 7.402e-07 \| 314 \| 0.423 \| 130 \| 64 \| 4 \| 20 \| 144 \| 2 \| 125 \| Lysozyme \| Lysozyme; afdb-uniprot50 \| AF-A0A397V392-F1-MODEL\_V4 \| 1.0 \| 1.257e-06 \| 314 \| 0.428 \| 105 \| 60 \| 0 \| 44 \| 148 \| 38 \| 142 \| Lysozyme \| Lysozyme; afdb-uniprot50 \| AF-A0A511BND2-F1-MODEL\_V4 \| 1.0 \| 2.307e-07 \| 314 \| 0.299 \| 157 \| 93 \| 6 \| 3 \| 145 \| 1 \| 154 \| Uncharacterized protein \| Uncharacterized protein; afdb-uniprot50 \| AF-X5M573-F1-MODEL\_V4 \| 1.0 \| 2.504e-06 \| 314 \| 0.432 \| 104 \| 59 \| 0 \| 41 \| 144 \| 16 \| 119 \| Lysozyme \| Lysozyme; afdb-uniprot50 \| AF-A0A2E6Y944-F1-MODEL\_V4 \| 1.0 \| 5.68e-07 \| 314 \| 0.283 \| 159 \| 93 \| 7 \| 1 \| 143 \| 1 \| 154 \| Lysozyme \| Lysozyme; afdb-uniprot50 \| AF-A0A6L9YUM6-F1-MODEL\_V4 \| 1.0 \| 1.554e-06 \| 314 \| 0.426 \| 129 \| 63 \| 4 \| 1 \| 122 \| 78 \| 202 \| Lysozyme \| Lysozyme; afdb-uniprot50 \| AF-A0A0Q4MLG9-F1-MODEL\_V4 \| 1.0 \| 1.51e-07 \| 314 \| 0.303 \| 178 \| 89 \| 3 \| 1 \| 144 \| 27 \| 203 \| Lysozyme \| Lysozyme; afdb-uniprot50 \| AF-A0A0F9HV13-F1-MODEL\_V4 \| 1.0 \| 2.075e-07 \| 314 \| 0.261 \| 157 \| 100 \| 7 \| 2 \| 146 \| 72 \| 224 \| Lysozyme \| Lysozyme; afdb-uniprot50 \| AF-A0A7Y7A7U2-F1-MODEL\_V4 \| 1.0 \| 5.68e-07 \| 314 \| 0.237 \| 156 \| 106 \| 3 \| 1 \| 144 \| 106 \| 260 \| Lysozyme \| Lysozyme; afdb-uniprot50 \| AF-A0A0A7KTN9-F1-MODEL\_V4 \| 1.0 \| 1.474e-06 \| 314 \| 0.302 \| 129 \| 87 \| 3 \| 1 \| 128 \| 54 \| 180 \| Lysozyme \| Lysozyme; afdb-uniprot50 \| AF-A0A5C7UR14-F1-MODEL\_V4 \| 1.0 \| 1.017e-06 \| 313 \| 0.326 \| 153 \| 90 \| 7 \| 1 \| 146 \| 4 \| 150 \| Lysozyme \| Lysozyme; afdb-uniprot50 \| AF-A0A212KMZ8-F1-MODEL\_V4 \| 1.0 \| 2.075e-07 \| 313 \| 0.268 \| 164 \| 94 \| 8 \| 2 \| 144 \| 3 \| 161 \| Lysozyme \| Lysozyme; afdb-uniprot50 \| AF-A0A495FC89-F1-MODEL\_V4 \| 1.0 \| 4.358e-07 \| 313 \| 0.284 \| 158 \| 94 \| 7 \| 4 \| 143 \| 20 \| 176 \| GH24 family phage-related lysozyme (Muramidase) \| GH24 family phage-related lysozyme (Muramidase); afdb-uniprot50 \| AF-A0A562T9D1-F1-MODEL\_V4 \| 1.0 \| 4.133e-07 \| 313 \| 0.301 \| 159 \| 90 \| 7 \| 1 \| 143 \| 1 \| 154 \| GH24 family phage-related lysozyme (Muramidase) \| GH24 family phage-related lysozyme (Muramidase); afdb-uniprot50 \| AF-A0A843HL85-F1-MODEL\_V4 \| 1.0 \| 3.171e-07 \| 312 \| 0.323 \| 139 \| 87 \| 3 \| 10 \| 144 \| 6 \| 141 \| Lysozyme \| Lysozyme; afdb-uniprot50 \| AF-A0A5J4V626-F1-MODEL\_V4 \| 1.0 \| 9.648e-07 \| 312 \| 0.298 \| 161 \| 92 \| 4 \| 1 \| 144 \| 13 \| 169 \| Lysozyme \| Lysozyme; afdb-uniprot50 \| AF-A0A530KLN7-F1-MODEL\_V4 \| 1.0 \| 4.133e-07 \| 312 \| 0.243 \| 156 \| 101 \| 6 \| 1 \| 144 \| 145 \| 295 \| Lysozyme \| Lysozyme; afdb-uniprot50 \| AF-A0A5Y0KEY1-F1-MODEL\_V4 \| 1.0 \| 5.989e-07 \| 311 \| 0.41 \| 134 \| 69 \| 5 \| 19 \| 147 \| 2 \| 130 \| Lysozyme \| Lysozyme; afdb-uniprot50 \| AF-A0A5C7P136-F1-MODEL\_V4 \| 1.0 \| 4.595e-07 \| 311 \| 0.26 \| 169 \| 103 \| 5 \| 1 \| 148 \| 11 \| 178 \| Lysozyme \| Lysozyme; afdb-uniprot50 \| AF-A0A4Q7FGN8-F1-MODEL\_V4 \| 1.0 \| 8.23e-07 \| 311 \| 0.254 \| 153 \| 100 \| 4 \| 6 \| 144 \| 2 \| 154 \| Lysozyme \| Lysozyme; afdb-uniprot50 \| AF-A6UN41-F1-MODEL\_V4 \| 1.0 \| 9.648e-07 \| 311 \| 0.339 \| 171 \| 82 \| 9 \| 1 \| 144 \| 56 \| 222 \| Lysozyme \| Lysozyme; afdb-uniprot50 \| AF-A0A1H7MLK9-F1-MODEL\_V4 \| 1.0 \| 1.073e-06 \| 310 \| 0.282 \| 131 \| 88 \| 3 \| 15 \| 145 \| 4 \| 128 \| Lysozyme \| Lysozyme; afdb-uniprot50 \| AF-A0A3B0JDC7-F1-MODEL\_V4 \| 1.0 \| 5.989e-07 \| 310 \| 0.32 \| 131 \| 87 \| 1 \| 16 \| 146 \| 3 \| 131 \| Lysozyme \| Lysozyme; afdb-uniprot50 \| AF-A0A524BEN0-F1-MODEL\_V4 \| 1.0 \| 2.252e-06 \| 310 \| 0.391 \| 115 \| 66 \| 3 \| 3 \| 117 \| 67 \| 177 \| Lysozyme \| Lysozyme; afdb-uniprot50 \| AF-A0A2U2P948-F1-MODEL\_V4 \| 1.0 \| 7.02e-07 \| 310 \| 0.329 \| 155 \| 88 \| 4 \| 1 \| 145 \| 83 \| 231 \| Lysozyme \| Lysozyme; afdb-uniprot50 \| AF-A0A1I3VR79-F1-MODEL\_V4 \| 1.0 \| 1.921e-06 \| 310 \| 0.259 \| 154 \| 101 \| 5 \| 1 \| 144 \| 95 \| 245 \| Lysozyme \| Lysozyme; afdb-uniprot50 \| AF-A0A3A6NXV1-F1-MODEL\_V4 \| 1.0 \| 7.02e-07 \| 309 \| 0.254 \| 157 \| 101 \| 6 \| 1 \| 145 \| 9 \| 161 \| Lysozyme \| Lysozyme; afdb-uniprot50 \| AF-A0A3A1Y9X2-F1-MODEL\_V4 \| 1.0 \| 7.805e-07 \| 309 \| 0.272 \| 147 \| 98 \| 5 \| 5 \| 146 \| 35 \| 177 \| Lysozyme \| Lysozyme; afdb-uniprot50 \| AF-A0A3M3YCV8-F1-MODEL\_V4 \| 1.0 \| 8.23e-07 \| 309 \| 0.294 \| 153 \| 89 \| 6 \| 6 \| 144 \| 32 \| 179 \| Lysozyme \| Lysozyme; afdb-uniprot50 \| AF-A0A0G3XM08-F1-MODEL\_V4 \| 1.0 \| 7.02e-07 \| 309 \| 0.282 \| 152 \| 98 \| 6 \| 1 \| 145 \| 28 \| 175 \| Lysozyme \| Lysozyme; afdb-uniprot50 \| AF-A0A4S5JEC4-F1-MODEL\_V4 \| 1.0 \| 3.343e-07 \| 309 \| 0.318 \| 160 \| 88 \| 6 \| 1 \| 143 \| 1 \| 156 \| Lysozyme \| Lysozyme; afdb-uniprot50 \| AF-A0A2I1DVJ3-F1-MODEL\_V4 \| 1.0 \| 1.554e-06 \| 308 \| 0.403 \| 109 \| 59 \| 1 \| 38 \| 146 \| 2 \| 104 \| Lysozyme \| Lysozyme; afdb-uniprot50 \| AF-A0A3B0JB27-F1-MODEL\_V4 \| 1.0 \| 2.504e-06 \| 308 \| 0.375 \| 112 \| 70 \| 0 \| 33 \| 144 \| 3 \| 114 \| Lysozyme \| Lysozyme; afdb-uniprot50 \| AF-A0A2W5R3D9-F1-MODEL\_V4 \| 1.0 \| 1.679e-07 \| 308 \| 0.281 \| 160 \| 94 \| 7 \| 5 \| 153 \| 38 \| 187 \| Lysozyme \| Lysozyme; afdb-uniprot50 \| AF-A3YYR1-F1-MODEL\_V4 \| 1.0 \| 3.92e-07 \| 308 \| 0.292 \| 154 \| 90 \| 5 \| 5 \| 143 \| 504 \| 653 \| Lysozyme \| Lysozyme; afdb-uniprot50 \| AF-A0A0H0XNY4-F1-MODEL\_V4 \| 1.0 \| 7.402e-07 \| 307 \| 0.328 \| 152 \| 91 \| 6 \| 1 \| 145 \| 6 \| 153 \| Lysozyme \| Lysozyme; afdb-uniprot50 \| AF-A0A6B4HYB8-F1-MODEL\_V4 \| 1.0 \| 4.845e-07 \| 307 \| 0.272 \| 154 \| 98 \| 5 \| 3 \| 145 \| 109 \| 259 \| Lysozyme \| Lysozyme; afdb-uniprot50 \| AF-A0A1T2B2A3-F1-MODEL\_V4 \| 1.0 \| 8.23e-07 \| 306 \| 0.259 \| 154 \| 100 \| 5 \| 1 \| 144 \| 115 \| 264 \| Lysozyme \| Lysozyme; afdb-uniprot50 \| AF-A0A2P8FP63-F1-MODEL\_V4 \| 1.0 \| 2.565e-07 \| 305 \| 0.267 \| 157 \| 95 \| 7 \| 2 \| 143 \| 6 \| 157 \| Lysozyme \| Lysozyme; afdb-uniprot50 \| AF-A0A512DJC9-F1-MODEL\_V4 \| 1.0 \| 6.314e-07 \| 305 \| 0.311 \| 186 \| 83 \| 5 \| 1 \| 143 \| 8 \| 191 \| Lysozyme \| Lysozyme; afdb-uniprot50 \| AF-A0A553DTU5-F1-MODEL\_V4 \| 1.0 \| 5.989e-07 \| 305 \| 0.303 \| 158 \| 91 \| 5 \| 1 \| 145 \| 631 \| 782 \| Lysozyme \| Lysozyme; afdb-uniprot50 \| AF-A0A4U0NZJ7-F1-MODEL\_V4 \| 1.0 \| 2.136e-06 \| 304 \| 0.246 \| 142 \| 97 \| 4 \| 5 \| 139 \| 28 \| 166 \| Lysozyme \| Lysozyme; afdb-uniprot50 \| AF-R5ZZA4-F1-MODEL\_V4 \| 1.0 \| 5.108e-07 \| 302 \| 0.324 \| 157 \| 82 \| 6 \| 6 \| 142 \| 12 \| 164 \| Lysozyme \| Lysozyme; afdb-uniprot50 \| AF-A0A3N7ELR7-F1-MODEL\_V4 \| 1.0 \| 2.136e-06 \| 302 \| 0.272 \| 154 \| 99 \| 4 \| 4 \| 147 \| 39 \| 189 \| Lysozyme \| Lysozyme; afdb-uniprot50 \| AF-A0A327JFF9-F1-MODEL\_V4 \| 1.0 \| 1.554e-06 \| 302 \| 0.265 \| 166 \| 101 \| 7 \| 1 \| 150 \| 3 \| 163 \| PG\_binding\_1 domain-containing protein \| PG\_binding\_1 domain-containing protein; afdb-uniprot50 \| AF-A0A2Z3H7L7-F1-MODEL\_V4 \| 1.0 \| 6.658e-07 \| 301 \| 0.304 \| 161 \| 91 \| 7 \| 1 \| 144 \| 5 \| 161 \| Lysozyme \| Lysozyme; afdb-uniprot50 \| AF-A0A346MWA4-F1-MODEL\_V4 \| 1.0 \| 4.358e-07 \| 301 \| 0.266 \| 169 \| 103 \| 8 \| 6 \| 157 \| 26 \| 190 \| Lysozyme \| Lysozyme; afdb-uniprot50 \| AF-A0A1Z8NXD5-F1-MODEL\_V4 \| 1.0 \| 2.198e-05 \| 300 \| 0.444 \| 81 \| 45 \| 0 \| 63 \| 143 \| 2 \| 82 \| Lysozyme \| Lysozyme; afdb-uniprot50 \| AF-A0A7I8I3M8-F1-MODEL\_V4 \| 1.0 \| 1.017e-06 \| 300 \| 0.343 \| 128 \| 77 \| 4 \| 19 \| 144 \| 20 \| 142 \| Lysozyme \| Lysozyme; afdb-uniprot50 \| AF-A0A2V3UDT3-F1-MODEL\_V4 \| 1.0 \| 5.989e-07 \| 300 \| 0.274 \| 171 \| 90 \| 7 \| 3 \| 144 \| 1 \| 166 \| Lysozyme \| Lysozyme; afdb-uniprot50 \| AF-A0A5C7PXY6-F1-MODEL\_V4 \| 1.0 \| 3.095e-06 \| 299 \| 0.263 \| 148 \| 98 \| 4 \| 6 \| 144 \| 34 \| 179 \| Lysozyme \| Lysozyme; afdb-uniprot50 \| AF-A0A6P0ZGF8-F1-MODEL\_V4 \| 1.0 \| 4.986e-06 \| 298 \| 0.434 \| 99 \| 56 \| 0 \| 50 \| 148 \| 7 \| 105 \| Lysozyme \| Lysozyme; afdb-uniprot50 \| AF-A0A2A4V774-F1-MODEL\_V4 \| 1.0 \| 1.193e-06 \| 298 \| 0.298 \| 144 \| 94 \| 2 \| 7 \| 144 \| 85 \| 227 \| Lysozyme \| Lysozyme; afdb-uniprot50 \| AF-E6YK95-F1-MODEL\_V4 \| 1.0 \| 1.047e-05 \| 297 \| 0.43 \| 93 \| 53 \| 0 \| 51 \| 143 \| 40 \| 132 \| Lysozyme \| Lysozyme; afdb-uniprot50 \| AF-A0A059DRI6-F1-MODEL\_V4 \| 1.0 \| 7.805e-07 \| 297 \| 0.242 \| 169 \| 110 \| 7 \| 1 \| 153 \| 1 \| 167 \| Uncharacterized protein \| Uncharacterized protein; afdb-uniprot50 \| AF-K4XZD7-F1-MODEL\_V4 \| 1.0 \| 1.164e-05 \| 296 \| 0.364 \| 96 \| 61 \| 0 \| 49 \| 144 \| 1 \| 96 \| Lysozyme \| Lysozyme; afdb-uniprot50 \| AF-A0A6L3SZ47-F1-MODEL\_V4 \| 1.0 \| 1.257e-06 \| 296 \| 0.294 \| 129 \| 84 \| 4 \| 4 \| 129 \| 16 \| 140 \| Lysozyme \| Lysozyme; afdb-uniprot50 \| AF-A0A829YXQ3-F1-MODEL\_V4 \| 1.0 \| 8.47e-06 \| 296 \| 0.387 \| 98 \| 60 \| 0 \| 47 \| 144 \| 1 \| 98 \| Uncharacterized protein \| Uncharacterized protein; afdb-uniprot50 \| AF-A0A7W9R3L0-F1-MODEL\_V4 \| 1.0 \| 9.648e-07 \| 296 \| 0.296 \| 152 \| 85 \| 7 \| 6 \| 145 \| 17 \| 158 \| Lysozyme \| Lysozyme; afdb-uniprot50 \| AF-A0A6D2B0C1-F1-MODEL\_V4 \| 1.0 \| 1.875e-05 \| 295 \| 0.666 \| 72 \| 24 \| 0 \| 73 \| 144 \| 2 \| 73 \| Lysozyme \| Lysozyme; afdb-uniprot50 \| AF-A0A4R0PG77-F1-MODEL\_V4 \| 1.0 \| 4.986e-06 \| 295 \| 0.425 \| 108 \| 56 \| 1 \| 46 \| 147 \| 2 \| 109 \| Lysozyme \| Lysozyme; afdb-uniprot50 \| AF-A0A0Q8B2U6-F1-MODEL\_V4 \| 1.0 \| 2.935e-06 \| 295 \| 0.248 \| 153 \| 98 \| 7 \| 6 \| 144 \| 43 \| 192 \| Lysozyme \| Lysozyme; afdb-uniprot50 \| AF-A0A7M1L5D9-F1-MODEL\_V4 \| 1.0 \| 2.252e-06 \| 295 \| 0.24 \| 154 \| 104 \| 5 \| 4 \| 144 \| 51 \| 204 \| Lysozyme \| Lysozyme; afdb-uniprot50 \| AF-A0A847GG99-F1-MODEL\_V4 \| 1.0 \| 1.822e-06 \| 294 \| 0.412 \| 114 \| 61 \| 2 \| 41 \| 148 \| 3 \| 116 \| Lysozyme \| Lysozyme; afdb-uniprot50 \| AF-A0A1E4JHV5-F1-MODEL\_V4 \| 1.0 \| 9.417e-06 \| 294 \| 0.257 \| 140 \| 99 \| 4 \| 6 \| 144 \| 18 \| 153 \| Lysozyme \| Lysozyme; afdb-uniprot50 \| AF-A0A1L6JBV4-F1-MODEL\_V4 \| 1.0 \| 2.64e-06 \| 294 \| 0.251 \| 147 \| 93 \| 6 \| 6 \| 143 \| 26 \| 164 \| Lysozyme \| Lysozyme; afdb-uniprot50 \| AF-A0A4R7BD68-F1-MODEL\_V4 \| 1.0 \| 1.474e-06 \| 294 \| 0.27 \| 174 \| 96 \| 5 \| 1 \| 145 \| 53 \| 224 \| Lysozyme \| Lysozyme; afdb-uniprot50 \| AF-A0A0Q4ND89-F1-MODEL\_V4 \| 1.0 \| 1.131e-06 \| 293 \| 0.314 \| 159 \| 87 \| 6 \| 1 \| 145 \| 371 \| 521 \| Lysozyme \| Lysozyme; afdb-uniprot50 \| AF-A0A0E3BDW0-F1-MODEL\_V4 \| 1.0 \| 1.728e-06 \| 292 \| 0.236 \| 144 \| 101 \| 4 \| 10 \| 144 \| 3 \| 146 \| Lysozyme \| Lysozyme; afdb-uniprot50 \| AF-A0A7Y4BU78-F1-MODEL\_V4 \| 1.0 \| 4.729e-06 \| 291 \| 0.383 \| 120 \| 66 \| 2 \| 33 \| 145 \| 1 \| 119 \| Lysozyme \| Lysozyme; afdb-uniprot50 \| AF-A0A1Y3B3E7-F1-MODEL\_V4 \| 1.0 \| 1.365e-05 \| 291 \| 0.478 \| 94 \| 48 \| 1 \| 55 \| 148 \| 160 \| 252 \| Galectin \| Galectin; afdb-uniprot50 \| AF-A0A679IQX3-F1-MODEL\_V4 \| 1.0 \| 5.845e-06 \| 290 \| 0.373 \| 115 \| 68 \| 3 \| 3 \| 117 \| 7 \| 117 \| Lysozyme \| Lysozyme; afdb-uniprot50 \| AF-A0A7Z8G7Z5-F1-MODEL\_V4 \| 1.0 \| 4.485e-06 \| 290 \| 0.366 \| 109 \| 64 \| 3 \| 8 \| 114 \| 18 \| 123 \| Lysozyme \| Lysozyme; afdb-uniprot50 \| AF-A0A447N5D3-F1-MODEL\_V4 \| 1.0 \| 4.729e-06 \| 290 \| 0.328 \| 146 \| 94 \| 3 \| 1 \| 146 \| 27 \| 168 \| Lysozyme \| Lysozyme; afdb-uniprot50 \| AF-A0A1C3FDG7-F1-MODEL\_V4 \| 1.0 \| 1.104e-05 \| 288 \| 0.41 \| 100 \| 54 \| 2 \| 49 \| 144 \| 6 \| 104 \| Lysozyme \| Lysozyme; afdb-uniprot50 \| AF-A0A380WB60-F1-MODEL\_V4 \| 1.0 \| 1.326e-06 \| 288 \| 0.224 \| 174 \| 111 \| 7 \| 1 \| 155 \| 10 \| 178 \| Lysozyme \| Lysozyme; afdb-uniprot50 \| AF-A0A1C7D850-F1-MODEL\_V4 \| 1.0 \| 2.252e-06 \| 287 \| 0.322 \| 161 \| 95 \| 8 \| 1 \| 151 \| 88 \| 244 \| Lysozyme \| Lysozyme; afdb-uniprot50 \| AF-A0A6A8A260-F1-MODEL\_V4 \| 1.0 \| 3.441e-06 \| 286 \| 0.342 \| 108 \| 66 \| 1 \| 37 \| 144 \| 7 \| 109 \| Lysozyme \| Lysozyme; afdb-uniprot50 \| AF-A0A2R5EFM4-F1-MODEL\_V4 \| 1.0 \| 6.163e-06 \| 285 \| 0.202 \| 158 \| 111 \| 4 \| 1 \| 145 \| 11 \| 166 \| Lysozyme \| Lysozyme; afdb-uniprot50 \| AF-A0A440JH49-F1-MODEL\_V4 \| 1.0 \| 3.095e-06 \| 284 \| 0.258 \| 155 \| 98 \| 6 \| 5 \| 146 \| 126 \| 276 \| Lysozyme \| Lysozyme; afdb-uniprot50 \| AF-A0A4R0E7W8-F1-MODEL\_V4 \| 1.0 \| 1.326e-06 \| 284 \| 0.254 \| 165 \| 103 \| 5 \| 1 \| 145 \| 553 \| 717 \| Lysozyme \| Lysozyme; afdb-uniprot50 \| AF-A0A149TPR5-F1-MODEL\_V4 \| 1.0 \| 3.095e-06 \| 283 \| 0.38 \| 113 \| 64 \| 3 \| 37 \| 145 \| 11 \| 121 \| Lysozyme \| Lysozyme; afdb-uniprot50 \| AF-A0A823BTY6-F1-MODEL\_V4 \| 1.0 \| 3.441e-06 \| 283 \| 0.301 \| 156 \| 93 \| 6 \| 4 \| 146 \| 22 \| 174 \| Lysozyme \| Lysozyme; afdb-uniprot50 \| AF-A0A351BF90-F1-MODEL\_V4 \| 1.0 \| 4.253e-06 \| 283 \| 0.307 \| 153 \| 90 \| 5 \| 1 \| 143 \| 190 \| 336 \| Lysozyme \| Lysozyme; afdb-uniprot50 \| AF-A0A4Q7BNM2-F1-MODEL\_V4 \| 1.0 \| 1.073e-06 \| 283 \| 0.278 \| 165 \| 95 \| 6 \| 1 \| 145 \| 467 \| 627 \| Lysozyme \| Lysozyme; afdb-uniprot50 \| AF-A0A3S4GC52-F1-MODEL\_V4 \| 1.0 \| 1.921e-06 \| 282 \| 0.308 \| 133 \| 79 \| 3 \| 7 \| 129 \| 19 \| 148 \| Lysozyme \| Lysozyme; afdb-uniprot50 \| AF-A0A142JHX9-F1-MODEL\_V4 \| 1.0 \| 4.034e-06 \| 282 \| 0.308 \| 149 \| 81 \| 6 \| 8 \| 143 \| 19 \| 158 \| Lysozyme \| Lysozyme; afdb-uniprot50 \| AF-A0A7W4RHC5-F1-MODEL\_V4 \| 1.0 \| 3.628e-06 \| 282 \| 0.212 \| 165 \| 107 \| 7 \| 1 \| 143 \| 13 \| 176 \| GH24 family phage-related lysozyme (Muramidase) \| GH24 family phage-related lysozyme (Muramidase); afdb-uniprot50 \| AF-A0A246GF81-F1-MODEL\_V4 \| 1.0 \| 2.252e-06 \| 282 \| 0.316 \| 158 \| 89 \| 5 \| 1 \| 145 \| 137 \| 288 \| Lysozyme \| Lysozyme; afdb-uniprot50 \| AF-A0A704CZH1-F1-MODEL\_V4 \| 1.0 \| 6.163e-06 \| 281 \| 0.533 \| 103 \| 44 \| 1 \| 28 \| 126 \| 1 \| 103 \| Lysozyme \| Lysozyme; afdb-uniprot50 \| AF-A0A1F3IZE5-F1-MODEL\_V4 \| 1.0 \| 1.474e-06 \| 281 \| 0.313 \| 150 \| 88 \| 5 \| 1 \| 143 \| 328 \| 469 \| Lysozyme \| Lysozyme; afdb-uniprot50 \| AF-A0A3N7ETY2-F1-MODEL\_V4 \| 1.0 \| 4.034e-06 \| 280 \| 0.432 \| 118 \| 62 \| 1 \| 33 \| 145 \| 2 \| 119 \| Lysozyme \| Lysozyme; afdb-uniprot50 \| AF-A0A7R9MBY2-F1-MODEL\_V4 \| 1.0 \| 9.648e-07 \| 280 \| 0.256 \| 156 \| 96 \| 5 \| 7 \| 144 \| 9 \| 162 \| Lysozyme \| Lysozyme; afdb-uniprot50 \| AF-A0A3T4D307-F1-MODEL\_V4 \| 1.0 \| 2.375e-06 \| 280 \| 0.294 \| 170 \| 91 \| 7 \| 3 \| 146 \| 9 \| 175 \| Lysozyme \| Lysozyme; afdb-uniprot50 \| AF-A0A5R8ZQ96-F1-MODEL\_V4 \| 1.0 \| 2.375e-06 \| 279 \| 0.235 \| 153 \| 98 \| 6 \| 6 \| 144 \| 26 \| 173 \| Lysozyme \| Lysozyme; afdb-uniprot50 \| AF-A0A1G4RXF8-F1-MODEL\_V4 \| 1.0 \| 4.729e-06 \| 279 \| 0.217 \| 156 \| 110 \| 5 \| 2 \| 151 \| 26 \| 175 \| Lysozyme \| Lysozyme; afdb-uniprot50 \| AF-A0A3T9QUX1-F1-MODEL\_V4 \| 1.0 \| 4.152e-05 \| 278 \| 0.363 \| 88 \| 56 \| 0 \| 56 \| 143 \| 2 \| 89 \| Lysozyme \| Lysozyme; afdb-uniprot50 \| AF-A0A1X7MCZ0-F1-MODEL\_V4 \| 1.0 \| 1.227e-05 \| 278 \| 0.383 \| 107 \| 65 \| 1 \| 38 \| 144 \| 1 \| 106 \| Lysozyme \| Lysozyme; afdb-uniprot50 \| AF-A0A060C0Y3-F1-MODEL\_V4 \| 1.0 \| 4.034e-06 \| 278 \| 0.377 \| 127 \| 71 \| 5 \| 22 \| 144 \| 1 \| 123 \| Lysozyme \| Lysozyme; afdb-uniprot50 \| AF-A3VBD6-F1-MODEL\_V4 \| 1.0 \| 1.822e-06 \| 278 \| 0.291 \| 161 \| 94 \| 6 \| 7 \| 150 \| 122 \| 279 \| Lysozyme \| Lysozyme; afdb-uniprot50 \| AF-A0A550KMS5-F1-MODEL\_V4 \| 1.0 \| 2.375e-06 \| 278 \| 0.267 \| 157 \| 97 \| 5 \| 5 \| 157 \| 9 \| 151 \| Lysozyme \| Lysozyme; afdb-uniprot50 \| AF-A0A433PHE8-F1-MODEL\_V4 \| 1.0 \| 4.485e-06 \| 278 \| 0.203 \| 157 \| 108 \| 7 \| 1 \| 147 \| 348 \| 497 \| Lysozyme \| Lysozyme; afdb-uniprot50 \| AF-D8MV64-F1-MODEL\_V4 \| 1.0 \| 1.875e-05 \| 277 \| 0.434 \| 99 \| 53 \| 2 \| 47 \| 144 \| 1 \| 97 \| Lysozyme \| Lysozyme; afdb-uniprot50 \| AF-A0A147GWA0-F1-MODEL\_V4 \| 1.0 \| 2.504e-06 \| 277 \| 0.246 \| 166 \| 99 \| 8 \| 4 \| 144 \| 23 \| 187 \| Uncharacterized protein \| Uncharacterized protein; afdb-uniprot50 \| AF-A0A7C2WUQ3-F1-MODEL\_V4 \| 1.0 \| 7.618e-06 \| 277 \| 0.233 \| 150 \| 102 \| 3 \| 6 \| 143 \| 45 \| 193 \| Lysozyme \| Lysozyme; afdb-uniprot50 \| AF-A0A7R8N0H2-F1-MODEL\_V4 \| 1.0 \| 2.865e-05 \| 276 \| 0.371 \| 97 \| 61 \| 0 \| 46 \| 142 \| 3 \| 99 \| Lysozyme \| Lysozyme; afdb-uniprot50 \| AF-A0A4V5MMW7-F1-MODEL\_V4 \| 1.0 \| 5.544e-06 \| 276 \| 0.234 \| 149 \| 99 \| 5 \| 3 \| 139 \| 87 \| 232 \| Lysozyme \| Lysozyme; afdb-uniprot50 \| AF-A0A448SSY8-F1-MODEL\_V4 \| 1.0 \| 1.164e-05 \| 275 \| 0.339 \| 106 \| 67 \| 1 \| 38 \| 143 \| 11 \| 113 \| Lysozyme \| Lysozyme; afdb-uniprot50 \| AF-A0A285CW41-F1-MODEL\_V4 \| 1.0 \| 1.517e-05 \| 275 \| 0.342 \| 108 \| 70 \| 1 \| 40 \| 146 \| 14 \| 121 \| Lysozyme \| Lysozyme; afdb-uniprot50 \| AF-A0A376RU32-F1-MODEL\_V4 \| 1.0 \| 1.294e-05 \| 275 \| 0.367 \| 117 \| 70 \| 3 \| 1 \| 117 \| 27 \| 139 \| Lysozyme \| Lysozyme; afdb-uniprot50 \| AF-J1J5J1-F1-MODEL\_V4 \| 1.0 \| 2.444e-05 \| 275 \| 0.406 \| 96 \| 56 \| 1 \| 51 \| 145 \| 6 \| 101 \| Lysozyme \| Lysozyme; afdb-uniprot50 \| AF-A0A431HNJ6-F1-MODEL\_V4 \| 1.0 \| 8.033e-06 \| 275 \| 0.42 \| 100 \| 52 \| 2 \| 53 \| 146 \| 2 \| 101 \| Lysozyme \| Lysozyme; afdb-uniprot50 \| AF-A0A377PMX4-F1-MODEL\_V4 \| 1.0 \| 2.318e-05 \| 274 \| 0.34 \| 100 \| 66 \| 0 \| 44 \| 143 \| 4 \| 103 \| Lysozyme \| Lysozyme; afdb-uniprot50 \| AF-A0A147IZ09-F1-MODEL\_V4 \| 1.0 \| 4.034e-06 \| 274 \| 0.284 \| 158 \| 89 \| 5 \| 6 \| 143 \| 23 \| 176 \| Lysozyme \| Lysozyme; afdb-uniprot50 \| AF-A0A2G1YJ53-F1-MODEL\_V4 \| 1.0 \| 9.929e-06 \| 274 \| 0.245 \| 151 \| 102 \| 7 \| 2 \| 146 \| 49 \| 193 \| Lysozyme \| Lysozyme; afdb-uniprot50 \| AF-A0A7R9KV78-F1-MODEL\_V4 \| 1.0 \| 3.734e-05 \| 274 \| 0.432 \| 97 \| 54 \| 1 \| 49 \| 145 \| 2 \| 97 \| Lysozyme \| Lysozyme; afdb-uniprot50 \| AF-A0A743U3J8-F1-MODEL\_V4 \| 1.0 \| 1.778e-05 \| 273 \| 0.319 \| 122 \| 79 \| 3 \| 4 \| 125 \| 30 \| 147 \| Lysozyme \| Lysozyme; afdb-uniprot50 \| AF-A0A7W0BQY6-F1-MODEL\_V4 \| 1.0 \| 2.504e-06 \| 273 \| 0.204 \| 171 \| 113 \| 8 \| 1 \| 159 \| 9 \| 168 \| Lysozyme \| Lysozyme; afdb-uniprot50 \| AF-A0A397VV53-F1-MODEL\_V4 \| 1.0 \| 8.47e-06 \| 273 \| 0.317 \| 123 \| 72 \| 4 \| 2 \| 117 \| 71 \| 188 \| Lysozyme \| Lysozyme; afdb-uniprot50 \| AF-A0A062IRX0-F1-MODEL\_V4 \| 1.0 \| 6.163e-06 \| 272 \| 0.183 \| 158 \| 114 \| 7 \| 1 \| 145 \| 12 \| 167 \| Phage lysozyme family protein \| Phage lysozyme family protein; afdb-uniprot50 \| AF-A3Z115-F1-MODEL\_V4 \| 1.0 \| 1.047e-05 \| 272 \| 0.44 \| 93 \| 51 \| 1 \| 53 \| 144 \| 3 \| 95 \| Lysozyme \| Lysozyme; afdb-uniprot50 \| AF-U9UUS9-F1-MODEL\_V4 \| 1.0 \| 1.687e-05 \| 271 \| 0.333 \| 111 \| 67 \| 2 \| 1 \| 108 \| 5 \| 111 \| Lysozyme \| Lysozyme; afdb-uniprot50 \| AF-A0A827P9W5-F1-MODEL\_V4 \| 1.0 \| 8.931e-06 \| 271 \| 0.298 \| 114 \| 72 \| 3 \| 6 \| 114 \| 21 \| 131 \| Lysozyme \| Lysozyme; afdb-uniprot50 \| AF-A0A7T5K5I0-F1-MODEL\_V4 \| 1.0 \| 2.717e-05 \| 271 \| 0.327 \| 110 \| 71 \| 2 \| 7 \| 116 \| 19 \| 125 \| Lysozyme \| Lysozyme; afdb-uniprot50 \| AF-A0A0F9SL40-F1-MODEL\_V4 \| 1.0 \| 1.164e-05 \| 271 \| 0.209 \| 167 \| 108 \| 5 \| 2 \| 144 \| 114 \| 280 \| Uncharacterized protein \| Uncharacterized protein; afdb-uniprot50 \| AF-A0A1Y1XFR8-F1-MODEL\_V4 \| 1.0 \| 1.921e-06 \| 270 \| 0.284 \| 158 \| 92 \| 8 \| 4 \| 152 \| 1 \| 146 \| Lysozyme \| Lysozyme; afdb-uniprot50 \| AF-A0A809ZZ30-F1-MODEL\_V4 \| 1.0 \| 1.294e-05 \| 270 \| 0.243 \| 152 \| 105 \| 4 \| 2 \| 143 \| 14 \| 165 \| Uncharacterized protein \| Uncharacterized protein; afdb-uniprot50 \| AF-A0A4R0DJ46-F1-MODEL\_V4 \| 1.0 \| 1.047e-05 \| 269 \| 0.235 \| 157 \| 104 \| 7 \| 2 \| 145 \| 22 \| 175 \| Lysozyme \| Lysozyme; afdb-uniprot50 \| AF-A0A4Q5R9M0-F1-MODEL\_V4 \| 1.0 \| 3.441e-06 \| 269 \| 0.263 \| 190 \| 95 \| 5 \| 3 \| 149 \| 106 \| 293 \| Lysozyme \| Lysozyme; afdb-uniprot50 \| AF-A0A4Y6U9J2-F1-MODEL\_V4 \| 1.0 \| 1.554e-06 \| 268 \| 0.272 \| 165 \| 95 \| 6 \| 6 \| 148 \| 15 \| 176 \| Lysozyme \| Lysozyme; afdb-uniprot50 \| AF-A0A5C8RYA1-F1-MODEL\_V4 \| 1.0 \| 4.867e-05 \| 267 \| 0.37 \| 100 \| 62 \| 1 \| 46 \| 144 \| 2 \| 101 \| Lysozyme \| Lysozyme; afdb-uniprot50 \| AF-A0A0P6WR43-F1-MODEL\_V4 \| 1.0 \| 3.441e-06 \| 266 \| 0.273 \| 172 \| 88 \| 6 \| 6 \| 143 \| 25 \| 193 \| Lysozyme \| Lysozyme; afdb-uniprot50 \| AF-C1D947-F1-MODEL\_V4 \| 1.0 \| 1.517e-05 \| 265 \| 0.411 \| 102 \| 59 \| 1 \| 44 \| 145 \| 1 \| 101 \| Lysozyme \| Lysozyme; afdb-uniprot50 \| AF-A0A0H5Q1T3-F1-MODEL\_V4 \| 1.0 \| 3.021e-05 \| 265 \| 0.231 \| 147 \| 105 \| 4 \| 1 \| 145 \| 17 \| 157 \| Uncharacterized protein \| Uncharacterized protein; afdb-uniprot50 \| AF-A0A431IGY8-F1-MODEL\_V4 \| 1.0 \| 4.034e-06 \| 265 \| 0.281 \| 160 \| 98 \| 6 \| 5 \| 150 \| 6 \| 162 \| Lysozyme \| Lysozyme; afdb-uniprot50 \| AF-A0A4R3UTK5-F1-MODEL\_V4 \| 1.0 \| 0.0001136 \| 264 \| 0.413 \| 87 \| 51 \| 0 \| 60 \| 146 \| 4 \| 90 \| Lysozyme \| Lysozyme; afdb-uniprot50 \| AF-A0A4R0D3N4-F1-MODEL\_V4 \| 1.0 \| 1.517e-05 \| 264 \| 0.227 \| 158 \| 106 \| 7 \| 1 \| 145 \| 23 \| 177 \| Lysozyme \| Lysozyme; afdb-uniprot50 \| AF-A0A7G7M048-F1-MODEL\_V4 \| 1.0 \| 5.258e-06 \| 264 \| 0.243 \| 152 \| 98 \| 7 \| 16 \| 158 \| 65 \| 208 \| Lysozyme \| Lysozyme; afdb-uniprot50 \| AF-A0A077FK68-F1-MODEL\_V4 \| 1.0 \| 0.0001481 \| 262 \| 0.379 \| 79 \| 49 \| 0 \| 65 \| 143 \| 2 \| 80 \| Lysozyme \| Lysozyme; afdb-uniprot50 \| AF-A0A0Q4I8D7-F1-MODEL\_V4 \| 1.0 \| 9.417e-06 \| 262 \| 0.351 \| 131 \| 76 \| 5 \| 18 \| 145 \| 9 \| 133 \| Lysozyme \| Lysozyme; afdb-uniprot50 \| AF-V2U938-F1-MODEL\_V4 \| 1.0 \| 1.365e-05 \| 262 \| 0.197 \| 157 \| 111 \| 6 \| 2 \| 145 \| 13 \| 167 \| Lysozyme \| Lysozyme; afdb-uniprot50 \| AF-A0A845M958-F1-MODEL\_V4 \| 1.0 \| 1.365e-05 \| 262 \| 0.291 \| 161 \| 97 \| 6 \| 1 \| 146 \| 116 \| 274 \| Glycoside hydrolase family protein \| Glycoside hydrolase family protein; afdb-uniprot50 \| AF-A0A0B5DZ45-F1-MODEL\_V4 \| 1.0 \| 8.931e-06 \| 261 \| 0.192 \| 151 \| 109 \| 5 \| 2 \| 144 \| 84 \| 229 \| Lysozyme \| Lysozyme; afdb-uniprot50 \| AF-B5ZDC6-F1-MODEL\_V4 \| 1.0 \| 0.0001198 \| 260 \| 0.376 \| 85 \| 52 \| 1 \| 61 \| 144 \| 7 \| 91 \| Lysozyme \| Lysozyme; afdb-uniprot50 \| AF-A0A561HF46-F1-MODEL\_V4 \| 1.0 \| 4.986e-06 \| 260 \| 0.4 \| 115 \| 64 \| 2 \| 37 \| 146 \| 2 \| 116 \| Lysozyme \| Lysozyme; afdb-uniprot50 \| AF-A0A561HYA6-F1-MODEL\_V4 \| 1.0 \| 1.164e-05 \| 259 \| 0.405 \| 111 \| 63 \| 2 \| 37 \| 146 \| 6 \| 114 \| Lysozyme \| Lysozyme; afdb-uniprot50 \| AF-A0A7C9P2F6-F1-MODEL\_V4 \| 1.0 \| 2.444e-05 \| 259 \| 0.282 \| 124 \| 83 \| 3 \| 24 \| 143 \| 2 \| 123 \| Lysozyme \| Lysozyme; afdb-uniprot50 \| AF-A0A7Z1UH38-F1-MODEL\_V4 \| 1.0 \| 2.444e-05 \| 257 \| 0.282 \| 152 \| 90 \| 7 \| 7 \| 144 \| 104 \| 250 \| Lysozyme \| Lysozyme; afdb-uniprot50 \| AF-A0A2P4P1I1-F1-MODEL\_V4 \| 1.0 \| 3.734e-05 \| 256 \| 0.333 \| 108 \| 65 \| 2 \| 4 \| 108 \| 1 \| 104 \| Lysozyme \| Lysozyme; afdb-uniprot50 \| AF-A0A349FCC2-F1-MODEL\_V4 \| 1.0 \| 5.705e-05 \| 256 \| 0.398 \| 103 \| 55 \| 1 \| 50 \| 145 \| 1 \| 103 \| Lysozyme \| Lysozyme; afdb-uniprot50 \| AF-A0A0Q7DQU4-F1-MODEL\_V4 \| 1.0 \| 7.618e-06 \| 256 \| 0.317 \| 151 \| 86 \| 6 \| 4 \| 146 \| 48 \| 189 \| Lysozyme \| Lysozyme; afdb-uniprot50 \| AF-A0A2G6GNA8-F1-MODEL\_V4 \| 1.0 \| 6.499e-06 \| 256 \| 0.278 \| 147 \| 93 \| 8 \| 7 \| 146 \| 55 \| 195 \| Uncharacterized protein \| Uncharacterized protein; afdb-uniprot50 \| AF-A0A4U2U973-F1-MODEL\_V4 \| 1.0 \| 0.0001404 \| 254 \| 0.306 \| 98 \| 68 \| 0 \| 49 \| 146 \| 1 \| 98 \| Lysozyme \| Lysozyme; afdb-uniprot50 \| AF-Q2N9G4-F1-MODEL\_V4 \| 1.0 \| 2.444e-05 \| 254 \| 0.276 \| 152 \| 99 \| 6 \| 1 \| 145 \| 35 \| 182 \| Lysozyme \| Lysozyme; afdb-uniprot50 \| AF-A0A226EDT1-F1-MODEL\_V4 \| 1.0 \| 4.152e-05 \| 254 \| 0.396 \| 106 \| 59 \| 3 \| 46 \| 148 \| 26 \| 129 \| Lysozyme \| Lysozyme; afdb-uniprot50 \| AF-A0A484QZB0-F1-MODEL\_V4 \| 1.0 \| 6.688e-05 \| 254 \| 0.416 \| 96 \| 56 \| 0 \| 49 \| 144 \| 135 \| 230 \| Lysozyme \| Lysozyme; afdb-uniprot50 \| AF-A0A4V6KVI0-F1-MODEL\_V4 \| 1.0 \| 5.132e-05 \| 253 \| 0.397 \| 98 \| 57 \| 1 \| 53 \| 150 \| 2 \| 97 \| Lysozyme \| Lysozyme; afdb-uniprot50 \| AF-A0A0G3S5Y9-F1-MODEL\_V4 \| 1.0 \| 1.439e-05 \| 253 \| 0.256 \| 152 \| 97 \| 5 \| 1 \| 147 \| 1 \| 141 \| Lysozyme \| Lysozyme; afdb-uniprot50 \| AF-A0A7V7Z6A1-F1-MODEL\_V4 \| 1.0 \| 6.163e-06 \| 253 \| 0.402 \| 139 \| 64 \| 3 \| 24 \| 147 \| 2 \| 136 \| Lysozyme \| Lysozyme; afdb-uniprot50 \| AF-A0A5W2M167-F1-MODEL\_V4 \| 1.0 \| 0.0001022 \| 252 \| 0.391 \| 97 \| 58 \| 1 \| 49 \| 145 \| 1 \| 96 \| Lysozyme \| Lysozyme; afdb-uniprot50 \| AF-A0A317PF33-F1-MODEL\_V4 \| 1.0 \| 2.198e-05 \| 252 \| 0.232 \| 159 \| 101 \| 7 \| 1 \| 143 \| 1 \| 154 \| GH24 family phage-related lysozyme (Muramidase) \| GH24 family phage-related lysozyme (Muramidase); afdb-uniprot50 \| AF-A0A0Q8U0X3-F1-MODEL\_V4 \| 1.0 \| 3.359e-05 \| 251 \| 0.216 \| 166 \| 104 \| 6 \| 4 \| 144 \| 19 \| 183 \| Lysozyme \| Lysozyme; afdb-uniprot50 \| AF-A0A1M6SYR1-F1-MODEL\_V4 \| 1.0 \| 2.865e-05 \| 251 \| 0.304 \| 161 \| 87 \| 8 \| 3 \| 146 \| 41 \| 193 \| Lysozyme \| Lysozyme; afdb-uniprot50 \| AF-A0A0R3MW87-F1-MODEL\_V4 \| 1.0 \| 3.826e-06 \| 251 \| 0.244 \| 184 \| 97 \| 8 \| 1 \| 159 \| 9 \| 175 \| Uncharacterized protein \| Uncharacterized protein; afdb-uniprot50 \| AF-A0A1Z8LH42-F1-MODEL\_V4 \| 1.0 \| 0.0001646 \| 250 \| 0.333 \| 96 \| 62 \| 1 \| 50 \| 143 \| 11 \| 106 \| Lysozyme \| Lysozyme; afdb-uniprot50 \| AF-A0A841QL93-F1-MODEL\_V4 \| 1.0 \| 0.0003843 \| 249 \| 0.437 \| 80 \| 45 \| 0 \| 66 \| 145 \| 2 \| 81 \| GH24 family phage-related lysozyme (Muramidase) \| GH24 family phage-related lysozyme (Muramidase); afdb-uniprot50 \| AF-A0A4V3UX11-F1-MODEL\_V4 \| 1.0 \| 0.0001078 \| 249 \| 0.336 \| 104 \| 69 \| 0 \| 41 \| 144 \| 3 \| 106 \| Lysozyme \| Lysozyme; afdb-uniprot50 \| AF-A0A430D648-F1-MODEL\_V4 \| 1.0 \| 1.227e-05 \| 247 \| 0.231 \| 164 \| 97 \| 5 \| 6 \| 143 \| 21 \| 181 \| Lysozyme \| Lysozyme; afdb-uniprot50 \| AF-A0A3M9Z534-F1-MODEL\_V4 \| 1.0 \| 1.921e-06 \| 247 \| 0.335 \| 137 \| 72 \| 4 \| 24 \| 146 \| 130 \| 261 \| Lysozyme \| Lysozyme; afdb-uniprot50 \| AF-A0A4V1RG86-F1-MODEL\_V4 \| 1.0 \| 0.0001646 \| 246 \| 0.333 \| 96 \| 64 \| 0 \| 49 \| 144 \| 1 \| 96 \| Lysozyme \| Lysozyme; afdb-uniprot50 \| AF-A0A1N6UYK7-F1-MODEL\_V4 \| 1.0 \| 1.517e-05 \| 246 \| 0.22 \| 172 \| 109 \| 9 \| 4 \| 156 \| 12 \| 177 \| Lysozyme \| Lysozyme; afdb-uniprot50 \| AF-A0A5E8P0K9-F1-MODEL\_V4 \| 1.0 \| 3.542e-05 \| 245 \| 0.336 \| 104 \| 61 \| 1 \| 41 \| 144 \| 52 \| 147 \| Lysozyme \| Lysozyme; afdb-uniprot50 \| AF-A0A1B3PFJ1-F1-MODEL\_V4 \| 1.0 \| 6.343e-05 \| 245 \| 0.186 \| 166 \| 110 \| 6 \| 2 \| 143 \| 22 \| 186 \| Putative phage lysozyme \| Putative phage lysozyme; afdb-uniprot50 \| AF-A0A3M9Z3J4-F1-MODEL\_V4 \| 1.0 \| 1.778e-05 \| 243 \| 0.396 \| 111 \| 58 \| 2 \| 38 \| 144 \| 22 \| 127 \| Lysozyme \| Lysozyme; afdb-uniprot50 \| AF-A0A1A9WSJ5-F1-MODEL\_V4 \| 1.0 \| 7.052e-05 \| 243 \| 0.397 \| 93 \| 56 \| 0 \| 40 \| 132 \| 25 \| 117 \| Lysozyme \| Lysozyme; afdb-uniprot50 \| AF-A0A1G8LAP6-F1-MODEL\_V4 \| 1.0 \| 0.000183 \| 241 \| 0.444 \| 90 \| 49 \| 1 \| 57 \| 146 \| 2 \| 90 \| Lysozyme \| Lysozyme; afdb-uniprot50 \| AF-C6XGM8-F1-MODEL\_V4 \| 1.0 \| 0.0001332 \| 239 \| 0.333 \| 102 \| 66 \| 1 \| 47 \| 146 \| 1 \| 102 \| Lysozyme \| Lysozyme; afdb-uniprot50 \| AF-A0A143HSA8-F1-MODEL\_V4 \| 1.0 \| 0.0001481 \| 236 \| 0.327 \| 113 \| 74 \| 2 \| 33 \| 144 \| 3 \| 114 \| Lysozyme \| Lysozyme; afdb-uniprot50 \| AF-A0A6B8LZY5-F1-MODEL\_V4 \| 1.0 \| 2.577e-05 \| 235 \| 0.427 \| 117 \| 58 \| 3 \| 32 \| 144 \| 2 \| 113 \| Lysozyme \| Lysozyme; afdb-uniprot50 \| AF-A0A126V243-F1-MODEL\_V4 \| 1.0 \| 0.0004052 \| 234 \| 0.349 \| 83 \| 52 \| 1 \| 64 \| 144 \| 2 \| 84 \| Lysozyme \| Lysozyme; afdb-uniprot50 \| AF-G5LKD0-F1-MODEL\_V4 \| 1.0 \| 0.0001481 \| 233 \| 0.368 \| 103 \| 63 \| 1 \| 46 \| 146 \| 4 \| 106 \| Lysozyme \| Lysozyme; afdb-uniprot50 \| AF-A0A356CHN3-F1-MODEL\_V4 \| 1.0 \| 0.0003457 \| 232 \| 0.432 \| 81 \| 46 \| 0 \| 67 \| 147 \| 4 \| 84 \| Lysozyme \| Lysozyme; afdb-uniprot50 \| AF-A0A4Q3MM19-F1-MODEL\_V4 \| 1.0 \| 6.016e-05 \| 232 \| 0.237 \| 135 \| 91 \| 2 \| 22 \| 144 \| 28 \| 162 \| Lysozyme \| Lysozyme; afdb-uniprot50 \| AF-A0A839KL75-F1-MODEL\_V4 \| 1.0 \| 0.000807 \| 231 \| 0.416 \| 72 \| 42 \| 0 \| 73 \| 144 \| 2 \| 73 \| Lysozyme RrrD \| Lysozyme RrrD; afdb-uniprot50 \| AF-E4UDK1-F1-MODEL\_V4 \| 1.0 \| 0.0002146 \| 231 \| 0.298 \| 104 \| 67 \| 2 \| 47 \| 146 \| 1 \| 102 \| Lysozyme \| Lysozyme; afdb-uniprot50 \| AF-A0A2D0ANW9-F1-MODEL\_V4 \| 1.0 \| 7.052e-05 \| 231 \| 0.226 \| 159 \| 101 \| 5 \| 6 \| 145 \| 22 \| 177 \| Lysozyme \| Lysozyme; afdb-uniprot50 \| AF-A0A418WYF3-F1-MODEL\_V4 \| 1.0 \| 0.0001136 \| 229 \| 0.32 \| 106 \| 69 \| 2 \| 38 \| 142 \| 1 \| 104 \| Lysozyme \| Lysozyme; afdb-uniprot50 \| AF-A0A376PX23-F1-MODEL\_V4 \| 1.0 \| 0.0001022 \| 229 \| 0.289 \| 107 \| 68 \| 3 \| 6 \| 107 \| 21 \| 124 \| Lysozyme \| Lysozyme; afdb-uniprot50 \| AF-A4WPP1-F1-MODEL\_V4 \| 1.0 \| 5.411e-05 \| 228 \| 0.21 \| 152 \| 101 \| 6 \| 5 \| 144 \| 9 \| 153 \| Lysozyme \| Lysozyme; afdb-uniprot50 \| AF-A0A6L3SU82-F1-MODEL\_V4 \| 1.0 \| 0.0002035 \| 228 \| 0.366 \| 101 \| 59 \| 2 \| 46 \| 146 \| 4 \| 99 \| Lysozyme \| Lysozyme; afdb-uniprot50 \| AF-F4VC63-F1-MODEL\_V4 \| 1.0 \| 3.937e-05 \| 226 \| 0.264 \| 121 \| 76 \| 4 \| 6 \| 116 \| 21 \| 138 \| Lysozyme \| Lysozyme; afdb-uniprot50 \| AF-R9AZY7-F1-MODEL\_V4 \| 1.0 \| 0.0004505 \| 225 \| 0.558 \| 77 \| 34 \| 0 \| 62 \| 138 \| 4 \| 80 \| Lysozyme \| Lysozyme; afdb-uniprot50 \| AF-A0A8B2R8P4-F1-MODEL\_V4 \| 1.0 \| 0.000807 \| 224 \| 0.348 \| 89 \| 58 \| 0 \| 56 \| 144 \| 3 \| 91 \| Lysozyme \| Lysozyme; afdb-uniprot50 \| AF-A0A1X7MEG5-F1-MODEL\_V4 \| 1.0 \| 0.001524 \| 220 \| 0.452 \| 73 \| 40 \| 0 \| 71 \| 143 \| 3 \| 75 \| Lysozyme \| Lysozyme; afdb-uniprot50 \| AF-A0A5X2XK25-F1-MODEL\_V4 \| 1.0 \| 7.436e-05 \| 219 \| 0.359 \| 114 \| 67 \| 3 \| 37 \| 147 \| 3 \| 113 \| Lysozyme \| Lysozyme; afdb-uniprot50 \| AF-A0A774SGM1-F1-MODEL\_V4 \| 1.0 \| 0.000807 \| 218 \| 0.404 \| 84 \| 50 \| 0 \| 49 \| 132 \| 4 \| 87 \| Lysozyme \| Lysozyme; afdb-uniprot50 \| AF-A0A7H4K6A6-F1-MODEL\_V4 \| 1.0 \| 2.865e-05 \| 218 \| 0.216 \| 189 \| 103 \| 9 \| 3 \| 154 \| 6 \| 186 \| Lysozyme \| Lysozyme; afdb-uniprot50 \| AF-A0A702DCC3-F1-MODEL\_V4 \| 1.0 \| 0.0004751 \| 217 \| 0.295 \| 98 \| 66 \| 1 \| 7 \| 104 \| 21 \| 115 \| Lysozyme \| Lysozyme; afdb-uniprot50 \| AF-A0A0Q6AKV8-F1-MODEL\_V4 \| 1.0 \| 8.717e-05 \| 216 \| 0.293 \| 116 \| 78 \| 2 \| 33 \| 145 \| 6 \| 120 \| Lysozyme \| Lysozyme; afdb-uniprot50 \| AF-A0A7H4PFX8-F1-MODEL\_V4 \| 1.0 \| 4.152e-05 \| 216 \| 0.235 \| 157 \| 104 \| 4 \| 1 \| 153 \| 6 \| 150 \| Lysozyme \| Lysozyme; afdb-uniprot50 \| AF-A0A1Y1J4F4-F1-MODEL\_V4 \| 1.0 \| 0.0002146 \| 215 \| 0.215 \| 176 \| 105 \| 8 \| 1 \| 144 \| 1 \| 175 \| Putative phage lysozyme \| Putative phage lysozyme; afdb-uniprot50 \| AF-V2V153-F1-MODEL\_V4 \| 1.0 \| 0.0002262 \| 214 \| 0.209 \| 143 \| 98 \| 6 \| 15 \| 144 \| 163 \| 303 \| Lysozyme \| Lysozyme; afdb-uniprot50 \| AF-A0A2D9KYN3-F1-MODEL\_V4 \| 1.0 \| 0.0001646 \| 213 \| 0.226 \| 163 \| 102 \| 6 \| 4 \| 145 \| 13 \| 172 \| Lysozyme \| Lysozyme; afdb-uniprot50 \| AF-A0A0N0ZAJ6-F1-MODEL\_V4 \| 1.0 \| 0.0004751 \| 212 \| 0.35 \| 100 \| 62 \| 2 \| 5 \| 104 \| 9 \| 105 \| Lysozyme \| Lysozyme; afdb-uniprot50 \| AF-C4K3M0-F1-MODEL\_V4 \| 1.0 \| 0.0001198 \| 201 \| 0.362 \| 116 \| 65 \| 4 \| 34 \| 145 \| 4 \| 114 \| Lysozyme \| Lysozyme; afdb-uniprot50 \| AF-A0A356LHJ7-F1-MODEL\_V4 \| 1.0 \| 0.0005872 \| 199 \| 0.327 \| 110 \| 69 \| 4 \| 16 \| 124 \| 57 \| 162 \| Lysozyme \| Lysozyme; afdb-uniprot50 \| AF-A0A197JJ61-F1-MODEL\_V4 \| 1.0 \| 0.0007653 \| 199 \| 0.254 \| 122 \| 85 \| 2 \| 2 \| 120 \| 162 \| 280 \| Lysozyme \| Lysozyme; afdb-uniprot50 \| AF-A0A2X1B257-F1-MODEL\_V4 \| 1.0 \| 0.000807 \| 198 \| 0.297 \| 111 \| 77 \| 1 \| 50 \| 159 \| 3 \| 113 \| Phage-related lysozyme (Muraminidase) \| Phage-related lysozyme (Muraminidase); afdb-uniprot50 \| AF-A0A350PK98-F1-MODEL\_V4 \| 1.0 \| 0.0009975 \| 197 \| 0.314 \| 108 \| 68 \| 4 \| 39 \| 144 \| 1 \| 104 \| Lysozyme \| Lysozyme; afdb-uniprot50 \| AF-A0A248KFB5-F1-MODEL\_V4 \| 1.0 \| 0.0006192 \| 197 \| 0.301 \| 116 \| 76 \| 2 \| 29 \| 143 \| 5 \| 116 \| Lysozyme \| Lysozyme; afdb-uniprot50 \| AF-A0A1G4RIK7-F1-MODEL\_V4 \| 1.0 \| 0.0004273 \| 197 \| 0.186 \| 166 \| 100 \| 9 \| 6 \| 144 \| 46 \| 203 \| Lysozyme \| Lysozyme; afdb-uniprot50 \| AF-A0A5B7TXD1-F1-MODEL\_V4 \| 1.0 \| 0.002094 \| 195 \| 0.35 \| 97 \| 57 \| 2 \| 51 \| 143 \| 204 \| 298 \| Lysozyme \| Lysozyme; afdb-uniprot50 \| AF-A0A1S8Y709-F1-MODEL\_V4 \| 1.0 \| 0.001607 \| 190 \| 0.313 \| 86 \| 55 \| 1 \| 60 \| 145 \| 5 \| 86 \| Lysozyme \| Lysozyme; afdb-uniprot50 \| AF-M4UCN0-F1-MODEL\_V4 \| 1.0 \| 0.0006884 \| 188 \| 0.376 \| 85 \| 48 \| 2 \| 60 \| 144 \| 5 \| 84 \| Lysozyme \| Lysozyme; afdb-uniprot50 \| AF-A0A1Z8T0S5-F1-MODEL\_V4 \| 1.0 \| 0.001169 \| 188 \| 0.188 \| 159 \| 106 \| 5 \| 1 \| 143 \| 55 \| 206 \| Uncharacterized protein \| Uncharacterized protein; afdb-uniprot50 \| AF-Q3RDH2-F1-MODEL\_V4 \| 1.0 \| 0.0032 \| 186 \| 0.465 \| 73 \| 39 \| 0 \| 86 \| 158 \| 2 \| 74 \| Lysozyme \| Lysozyme; afdb-uniprot50 \| AF-A0A7W6CC09-F1-MODEL\_V4 \| 1.0 \| 0.001445 \| 186 \| 0.324 \| 108 \| 67 \| 4 \| 38 \| 143 \| 17 \| 120 \| GH24 family phage-related lysozyme (Muramidase) \| GH24 family phage-related lysozyme (Muramidase); afdb-uniprot50 \| AF-A0A854EPE1-F1-MODEL\_V4 \| 1.0 \| 0.0005282 \| 185 \| 0.253 \| 138 \| 80 \| 8 \| 3 \| 129 \| 13 \| 138 \| Glycosyl hydrolase \| Glycosyl hydrolase; afdb-uniprot50 \| AF-A0A447MPG6-F1-MODEL\_V4 \| 1.0 \| 0.004637 \| 182 \| 0.313 \| 83 \| 53 \| 1 \| 63 \| 145 \| 2 \| 80 \| Lysozyme \| Lysozyme; afdb-uniprot50 \| AF-A0A4V1RNB0-F1-MODEL\_V4 \| 1.0 \| 0.0007653 \| 182 \| 0.23 \| 152 \| 75 \| 4 \| 1 \| 148 \| 343 \| 456 \| Uncharacterized protein \| Uncharacterized protein; afdb-uniprot50 \| AF-V6MKU5-F1-MODEL\_V4 \| 1.0 \| 0.0001332 \| 162 \| 0.286 \| 164 \| 80 \| 4 \| 15 \| 145 \| 13 \| 172 \| Lysozyme \| Lysozyme; afdb-uniprot50 \| AF-A0A5M3PTT5-F1-MODEL\_V4 \| 1.0 \| 0.009234 \| 160 \| 0.337 \| 83 \| 50 \| 2 \| 64 \| 146 \| 2 \| 79 \| Lysozyme \| Lysozyme; afdb-uniprot50 \| AF-A0A4Q3MGT4-F1-MODEL\_V4 \| 1.0 \| 0.009234 \| 153 \| 0.176 \| 142 \| 100 \| 6 \| 18 \| 144 \| 2 \| 141 \| Lysozyme \| Lysozyme; afdb-uniprot50 \| AF-A0A6P0BZY4-F1-MODEL\_V4 \| 1.0 \| 0.004889 \| 130 \| 0.234 \| 141 \| 88 \| 4 \| 20 \| 145 \| 3 \| 138 \| Lysozyme \| Lysozyme; afdb-uniprot50 \| AF-A0A418VWW6-F1-MODEL\_V4 \| 1.0 \| 0.0003457 \| 114 \| 0.252 \| 174 \| 109 \| 6 \| 1 \| 159 \| 2 \| 169 \| PG\_binding\_1 domain-containing protein \| PG\_binding\_1 domain-containing protein |
| Top keywords  (threshold 1.00e-02 (evalue)) | **Lysozyme, T4, Endolysin, receptor, chimera, hydrolase, Glycoside, type, Glucagon\_like, peptide** |
| Output files | ../../similar\_structures/64\_FANPEZAQ\_CDS\_0064\_afdb-proteome\_foldseek.tsv ../../similar\_structures/64\_FANPEZAQ\_CDS\_0064\_afdb-uniprot50\_foldseek.tsv ../../similar\_structures/64\_FANPEZAQ\_CDS\_0064\_merged.svg ../../similar\_structures/64\_FANPEZAQ\_CDS\_0064\_pdb\_foldseek.tsv |

  
  
  

Return to summary | Go to previous | Go to next

  


---

**Sequence/structure alignments coloring**  
Each object in the alignment figures is colored according to its E-value following this color coding:

1e-100
10
