## Supplementary material for "Completing the BASEL phage collection to unlock hidden diversity for systematic exploration of phage-host interactions": Data S2: 65.html

FANPEZAQ\_CDS\_0065


Return to summary | Go to previous | Go to next

|  |  |
| --- | --- |
| FANPEZAQ\_CDS\_0065  Page creation date: 02 Sep 2024, 12:00  Project folder: n/a  Input sequences file: Escherichia\_virus\_HeidiAbel.gb | duf2514 lysis domain\_containing fragment lysozyme rz bacteriophage putative spanin endopeptidase prophage phage membrane rz\_like system i\_spanin assembly inner phage\_tail associated hypothetical lambda pspph06 gp55 rac |

### Sequence information

|  |  |
| --- | --- |
| Name | FANPEZAQ\_CDS\_0065  65\_FANPEZAQ\_CDS\_0065 (pipeline id) |
| Imported annotations | Escherichia\_virus\_HeidiAbel Bas97 |
| Protein sequence | MMDKYATPVKLIVAIIIAFALVAFGYVSGFSKSDREWQAKWSARDAADAQAHQQFTEQQR RIEQERQGAIDAIQEQAQQDIATAQRNAAIAAAESKRLQDGIADAITRLQADSGNPGATI SSKTRASNSSLLAELFREIDTAAGIYASEADRARAAGLTCERAYDAVRATSQQ |
| Number of residues | 173 |
| Molecular weight (Da) | 18737.58 |
| Output files | ../../query\_sequences/65\_FANPEZAQ\_CDS\_0065.fasta |

### Putative domain architecture and protein family

#### Search results (HHblits)^1^

|  |  |
| --- | --- |
| Domain family databases searched | Pfam, Ncbi-cd, Cath, Phrogs |
| Results, scheme(s)  (Top layers only; threshold 1.00e-03 (evalue)) | xml version="1.0" encoding="utf-8" standalone="no"?       2024-09-02T21:08:26.615985 image/svg+xml   Matplotlib v3.7.2, https://matplotlib.org/ |
| Results, table  (E-value ≤ 1.00e-03 (evalue)) | db \| id \| prob \| evalue \| pvalue \| score \| cols \| query \| query\_len \| template \| template\_len \| name \| description; pfam \| PF10721 \| 99.7 \| 1e-21 \| 1.9e-25 \| 142.2 \| 139 \| (16, 168) \| 173 \| (8, 150) \| 151 \| DUF2514 \| Protein of unknown function (DUF2514); pfam \| PF03245 \| 98.9 \| 9.2e-14 \| 1.8e-17 \| 96.2 \| 108 \| (52, 167) \| 173 \| (15, 124) \| 126 \| Phage\_lysis \| Bacteriophage Rz lysis protein; phrogs \| 312 \| 100.0 \| 2.2e-35 \| 2.8e-39 \| 229.2 \| 145 \| (19, 172) \| 173 \| (18, 166) \| 170 \| Rz-like spanin \| Rz-like spanin; Category: lysis; NC\_027388\_p53; phrogs \| 1327 \| 99.8 \| 7.2e-25 \| 8.9e-29 \| 170.6 \| 163 \| (2, 172) \| 173 \| (1, 182) \| 188 \| Rz-like spanin \| Rz-like spanin; Category: lysis; KY000219\_p64; phrogs \| 14926 \| 98.9 \| 1.2e-13 \| 1.3e-17 \| 91.1 \| 81 \| (93, 173) \| 173 \| (2, 84) \| 89 \| NA \| NA; Category: unknown function; p303683 VI\_03517; phrogs \| 5777 \| 98.8 \| 7e-13 \| 8.3e-17 \| 99.3 \| 105 \| (56, 171) \| 173 \| (55, 167) \| 169 \| NA \| NA; Category: unknown function; p139219 VI\_05583; phrogs \| 17598 \| 98.0 \| 2.3e-09 \| 2.6e-13 \| 68.0 \| 63 \| (16, 78) \| 173 \| (15, 77) \| 77 \| Rz-like spanin \| Rz-like spanin; Category: lysis; p348110 VI\_03518; phrogs \| 2466 \| 97.6 \| 3.7e-08 \| 4.8e-12 \| 73.4 \| 140 \| (21, 171) \| 173 \| (9, 162) \| 163 \| Rz-like spanin \| Rz-like spanin; Category: lysis; NC\_011589\_p12; phrogs \| 36833 \| 96.7 \| 3.8e-06 \| 4.2e-10 \| 53.0 \| 43 \| (127, 169) \| 173 \| (34, 76) \| 80 \| NA \| NA; Category: unknown function; p236003 VI\_10093; phrogs \| 1026 \| 94.6 \| 0.00055 \| 6.9e-08 \| 46.1 \| 39 \| (1, 39) \| 173 \| (1, 40) \| 101 \| Rz-like spanin \| Rz-like spanin; Category: lysis; MG897800\_p45 |
| Top keywords  (threshold 1.00e-03 (evalue)) | **lysis, Rz\_like, spanin, DUF2514, Bacteriophage, Rz, NC\_027388\_p53, KY000219\_p64, p303683, VI\_03517** |
| Output files | ../../domain\_architecture/65\_FANPEZAQ\_CDS\_0065\_cath.hhr ../../domain\_architecture/65\_FANPEZAQ\_CDS\_0065\_merged.svg ../../domain\_architecture/65\_FANPEZAQ\_CDS\_0065\_ncbi-cd.hhr ../../domain\_architecture/65\_FANPEZAQ\_CDS\_0065\_pfam.hhr ../../domain\_architecture/65\_FANPEZAQ\_CDS\_0065\_phrogs.hhr |

### Identical protein sequences/structures

#### Search results

|  |  |
| --- | --- |
| Protein sequence databases searched | Pdb, Swissprot, Refseq |
| Identical proteins found | -- |
| Top keywords | -- |
| Output files | -- |

### Similar protein sequences/structures

#### Sequence similarity search results (HHblits)^1^

|  |  |
| --- | --- |
| Sequence databases searched | Uniclust, Pdb70 |
| Results, scheme(s)  (Top layers only, threshold 1.00e-03 (evalue)) | xml version="1.0" encoding="utf-8" standalone="no"?       2024-09-02T21:08:55.722460 image/svg+xml   Matplotlib v3.7.2, https://matplotlib.org/ |
| Results, table(s)  (threshold 1.00e-03 (evalue)) | db \| id \| prob \| evalue \| pvalue \| score \| cols \| query \| query\_len \| template \| template\_len \| name \| description; uniclust \| UniRef100\_A0A077KH21 \| 100.0 \| 3.4e-36 \| 7.9e-42 \| 236.0 \| 172 \| (1, 172) \| 173 \| (12, 183) \| 204 \| DUF2514 domain-containing protein \| DUF2514 domain-containing protein; uniclust \| UniRef100\_A0A076LNF4 \| 100.0 \| 2e-35 \| 4.4e-41 \| 225.7 \| 165 \| (7, 171) \| 173 \| (15, 183) \| 193 \| Phage-tail assembly proteins like protein \| Phage-tail assembly proteins like protein; uniclust \| UniRef100\_A0A059VF51 \| 100.0 \| 5.1e-35 \| 1.1e-40 \| 223.2 \| 171 \| (1, 171) \| 173 \| (4, 175) \| 202 \| DUF2514 domain-containing protein \| DUF2514 domain-containing protein; uniclust \| UniRef100\_A0A061KWJ5 \| 100.0 \| 6.3e-34 \| 1.3e-39 \| 219.6 \| 167 \| (2, 172) \| 173 \| (21, 189) \| 212 \| DUF2514 domain-containing protein \| DUF2514 domain-containing protein; uniclust \| UniRef100\_A0A063BGS6 \| 99.9 \| 6.2e-31 \| 1.3e-36 \| 200.5 \| 159 \| (1, 171) \| 173 \| (19, 177) \| 198 \| DUF2514 domain-containing protein \| DUF2514 domain-containing protein; uniclust \| UniRef100\_A0A072TDR1 \| 99.9 \| 3.4e-30 \| 6.6e-36 \| 190.9 \| 165 \| (8, 172) \| 173 \| (9, 176) \| 181 \| DUF2514 family protein (Fragment) \| DUF2514 family protein (Fragment); uniclust \| UniRef100\_A0A022G2T6 \| 99.9 \| 7.7e-29 \| 1.6e-34 \| 197.0 \| 147 \| (19, 172) \| 173 \| (71, 217) \| 251 \| DUF2514 domain-containing protein \| DUF2514 domain-containing protein; uniclust \| UniRef100\_A0A061JLI3 \| 99.9 \| 4.3e-28 \| 8.6e-34 \| 180.3 \| 166 \| (2, 171) \| 173 \| (4, 169) \| 173 \| Endopeptidase \| Endopeptidase; uniclust \| UniRef100\_A0A1B7HMC9 \| 99.9 \| 5.8e-27 \| 1.2e-32 \| 176.8 \| 150 \| (22, 171) \| 173 \| (31, 182) \| 185 \| Putative phage-tail assembly protein \| Putative phage-tail assembly protein; uniclust \| UniRef100\_A0A2R7NV60 \| 99.9 \| 9e-27 \| 1.9e-32 \| 178.1 \| 168 \| (5, 172) \| 173 \| (10, 178) \| 185 \| DUF2514 domain-containing protein \| DUF2514 domain-containing protein; uniclust \| UniRef100\_A0A077FAV2 \| 99.9 \| 2.6e-26 \| 5.6e-32 \| 181.6 \| 144 \| (28, 172) \| 173 \| (64, 207) \| 227 \| DUF2514 family protein \| DUF2514 family protein; uniclust \| UniRef100\_A0A1G6P3N9 \| 99.8 \| 2.4e-24 \| 4.9e-30 \| 165.3 \| 124 \| (47, 172) \| 173 \| (63, 186) \| 192 \| DUF2514 family protein \| DUF2514 family protein; uniclust \| UniRef100\_A0A0A7KTN9 \| 99.8 \| 5e-23 \| 9.1e-29 \| 165.0 \| 147 \| (24, 170) \| 173 \| (235, 385) \| 391 \| Lysozyme \| Lysozyme; uniclust \| UniRef100\_A0A5N8AHL5 \| 99.8 \| 3.3e-22 \| 6.7e-28 \| 152.5 \| 168 \| (5, 172) \| 173 \| (10, 186) \| 196 \| DUF2514 family protein \| DUF2514 family protein; uniclust \| UniRef100\_A0A3M3B9T7 \| 99.8 \| 5.3e-22 \| 9.8e-28 \| 150.7 \| 157 \| (12, 168) \| 173 \| (85, 241) \| 242 \| Prophage PssSM-03, Orf7 \| Prophage PssSM-03, Orf7; uniclust \| UniRef100\_A0A126Z7Q2 \| 99.8 \| 9.7e-22 \| 1.9e-27 \| 147.3 \| 171 \| (1, 171) \| 173 \| (7, 181) \| 182 \| DUF2514 domain-containing protein \| DUF2514 domain-containing protein; uniclust \| UniRef100\_A0A3S0EZ37 \| 99.7 \| 3.4e-21 \| 7.3e-27 \| 151.7 \| 167 \| (4, 171) \| 173 \| (21, 189) \| 206 \| DUF2514 family protein \| DUF2514 family protein; uniclust \| UniRef100\_A0A1G8MFE6 \| 99.7 \| 3e-20 \| 6e-26 \| 140.8 \| 159 \| (7, 171) \| 173 \| (10, 168) \| 187 \| DUF2514 domain-containing protein \| DUF2514 domain-containing protein; uniclust \| UniRef100\_A0A6J5L3W2 \| 99.7 \| 4.3e-20 \| 9e-26 \| 143.1 \| 154 \| (1, 172) \| 173 \| (1, 157) \| 190 \| Spanin, inner membrane subunit \| Spanin, inner membrane subunit; uniclust \| UniRef100\_UPI000F0145F3 \| 99.7 \| 5.9e-20 \| 1.1e-25 \| 134.3 \| 169 \| (2, 170) \| 173 \| (1, 169) \| 171 \| DUF2514 family protein \| DUF2514 family protein; uniclust \| UniRef100\_A0A072T468 \| 99.7 \| 1.1e-19 \| 2.2e-25 \| 142.2 \| 121 \| (51, 171) \| 173 \| (68, 188) \| 216 \| DUF2514 family protein \| DUF2514 family protein; uniclust \| UniRef100\_A0A1X0YIG9 \| 99.7 \| 1.4e-19 \| 2.7e-25 \| 134.9 \| 160 \| (4, 167) \| 173 \| (7, 168) \| 168 \| DUF2514 domain-containing protein (Fragment) \| DUF2514 domain-containing protein (Fragment); uniclust \| UniRef100\_A0A2P1VUX7 \| 99.7 \| 1.7e-19 \| 3.1e-25 \| 135.3 \| 167 \| (3, 169) \| 173 \| (10, 186) \| 203 \| DUF2514 domain-containing protein \| DUF2514 domain-containing protein; uniclust \| UniRef100\_A0A6N3TCN9 \| 99.7 \| 1.8e-19 \| 3.2e-25 \| 136.2 \| 143 \| (23, 168) \| 173 \| (81, 223) \| 224 \| DUF2514 domain-containing protein \| DUF2514 domain-containing protein; uniclust \| UniRef100\_A0A0P9JKE3 \| 99.6 \| 2.2e-19 \| 4.4e-25 \| 132.1 \| 123 \| (48, 172) \| 173 \| (12, 134) \| 138 \| DUF2514 domain-containing protein (Fragment) \| DUF2514 domain-containing protein (Fragment); uniclust \| UniRef100\_A0A0J6KI14 \| 99.6 \| 4.1e-19 \| 8.6e-25 \| 138.0 \| 163 \| (1, 172) \| 173 \| (14, 177) \| 189 \| Lysozyme \| Lysozyme; uniclust \| UniRef100\_A0A2P0N9U2 \| 99.6 \| 1.4e-18 \| 2.8e-24 \| 130.6 \| 169 \| (1, 171) \| 173 \| (1, 169) \| 170 \| Endopeptidase \| Endopeptidase; uniclust \| UniRef100\_A0A0C1YJT8 \| 99.6 \| 2.2e-18 \| 4.6e-24 \| 131.5 \| 155 \| (4, 166) \| 173 \| (8, 162) \| 165 \| Lysozyme \| Lysozyme; uniclust \| UniRef100\_A0A0B5CPB3 \| 99.6 \| 6.3e-18 \| 1.3e-23 \| 128.1 \| 158 \| (1, 163) \| 173 \| (1, 158) \| 168 \| Phage associated membrane protein \| Phage associated membrane protein; uniclust \| UniRef100\_A0A1T1ANQ7 \| 99.6 \| 6.7e-18 \| 1.4e-23 \| 129.0 \| 162 \| (2, 172) \| 173 \| (1, 162) \| 169 \| Lysis protein \| Lysis protein; uniclust \| UniRef100\_C5CJR8 \| 99.5 \| 1.1e-17 \| 2.2e-23 \| 126.2 \| 138 \| (33, 171) \| 173 \| (37, 174) \| 179 \| DUF2514 family protein \| DUF2514 family protein; uniclust \| UniRef100\_Q2T6D5 \| 99.5 \| 1.2e-17 \| 2.2e-23 \| 126.5 \| 158 \| (2, 171) \| 173 \| (53, 210) \| 212 \| Gp23 \| Gp23; uniclust \| UniRef100\_A0A2A2JXU9 \| 99.5 \| 3.2e-17 \| 5.9e-23 \| 129.1 \| 157 \| (14, 170) \| 173 \| (138, 294) \| 302 \| lysozyme \| lysozyme; uniclust \| UniRef100\_A0A072T277 \| 99.5 \| 7.2e-17 \| 1.5e-22 \| 126.9 \| 151 \| (7, 169) \| 173 \| (26, 183) \| 205 \| DUF2514 family protein \| DUF2514 family protein; uniclust \| UniRef100\_A0A009ZWR4 \| 99.5 \| 7.7e-17 \| 1.6e-22 \| 127.2 \| 159 \| (4, 170) \| 173 \| (36, 199) \| 204 \| Uncharacterized protein \| Uncharacterized protein; uniclust \| UniRef100\_A0A072T346 \| 99.4 \| 2.5e-16 \| 5.2e-22 \| 123.4 \| 168 \| (1, 170) \| 173 \| (5, 179) \| 200 \| DUF2514 family protein \| DUF2514 family protein; uniclust \| UniRef100\_A0A076PZD4 \| 99.4 \| 3.9e-16 \| 8.6e-22 \| 123.9 \| 123 \| (43, 172) \| 173 \| (53, 177) \| 201 \| Lysozyme \| Lysozyme; uniclust \| UniRef100\_UPI000D0449AE \| 99.4 \| 5.6e-16 \| 1e-21 \| 124.0 \| 157 \| (2, 158) \| 173 \| (5, 167) \| 338 \| DUF2514 family protein \| DUF2514 family protein; uniclust \| UniRef100\_A0A0J5GKH4 \| 99.4 \| 8e-16 \| 1.8e-21 \| 124.5 \| 143 \| (20, 170) \| 173 \| (42, 188) \| 225 \| Lysozyme \| Lysozyme; uniclust \| UniRef100\_A0A2T5HJ49 \| 99.4 \| 9.3e-16 \| 1.8e-21 \| 125.9 \| 158 \| (6, 170) \| 173 \| (188, 345) \| 347 \| Putative chitinase \| Putative chitinase; uniclust \| UniRef100\_A0A031HSV0 \| 99.4 \| 1e-15 \| 2e-21 \| 115.8 \| 166 \| (1, 168) \| 173 \| (1, 169) \| 186 \| Uncharacterized protein \| Uncharacterized protein; uniclust \| UniRef100\_A0A376THA0 \| 99.3 \| 3.3e-15 \| 6.2e-21 \| 111.7 \| 154 \| (20, 173) \| 173 \| (9, 164) \| 181 \| Putative prophage protein \| Putative prophage protein; uniclust \| UniRef100\_UPI001FB7112E \| 99.3 \| 4.5e-15 \| 8.2e-21 \| 109.5 \| 118 \| (55, 172) \| 173 \| (54, 171) \| 175 \| DUF2514 domain-containing protein \| DUF2514 domain-containing protein; uniclust \| UniRef100\_A0A096F537 \| 99.3 \| 4.1e-15 \| 8.8e-21 \| 115.1 \| 128 \| (32, 167) \| 173 \| (37, 164) \| 175 \| Endopeptidase \| Endopeptidase; uniclust \| UniRef100\_A0A737M960 \| 99.3 \| 5.1e-15 \| 9.7e-21 \| 105.2 \| 116 \| (56, 171) \| 173 \| (4, 121) \| 125 \| DUF2514 family protein (Fragment) \| DUF2514 family protein (Fragment); uniclust \| UniRef100\_A0A0T9L9I2 \| 99.3 \| 5.2e-15 \| 1.1e-20 \| 117.0 \| 160 \| (3, 169) \| 173 \| (9, 176) \| 195 \| Bacteriophage lysis protein \| Bacteriophage lysis protein; uniclust \| UniRef100\_A0A5E6S0Z6 \| 99.3 \| 9.2e-15 \| 1.7e-20 \| 105.2 \| 127 \| (12, 138) \| 173 \| (11, 139) \| 139 \| DUF2514 domain-containing protein \| DUF2514 domain-containing protein; uniclust \| UniRef100\_A0A077LBX1 \| 99.3 \| 1.3e-14 \| 2.8e-20 \| 117.1 \| 129 \| (33, 169) \| 173 \| (61, 194) \| 217 \| Prophage PSPPH06, lysis protein \| Prophage PSPPH06, lysis protein; uniclust \| UniRef100\_A0A0N0GNZ7 \| 99.3 \| 2.1e-14 \| 4.1e-20 \| 108.9 \| 161 \| (3, 170) \| 173 \| (5, 167) \| 169 \| Bacteriophage lysis protein \| Bacteriophage lysis protein; uniclust \| UniRef100\_A0A0Q4GQ76 \| 99.3 \| 2.4e-14 \| 4.5e-20 \| 105.9 \| 167 \| (1, 168) \| 173 \| (1, 175) \| 176 \| DUF2514 domain-containing protein \| DUF2514 domain-containing protein; uniclust \| UniRef100\_UPI000F06DD48 \| 99.2 \| 6.7e-14 \| 1.2e-19 \| 97.7 \| 116 \| (25, 140) \| 173 \| (1, 116) \| 116 \| DUF2514 family protein \| DUF2514 family protein; uniclust \| UniRef100\_UPI00065FE3D1 \| 99.2 \| 1.3e-13 \| 2.5e-19 \| 100.9 \| 143 \| (15, 157) \| 173 \| (17, 161) \| 161 \| DUF2514 family protein \| DUF2514 family protein; uniclust \| UniRef100\_A0A0E3T9S8 \| 99.1 \| 2.6e-13 \| 5e-19 \| 104.5 \| 165 \| (2, 171) \| 173 \| (32, 196) \| 201 \| Tail assembly protein \| Tail assembly protein; uniclust \| UniRef100\_UPI00077A1791 \| 99.1 \| 3e-13 \| 5.9e-19 \| 99.2 \| 105 \| (64, 171) \| 173 \| (31, 135) \| 141 \| DUF2514 family protein \| DUF2514 family protein; uniclust \| UniRef100\_A0A2I6PHU9 \| 99.1 \| 4.2e-13 \| 8.1e-19 \| 102.1 \| 122 \| (46, 170) \| 173 \| (44, 165) \| 178 \| DUF2514 family protein \| DUF2514 family protein; uniclust \| UniRef100\_UPI0021A505FF \| 99.1 \| 4.9e-13 \| 9e-19 \| 96.4 \| 98 \| (72, 170) \| 173 \| (32, 129) \| 143 \| DUF2514 domain-containing protein \| DUF2514 domain-containing protein; uniclust \| UniRef100\_A0A447M6P4 \| 99.1 \| 5.1e-13 \| 1e-18 \| 97.9 \| 117 \| (56, 172) \| 173 \| (6, 124) \| 136 \| Protein gp55 \| Protein gp55; uniclust \| UniRef100\_A0A447N8Z3 \| 99.1 \| 6.1e-13 \| 1.1e-18 \| 97.9 \| 121 \| (52, 172) \| 173 \| (18, 140) \| 151 \| Protein gp55 \| Protein gp55; uniclust \| UniRef100\_A0A059KU98 \| 99.0 \| 1.3e-12 \| 2.6e-18 \| 100.1 \| 119 \| (51, 171) \| 173 \| (70, 188) \| 190 \| Endopeptidase \| Endopeptidase; uniclust \| UniRef100\_UPI001F13E5F8 \| 99.0 \| 1.7e-12 \| 3.1e-18 \| 99.4 \| 122 \| (46, 169) \| 173 \| (92, 213) \| 215 \| DUF2514 family protein \| DUF2514 family protein; uniclust \| UniRef100\_A0A031GZS8 \| 99.0 \| 1.5e-12 \| 3.2e-18 \| 105.8 \| 114 \| (51, 171) \| 173 \| (97, 215) \| 230 \| Bacteriophage lysis protein \| Bacteriophage lysis protein; uniclust \| UniRef100\_A0A418X1B9 \| 99.0 \| 2.1e-12 \| 4.1e-18 \| 98.4 \| 161 \| (2, 172) \| 173 \| (1, 163) \| 172 \| Lysis protein \| Lysis protein; uniclust \| UniRef100\_A0A6J5N0Y4 \| 99.0 \| 2.3e-12 \| 4.6e-18 \| 97.9 \| 154 \| (1, 166) \| 173 \| (3, 157) \| 158 \| Uncharacterized protein \| Uncharacterized protein; uniclust \| UniRef100\_UPI001E623135 \| 99.0 \| 3.1e-12 \| 5.7e-18 \| 89.8 \| 111 \| (58, 168) \| 173 \| (6, 118) \| 119 \| DUF2514 family protein \| DUF2514 family protein; uniclust \| UniRef100\_A0A612N5Q1 \| 99.0 \| 4.3e-12 \| 8.1e-18 \| 90.3 \| 107 \| (4, 110) \| 173 \| (7, 117) \| 125 \| DUF2514 family protein (Fragment) \| DUF2514 family protein (Fragment); uniclust \| UniRef100\_A0A3N7DP53 \| 98.9 \| 8.8e-12 \| 1.6e-17 \| 97.4 \| 107 \| (61, 169) \| 173 \| (141, 247) \| 248 \| DUF2514 domain-containing protein \| DUF2514 domain-containing protein; uniclust \| UniRef100\_A0A0E3GMK9 \| 98.9 \| 9.8e-12 \| 1.9e-17 \| 95.0 \| 109 \| (60, 171) \| 173 \| (62, 170) \| 180 \| Lysin \| Lysin; uniclust \| UniRef100\_A0A7G9RJE6 \| 98.9 \| 1.1e-11 \| 2e-17 \| 91.8 \| 95 \| (76, 171) \| 173 \| (48, 142) \| 165 \| DUF2514 family protein \| DUF2514 family protein; uniclust \| UniRef100\_UPI000A6BEC42 \| 98.9 \| 1.4e-11 \| 2.6e-17 \| 89.3 \| 116 \| (55, 171) \| 173 \| (19, 134) \| 144 \| DUF2514 family protein \| DUF2514 family protein; uniclust \| UniRef100\_UPI000209A3A1 \| 98.9 \| 1.5e-11 \| 2.9e-17 \| 85.1 \| 94 \| (8, 101) \| 173 \| (5, 101) \| 103 \| DUF2514 family protein \| DUF2514 family protein; uniclust \| UniRef100\_UPI0009BD088E \| 98.8 \| 2.6e-11 \| 4.7e-17 \| 85.4 \| 103 \| (9, 111) \| 173 \| (9, 111) \| 118 \| DUF2514 domain-containing protein \| DUF2514 domain-containing protein; uniclust \| UniRef100\_A0A736RDN2 \| 98.8 \| 4.6e-11 \| 8.6e-17 \| 86.5 \| 105 \| (2, 110) \| 173 \| (5, 109) \| 130 \| DUF2514 family protein (Fragment) \| DUF2514 family protein (Fragment); uniclust \| UniRef100\_UPI000F846E2F \| 98.8 \| 4.6e-11 \| 8.8e-17 \| 78.6 \| 57 \| (116, 172) \| 173 \| (13, 69) \| 71 \| DUF2514 family protein \| DUF2514 family protein; uniclust \| UniRef100\_A0A0H3LPM7 \| 98.8 \| 5.5e-11 \| 1.2e-16 \| 93.3 \| 149 \| (1, 169) \| 173 \| (10, 162) \| 176 \| Uncharacterized protein \| Uncharacterized protein; uniclust \| UniRef100\_UPI001BA9F773 \| 98.8 \| 6.4e-11 \| 1.2e-16 \| 84.8 \| 106 \| (54, 164) \| 173 \| (23, 128) \| 130 \| DUF2514 family protein \| DUF2514 family protein; uniclust \| UniRef100\_A0A7Z3BLE0 \| 98.8 \| 6.9e-11 \| 1.3e-16 \| 82.8 \| 90 \| (2, 91) \| 173 \| (1, 90) \| 108 \| DUF2514 family protein \| DUF2514 family protein; uniclust \| UniRef100\_A0A329HWK1 \| 98.7 \| 9.5e-11 \| 2e-16 \| 93.3 \| 113 \| (51, 170) \| 173 \| (64, 183) \| 197 \| Lysis protein \| Lysis protein; uniclust \| UniRef100\_UPI001CF29D96 \| 98.7 \| 1.6e-10 \| 2.8e-16 \| 80.3 \| 100 \| (72, 171) \| 173 \| (3, 104) \| 106 \| DUF2514 family protein \| DUF2514 family protein; uniclust \| UniRef100\_A0A965JDW4 \| 98.7 \| 1.6e-10 \| 3e-16 \| 87.0 \| 158 \| (2, 168) \| 173 \| (1, 161) \| 164 \| DUF2514 family protein \| DUF2514 family protein; uniclust \| UniRef100\_A0KJY0 \| 98.7 \| 2.1e-10 \| 3.9e-16 \| 88.9 \| 87 \| (19, 105) \| 173 \| (81, 171) \| 223 \| Uncharacterized protein \| Uncharacterized protein; uniclust \| UniRef100\_UPI001C594AF0 \| 98.7 \| 2.2e-10 \| 4e-16 \| 85.8 \| 160 \| (4, 170) \| 173 \| (7, 166) \| 168 \| DUF2514 family protein \| DUF2514 family protein; uniclust \| UniRef100\_UPI00033CB928 \| 98.6 \| 3.1e-10 \| 5.7e-16 \| 82.7 \| 90 \| (4, 93) \| 173 \| (7, 96) \| 143 \| DUF2514 family protein \| DUF2514 family protein; uniclust \| UniRef100\_A0A431IGX4 \| 98.6 \| 3.1e-10 \| 6.2e-16 \| 88.1 \| 145 \| (17, 171) \| 173 \| (21, 165) \| 171 \| DUF2514 family protein \| DUF2514 family protein; uniclust \| UniRef100\_A0A0P0HS49 \| 98.6 \| 3.4e-10 \| 6.4e-16 \| 85.7 \| 105 \| (63, 170) \| 173 \| (57, 161) \| 171 \| Endopeptidase Rz \| Endopeptidase Rz; uniclust \| UniRef100\_A0A379W8I7 \| 98.6 \| 3.6e-10 \| 6.9e-16 \| 81.2 \| 104 \| (3, 110) \| 173 \| (6, 109) \| 119 \| Exported phage protein \| Exported phage protein; uniclust \| UniRef100\_A0A9E6U0C0 \| 98.6 \| 4.5e-10 \| 8.2e-16 \| 78.1 \| 74 \| (95, 168) \| 173 \| (32, 105) \| 106 \| DUF2514 domain-containing protein \| DUF2514 domain-containing protein; uniclust \| UniRef100\_UPI000EA8E18C \| 98.6 \| 4.6e-10 \| 8.4e-16 \| 81.3 \| 117 \| (52, 169) \| 173 \| (19, 135) \| 136 \| DUF2514 family protein \| DUF2514 family protein; uniclust \| UniRef100\_A0A0H5AUG6 \| 98.6 \| 4.6e-10 \| 9.8e-16 \| 89.7 \| 138 \| (26, 170) \| 173 \| (26, 175) \| 188 \| Lysozyme \| Lysozyme; uniclust \| UniRef100\_UPI00027B3473 \| 98.6 \| 6e-10 \| 1.1e-15 \| 75.2 \| 72 \| (95, 170) \| 173 \| (16, 87) \| 89 \| DUF2514 family protein \| DUF2514 family protein; uniclust \| UniRef100\_A0A010SYQ9 \| 98.6 \| 6.1e-10 \| 1.3e-15 \| 91.5 \| 140 \| (25, 171) \| 173 \| (45, 198) \| 229 \| Lysozyme \| Lysozyme; uniclust \| UniRef100\_A0A5A9ZEY6 \| 98.5 \| 7.7e-10 \| 1.5e-15 \| 84.5 \| 148 \| (4, 163) \| 173 \| (5, 153) \| 157 \| DUF2514 family protein \| DUF2514 family protein; uniclust \| UniRef100\_F0FWP2 \| 98.5 \| 1.3e-09 \| 2.3e-15 \| 72.1 \| 54 \| (119, 172) \| 173 \| (14, 67) \| 75 \| DUF2514 family protein (Fragment) \| DUF2514 family protein (Fragment); uniclust \| UniRef100\_A0A3B9Q8I9 \| 98.5 \| 1.2e-09 \| 2.5e-15 \| 84.2 \| 145 \| (5, 153) \| 173 \| (3, 149) \| 159 \| DUF2514 domain-containing protein \| DUF2514 domain-containing protein; uniclust \| UniRef100\_A0A7H1MBE5 \| 98.5 \| 1.7e-09 \| 3.3e-15 \| 83.3 \| 157 \| (1, 163) \| 173 \| (1, 167) \| 174 \| Uncharacterized protein \| Uncharacterized protein; uniclust \| UniRef100\_A0A935YVG0 \| 98.4 \| 2.5e-09 \| 4.5e-15 \| 80.0 \| 99 \| (65, 166) \| 173 \| (66, 164) \| 165 \| DUF2514 family protein \| DUF2514 family protein; uniclust \| UniRef100\_UPI001F0836F7 \| 98.4 \| 2.6e-09 \| 4.7e-15 \| 71.5 \| 78 \| (1, 78) \| 173 \| (1, 82) \| 83 \| DUF2514 family protein \| DUF2514 family protein; uniclust \| UniRef100\_A0A3S0F432 \| 98.4 \| 2.7e-09 \| 4.9e-15 \| 81.7 \| 115 \| (46, 168) \| 173 \| (79, 193) \| 195 \| DUF2514 family protein \| DUF2514 family protein; uniclust \| UniRef100\_UPI001454EF55 \| 98.4 \| 3.2e-09 \| 5.9e-15 \| 75.1 \| 71 \| (21, 91) \| 173 \| (25, 99) \| 110 \| DUF2514 family protein \| DUF2514 family protein; uniclust \| UniRef100\_A0A965UN57 \| 98.4 \| 3.2e-09 \| 6.1e-15 \| 79.6 \| 157 \| (1, 173) \| 173 \| (1, 157) \| 158 \| Uncharacterized protein \| Uncharacterized protein; uniclust \| UniRef100\_A0A377XDG9 \| 98.4 \| 4.5e-09 \| 8.2e-15 \| 83.7 \| 73 \| (93, 168) \| 173 \| (187, 259) \| 260 \| Putative glycoside hydrolase \| Putative glycoside hydrolase; uniclust \| UniRef100\_A0A5B0BND6 \| 98.3 \| 5.6e-09 \| 1.1e-14 \| 62.9 \| 41 \| (130, 170) \| 173 \| (2, 42) \| 43 \| DUF2514 family protein \| DUF2514 family protein; uniclust \| UniRef100\_A0A3S4K6X8 \| 98.3 \| 5.8e-09 \| 1.1e-14 \| 74.6 \| 78 \| (96, 173) \| 173 \| (21, 100) \| 107 \| Putative prophage protein \| Putative prophage protein; uniclust \| UniRef100\_A0A085JGU0 \| 98.3 \| 5.5e-09 \| 1.2e-14 \| 83.9 \| 116 \| (47, 170) \| 173 \| (53, 170) \| 183 \| Rz family phage lysis protein \| Rz family phage lysis protein; uniclust \| UniRef100\_A0A540WHN8 \| 98.3 \| 6.5e-09 \| 1.2e-14 \| 74.0 \| 96 \| (74, 169) \| 173 \| (8, 108) \| 118 \| DUF2514 family protein (Fragment) \| DUF2514 family protein (Fragment); uniclust \| UniRef100\_UPI002113A69C \| 98.3 \| 8.2e-09 \| 1.5e-14 \| 71.4 \| 85 \| (84, 168) \| 173 \| (13, 98) \| 99 \| DUF2514 family protein \| DUF2514 family protein; uniclust \| UniRef100\_A0A5T8J8N9 \| 98.3 \| 1.1e-08 \| 2.1e-14 \| 69.5 \| 74 \| (3, 76) \| 173 \| (6, 83) \| 85 \| DUF2514 domain-containing protein (Fragment) \| DUF2514 domain-containing protein (Fragment); uniclust \| UniRef100\_A0A433SCY5 \| 98.3 \| 1.1e-08 \| 2.2e-14 \| 78.8 \| 152 \| (2, 167) \| 173 \| (1, 159) \| 168 \| Uncharacterized protein \| Uncharacterized protein; uniclust \| UniRef100\_A0A1E8PNC0 \| 98.3 \| 1.2e-08 \| 2.2e-14 \| 79.0 \| 157 \| (6, 170) \| 173 \| (11, 172) \| 178 \| Bacteriophage lysis protein \| Bacteriophage lysis protein; uniclust \| UniRef100\_A0A6J5KTC4 \| 98.3 \| 1.3e-08 \| 2.4e-14 \| 75.4 \| 112 \| (20, 168) \| 173 \| (18, 129) \| 130 \| Spanin, inner membrane subunit \| Spanin, inner membrane subunit; uniclust \| UniRef100\_UPI00051BC885 \| 98.3 \| 1.4e-08 \| 2.5e-14 \| 72.9 \| 86 \| (25, 110) \| 173 \| (34, 119) \| 122 \| DUF2514 family protein \| DUF2514 family protein; uniclust \| UniRef100\_A0A6G6IUM5 \| 98.2 \| 1.6e-08 \| 3e-14 \| 70.6 \| 77 \| (95, 171) \| 173 \| (21, 99) \| 101 \| DUF2514 family protein \| DUF2514 family protein; uniclust \| UniRef100\_A0A1E8CG11 \| 98.2 \| 1.6e-08 \| 3.3e-14 \| 80.0 \| 153 \| (8, 168) \| 173 \| (13, 173) \| 179 \| Lysis protein \| Lysis protein; uniclust \| UniRef100\_UPI001C0D8754 \| 98.2 \| 2e-08 \| 3.7e-14 \| 74.7 \| 144 \| (9, 162) \| 173 \| (5, 148) \| 155 \| DUF2514 family protein \| DUF2514 family protein; uniclust \| UniRef100\_UPI000E69ED77 \| 98.2 \| 2.1e-08 \| 3.8e-14 \| 66.4 \| 59 \| (113, 171) \| 173 \| (7, 65) \| 76 \| DUF2514 family protein \| DUF2514 family protein; uniclust \| UniRef100\_A0A255ZJS3 \| 98.2 \| 2.4e-08 \| 4.7e-14 \| 76.8 \| 154 \| (1, 172) \| 173 \| (1, 154) \| 163 \| Lysis protein \| Lysis protein; uniclust \| UniRef100\_UPI001E6220F1 \| 98.2 \| 3.2e-08 \| 6e-14 \| 67.7 \| 56 \| (113, 168) \| 173 \| (29, 84) \| 87 \| DUF2514 family protein \| DUF2514 family protein; uniclust \| UniRef100\_A0A136Q9N2 \| 98.1 \| 3.3e-08 \| 6.3e-14 \| 70.8 \| 101 \| (6, 106) \| 173 \| (1, 101) \| 109 \| Uncharacterized protein (Fragment) \| Uncharacterized protein (Fragment); uniclust \| UniRef100\_A0A0T9Q0I4 \| 98.1 \| 4.1e-08 \| 7.9e-14 \| 76.6 \| 147 \| (15, 166) \| 173 \| (22, 172) \| 185 \| Protein of uncharacterized function (DUF2514) \| Protein of uncharacterized function (DUF2514); uniclust \| UniRef100\_A0A2W5L444 \| 98.1 \| 4.3e-08 \| 8e-14 \| 71.8 \| 124 \| (3, 136) \| 173 \| (4, 127) \| 127 \| DUF2514 domain-containing protein (Fragment) \| DUF2514 domain-containing protein (Fragment); uniclust \| UniRef100\_A0A3B8P7Q2 \| 98.1 \| 4.3e-08 \| 8.1e-14 \| 66.1 \| 49 \| (121, 169) \| 173 \| (28, 76) \| 78 \| DUF2514 domain-containing protein (Fragment) \| DUF2514 domain-containing protein (Fragment); uniclust \| UniRef100\_A0A6J5KHQ3 \| 98.1 \| 4.3e-08 \| 8.4e-14 \| 75.3 \| 147 \| (4, 165) \| 173 \| (11, 157) \| 158 \| Uncharacterized protein \| Uncharacterized protein; uniclust \| UniRef100\_A0A3M2U176 \| 98.1 \| 5.1e-08 \| 9.5e-14 \| 65.4 \| 69 \| (8, 76) \| 173 \| (6, 77) \| 78 \| Uncharacterized protein (Fragment) \| Uncharacterized protein (Fragment); uniclust \| UniRef100\_A0A5R1YM75 \| 98.1 \| 5.4e-08 \| 1e-13 \| 65.0 \| 56 \| (117, 172) \| 173 \| (9, 64) \| 76 \| DUF2514 domain-containing protein (Fragment) \| DUF2514 domain-containing protein (Fragment); uniclust \| UniRef100\_A0A4Q5VW04 \| 98.1 \| 6e-08 \| 1.1e-13 \| 69.8 \| 87 \| (75, 167) \| 173 \| (36, 122) \| 122 \| DUF2514 family protein \| DUF2514 family protein; uniclust \| UniRef100\_F0FWN7 \| 98.1 \| 7.6e-08 \| 1.4e-13 \| 68.1 \| 101 \| (2, 110) \| 173 \| (1, 101) \| 109 \| Putative bacteriophage protein (Fragment) \| Putative bacteriophage protein (Fragment); uniclust \| UniRef100\_A0A533I619 \| 98.0 \| 7.9e-08 \| 1.5e-13 \| 72.0 \| 138 \| (32, 169) \| 173 \| (20, 157) \| 158 \| DUF2514 family protein \| DUF2514 family protein; uniclust \| UniRef100\_A0A6J5LN54 \| 98.0 \| 8.6e-08 \| 1.6e-13 \| 73.1 \| 121 \| (32, 169) \| 173 \| (29, 154) \| 158 \| Spanin, inner membrane subunit \| Spanin, inner membrane subunit; uniclust \| UniRef100\_A0A7Z3BS56 \| 98.0 \| 1.2e-07 \| 2.2e-13 \| 62.3 \| 58 \| (114, 171) \| 173 \| (8, 65) \| 68 \| DUF2514 family protein \| DUF2514 family protein; uniclust \| UniRef100\_A0A068Z3N6 \| 98.0 \| 1.1e-07 \| 2.4e-13 \| 76.3 \| 114 \| (50, 171) \| 173 \| (47, 161) \| 173 \| Lysis protein \| Lysis protein; uniclust \| UniRef100\_A0A1W6JT24 \| 98.0 \| 1.3e-07 \| 2.4e-13 \| 68.7 \| 96 \| (10, 105) \| 173 \| (10, 105) \| 128 \| Uncharacterized protein \| Uncharacterized protein; uniclust \| UniRef100\_A0A447M6N0 \| 98.0 \| 1.3e-07 \| 2.5e-13 \| 65.4 \| 51 \| (121, 171) \| 173 \| (35, 85) \| 87 \| Phage-tail assembly protein \| Phage-tail assembly protein; uniclust \| UniRef100\_A0A136Q9T1 \| 98.0 \| 1.4e-07 \| 2.6e-13 \| 66.1 \| 80 \| (90, 170) \| 173 \| (6, 85) \| 95 \| DUF2514 family protein (Fragment) \| DUF2514 family protein (Fragment); uniclust \| UniRef100\_A0A0A8F3R3 \| 98.0 \| 1.5e-07 \| 2.7e-13 \| 73.5 \| 159 \| (2, 167) \| 173 \| (40, 202) \| 208 \| Phage associated membrane protein \| Phage associated membrane protein; uniclust \| UniRef100\_UPI00111B1EBA \| 98.0 \| 1.7e-07 \| 3.1e-13 \| 67.3 \| 85 \| (26, 110) \| 173 \| (15, 99) \| 113 \| DUF2514 family protein \| DUF2514 family protein; uniclust \| UniRef100\_A0A6J5KRF4 \| 97.9 \| 2e-07 \| 3.6e-13 \| 69.8 \| 121 \| (42, 168) \| 173 \| (32, 152) \| 154 \| Spanin, inner membrane subunit \| Spanin, inner membrane subunit; uniclust \| UniRef100\_A0A7J5P9B2 \| 97.9 \| 2e-07 \| 3.6e-13 \| 61.5 \| 70 \| (96, 168) \| 173 \| (2, 71) \| 72 \| DUF2514 family protein \| DUF2514 family protein; uniclust \| UniRef100\_UPI001AFE8EB9 \| 97.9 \| 2.5e-07 \| 4.7e-13 \| 72.2 \| 138 \| (29, 172) \| 173 \| (33, 181) \| 185 \| hypothetical protein \| hypothetical protein; uniclust \| UniRef100\_A0A2W5FEN7 \| 97.9 \| 2.6e-07 \| 4.7e-13 \| 55.9 \| 42 \| (130, 171) \| 173 \| (1, 42) \| 46 \| DUF2514 domain-containing protein (Fragment) \| DUF2514 domain-containing protein (Fragment); uniclust \| UniRef100\_A0A0G9H235 \| 97.8 \| 4.1e-07 \| 7.5e-13 \| 65.9 \| 104 \| (62, 168) \| 173 \| (20, 123) \| 123 \| DUF2514 domain-containing protein (Fragment) \| DUF2514 domain-containing protein (Fragment); uniclust \| UniRef100\_A0A8T5ZKZ9 \| 97.8 \| 3.9e-07 \| 7.8e-13 \| 64.3 \| 63 \| (24, 86) \| 173 \| (13, 75) \| 87 \| DUF2514 family protein (Fragment) \| DUF2514 family protein (Fragment); uniclust \| UniRef100\_A0A2S0MB23 \| 97.8 \| 4.4e-07 \| 8.7e-13 \| 72.3 \| 130 \| (33, 164) \| 173 \| (37, 171) \| 190 \| DUF2514 domain-containing protein \| DUF2514 domain-containing protein; uniclust \| UniRef100\_A0A2K9VHR6 \| 97.8 \| 4.8e-07 \| 8.8e-13 \| 68.7 \| 159 \| (6, 169) \| 173 \| (3, 166) \| 167 \| Endopeptidase \| Endopeptidase; uniclust \| UniRef100\_A0A8I1RBJ4 \| 97.8 \| 5.8e-07 \| 1.1e-12 \| 68.3 \| 132 \| (29, 167) \| 173 \| (29, 160) \| 167 \| DUF2514 family protein \| DUF2514 family protein; uniclust \| UniRef100\_A0A178MV11 \| 97.8 \| 5.6e-07 \| 1.1e-12 \| 70.2 \| 154 \| (5, 164) \| 173 \| (2, 159) \| 180 \| Uncharacterized protein \| Uncharacterized protein; uniclust \| UniRef100\_A0A526VKI8 \| 97.8 \| 6.3e-07 \| 1.2e-12 \| 71.0 \| 134 \| (21, 162) \| 173 \| (23, 156) \| 195 \| DUF2514 family protein \| DUF2514 family protein; uniclust \| UniRef100\_A0A068QVD3 \| 97.8 \| 5.6e-07 \| 1.2e-12 \| 75.3 \| 76 \| (28, 110) \| 173 \| (54, 136) \| 230 \| Putative endopeptidase \| Putative endopeptidase; uniclust \| UniRef100\_A0A7H0HJ18 \| 97.8 \| 6.3e-07 \| 1.2e-12 \| 68.7 \| 107 \| (55, 169) \| 173 \| (27, 133) \| 144 \| Uncharacterized protein \| Uncharacterized protein; uniclust \| UniRef100\_UPI001D0EDDCC \| 97.8 \| 7e-07 \| 1.3e-12 \| 59.8 \| 50 \| (120, 169) \| 173 \| (25, 74) \| 77 \| DUF2514 family protein \| DUF2514 family protein; uniclust \| UniRef100\_A0A6N4T4E4 \| 97.8 \| 6.9e-07 \| 1.3e-12 \| 68.7 \| 116 \| (50, 172) \| 173 \| (45, 163) \| 168 \| Uncharacterized protein \| Uncharacterized protein; uniclust \| UniRef100\_A0A061YD13 \| 97.7 \| 6.3e-07 \| 1.4e-12 \| 74.8 \| 111 \| (50, 168) \| 173 \| (76, 188) \| 225 \| Endopeptidase (Lysis protein) from bacteriophage origin \| Endopeptidase (Lysis protein) from bacteriophage origin; uniclust \| UniRef100\_UPI00034B3AB5 \| 97.7 \| 7.5e-07 \| 1.4e-12 \| 70.5 \| 102 \| (2, 103) \| 173 \| (1, 106) \| 197 \| lysis system i-spanin subunit Rz \| lysis system i-spanin subunit Rz; uniclust \| UniRef100\_A0A8S5NLS2 \| 97.7 \| 8.9e-07 \| 1.7e-12 \| 69.5 \| 94 \| (7, 104) \| 173 \| (5, 98) \| 170 \| Uncharacterized protein \| Uncharacterized protein; uniclust \| UniRef100\_A0A6G6IV19 \| 97.7 \| 1.1e-06 \| 2e-12 \| 56.4 \| 55 \| (2, 56) \| 173 \| (1, 55) \| 61 \| Uncharacterized protein \| Uncharacterized protein; uniclust \| UniRef100\_A0A0P9PJB1 \| 97.7 \| 1.1e-06 \| 2.1e-12 \| 53.9 \| 44 \| (130, 173) \| 173 \| (1, 44) \| 47 \| Uncharacterized protein \| Uncharacterized protein; uniclust \| UniRef100\_UPI000A747C3A \| 97.7 \| 1.3e-06 \| 2.4e-12 \| 58.3 \| 48 \| (121, 168) \| 173 \| (9, 56) \| 72 \| DUF2514 family protein \| DUF2514 family protein; uniclust \| UniRef100\_UPI0012FD249F \| 97.6 \| 1.5e-06 \| 2.8e-12 \| 57.8 \| 70 \| (36, 105) \| 173 \| (2, 71) \| 73 \| DUF2514 family protein \| DUF2514 family protein; uniclust \| UniRef100\_A0A270P9R3 \| 97.6 \| 1.5e-06 \| 2.8e-12 \| 60.6 \| 48 \| (121, 168) \| 173 \| (48, 95) \| 96 \| DUF2514 domain-containing protein \| DUF2514 domain-containing protein; uniclust \| UniRef100\_A0A9E0KB07 \| 97.6 \| 1.6e-06 \| 2.9e-12 \| 63.3 \| 101 \| (57, 165) \| 173 \| (10, 110) \| 113 \| Uncharacterized protein \| Uncharacterized protein; uniclust \| UniRef100\_UPI001E39CCA4 \| 97.6 \| 1.6e-06 \| 2.9e-12 \| 58.3 \| 51 \| (119, 169) \| 173 \| (24, 74) \| 76 \| DUF2514 family protein \| DUF2514 family protein; uniclust \| UniRef100\_A0A965PEI7 \| 97.6 \| 1.7e-06 \| 3.2e-12 \| 66.0 \| 102 \| (56, 168) \| 173 \| (47, 148) \| 150 \| Uncharacterized protein \| Uncharacterized protein; uniclust \| UniRef100\_UPI0021147FA3 \| 97.6 \| 1.9e-06 \| 3.5e-12 \| 60.5 \| 72 \| (98, 169) \| 173 \| (24, 97) \| 99 \| DUF2514 family protein \| DUF2514 family protein; uniclust \| UniRef100\_UPI001F43A9C7 \| 97.6 \| 1.9e-06 \| 3.6e-12 \| 59.6 \| 56 \| (115, 170) \| 173 \| (31, 86) \| 87 \| DUF2514 family protein \| DUF2514 family protein; uniclust \| UniRef100\_A0A315BGF8 \| 97.6 \| 2e-06 \| 3.7e-12 \| 66.3 \| 122 \| (43, 172) \| 173 \| (48, 175) \| 180 \| Lysozyme \| Lysozyme; uniclust \| UniRef100\_A0A0D0IWK6 \| 97.6 \| 1.9e-06 \| 3.8e-12 \| 70.9 \| 132 \| (25, 168) \| 173 \| (55, 190) \| 213 \| Lysozyme \| Lysozyme; uniclust \| UniRef100\_A0A0W7YSA5 \| 97.6 \| 1.9e-06 \| 3.8e-12 \| 70.2 \| 148 \| (5, 164) \| 173 \| (24, 175) \| 200 \| Lysis protein \| Lysis protein; uniclust \| UniRef100\_A0A6M9YZG7 \| 97.6 \| 2.1e-06 \| 3.9e-12 \| 65.7 \| 133 \| (35, 168) \| 173 \| (36, 169) \| 170 \| DUF2514 domain-containing protein \| DUF2514 domain-containing protein; uniclust \| UniRef100\_A0A088F9X7 \| 97.6 \| 1.9e-06 \| 4e-12 \| 69.2 \| 130 \| (15, 163) \| 173 \| (30, 159) \| 177 \| Phage lambda Rz-like lysis protein \| Phage lambda Rz-like lysis protein; uniclust \| UniRef100\_A0A2K4KXJ6 \| 97.6 \| 2.5e-06 \| 4.6e-12 \| 55.7 \| 44 \| (126, 169) \| 173 \| (22, 65) \| 66 \| DUF2514 domain-containing protein \| DUF2514 domain-containing protein; uniclust \| UniRef100\_A0A918U7Z6 \| 97.5 \| 2.5e-06 \| 4.9e-12 \| 67.5 \| 107 \| (58, 170) \| 173 \| (61, 167) \| 178 \| Uncharacterized protein \| Uncharacterized protein; uniclust \| UniRef100\_A0A031IWR0 \| 97.5 \| 2.7e-06 \| 5.8e-12 \| 70.1 \| 118 \| (44, 169) \| 173 \| (62, 181) \| 200 \| Prophage PSPPH06, lysis protein \| Prophage PSPPH06, lysis protein; uniclust \| UniRef100\_A0A068R0F6 \| 97.5 \| 2.7e-06 \| 6e-12 \| 70.7 \| 108 \| (54, 170) \| 173 \| (86, 194) \| 206 \| Putative Rac prophage prophage lambda endopeptidase \| Putative Rac prophage prophage lambda endopeptidase; uniclust \| UniRef100\_UPI001F2FD331 \| 97.5 \| 3.3e-06 \| 6.1e-12 \| 60.0 \| 95 \| (72, 169) \| 173 \| (8, 103) \| 105 \| DUF2514 family protein \| DUF2514 family protein; uniclust \| UniRef100\_A0A7Z2JFM2 \| 97.5 \| 3.4e-06 \| 6.3e-12 \| 56.9 \| 45 \| (126, 170) \| 173 \| (29, 73) \| 78 \| DUF2514 family protein \| DUF2514 family protein; uniclust \| UniRef100\_A0A5C8IZU8 \| 97.5 \| 3.5e-06 \| 6.4e-12 \| 64.9 \| 155 \| (2, 165) \| 173 \| (6, 174) \| 176 \| DUF2514 family protein \| DUF2514 family protein; uniclust \| UniRef100\_A0A8S5QU50 \| 97.5 \| 3.4e-06 \| 6.7e-12 \| 66.1 \| 138 \| (5, 155) \| 173 \| (15, 152) \| 163 \| Uncharacterized protein \| Uncharacterized protein; uniclust \| UniRef100\_UPI001D1909C2 \| 97.5 \| 3.7e-06 \| 6.9e-12 \| 59.3 \| 51 \| (120, 170) \| 173 \| (48, 98) \| 100 \| DUF2514 family protein \| DUF2514 family protein; uniclust \| UniRef100\_A0A067Y1K9 \| 97.5 \| 3.6e-06 \| 7.7e-12 \| 68.3 \| 74 \| (26, 99) \| 173 \| (25, 98) \| 180 \| Rz/RzI spanin protein \| Rz/RzI spanin protein; uniclust \| UniRef100\_A0A0B6S1Z9 \| 97.4 \| 4.9e-06 \| 1e-11 \| 70.2 \| 115 \| (49, 170) \| 173 \| (96, 218) \| 245 \| Putative bacteriophage lysis protein \| Putative bacteriophage lysis protein; uniclust \| UniRef100\_A0A1H9E2Y4 \| 97.4 \| 5.8e-06 \| 1.1e-11 \| 62.7 \| 90 \| (77, 168) \| 173 \| (62, 151) \| 157 \| Bacteriophage Rz lysis protein \| Bacteriophage Rz lysis protein; uniclust \| UniRef100\_A0A0S2F7F8 \| 97.4 \| 6.6e-06 \| 1.4e-11 \| 66.8 \| 135 \| (24, 165) \| 173 \| (28, 163) \| 175 \| Putative gp12 \| Putative gp12; uniclust \| UniRef100\_A0A0J6C284 \| 97.3 \| 9e-06 \| 1.8e-11 \| 64.5 \| 130 \| (16, 155) \| 173 \| (23, 152) \| 174 \| Uncharacterized protein \| Uncharacterized protein; uniclust \| UniRef100\_UPI001116F226 \| 97.3 \| 1.1e-05 \| 1.9e-11 \| 55.9 \| 74 \| (4, 81) \| 173 \| (7, 80) \| 88 \| DUF2514 family protein \| DUF2514 family protein; uniclust \| UniRef100\_UPI001E5BBD03 \| 97.3 \| 1.1e-05 \| 2e-11 \| 59.0 \| 104 \| (55, 166) \| 173 \| (8, 115) \| 118 \| lysis system i-spanin subunit Rz \| lysis system i-spanin subunit Rz; uniclust \| UniRef100\_A0A3N8D9L6 \| 97.3 \| 1.1e-05 \| 2e-11 \| 55.7 \| 81 \| (58, 144) \| 173 \| (6, 86) \| 87 \| DUF2514 family protein \| DUF2514 family protein; uniclust \| UniRef100\_A0A023KKM3 \| 97.3 \| 1.1e-05 \| 2.3e-11 \| 67.5 \| 54 \| (50, 110) \| 173 \| (68, 121) \| 213 \| Lysis protein \| Lysis protein; uniclust \| UniRef100\_A0A0A1IVS7 \| 97.3 \| 1.2e-05 \| 2.4e-11 \| 64.3 \| 157 \| (2, 163) \| 173 \| (1, 163) \| 180 \| Uncharacterized protein \| Uncharacterized protein; uniclust \| UniRef100\_A0A370FC04 \| 97.3 \| 1.4e-05 \| 2.6e-11 \| 62.1 \| 151 \| (14, 171) \| 173 \| (13, 167) \| 181 \| Bacteriophage Rz lysis protein \| Bacteriophage Rz lysis protein; uniclust \| UniRef100\_UPI0009304A3F \| 97.2 \| 1.8e-05 \| 3.3e-11 \| 57.5 \| 78 \| (93, 170) \| 173 \| (32, 111) \| 116 \| DUF2514 family protein \| DUF2514 family protein; uniclust \| UniRef100\_A0A2Z3IMC6 \| 97.2 \| 1.8e-05 \| 3.4e-11 \| 63.3 \| 152 \| (13, 169) \| 173 \| (54, 209) \| 222 \| Lysozyme \| Lysozyme; uniclust \| UniRef100\_UPI00207BF91A \| 97.2 \| 2e-05 \| 3.7e-11 \| 49.2 \| 45 \| (129, 173) \| 173 \| (5, 49) \| 50 \| DUF2514 family protein \| DUF2514 family protein; uniclust \| UniRef100\_A0A965PHN2 \| 97.2 \| 2.1e-05 \| 3.8e-11 \| 57.5 \| 98 \| (1, 103) \| 173 \| (1, 98) \| 120 \| Uncharacterized protein (Fragment) \| Uncharacterized protein (Fragment); uniclust \| UniRef100\_UPI00195B7822 \| 97.2 \| 2.2e-05 \| 4e-11 \| 55.3 \| 91 \| (54, 144) \| 173 \| (2, 94) \| 96 \| DUF2514 family protein \| DUF2514 family protein; uniclust \| UniRef100\_A0A212IIM0 \| 97.2 \| 2.3e-05 \| 4.3e-11 \| 50.8 \| 43 \| (128, 170) \| 173 \| (4, 46) \| 57 \| DUF2514 domain-containing protein \| DUF2514 domain-containing protein; uniclust \| UniRef100\_UPI0021AB3194 \| 97.1 \| 2.9e-05 \| 5.3e-11 \| 56.3 \| 83 \| (56, 144) \| 173 \| (30, 112) \| 113 \| DUF2514 family protein \| DUF2514 family protein; uniclust \| UniRef100\_UPI001E3A12C5 \| 97.1 \| 2.9e-05 \| 5.4e-11 \| 54.0 \| 67 \| (34, 100) \| 173 \| (3, 70) \| 82 \| DUF2514 domain-containing protein \| DUF2514 domain-containing protein; uniclust \| UniRef100\_A0A1I6GE33 \| 97.1 \| 3.1e-05 \| 5.8e-11 \| 59.3 \| 102 \| (58, 169) \| 173 \| (47, 148) \| 153 \| Bacteriophage Rz lysis protein \| Bacteriophage Rz lysis protein; uniclust \| UniRef100\_A0A1G6NB34 \| 97.1 \| 2.9e-05 \| 6e-11 \| 62.9 \| 137 \| (13, 165) \| 173 \| (17, 160) \| 172 \| Bacteriophage Rz lysis protein \| Bacteriophage Rz lysis protein; uniclust \| UniRef100\_A0A084YZU5 \| 97.1 \| 3.3e-05 \| 6.6e-11 \| 60.2 \| 75 \| (59, 140) \| 173 \| (56, 135) \| 143 \| Phage lysis protein (Fragment) \| Phage lysis protein (Fragment); uniclust \| UniRef100\_A0A0N9ER84 \| 97.1 \| 3.7e-05 \| 7e-11 \| 60.8 \| 36 \| (131, 166) \| 173 \| (140, 175) \| 189 \| Putative Rz lysis protein \| Putative Rz lysis protein; uniclust \| UniRef100\_A0A1H8TC25 \| 97.1 \| 3.6e-05 \| 7.2e-11 \| 64.0 \| 159 \| (5, 167) \| 173 \| (7, 169) \| 214 \| Bacteriophage Rz lysis protein \| Bacteriophage Rz lysis protein; uniclust \| UniRef100\_UPI001F11D6A6 \| 97.1 \| 4.1e-05 \| 7.5e-11 \| 50.3 \| 48 \| (122, 169) \| 173 \| (5, 52) \| 64 \| DUF2514 family protein \| DUF2514 family protein; uniclust \| UniRef100\_A0A0Q6W2Z3 \| 97.0 \| 4.3e-05 \| 7.9e-11 \| 59.1 \| 100 \| (69, 169) \| 173 \| (70, 169) \| 170 \| Lysozyme \| Lysozyme; uniclust \| UniRef100\_A0A965RZC8 \| 97.0 \| 5e-05 \| 9.2e-11 \| 55.7 \| 87 \| (17, 110) \| 173 \| (11, 97) \| 120 \| DUF2514 family protein (Fragment) \| DUF2514 family protein (Fragment); uniclust \| UniRef100\_A0A3S4G381 \| 97.0 \| 5.3e-05 \| 1e-10 \| 60.5 \| 60 \| (20, 79) \| 173 \| (29, 88) \| 184 \| Phage-tail assembly protein \| Phage-tail assembly protein; uniclust \| UniRef100\_A0A137YJG3 \| 97.0 \| 6.3e-05 \| 1.2e-10 \| 59.6 \| 106 \| (58, 171) \| 173 \| (74, 179) \| 182 \| DUF2514 family protein \| DUF2514 family protein; uniclust \| UniRef100\_UPI00057A4F47 \| 96.9 \| 7.2e-05 \| 1.3e-10 \| 52.4 \| 67 \| (3, 69) \| 173 \| (10, 84) \| 85 \| hypothetical protein \| hypothetical protein; uniclust \| UniRef100\_UPI0020C8520B \| 96.9 \| 7.5e-05 \| 1.4e-10 \| 53.3 \| 59 \| (114, 172) \| 173 \| (27, 85) \| 100 \| DUF2514 family protein \| DUF2514 family protein; uniclust \| UniRef100\_UPI001E381099 \| 96.9 \| 8.1e-05 \| 1.5e-10 \| 52.8 \| 72 \| (39, 110) \| 173 \| (6, 77) \| 97 \| DUF2514 family protein \| DUF2514 family protein; uniclust \| UniRef100\_A0A9E0Q5U7 \| 96.9 \| 8.2e-05 \| 1.5e-10 \| 58.9 \| 119 \| (45, 170) \| 173 \| (58, 177) \| 191 \| Uncharacterized protein \| Uncharacterized protein; uniclust \| UniRef100\_A0A3A1YQA2 \| 96.9 \| 9e-05 \| 1.6e-10 \| 52.7 \| 90 \| (47, 138) \| 173 \| (7, 96) \| 98 \| Uncharacterized protein \| Uncharacterized protein; uniclust \| UniRef100\_UPI0021F33334 \| 96.9 \| 9.5e-05 \| 1.7e-10 \| 49.9 \| 41 \| (130, 170) \| 173 \| (1, 41) \| 73 \| DUF2514 domain-containing protein \| DUF2514 domain-containing protein; uniclust \| UniRef100\_A0A198GPX5 \| 96.9 \| 8.4e-05 \| 1.7e-10 \| 60.9 \| 106 \| (55, 168) \| 173 \| (55, 162) \| 181 \| Endopeptidase \| Endopeptidase; uniclust \| UniRef100\_UPI0021BECF36 \| 96.9 \| 0.00011 \| 1.9e-10 \| 47.5 \| 34 \| (138, 171) \| 173 \| (12, 45) \| 57 \| DUF2514 domain-containing protein \| DUF2514 domain-containing protein; uniclust \| UniRef100\_A0A0E3BUQ5 \| 96.9 \| 9.3e-05 \| 2e-10 \| 62.8 \| 99 \| (1, 106) \| 173 \| (1, 109) \| 223 \| Lysis protein \| Lysis protein; uniclust \| UniRef100\_A0A6J5NGR5 \| 96.8 \| 0.00011 \| 2.1e-10 \| 56.6 \| 153 \| (5, 166) \| 173 \| (1, 160) \| 163 \| Spanin, inner membrane subunit \| Spanin, inner membrane subunit; uniclust \| UniRef100\_UPI001F37191F \| 96.8 \| 0.00013 \| 2.4e-10 \| 49.2 \| 37 \| (134, 170) \| 173 \| (34, 70) \| 72 \| DUF2514 family protein \| DUF2514 family protein; uniclust \| UniRef100\_A0A2S1GSJ3 \| 96.8 \| 0.00014 \| 2.8e-10 \| 58.9 \| 136 \| (13, 159) \| 173 \| (17, 156) \| 176 \| Rz lysis protein \| Rz lysis protein; uniclust \| UniRef100\_UPI00037C877F \| 96.8 \| 0.00016 \| 2.9e-10 \| 56.3 \| 147 \| (15, 169) \| 173 \| (11, 161) \| 170 \| lysis system i-spanin subunit Rz \| lysis system i-spanin subunit Rz; uniclust \| UniRef100\_A0A4Q3MIQ8 \| 96.7 \| 0.00017 \| 3.1e-10 \| 54.9 \| 99 \| (55, 171) \| 173 \| (36, 134) \| 136 \| Uncharacterized protein \| Uncharacterized protein; uniclust \| UniRef100\_UPI0020A19267 \| 96.7 \| 0.00018 \| 3.3e-10 \| 48.0 \| 48 \| (123, 170) \| 173 \| (14, 61) \| 67 \| DUF2514 family protein \| DUF2514 family protein; uniclust \| UniRef100\_UPI001377749F \| 96.7 \| 0.00018 \| 3.3e-10 \| 46.3 \| 50 \| (23, 72) \| 173 \| (2, 51) \| 56 \| DUF2514 family protein \| DUF2514 family protein; uniclust \| UniRef100\_A0A0P9R5I7 \| 96.7 \| 0.00018 \| 3.4e-10 \| 53.4 \| 41 \| (131, 171) \| 173 \| (1, 41) \| 125 \| Uncharacterized protein \| Uncharacterized protein; uniclust \| UniRef100\_A0A0C1HF82 \| 96.7 \| 0.00016 \| 3.5e-10 \| 57.7 \| 67 \| (12, 85) \| 173 \| (18, 84) \| 144 \| Phage associated protein \| Phage associated protein; uniclust \| UniRef100\_A0A4Q7NCI6 \| 96.7 \| 0.00019 \| 3.5e-10 \| 56.2 \| 110 \| (57, 168) \| 173 \| (63, 176) \| 177 \| Uncharacterized protein DUF2514 \| Uncharacterized protein DUF2514; uniclust \| UniRef100\_UPI0022208F3E \| 96.7 \| 0.00021 \| 3.8e-10 \| 61.1 \| 102 \| (62, 170) \| 173 \| (238, 342) \| 348 \| lysis system i-spanin subunit Rz \| lysis system i-spanin subunit Rz; uniclust \| UniRef100\_A0A0A1H6K0 \| 96.7 \| 0.00019 \| 3.8e-10 \| 59.8 \| 73 \| (8, 84) \| 173 \| (16, 88) \| 199 \| Uncharacterized protein \| Uncharacterized protein; uniclust \| UniRef100\_A4EKQ5 \| 96.7 \| 0.00021 \| 3.9e-10 \| 54.5 \| 111 \| (51, 171) \| 173 \| (36, 146) \| 148 \| Uncharacterized protein \| Uncharacterized protein; uniclust \| UniRef100\_A0A1B2LZW7 \| 96.7 \| 0.00021 \| 3.9e-10 \| 51.6 \| 87 \| (71, 165) \| 173 \| (5, 91) \| 96 \| Uncharacterized protein \| Uncharacterized protein; uniclust \| UniRef100\_A0A0R0A6K2 \| 96.7 \| 0.00022 \| 4.5e-10 \| 59.6 \| 84 \| (62, 154) \| 173 \| (83, 171) \| 201 \| Lysozyme \| Lysozyme; uniclust \| UniRef100\_UPI001ED9A4F7 \| 96.6 \| 0.00026 \| 4.8e-10 \| 46.7 \| 59 \| (86, 144) \| 173 \| (3, 63) \| 63 \| DUF2514 family protein \| DUF2514 family protein; uniclust \| UniRef100\_A0A1E3ZLE2 \| 96.6 \| 0.00027 \| 5.2e-10 \| 57.1 \| 120 \| (47, 170) \| 173 \| (55, 174) \| 185 \| Lysozyme \| Lysozyme; uniclust \| UniRef100\_UPI001F078139 \| 96.6 \| 0.00028 \| 5.2e-10 \| 47.0 \| 53 \| (26, 78) \| 173 \| (13, 65) \| 66 \| DUF2514 family protein \| DUF2514 family protein; uniclust \| UniRef100\_UPI001E36A577 \| 96.6 \| 0.00031 \| 5.7e-10 \| 55.8 \| 91 \| (56, 153) \| 173 \| (82, 174) \| 193 \| DUF2514 family protein \| DUF2514 family protein; uniclust \| UniRef100\_A0A142JMY0 \| 96.6 \| 0.00032 \| 6.5e-10 \| 58.0 \| 105 \| (63, 171) \| 173 \| (69, 178) \| 193 \| Lysozyme \| Lysozyme; uniclust \| UniRef100\_A0A9E9KSP8 \| 96.6 \| 0.00035 \| 6.7e-10 \| 55.9 \| 89 \| (74, 169) \| 173 \| (76, 170) \| 172 \| Lysis system i-spanin subunit Rz \| Lysis system i-spanin subunit Rz; uniclust \| UniRef100\_UPI000CC96F41 \| 96.6 \| 0.00035 \| 6.8e-10 \| 59.6 \| 137 \| (25, 170) \| 173 \| (55, 192) \| 270 \| hypothetical protein \| hypothetical protein; uniclust \| UniRef100\_A0A2S1GTV9 \| 96.5 \| 0.00038 \| 7.2e-10 \| 55.7 \| 100 \| (70, 170) \| 173 \| (79, 178) \| 182 \| Rz lysis protein \| Rz lysis protein; uniclust \| UniRef100\_A0A519FV23 \| 96.5 \| 0.00038 \| 7.3e-10 \| 56.1 \| 86 \| (81, 170) \| 173 \| (83, 168) \| 187 \| DUF2514 family protein \| DUF2514 family protein; uniclust \| UniRef100\_UPI0001F06794 \| 96.5 \| 0.00041 \| 7.6e-10 \| 48.6 \| 81 \| (77, 157) \| 173 \| (5, 87) \| 87 \| DUF2514 family protein \| DUF2514 family protein; uniclust \| UniRef100\_UPI00145608D9 \| 96.5 \| 0.00042 \| 7.7e-10 \| 54.7 \| 116 \| (52, 172) \| 173 \| (54, 172) \| 181 \| lysis system i-spanin subunit Rz \| lysis system i-spanin subunit Rz; uniclust \| UniRef100\_A0A1A9RFJ3 \| 96.5 \| 0.00039 \| 8e-10 \| 54.6 \| 38 \| (1, 42) \| 173 \| (11, 48) \| 135 \| Phage associated protein \| Phage associated protein; uniclust \| UniRef100\_A0A2U3CZ75 \| 96.5 \| 0.00049 \| 8.9e-10 \| 47.0 \| 52 \| (120, 171) \| 173 \| (20, 71) \| 74 \| DUF2524 domain-containing protein \| DUF2524 domain-containing protein; uniclust \| UniRef100\_A0A0F5P9U3 \| 96.5 \| 0.00045 \| 8.9e-10 \| 55.4 \| 98 \| (1, 102) \| 173 \| (1, 102) \| 160 \| Uncharacterized protein \| Uncharacterized protein; uniclust \| UniRef100\_A0A0J6IL90 \| 96.5 \| 0.00045 \| 9e-10 \| 54.6 \| 96 \| (69, 171) \| 173 \| (26, 128) \| 142 \| Lysozyme \| Lysozyme; uniclust \| UniRef100\_A0A1D8IVC9 \| 96.5 \| 0.00047 \| 9.3e-10 \| 56.7 \| 104 \| (1, 110) \| 173 \| (4, 114) \| 186 \| DUF2514 family protein \| DUF2514 family protein; uniclust \| UniRef100\_UPI0018D42AA6 \| 96.5 \| 0.00052 \| 9.6e-10 \| 53.2 \| 74 \| (1, 85) \| 173 \| (46, 119) \| 159 \| hypothetical protein \| hypothetical protein; uniclust \| UniRef100\_UPI000AD9CA16 \| 96.4 \| 0.00057 \| 1e-09 \| 48.4 \| 76 \| (10, 85) \| 173 \| (7, 82) \| 91 \| DUF2514 family protein \| DUF2514 family protein; uniclust \| UniRef100\_A0A853F7S4 \| 96.4 \| 0.00057 \| 1e-09 \| 49.7 \| 86 \| (82, 169) \| 173 \| (17, 102) \| 106 \| DUF2514 family protein \| DUF2514 family protein; uniclust \| UniRef100\_A0A6I1VLR2 \| 96.4 \| 0.00059 \| 1.1e-09 \| 46.3 \| 51 \| (119, 169) \| 173 \| (16, 66) \| 72 \| DUF2514 family protein \| DUF2514 family protein; uniclust \| UniRef100\_A0A6L6TIH4 \| 96.4 \| 0.00059 \| 1.1e-09 \| 44.6 \| 40 \| (129, 168) \| 173 \| (19, 58) \| 59 \| DUF2514 family protein \| DUF2514 family protein; uniclust \| UniRef100\_A0A2D0K6M7 \| 96.4 \| 0.00054 \| 1.1e-09 \| 53.0 \| 49 \| (55, 110) \| 173 \| (46, 94) \| 123 \| Peptidase \| Peptidase; uniclust \| UniRef100\_A0A923JVT3 \| 96.4 \| 0.00061 \| 1.1e-09 \| 38.7 \| 27 \| (144, 170) \| 173 \| (2, 28) \| 30 \| DUF2514 domain-containing protein \| DUF2514 domain-containing protein; uniclust \| UniRef100\_UPI0022FEA43E \| 96.4 \| 0.00061 \| 1.1e-09 \| 46.8 \| 50 \| (20, 69) \| 173 \| (18, 67) \| 73 \| DUF2514 family protein \| DUF2514 family protein; uniclust \| UniRef100\_A0A7G8VCI8 \| 96.4 \| 0.00064 \| 1.2e-09 \| 47.7 \| 52 \| (119, 170) \| 173 \| (22, 73) \| 86 \| DUF2514 family protein \| DUF2514 family protein; uniclust \| UniRef100\_A0A3N8D384 \| 96.4 \| 0.00064 \| 1.2e-09 \| 54.5 \| 96 \| (69, 169) \| 173 \| (63, 161) \| 171 \| Lysis protein \| Lysis protein; uniclust \| UniRef100\_A0A2H4JG77 \| 96.4 \| 0.00061 \| 1.2e-09 \| 53.7 \| 102 \| (2, 110) \| 173 \| (1, 109) \| 139 \| Lysis protein (Fragment) \| Lysis protein (Fragment); uniclust \| UniRef100\_A0A0E3BVE5 \| 96.4 \| 0.00067 \| 1.2e-09 \| 48.8 \| 76 \| (93, 170) \| 173 \| (14, 90) \| 99 \| Lysozyme \| Lysozyme; uniclust \| UniRef100\_A0A0E2NG58 \| 96.4 \| 0.00064 \| 1.3e-09 \| 53.0 \| 36 \| (14, 49) \| 173 \| (22, 57) \| 129 \| Uncharacterized protein \| Uncharacterized protein; uniclust \| UniRef100\_A0A378XQ82 \| 96.4 \| 0.0007 \| 1.3e-09 \| 53.5 \| 115 \| (37, 157) \| 173 \| (40, 159) \| 180 \| Protein of uncharacterized function (DUF2514) \| Protein of uncharacterized function (DUF2514); uniclust \| UniRef100\_A0A1X3DHC2 \| 96.4 \| 0.00073 \| 1.3e-09 \| 53.0 \| 93 \| (9, 110) \| 173 \| (2, 94) \| 164 \| Lysozyme \| Lysozyme; uniclust \| UniRef100\_A0A077NBU8 \| 96.4 \| 0.00067 \| 1.4e-09 \| 55.8 \| 50 \| (54, 110) \| 173 \| (53, 102) \| 174 \| Putative Rac prophage prophage lambda endopeptidase \| Putative Rac prophage prophage lambda endopeptidase; uniclust \| UniRef100\_A0A022PKZ8 \| 96.3 \| 0.00076 \| 1.5e-09 \| 57.8 \| 29 \| (6, 34) \| 173 \| (43, 71) \| 250 \| Lysis protein \| Lysis protein; uniclust \| UniRef100\_UPI000442563A \| 96.3 \| 0.00089 \| 1.6e-09 \| 43.9 \| 38 \| (14, 51) \| 173 \| (10, 47) \| 59 \| DUF2514 family protein \| DUF2514 family protein; uniclust \| UniRef100\_A0A074TS68 \| 96.3 \| 0.00084 \| 1.7e-09 \| 50.1 \| 32 \| (1, 32) \| 173 \| (2, 33) \| 102 \| Uncharacterized protein \| Uncharacterized protein; uniclust \| UniRef100\_A0A0J8FPI0 \| 96.3 \| 0.00092 \| 1.8e-09 \| 53.8 \| 125 \| (25, 165) \| 173 \| (21, 147) \| 159 \| Lysozyme \| Lysozyme; uniclust \| UniRef100\_A0A543AK70 \| 96.3 \| 0.001 \| 1.8e-09 \| 48.7 \| 47 \| (121, 167) \| 173 \| (43, 89) \| 108 \| Uncharacterized protein DUF2514 \| Uncharacterized protein DUF2514; uniclust \| UniRef100\_A0A923GH54 \| 96.3 \| 0.001 \| 1.9e-09 \| 48.8 \| 62 \| (15, 76) \| 173 \| (6, 67) \| 110 \| DUF2514 family protein \| DUF2514 family protein |
| Top keywords  (threshold 1.00e-03 (evalue)) | **DUF2514, lysis, domain\_containing, Fragment, Lysozyme, Rz, Putative, Bacteriophage, Endopeptidase, prophage** |
| Output files | ../../similar\_sequences/65\_FANPEZAQ\_CDS\_0065\_merged.svg ../../similar\_sequences/65\_FANPEZAQ\_CDS\_0065\_pdb70.a3m ../../similar\_sequences/65\_FANPEZAQ\_CDS\_0065\_pdb70.hhr ../../similar\_sequences/65\_FANPEZAQ\_CDS\_0065\_uniclust.a3m ../../similar\_sequences/65\_FANPEZAQ\_CDS\_0065\_uniclust.hhr |

#### Structure prediction (AlphaFold)^2^

|  |  |
| --- | --- |
| Stats | xml version="1.0" encoding="utf-8" standalone="no"?       2024-09-02T21:09:59.965739 image/svg+xml   Matplotlib v3.7.2, https://matplotlib.org/ |
| Predicted structure | **NGL Viewer Controls:**  - Center: *Left-Click* - Rotate: *Left-Click + Drag* - Translate: *Right-Click + Drag* - Zoom: *Shift + Left-Click + Drag* |
| Output files | ../../predicted\_structures/65\_FANPEZAQ\_CDS\_0065/features.pkl ../../predicted\_structures/65\_FANPEZAQ\_CDS\_0065/ranked\_0.pdb ../../predicted\_structures/65\_FANPEZAQ\_CDS\_0065/ranked\_0\_plots.svg ../../predicted\_structures/65\_FANPEZAQ\_CDS\_0065/result\_model\_1\_ptm\_pred\_0.pkl |

#### Structure similarity search results (Foldseek)^3^

|  |  |
| --- | --- |
| Structure databases searched | Pdb, Afdb-proteome, Afdb-uniprot50 |
| Results, scheme(s)  (Top layers only, threshold 1.00e-02 (evalue)) | xml version="1.0" encoding="utf-8" standalone="no"?       2024-09-02T21:11:34.177487 image/svg+xml   Matplotlib v3.7.2, https://matplotlib.org/ |
| Results, table  (threshold 1.00e-02 (evalue)) | db \| id \| prob \| evalue \| bits \| fident \| alnlen \| mismatch \| gapopen \| qstart \| qend \| tstart \| tend \| name \| description; afdb-uniprot50 \| AF-A0A423A0K5-F1-MODEL\_V4 \| 1.0 \| 8.468e-11 \| 391 \| 0.687 \| 173 \| 53 \| 1 \| 1 \| 172 \| 1 \| 173 \| DUF2514 family protein \| DUF2514 family protein; afdb-uniprot50 \| AF-A0A7H9F4S6-F1-MODEL\_V4 \| 1.0 \| 0.001541 \| 188 \| 0.343 \| 166 \| 109 \| 0 \| 2 \| 167 \| 10 \| 175 \| DUF2514 family protein \| DUF2514 family protein; afdb-uniprot50 \| AF-A0A0A7KTN9-F1-MODEL\_V4 \| 1.0 \| 0.001139 \| 184 \| 0.343 \| 166 \| 107 \| 1 \| 10 \| 173 \| 223 \| 388 \| Lysozyme \| Lysozyme |
| Top keywords  (threshold 1.00e-02 (evalue)) | **DUF2514, Lysozyme** |
| Output files | ../../similar\_structures/65\_FANPEZAQ\_CDS\_0065\_afdb-proteome\_foldseek.tsv ../../similar\_structures/65\_FANPEZAQ\_CDS\_0065\_afdb-uniprot50\_foldseek.tsv ../../similar\_structures/65\_FANPEZAQ\_CDS\_0065\_merged.svg ../../similar\_structures/65\_FANPEZAQ\_CDS\_0065\_pdb\_foldseek.tsv |

  
  
  

Return to summary | Go to previous | Go to next

  


---

**Sequence/structure alignments coloring**  
Each object in the alignment figures is colored according to its E-value following this color coding:

1e-100
10
