## Supplementary material for "Completing the BASEL phage collection to unlock hidden diversity for systematic exploration of phage-host interactions": Data S2: index.html

Summary


### Escherichia virus HeidiAbel

Report creation date: 02 Sep 2024, 12:00
  
Project folder: n/a
  
Input sequences file: Escherichia\_virus\_HeidiAbel.gb

**More information (click to expand)**

**Table legend:**  
Entry name = Name of the protein sequence imported from the input file. Links to entry’s highlight page.Click on the entry's name to see detailed information.  
DA = ‘Domain Architecture’. The displayed value indicates the best score obtained by the search algorithm (HHblits/HH-suite3)^1^. The cell is colored in green if the value is better than the treshold 1.00e-03  
IP = ‘Identical Proteins’. Value indicates how many proteins identical to the entry were found in sequence/structure databases. The cell is green if the value is greater than 0.  
SeS = ‘Sequence Similarity’. Value indicates the best score obtained by the search algorithm (HHblits/HH-suite3)^1^. The cell is green if the value is better than the threshold 1.00e-03  
StP = ‘Structure Prediction’. Value indicates the confidence score (= average pLDDT) obtained by the structure prediction algorithm (AlphaFold)^2^. The cell is green if the value is greater than 70 %.  
StS = ‘Structure Similarity’. Value indicates the best score obtained by the search algorithm (FoldSeek)^3^. The cell is green if the value is better than the threshold 1.00e-02  
Keywords = Major keywords retrieved from the analysis of DA, IP, SeS, StP and StS results. Annotations imported from the input file are not considered.  
Click on "Toggle Content" to display more information.

**Sequence/structure alignments coloring**  
Each object in the alignment figures is colored according to its E-value following this color coding:

1e-100
10

**Contact/Questions:**  

  
  

| # | Entry name | DA | IP | SeS | StP | StS | Keywords Toggle Content |
| --- | --- | --- | --- | --- | --- | --- | --- |
| 1 | FANPEZAQ\_CDS\_0001 Locus: [81:525](+) Seq: MWRKDGCPVVTMGGRGKE... Seq Len: 147 aa MW: 16.4 kDa | 2.1e-24 | 0 | 1.7e-28 | 82.0 | 4.4e-12 | terminase, small, packaging, dna, nu1, phage, hypothetical, domain\_containing, fragment, helix\_turn\_helix, duf1441, hth, merr\_type, dna\_packaging, elements, external, origin, putative, dna\_binding, transcriptional **DA** \| packaging, terminase, small, head, and, DUF1441, Phage, DNA, Nu1, p69558, VI\_02299, p277333, VI\_02593, p302384, VI\_08027, p410932, VI\_07810, p435585, VI\_08997, p386363; **IP** \|; **SeS** \| Terminase, small, DNA, packaging, hypothetical, Phage, Nu1, domain\_containing, Fragment, Helix\_turn\_helix, HTH, merR\_type, DUF1441, Elements, external, origin, DNA\_packaging, Putative, transcriptional, regulator; **StS** \| Terminase, Nu1, small, DNA, packaging, Phage, DNA\_packaging, Protoporphyrinogen, oxidase, homolog, Helix\_turn\_helix, domain\_containing, Putative, gpNu1, Excisionase, DNA\_binding, Bll5240 |
| 2 | FANPEZAQ\_CDS\_0002 Locus: [484:2533](+) Seq: MAMTKGATVTAKNNPFAN... Seq Len: 682 aa MW: 76.7 kDa | 1.0e-128 | 0 | 9.0e-146 | 82.8 | 4.1e-94 | terminase, large, phage, helicase, gpa, rna, dna, fragment, a, atp\_dependent, domain\_containing, atpase, in, c, complex, e, and, yes, adp, tail **DA** \| helicase, a, in, yes, RNA, and, Engineered, the, DNA, i, escherichia, coli, ATP\_dependent, Alpha, containing, Organism\_taxid, Beta, Expressed, nucleotide, ARCH; **IP** \|; **SeS** \| terminase, large, Phage, helicase, RNA, GpA, Fragment, ATPase, domain\_containing, DNA, complex, C, ADP, E, ATP\_dependent, HYDROLASE, endonuclease, factor, splicing, DEAD\_box; **StS** \| terminase, Phage, large, GpA, tail, GpA\_like, assembly, Bacteriophage, DNA, Putative, packaging, domain\_containing, helicase, prophage, Gene, Crossover, junction, endodeoxyribonuclease, RuvC, Primosome |
| 3 | FANPEZAQ\_CDS\_0003 Locus: [2539:3079](+) Seq: MKCSQLPQKITAGLTFES... Seq Len: 179 aa MW: 20.0 kDa | 2.4e-42 | 0 | 1.8e-51 | 81.0 | 1.6e-22 | domain\_containing, bppu\_n, light, phage, receptor, fab, duf2479, hypothetical, bppu, dolichyl\_diphosphooligosaccharide\_\_protein, glycosyltransferase, n\_terminal, immunoglobulin, fibronectin, motile, sperm, tail, type, repeat, and **DA** \| head\_tail, connector, joining, NC\_009016\_p5, NC\_024367\_p52, adaptor, p69560, VI\_02299, p130612, VI\_06329, p166236, VI\_12298; **IP** \|; **SeS** \| domain\_containing, hypothetical, BppU, N\_terminal, Phage, tail, Fragment, Peptidylprolyl, isomerase, Primosomal, replication, PriB, PriC, DUF6148, GpW, associated, Putative, DUF1254, portal, Preprotein; **StS** \| domain\_containing, BppU\_N, light, receptor, Fab, DUF2479, Dolichyl\_diphosphooligosaccharide\_\_protein, glycosyltransferase, Phage, immunoglobulin, Fibronectin, Motile, sperm, type, repeat, and, Ig\_like, antibody, heavy, cell |
| 4 | FANPEZAQ\_CDS\_0004 Locus: [3036:4635](+) Seq: MQSIRAQSTGEPSLMKLW... Seq Len: 532 aa MW: 60.1 kDa | 3.1e-59 | 0 | 4.0e-104 | 82.5 | 1.9e-69 | portal, phage, lambda, fragment, capsid, putative, duf935, head, prophage, and, packaging, domain\_containing, bacteriophage, minor, terminase, head\_tail, phage\_related, duf4055, spp1, preconnector **DA** \| portal, head, and, packaging, Phage, DUF935, lambda, DUF1073, Bacteriophage, T4\_like, Gp20, SPP1, Gp6\_like, p173499, VI\_06341, NC\_023612\_p3, p353614, VI\_06430, p213127, VI\_06558; **IP** \|; **SeS** \| portal, Phage, Fragment, lambda, Capsid, Putative, prophage, terminase, Minor, domain\_containing, ParB, Head\_tail, preconnector, protease, Plasmid, partitioning, Bacteriophage\_related, large, ATP\_dependent, Clp; **StS** \| portal, Phage, lambda, DUF935, Putative, Capsid, prophage, domain\_containing, Bacteriophage, Phage\_related, DUF4055, HK97, Phage\_Mu\_F, Rcc01684, UPPER, COLLAR, gp20, connector, tail, Gene |
| 5 | FANPEZAQ\_CDS\_0005 Locus: [4612:6661](+) Seq: MQKNPLNSIIGAEFTRKS... Seq Len: 682 aa MW: 73.9 kDa | 9.7e-57 | 0 | 2.5e-93 | 70.6 | 4.0e-91 | capsid, phage, major, hk97, protease, prohead, head, peptidase, fragment, putative, u35, caudovirus, bacteriophage, and, packaging, domain\_containing, virus, gp36, gpt, phage\_related **DA** \| head, and, packaging, major, protease, maturation, capsid, Phage, gp5, serine, bacteriophage, in, Caudovirus, prohead, Putative, XkdF, Lactococcus, lactis, P2, DUF4043; **IP** \|; **SeS** \| Phage, capsid, major, HK97, prohead, protease, Fragment, Peptidase, U35, Bacteriophage, VIRUS, Caudovirus, Putative, GpT, domain\_containing, Mu, head, Phage\_related, phi\_C31, gp36; **StS** \| capsid, major, Phage, HK97, prohead, protease, Putative, Peptidase, head, Caudovirus, gp36, U35, Phage\_related, domain\_containing, phi\_C31, capsid\_like, Mu\_like\_gpT, prophage, Maritimacin, TIGR01554 |
| 6 | FANPEZAQ\_CDS\_0006 Locus: [6712:7015](+) Seq: MSGTYVLKLTSAIAISGE... Seq Len: 100 aa MW: 10.8 kDa | 1.7e-21 | 0 | 2.9e-19 | 74.8 | 6.2e-10 | domain\_containing, rho, termination, factor, n\_terminal, heh, lem, fragment, transcription, hypothetical, sap, phage, rho\_n, ribosomal, 50s, endonuclease, vii, mu\_like, prophage, flumu **DA** \| and, A, Transcription, in, endonuclease, VII, Alpha, yes, Termination, Factor, Rho, Rna\_binding, T4, recombination, LEM\_like, LAP2, isoform, gp49, Engineered, Organism\_taxid; **IP** \|; **SeS** \| Rho, termination, domain\_containing, factor, N\_terminal, HeH, LEM, Fragment, hypothetical, Transcription, Phage, SAP, ribosomal, 50S, L21, Mu\_like, prophage, FluMu, Tail, factor\_like; **StS** \| domain\_containing, Rho\_N, Rho, termination, factor, N\_terminal, SAP, factor\_like, containing, Contig\_80, whole, genome, shotgun, sequence, Helix\_hairpin\_helix, Putative, membrane, ycf1 |
| 7 | FANPEZAQ\_CDS\_0007 Locus: [7004:7100](+) Seq: MTNNATRHICNGSGRASN... Seq Len: 31 aa MW: 3.6 kDa | 1.5e-01 | 0 | 1.5e-01 | 64.0 | -- | **DA** \|; **IP** \|; **SeS** \|; **StS** \| |
| 8 | FANPEZAQ\_CDS\_0008 Locus: [7103:8138](-) Seq: MKTSALSLMGINCSVIGR... Seq Len: 344 aa MW: 37.9 kDa | 2.3e+00 | 0 | 7.9e+00 | 27.3 | 5.0e-02 | **DA** \|; **IP** \|; **SeS** \|; **StS** \| |
| 9 | FANPEZAQ\_CDS\_0009 Locus: [8106:8244](+) Seq: MPIKDKAEVFIEDEIFTQ... Seq Len: 45 aa MW: 5.6 kDa | 1.6e+00 | 0 | 3.0e-17 | 87.9 | -- | phage, hypothetical, tail, portal, fragment **DA** \|; **IP** \|; **SeS** \| Phage, hypothetical, tail, portal, Fragment; **StS** \| |
| 10 | FANPEZAQ\_CDS\_0010 Locus: [8243:8813](+) Seq: MNADTHIDLEVLHNAIVA... Seq Len: 189 aa MW: 21.3 kDa | 1.7e-39 | 0 | 1.8e-45 | 90.7 | 2.2e-23 | hypothetical, gp37, phage, duf1834, tail, prophage, domain\_containing, mu\_like, minor, u, fragment, putative, duf3168, ankyrin, phage\_related, duf4128, associated, gp37\_like, duf806, atp\_binding **DA** \| portal, head, and, packaging, p140225, VI\_01698, p377723, VI\_06234, p190981, VI\_07687, p352299, VI\_01044, tail, terminator, connector, p329161, VI\_04355, p338775, VI\_10901; **IP** \|; **SeS** \| hypothetical, Phage, DUF1834, Gp37, domain\_containing, Fragment, tail, Mu\_like, prophage, DUF3168, related, Ankyrin, Putative, phage\_like, Phage\_base\_V, Phage\_related, DUF4113, Bacteriophage, lambda, GpZ; **StS** \| Gp37, DUF1834, tail, Phage, prophage, Minor, U, Putative, Mu\_like, DUF4128, associated, Phage\_related, Gp37\_like, Hypothetical, DUF806, ATP\_binding, cytoplasmic, FluMu, Ankyrin, domain\_containing |
| 11 | FANPEZAQ\_CDS\_0011 Locus: [8809:9463](+) Seq: MSADVSELNRQLNNVVRI... Seq Len: 217 aa MW: 22.3 kDa | 2.8e-25 | 0 | 1.1e-45 | 93.4 | 2.7e-23 | baseplate, assembly, phage, v, type, secretion, domain\_containing, vi, system, vgr, ob\_fold, gp5, phage\_base\_v, vgrg, tip, phage\_related, fragment, rhs, gpv, p2 **DA** \| Beta, yes, Baseplate, a, Engineered, escherichia, coli, OB, Organism\_taxid, Expressed, in, Expression\_system\_taxid, ARCH, phage, assembly, v, Synonym, p2, Gene, Mainly; **IP** \|; **SeS** \| baseplate, assembly, Phage, V, Type, secretion, VI, domain\_containing, Vgr, system, OB\_fold, Gp5, Fragment, Rhs, Phage\_related, beta\_helix, Putative, Gp138, element, VgrG; **StS** \| baseplate, assembly, V, Phage, Phage\_base\_V, domain\_containing, Type, secretion, VI, system, VgrG, tip, gpV, Phage\_related, P2, Rhs, Putative, Vgr, element, in |
| 12 | FANPEZAQ\_CDS\_0012 Locus: [9516:9855](+) Seq: MKGTNAATGAPLDGMDHL... Seq Len: 112 aa MW: 12.1 kDa | 3.2e-27 | 0 | 2.1e-30 | 93.6 | 4.4e-18 | domain\_containing, gpw\_gp25, baseplate, gp25, gpw, assembly, irad, gp25\_like, phage, lysozyme, w, putative, 25\_like, duf2634, gene, wedge, secretion, type, fragment, vi **DA** \| baseplate, tail, wedge, A, Nuclear, Transport, Factor, gp25, Phage, lysozyme, Engineered, yes, Organism\_taxid, Gene, Expressed, in, escherichia, coli, Expression\_system\_taxid, Alpha; **IP** \|; **SeS** \| domain\_containing, IraD, Gp25\_like, Baseplate, GPW, gp25, assembly, Phage, Fragment, DUF2634, secretion, wedge, Type, lysozyme, VI, W, Putative, system, 25\_like, Gene; **StS** \| domain\_containing, GPW\_gp25, Baseplate, assembly, gp25, GPW, Phage, lysozyme, W, 25\_like, Putative, Gene, DUF2634, tail, wedge, Integrase, prophage, Type, VI, secretion |
| 13 | FANPEZAQ\_CDS\_0013 Locus: [9855:10758](+) Seq: MAGSYTVIDLSQLPAPTI... Seq Len: 300 aa MW: 31.8 kDa | 1.7e-36 | 0 | 3.9e-55 | 91.8 | 2.6e-43 | baseplate, domain\_containing, assembly, baseplate\_j, j\_like, j, gp47, phage, putative, phage\_related, tail, fragment, jaye, wedge, mu, bacteriophage, homolog, gp6, gpj, tube **DA** \| tail, baseplate, wedge, structural, head, and, packaging, virion, DUF276, J\_like, DUF2612, Phage, Tail\_P2\_I, KU686208\_p78, p428496, VI\_02892, p248291, VI\_09499, p256756, VI\_05872; **IP** \|; **SeS** \| Baseplate, J\_like, domain\_containing, J, gp47, assembly, Phage, Fragment, Putative, wedge, tail, gp6, Mu, T4, complex, baseplate\_tail, tube, pre\_attachment, Phage\_related, bacteriophage; **StS** \| Baseplate, assembly, domain\_containing, Baseplate\_J, J, gp47, Phage, J\_like, Phage\_related, Putative, JayE, tail, bacteriophage, Mu, homolog, GpJ, wedge, fiber, prophage, Probable |
| 14 | FANPEZAQ\_CDS\_0014 Locus: [10750:11368](+) Seq: MSDLLPPNATAQERALAN... Seq Len: 205 aa MW: 22.9 kDa | 4.3e-47 | 0 | 1.9e-54 | 82.7 | 4.1e-28 | tail, phage, i, p2, bacteriophage, formation, p2\_related, p2\_like, putative, fragment, domain\_containing, tail\_p2\_i, gpi, duf2313, baseplate, fiber, i\_like, j, prophage, assembly **DA** \| tail, VI\_08944, Phage, Tail\_P2\_I, Bacteriophage, Mu\_like, Gp48, DUF2612, p200917, VI\_08608, p214505, VI\_09983, p204625, VI\_11278, baseplate, p127197, VI\_01263, p52203, p284533, VI\_03834; **IP** \|; **SeS** \| tail, Phage, I, Fragment, P2, P2\_related, formation, Bacteriophage, domain\_containing, Putative, Tail\_P2\_I, P2\_like, GpI, fiber, baseplate, I\_like, J, tail\_like, assembly, J\_like; **StS** \| tail, Phage, I, P2, P2\_like, Bacteriophage, formation, Putative, P2\_related, Tail\_P2\_I, domain\_containing, GpI, DUF2313, prophage, baseplate, Phage\_related, assembly, fiber, J, I\_like |
| 15 | FANPEZAQ\_CDS\_0015 Locus: [11384:12656](+) Seq: MATYYTLLTKIGQARIAN... Seq Len: 423 aa MW: 45.0 kDa | 7.4e-53 | 0 | 1.2e-52 | 84.3 | 1.8e-31 | tail, phage, fiber, fibre, tail\_collar, domain\_containing, collar, putative, phage\_related, repeat, h, fragment, protein\_like, variable, bacteriophage, gph, repeat\_containing, probable, product, fiber\_like **DA** \| tail, fiber, collar, p300453, VI\_07590, p250465, VI\_02803, p390163, VI\_08186, p384764, VI\_09705, p13596, VI\_07483, p204627, VI\_11278, p306437, VI\_06250, p203938, VI\_11941, p394343; **IP** \|; **SeS** \| tail, Phage, fiber, fibre, tail\_collar, Fragment, domain\_containing, collar, Putative, Phage\_related, repeat, H, Variable, protein\_like, bacteriophage, repeat\_containing, adhesin, GpH, T7, Gp17; **StS** \| tail, Phage, fiber, fibre, tail\_collar, domain\_containing, Collar, Putative, Phage\_related, repeat, H, protein\_like, variable, GpH, repeat\_containing, bacteriophage, Peptidase, S74, Probable, DNA |
| 16 | FANPEZAQ\_CDS\_0016 Locus: [12659:13091](+) Seq: MSKYMFSPSTNAFYPVAL... Seq Len: 143 aa MW: 16.2 kDa | 6.4e-39 | 0 | 1.3e-50 | 94.3 | 3.8e-18 | tail, assembly, fiber, phage, chaperone, caudovirales, putative, fibre, lambda, gpk, prophage, virus, gp38, domain\_containing, fragment, hypothetical, duf4376, bacteriophage, cps\_53, kple1 **DA** \| tail, assembly, fiber, Helix, chaperone, Hairpins, Caudovirales, fibre, lambda, gpK, Phage, a, Engineered, yes, Chromobacterium, violaceum, Organism\_taxid, Gene, cv\_0426, Expressed; **IP** \|; **SeS** \| tail, assembly, fiber, Phage, chaperone, Caudovirales, Fragment, Putative, fibre, hypothetical, prophage, Virus, lambda, gpK, domain\_containing, gp38, DUF4376, G, Bacteriophage, CPS\_53; **StS** \| tail, assembly, fiber, Phage, chaperone, Caudovirales, fibre, Putative, prophage, lambda, gpK, Virus, gp38, domain\_containing, DUF4376, homolog, CPS\_53, KpLE1, Bacteriophage, TfaE |
| 17 | FANPEZAQ\_CDS\_0017 Locus: [13208:13874](+) Seq: MSQILEFWKGLSGAVLPF... Seq Len: 221 aa MW: 22.2 kDa | 4.4e-34 | 0 | 8.0e-41 | 83.3 | 3.9e-22 | tail, collar, domain\_containing, phage, fiber, microcystin\_dependent, hypothetical, baseplate, t4, wedge, gp6, complex, tube, putative, baseplate\_tail, pre\_attachment, fragment, phage\_related, side, microcystin **DA** \| tail, fiber, collar, p130317, VI\_07372, short, NC\_023006\_p17, NC\_019917\_p61, p284593, VI\_03272, KJ019100\_p265, p362065, VI\_01943, KU160641\_p20, p203462, VI\_04227, KU234532\_p33, p28392, VI\_07752, p276897; **IP** \|; **SeS** \| tail, Phage, collar, domain\_containing, fiber, Baseplate, T4, wedge, gp6, complex, baseplate\_tail, tube, pre\_attachment, Microcystin\_dependent, Fragment, Putative, Microcystin, dependent, STRUCTURAL, hypothetical; **StS** \| Collar, tail, domain\_containing, Phage, fiber, Hypothetical, Microcystin\_dependent, Putative, Side, Phage\_related, prophage, fibre, transporter, Short\_chain, protein\_like, Microcystin, dependent, fatty, acid, probable |
| 18 | FANPEZAQ\_CDS\_0018 Locus: [14011:15439](+) Seq: MAGTDFLHGVEVLEIDSG... Seq Len: 475 aa MW: 49.9 kDa | 2.6e-57 | 0 | 3.3e-79 | 85.4 | 1.0e-70 | tail, phage, sheath, domain\_containing, fragment, subtilisin\_like, fi, c\_terminal, phage\_sheath\_1c, major, phage\_sheath\_1, putative, prophage, monomer, outer, membrane, secretion, hypothetical, contractile, ragb **DA** \| tail, sheath, Phage, EvpB, VC\_A0108, yes, t4, Rossmann, fold, VI\_12002, Type, VI, secretion, system, TssC, VipB, DUF2586, C\_terminal, N\_terminal, gpW; **IP** \|; **SeS** \| tail, sheath, Phage, domain\_containing, Fragment, subtilisin\_like, C\_terminal, Major, FI, Putative, prophage, monomer, hypothetical, Structural, Afp1, Afp2, Afp3, Anti\_feeding, secretion, system; **StS** \| tail, Phage, sheath, domain\_containing, Phage\_sheath\_1C, Phage\_sheath\_1, FI, Major, Putative, subtilisin\_like, outer, membrane, prophage, RagB, SusD, nutrient, uptake, monomer, Phage\_related, contractile |
| 19 | FANPEZAQ\_CDS\_0019 Locus: [15438:15945](+) Seq: MAARDVLKNINLFVDGRG... Seq Len: 168 aa MW: 18.1 kDa | 3.0e-52 | 5 | 3.6e-56 | 84.1 | 2.2e-23 | tail, phage, tube, major, fii, putative, fragment, contractile, prophage, p2, sheath, bacteriophage, hypothetical, core, head, closure, domain\_containing, from, structural, tp901\_1 **DA** \| head, closure, connector, Phage, tail, tube, FII, p363376, VI\_00073, p268213, VI\_02784, p388731, VI\_08178; **IP** \| tail, Head, closure, Putative, phage, Major, tube; **SeS** \| tail, Phage, tube, major, FII, Fragment, sheath, Putative, contractile, P2, prophage, from, domain\_containing, pyocin, bacteriocin, STRUCTURAL, tail\_like, Hypothetical, Bacteriophage, GpFI; **StS** \| tail, Phage, tube, major, FII, Putative, prophage, contractile, P2, Bacteriophage, core, HYPOTHETICAL, TP901\_1, Probable, sheath, Protoporphyrinogen, oxidase, domain\_containing, Afp1, Baseplate |
| 20 | FANPEZAQ\_CDS\_0020 Locus: [16005:16290](+) Seq: MAKLPDYLKFNHETGHCD... Seq Len: 94 aa MW: 10.6 kDa | 3.3e-23 | 0 | 1.9e-26 | 94.5 | 9.5e-13 | tail, phage, assembly, or, e, chaperone, 14\_like, gp41, flumu, mu\_like, prophage, putative, fragment, bacteriophage, hypothetical, related, phage\_related, domain\_containing, tube, small **DA** \| tail, assembly, chaperone, or, Phage, proteins, E, NC\_003278\_p26, p131196, VI\_08606, p249034, VI\_04203, NC\_024369\_p21, KY421186\_p67, p161962, VI\_09828, p416168, VI\_03036, p338877, VI\_04185; **IP** \|; **SeS** \| tail, Phage, assembly, or, E, chaperone, 14\_like, Mu\_like, FluMu, gp41, prophage, Fragment, Putative, hypothetical, related, Phage\_related, domain\_containing, small, Phage\_like, bacteriophage; **StS** \| tail, assembly, Phage, or, E, chaperone, 14\_like, gp41, Putative, prophage, Bacteriophage, FluMu, Mu\_like, whole, genome, shotgun, sequence, related, ArsR, transcriptional |
| 21 | FANPEZAQ\_CDS\_0021 Locus: [16316:16421](+) Seq: MALASHTGWQLSEIQRLR... Seq Len: 34 aa MW: 4.1 kDa | 1.7e-03 | 0 | 4.6e-16 | 76.1 | -- | phage, gpe, tail, p2, tape, measure, chaperone, hypothetical **DA** \|; **IP** \|; **SeS** \| phage, GpE, tail, P2, Tape, measure, chaperone, hypothetical; **StS** \| |
| 22 | FANPEZAQ\_CDS\_0022 Locus: [16421:18779](+) Seq: MANKKLNATITIGGAVSS... Seq Len: 785 aa MW: 82.7 kDa | 4.4e-44 | 0 | 9.9e-84 | 66.6 | 3.0e-34 | tail, tape, measure, phage, tp901, domain\_containing, core, region, phagemin\_tail, length, phage\_related, minor, putative, fragment, tape\_measure, prophage, bacteriophage, chromosome, smc, determinator **DA** \| tail, length, tape, measure, p335142, VI\_05069, minor, KY203335\_p44, Phage\_related, p348226, VI\_02582, p290120, VI\_00584, p73956, VI\_01292, p160953, VI\_04898, p362597, VI\_05060, p177841; **IP** \|; **SeS** \| tail, Phage, tape, measure, TP901, domain\_containing, core, region, Phage\_related, Fragment, minor, length, tape\_measure, Putative, Membrane, bacteriophage, Chromosome, Transglycosylase, Smc, transmembrane; **StS** \| tail, Phage, tape, measure, TP901, domain\_containing, PhageMin\_Tail, core, region, Phage\_related, minor, Putative, length, tape\_measure, prophage, determinator, Bacteriophage, Chromosome, Lysostaphin, fiber |
| 23 | FANPEZAQ\_CDS\_0023 Locus: [18775:19186](+) Seq: MTSFTILTGQTNIMMMLG... Seq Len: 136 aa MW: 15.6 kDa | 9.5e-35 | 0 | 7.9e-42 | 82.2 | 1.3e-17 | phage, tail, domain\_containing, gpu, lysm, p2, u, tube, contractile, system, injection, n\_terminal, oxidoreductase, peptidoglycan\_binding, putative, fragment, bacteriophage, hypothetical, phage\_related, dna **DA** \| tail, endolysin, lysis, Phage, P2, GpU, Contractile, injection, system, tube, DUF6046, DNA, circularisation, N\_terminus, p258311, VI\_01887, p182850, VI\_05566, p422642, VI\_05571; **IP** \|; **SeS** \| Phage, tail, domain\_containing, Contractile, system, injection, N\_terminal, tube, GpU, LysM, Fragment, P2, U, Peptidoglycan\_binding, hypothetical, Oxidoreductase, Putative, tape, measure, Bacteriophage; **StS** \| Phage, tail, GpU, LysM, domain\_containing, P2, U, Oxidoreductase, Putative, peptidoglycan\_binding, bacteriophage, Phage\_related, prophage, DNA, fiber, DNA\_circ\_N, circulation, major, methyl\_accepting, chemotaxis |
| 24 | FANPEZAQ\_CDS\_0024 Locus: [19185:19395](+) Seq: MQTYITKDGDTADYIAWK... Seq Len: 69 aa MW: 7.4 kDa | 1.4e-26 | 0 | 7.2e-24 | 73.5 | 6.9e-09 | lysm, domain\_containing, tail, phage, x, peptidoglycan\_binding, prophage, p2\_like, fragment, bon, superfamily, putative, spore, coat, assembly, morphogenetic, safa, associated, with, spovid **DA** \| tail, baseplate, wedge, endolysin, lysis, hub, p199511, VI\_11453, MF360958\_p187, AY855346\_p20, MG428991\_p108, NC\_029013\_p171, p23038, VI\_12315, NC\_020871\_p93, NC\_005294\_p57, NC\_027132\_p2, p55048, VI\_12418; **IP** \|; **SeS** \| LysM, domain\_containing, peptidoglycan\_binding, tail, Phage, Fragment, BON, X, superfamily, Spore, coat, assembly, Morphogenetic, SafA, p2\_like, prophage, Putative, associated, with, SpoVID; **StS** \| tail, Phage, X, prophage, P2\_like, domain\_containing, LysM, Putative, peptidoglycan\_binding, component, inner, membrane, completion, Phage\_related, Glue, PA0627, Fels\_2, X\_like, Gp8, Conserved |
| 25 | FANPEZAQ\_CDS\_0025 Locus: [19556:20492](+) Seq: MPETGAELELFLGYDNWT... Seq Len: 311 aa MW: 34.1 kDa | 4.8e-40 | 0 | 4.5e-41 | 82.3 | 3.9e-43 | d, phage, late, control, type, secretion, vi, system, vgr, vgrg, rhs, domain\_containing, tip, tail, element, fragment, gene, ob\_fold, gp5, iv **DA** \| tail, Beta, Phage, Baseplate, fold, roll, Gene, escherichia, coli, Engineered, yes, Organism\_taxid, Expressed, in, ARCH, protein\_like, a, \_, b, kda; **IP** \|; **SeS** \| Type, secretion, VI, system, VgrG, Vgr, Fragment, Rhs, Phage, tip, D, OB\_fold, domain\_containing, Gp5, late, control, element, IV, tail, assembly; **StS** \| D, Phage, late, control, Type, secretion, Rhs, domain\_containing, tail, gene, Phage\_base\_V, element, VI, Vgr, system, VgrG, GPD, tip, IV, Putative |
| 26 | FANPEZAQ\_CDS\_0026 Locus: [20460:20568](+) Seq: MDLGTSLKKNSNKHCCNH... Seq Len: 35 aa MW: 3.8 kDa | 3.3e+00 | 0 | 7.8e-02 | 54.2 | -- | **DA** \|; **IP** \|; **SeS** \|; **StS** \| |
| 27 | FANPEZAQ\_CDS\_0027 Locus: [20586:21288](+) Seq: MNTSNKTLLQILVEELPK... Seq Len: 233 aa MW: 26.4 kDa | 2.9e-80 | 0 | 1.6e-69 | 66.8 | 7.8e-07 | hypothetical, domain\_containing, duf551, duf6378, morphogenetic, fragment, restriction, alleviation, lar, ky705409\_p32, nc\_031918\_p6, mg676466\_p62, nc\_004333\_p63, mf153391\_p29, he956707\_p19, sav\_like, kow, hnh, endonuclease, duf3310 **DA** \| KY705409\_p32, NC\_031918\_p6, MG676466\_p62, NC\_004333\_p63, MF153391\_p29, HE956707\_p19; **IP** \|; **SeS** \| hypothetical, domain\_containing, DUF551, DUF6378, Morphogenetic, Fragment, Restriction, alleviation, Lar, SaV\_like, KOW, HNH, endonuclease, DUF3310, Cupin, DUF3085, dATP, dGTP, diphosphohydrolase, N\_terminal; **StS** \| |
| 28 | FANPEZAQ\_CDS\_0028 Locus: [21368:22244](+) Seq: MATIGQLYKAGATEAKPR... Seq Len: 291 aa MW: 31.7 kDa | 8.5e-40 | 0 | 4.4e-41 | 87.5 | 2.4e-28 | parb, partition, spo0j, domain\_containing, repb, partitioning, chromosome, sulfiredoxin, chromosome\_partitioning, parb\_like, plasmid, probable, fragment, putative, and, dna, nuclease, dna\_binding, stage, sporulation **DA** \| and, ParB, DNA, ParB\_like, partition, RNA, nucleotide, metabolism, N\_terminal, plasmid, the, partitioning, a, in, system, pyrococcus, furiosus, hypothetical, nuclease, Conserved; **IP** \|; **SeS** \| ParB, domain\_containing, Sulfiredoxin, partition, partitioning, Chromosome, Spo0J, RepB, Fragment, chromosome\_partitioning, ParB\_like, Plasmid, Probable, Putative, Nucleoid, occlusion, HTH, Stage, sporulation, J; **StS** \| ParB, partition, Spo0J, RepB, domain\_containing, partitioning, Chromosome, chromosome\_partitioning, ParB\_like, Probable, Putative, Plasmid, Stage, sporulation, J, nuclease, DNA\_binding, segregation, regulator, transcriptional |
| 29 | FANPEZAQ\_CDS\_0029 Locus: [22240:22561](+) Seq: MKGELKAGGLALVYGLRV... Seq Len: 106 aa MW: 11.6 kDa | 4.2e-29 | 0 | 1.1e-35 | 74.4 | 4.6e-04 | hypothetical, domain\_containing, phage, periplasmic, kow, eef2k, atpase, duf2158, fragment, p143733, vi\_10070, nc\_018850\_p11, mh020244\_p60, carboxypeptidase, regulatory\_like, nitrogen, fixation, nifz, duf2171, snoal\_like **DA** \| p143733, VI\_10070, NC\_018850\_p11, MH020244\_p60; **IP** \|; **SeS** \| hypothetical, domain\_containing, Phage, Periplasmic, KOW, ATPase, DUF2158, Fragment, Carboxypeptidase, regulatory\_like, Nitrogen, fixation, NifZ, DUF2171, SnoaL\_like; **StS** \| Hypothetical, Eef2k, domain\_containing, PRPF4B, Peptidylprolyl, isomerase, DUF5641, Stress\_induced\_phosphoprotein, J |
| 30 | FANPEZAQ\_CDS\_0030 Locus: [22570:22783](+) Seq: MKLTKATDNGFNLWVVEH... Seq Len: 70 aa MW: 7.7 kDa | 9.4e-01 | 0 | 7.2e-38 | 76.8 | 6.0e-02 | **DA** \|; **IP** \|; **SeS** \|; **StS** \| |
| 31 | FANPEZAQ\_CDS\_0031 Locus: [22905:23097](+) Seq: MALFAVKVLHYHTLNHTL... Seq Len: 63 aa MW: 7.1 kDa | 9.8e-01 | 0 | 7.2e-03 | 60.1 | -- | **DA** \|; **IP** \|; **SeS** \|; **StS** \| |
| 32 | FANPEZAQ\_CDS\_0032 Locus: [23199:24411](+) Seq: MLTERQIQAAMRAVTSET... Seq Len: 403 aa MW: 45.0 kDa | 2.4e-35 | 0 | 2.1e-70 | 88.3 | 5.2e-36 | integrase, recombinase, domain\_containing, tyr, site\_specific, phage, duf4102, prophage, tyrosine\_type, tyrosine, xerd, putative, dna, cp4\_57, xerc, catalytic, dna\_binding, complex, a, tp **DA** \| integrase, catalytic, a, tp, ap, and, recombinase, yes, C\_terminal, Engineered, in, site\_specific, Dna, integration, excision, tyrosine, Mainly, Alpha, ARCH, Orthogonal; **IP** \|; **SeS** \| Integrase, recombinase, domain\_containing, Tyr, Site\_specific, Phage, DNA, XerD, complex, Prophage, Tyrosine, Putative, Fragment, RECOMBINATION, DNA\_binding, PROTEIN\_DNA, BINDING, Structural, CP4\_57, DUF4102; **StS** \| Integrase, recombinase, domain\_containing, Tyr, Site\_specific, DUF4102, Tyrosine\_type, Phage, Prophage, Tyrosine, Putative, XerD, CP4\_57, XerC, arm\_type, DNA\_binding, CPS\_53, Int, IntA, Core\_binding |
| 33 | FANPEZAQ\_CDS\_0033 Locus: [24423:24684](-) Seq: MTKAATNDQRKTPPKRGF... Seq Len: 86 aa MW: 9.7 kDa | 1.3e-01 | 0 | 1.8e-01 | 59.5 | 3.4e-02 | **DA** \|; **IP** \|; **SeS** \|; **StS** \| |
| 34 | FANPEZAQ\_CDS\_0034 Locus: [24661:25345](-) Seq: MDREGWLNALAAKFAPRF... Seq Len: 227 aa MW: 24.9 kDa | 2.6e-52 | 1 | 7.8e-50 | 84.7 | 1.2e-23 | sprt\_like, domain\_containing, fragment, sprt, acidic, repeat\_containing, metalloprotease, transcription, elongation, hypothetical, zinc\_dependent, spartan, putative, protein\_like, with, at, the, n, terminus, zinc **DA** \| MG670586\_p45, JQ512844\_p79, NC\_019538\_p220, NC\_029000\_p177, NC\_019401\_p436, NC\_029009\_p209, NC\_021327\_p65, NC\_028683\_p287, AP018399\_p45, NC\_015251\_p422, p367521, VI\_04587, NC\_017972\_p199; **IP** \| Hypothetical; **SeS** \| SprT\_like, domain\_containing, Fragment, Acidic, SprT, repeat\_containing, metalloprotease, hypothetical, Spartan, Putative, zinc\_dependent, protein\_like, with, at, the, N, terminus, Zinc, isoform, protease; **StS** \| SprT\_like, domain\_containing, SprT, Transcription, elongation, zinc\_dependent, metalloprotease, PH, M48, peptidase |
| 35 | FANPEZAQ\_CDS\_0035 Locus: [25443:25932](-) Seq: MAGSLNKVEIIGNLGNDP... Seq Len: 162 aa MW: 17.6 kDa | 6.1e-35 | 0 | 5.3e-45 | 81.6 | 8.9e-21 | single\_stranded, dna\_binding, binding, dna, single\_strand, replication, beta, a, primosomal, n, complex, mitochondrial, fold, yes, ob, fragment, single, engineered, rna, nucleic **DA** \| Beta, yes, Engineered, OB, fold, a, Organism\_taxid, Mainly, ARCH, Barrel, Dihydrolipoamide, Acetyltransferase, E2P, Nucleic, acid\_binding, proteins, escherichia, coli, Expressed, in; **IP** \|; **SeS** \| Single\_stranded, DNA\_binding, DNA, binding, Single\_strand, Fragment, Single, replication, complex, Primosomal, fold, n, SSB, STRANDED, OB, OB\_fold, VHH, antibody, RNA, MSE; **StS** \| DNA\_binding, Single\_stranded, Single\_strand, binding, replication, mitochondrial, Primosomal, n, A, complex, RNA, editing, RIM1, DNA, MP18, Hypothetical, factor, chloroplastic, RNA\_editing, putative |
| 36 | FANPEZAQ\_CDS\_0036 Locus: [25940:26252](-) Seq: MTTNSRTIAALFAVSRMV... Seq Len: 103 aa MW: 11.2 kDa | 1.4e-21 | 0 | 7.3e-18 | 76.1 | 2.3e-03 | dksa, c4\_type, domain\_containing, trar, transcriptional, regulator, zinc, finger, dnak, rna, conjugal, transfer, polymerase\_binding, transcription, factor, fragment, suppressor, molecular, chaperone, dna\_binding **DA** \| DksA\_like, zinc\_finger, other, MF490239\_p53; **IP** \|; **SeS** \| DksA, C4\_type, domain\_containing, TraR, transcriptional, regulator, zinc, finger, DnaK, RNA, Conjugal, transfer, polymerase\_binding, transcription, factor, Fragment, suppressor, Molecular, chaperone, DNA\_binding; **StS** \| |
| 37 | FANPEZAQ\_CDS\_0037 Locus: [26258:26555](-) Seq: MLVVMNVWLMYWNAPLTR... Seq Len: 98 aa MW: 11.2 kDa | 1.6e+00 | 0 | 2.2e-01 | 39.0 | 1.0e+00 | **DA** \|; **IP** \|; **SeS** \|; **StS** \| |
| 38 | FANPEZAQ\_CDS\_0038 Locus: [26548:26770](-) Seq: MAQPAQKQSTYLTPQELV... Seq Len: 73 aa MW: 8.1 kDa | 7.0e-15 | 0 | 4.9e-24 | 74.9 | 5.1e-07 | domain\_containing, helix\_turn\_helix, dna\_binding, regulator, hth\_17, transcriptional, excisionase, dna, binding, alpa, merr, phage, a, putative, tp, and, regulatory, ap, yes, cp **DA** \| A, tp, Regulator, and, ap, DNA, yes, cp, gp, Engineered, Multidrug\_efflux, Transporter, escherichia, excisionase, coli, Organism\_taxid, ARCH, Expressed, in, Expression\_system\_taxid; **IP** \|; **SeS** \| Helix\_turn\_helix, domain\_containing, regulator, transcriptional, DNA, excisionase, DNA\_binding, AlpA, binding, MerR, phage, regulatory, Complex, Transcription, Putative, activator, HTH\_type, HTH, CadR, MSE; **StS** \| domain\_containing, DNA\_binding, Helix\_turn\_helix, HTH\_17, Excisionase, transcriptional, regulator, DNA, binding, AlpA, Putative, MerR, Phage, regulatory, Xis, HTH, site\_specific, integrase\_resolvase, merR\_type, Terminase |
| 39 | FANPEZAQ\_CDS\_0039 Locus: [26783:27773](-) Seq: MAVINIRKAEREGARLVI... Seq Len: 329 aa MW: 36.6 kDa | 1.2e-29 | 0 | 2.4e-57 | 87.7 | 1.1e-41 | kaic, domain\_containing, aaa, atpase, dna, circadian, clock, reca, kinase, repair, recombination, fragment, and, atp, helicase, rad51, profile, rada, synthase, in **DA** \| and, DNA, in, nucleotide, the, a, RNA, metabolism, KaiC, is, repair, Alpha, Rad51, recombination, escherichia, coli, helicase, Organism\_taxid, Beta, ARCH; **IP** \|; **SeS** \| ATPase, domain\_containing, AAA, RecA, DNA, recombination, Fragment, KaiC, repair, profile, ATP, and, RadA, Circadian, kinase, homologous, clock, Complex, ATP\_binding, ADP; **StS** \| KaiC, domain\_containing, Circadian, clock, AAA, kinase, ATPase, DNA, repair, beta, ATP, synthase, helicase, RAD51, RecA, homolog, Replicative, Putative, mitochondrial, ATP\_binding |
| 40 | FANPEZAQ\_CDS\_0040 Locus: [27759:27939](-) Seq: MDALIYLVLVLFFYGCWF... Seq Len: 59 aa MW: 6.9 kDa | 6.2e-02 | 0 | 3.8e-02 | 70.2 | 2.1e+00 | **DA** \|; **IP** \|; **SeS** \|; **StS** \| |
| 41 | FANPEZAQ\_CDS\_0041 Locus: [27938:29057](-) Seq: MKPGVYEGIPNAEYHGGP... Seq Len: 372 aa MW: 40.8 kDa | 1.1e-63 | 0 | 3.3e-70 | 93.6 | 8.2e-48 | domain\_containing, duf3799, exodeoxyribonuclease, putative, pddexk\_like, e, d, pd\_, xk, dna, helicase, nuclease\_like, viii, exonuclease, pddexk\_1, nuclease, fragment, breaking\_rejoining, 3'\_5', endonuclease\_like **DA** \| Exonuclease, A, yes, Lambda, Engineered, escherichia, coli, Organism\_taxid, Gene, Expressed, in, Expression\_system\_taxid, Alpha, Beta, ARCH, Alpha\_Beta, Complex, nuclease, Strain, Atp\_dependent; **IP** \|; **SeS** \| domain\_containing, exodeoxyribonuclease, Putative, PDDEXK\_like, E, PD\_, D, XK, nuclease\_like, DNA, helicase, Fragment, Exonuclease, 3'\_5', endonuclease\_like, VIII, nuclease, ATP\_dependent, \_, hypothetical; **StS** \| domain\_containing, DUF3799, Exodeoxyribonuclease, DNA, helicase, PD\_, D, E, XK, VIII, PDDEXK\_1, nuclease, Exonuclease, breaking\_rejoining, nuclease\_like, Putative, superfamily, PDDEXK\_like, RecE, Cas4 |
| 42 | FANPEZAQ\_CDS\_0042 Locus: [29139:29682](-) Seq: MPNRIRERRQAAGMTLQD... Seq Len: 180 aa MW: 19.9 kDa | 4.3e-23 | 0 | 3.8e-37 | 86.3 | 3.1e-13 | transcriptional, domain\_containing, regulator, repressor, hth, cro, c1\_type, peptidase, helix\_turn\_helix, s24, phage, putative, c, peptidase\_s24, and, lexa, peptisase, transcription, s24\_like, dna **DA** \| transcription, regulation, transcriptional, DNA, repressor, regulator, A, and, Fragment, Umud, RNA, nucleotide, metabolism, binding, HTH, peptidase, yes, escherichia, coli, in; **IP** \|; **SeS** \| domain\_containing, repressor, transcriptional, regulator, HTH, cro, C1\_type, Peptidase, Helix\_turn\_helix, S24, LexA, Phage, Putative, DNA, TRANSCRIPTION, S26A, S26B, S26C, Structural, C; **StS** \| transcriptional, regulator, domain\_containing, HTH, repressor, cro, C1\_type, Helix\_turn\_helix, Peptidase, Phage, S24, Peptidase\_S24, Putative, C, and, peptisase, S24\_like, LexA, HTH\_type, with |
| 43 | FANPEZAQ\_CDS\_0043 Locus: [29675:29807](+) Seq: MASLQILNNHAEWTVTDT... Seq Len: 43 aa MW: 5.0 kDa | 8.9e-01 | 0 | 4.5e-01 | 42.0 | 2.5e-01 | **DA** \|; **IP** \|; **SeS** \|; **StS** \| |
| 44 | FANPEZAQ\_CDS\_0044 Locus: [29851:31267](+) Seq: MGVVLRDYQQDIIDKGRQ... Seq Len: 471 aa MW: 51.9 kDa | 6.2e-30 | 0 | 2.0e-108 | -- | -- | helicase, rna, dna, a, atp\_dependent, dead, box, and, in, the, hydrolase, i, deah, complex, c, yes, nucleotide, fragment, dead\_box, e **DA** \| helicase, a, and, in, the, DNA, RNA, i, yes, ATP\_dependent, nucleotide, Engineered, DEAD, box, containing, Alpha, Beta, ARCH, 3\_Layer, aba; **IP** \|; **SeS** \| helicase, RNA, DEAD, DNA, box, ATP\_dependent, DEAH, complex, C, HYDROLASE, E, ADP, domain\_containing, ATPase, Fragment, ATP\_binding, DEAD\_box, factor, restriction, Type; **StS** \| |
| 45 | FANPEZAQ\_CDS\_0045 Locus: [31356:31515](+) Seq: MNTTHDMGNNETIKTGVF... Seq Len: 52 aa MW: 5.8 kDa | 5.3e-39 | 0 | 1.2e-40 | 94.7 | 2.1e-06 | duf1391, domain\_containing, prophage, putative, phage, fragment, p287170, vi\_07176, p274931, vi\_10756, pf07151, ydaf, ydfa **DA** \| DUF1391, p287170, VI\_07176, p274931, VI\_10756; **IP** \|; **SeS** \| DUF1391, domain\_containing, prophage, Putative, Fragment, Phage, PF07151; **StS** \| DUF1391, domain\_containing, prophage, Phage, Putative, YdaF, YdfA |
| 46 | FANPEZAQ\_CDS\_0046 Locus: [31564:32212](+) Seq: MARLTAAIRDGIIANAIK... Seq Len: 215 aa MW: 24.2 kDa | 5.6e-54 | 0 | 1.7e-51 | 85.0 | 1.2e-09 | domain\_containing, nucleotide, modification, nmad5, associated, putative, fragment, hypothetical, nc\_028980\_p51, gp23, dna\_directed, dna, polymerase, a, palm, contig\_80, whole, genome, shotgun, sequence **DA** \| Nucleotide, modification, associated, NC\_028980\_p51; **IP** \|; **SeS** \| Nucleotide, modification, associated, domain\_containing, Nmad5, putative, Fragment, hypothetical, Gp23, DNA\_directed, DNA, polymerase, A, palm; **StS** \| Nmad5, domain\_containing, Contig\_80, whole, genome, shotgun, sequence |
| 47 | FANPEZAQ\_CDS\_0047 Locus: [32276:32669](+) Seq: MKKLIIAAFILAATSAAA... Seq Len: 130 aa MW: 14.4 kDa | 5.5e-87 | 0 | 4.9e-82 | 64.4 | 1.0e-02 | nc\_025430\_p8 **DA** \| NC\_025430\_p8; **IP** \|; **SeS** \|; **StS** \| |
| 48 | FANPEZAQ\_CDS\_0048 Locus: [32705:33176](+) Seq: MNYSGCWWFWLLALSWWG... Seq Len: 156 aa MW: 17.2 kDa | 1.9e-31 | 0 | 2.5e-41 | 82.3 | 7.4e-17 | vrr\_nuc, domain\_containing, nuclease, fragment, endonuclease, a, containing, hypothetical, putative, trna, dna, hydrolase, engineered, yes, organism\_taxid, expressed, in, escherichia, coli, expression\_system\_taxid **DA** \| a, Endonuclease, Trna, nuclease, Engineered, yes, Organism\_taxid, Expressed, in, escherichia, coli, Expression\_system\_taxid, Alpha, Beta, ARCH, 3\_Layer, aba, Sandwich, and, b; **IP** \|; **SeS** \| VRR\_NUC, domain\_containing, Nuclease, Fragment, containing, hypothetical, Endonuclease, Putative, HYDROLASE, DNA, Fanconi\_associated, Recombinase, RecB, complex, E, C, \_, norphogenetic, Phage, restriction; **StS** \| VRR\_NUC, domain\_containing, Penicillin\_binding, protein\_related, factor, A, Nuclease |
| 49 | FANPEZAQ\_CDS\_0049 Locus: [33251:33416](+) Seq: MMTIESIRRLFAEALVTR... Seq Len: 54 aa MW: 6.3 kDa | 9.4e-43 | 0 | 7.6e-35 | 81.8 | 1.9e+00 | nc\_025430\_p10 **DA** \| NC\_025430\_p10; **IP** \|; **SeS** \|; **StS** \| |
| 50 | FANPEZAQ\_CDS\_0050 Locus: [33412:33649](+) Seq: MNIFKVAVQRLVDAWKLS... Seq Len: 78 aa MW: 9.0 kDa | 1.0e-51 | 0 | 5.9e-48 | 61.1 | 9.6e-01 | nc\_025430\_p11 **DA** \| NC\_025430\_p11; **IP** \|; **SeS** \|; **StS** \| |
| 51 | FANPEZAQ\_CDS\_0051 Locus: [33645:33870](+) Seq: MNPVDFIKKQVMAELVKQ... Seq Len: 74 aa MW: 8.1 kDa | 6.7e-34 | 0 | 1.2e-33 | 89.1 | 3.8e-02 | hypothetical, phage, transcriptional, regulator, p119269, vi\_03293, cell, envelope, biogenesis, ompa, membrane, lipoprotein, duf4222, domain\_containing, cro, ci, coil, containing, lysis, system **DA** \| p119269, VI\_03293; **IP** \|; **SeS** \| hypothetical, Phage, Transcriptional, regulator, Cell, envelope, biogenesis, OmpA, Membrane, lipoprotein, DUF4222, domain\_containing, Cro, CI, Coil, containing, lysis, system, i\_spanin, Rz; **StS** \| |
| 52 | FANPEZAQ\_CDS\_0052 Locus: [33905:34139](+) Seq: MRPVLWAHCYYEAITMDE... Seq Len: 77 aa MW: 8.9 kDa | 7.7e-20 | 0 | 1.0e-18 | 81.9 | 6.9e-04 | transcriptional, regulator, hth, domain\_containing, cro, c1\_type, helix\_turn\_helix, xre, dna\_binding, repressor, hth\_type, transcription, dna, regulation, and, xre\_family, immr, dica, binding, rna **DA** \| transcription, regulation, DNA, transcriptional, and, regulator, binding, RNA, nucleotide, metabolism, repressor, HTH, integration, excision, terminase, small, head, packaging, CII\_like, excisionase; **IP** \|; **SeS** \| transcriptional, regulator, domain\_containing, HTH, cro, C1\_type, Helix\_turn\_helix, XRE, DNA\_binding, HTH\_type, repressor, XRE\_family, immR, DicA, Putative, contains, phage, Antitoxin, CI, LexA; **StS** \| |
| 53 | FANPEZAQ\_CDS\_0053 Locus: [34135:34366](+) Seq: MNACTWQPTTVTLQDGRE... Seq Len: 76 aa MW: 8.8 kDa | 9.5e-42 | 2 | 4.0e-30 | 93.0 | 1.6e-06 | hypothetical, ep3\_0015, domain\_containing, ky705409\_p11, p222118, vi\_08569, duf1289, kfra, n\_terminal, dna\_binding **DA** \| KY705409\_p11, p222118, VI\_08569; **IP** \| Hypothetical, ep3\_0015; **SeS** \| hypothetical, domain\_containing, DUF1289, KfrA, N\_terminal, DNA\_binding; **StS** \| |
| 54 | FANPEZAQ\_CDS\_0054 Locus: [34378:36688](+) Seq: MSAQQNYKDLTEQEIAEA... Seq Len: 769 aa MW: 86.4 kDa | 2.4e-76 | 0 | 2.0e-108 | 86.8 | 9.0e-74 | dna, domain\_containing, primase, virulence\_associated, e, replication, factor, licensing, toprim, helicase, putative, e\_like, atpase, p\_loop, and, virulence, mcm2, trac, mcm3, prict\_2 **DA** \| DNA, RNA, and, nucleotide, metabolism, primase, helicase, polymerase, A, ATPase, clamp, loader, E, in, P\_loop, yes, Topoisomerase, Vi, other, Virulence\_associated; **IP** \|; **SeS** \| primase, domain\_containing, DNA, Virulence\_associated, E, E\_like, Toprim, Fragment, helicase, ATPase, Putative, P\_loop, VapE, C, TraC, polymerase, N\_terminal, Phage, \_, plasmid; **StS** \| DNA, domain\_containing, E, Virulence\_associated, primase, replication, factor, licensing, helicase, Toprim, Putative, ATPase, P\_loop, MCM2, Virulence, MCM3, PriCT\_2, DUF3874, MCM6, TraC |
| 55 | FANPEZAQ\_CDS\_0055 Locus: [36820:36940](+) Seq: MIVVSGFPTVRPERTHYR... Seq Len: 39 aa MW: 4.7 kDa | 4.7e-01 | 0 | 4.0e+00 | 61.7 | -- | **DA** \|; **IP** \|; **SeS** \|; **StS** \| |
| 56 | FANPEZAQ\_CDS\_0056 Locus: [36958:37057](+) Seq: MGPLVNTMNNIIFIYAIG... Seq Len: 32 aa MW: 3.4 kDa | 2.0e-01 | 0 | 1.1e-01 | 91.7 | -- | **DA** \|; **IP** \|; **SeS** \|; **StS** \| |
| 57 | FANPEZAQ\_CDS\_0057 Locus: [37053:38106](+) Seq: MKQVTIDTVRERIAELEM... Seq Len: 350 aa MW: 39.2 kDa | 4.5e-24 | 0 | 4.4e-25 | 54.5 | 3.0e+00 | domain\_containing, duf551, hypothetical, phage, eaa, eaa1, putative, duf550, duf3850, ead, lar, restriction, alleviation, ea22\_like, fragment, p181809, vi\_03505, p158937, vi\_06901, conserved **DA** \| p181809, VI\_03505, p158937, VI\_06901; **IP** \|; **SeS** \| domain\_containing, DUF551, hypothetical, Phage, Eaa, Eaa1, Putative, DUF550, DUF3850, Ead, Lar, restriction, alleviation, Ea22\_like, Fragment, Conserved, 3'\_5', exoribonuclease, Molecular, chaperone; **StS** \| |
| 58 | FANPEZAQ\_CDS\_0058 Locus: [38125:38293](+) Seq: MARFCYAGETEAEAKTST... Seq Len: 55 aa MW: 5.7 kDa | 2.7e-01 | 0 | 7.1e-01 | 51.7 | -- | **DA** \|; **IP** \|; **SeS** \|; **StS** \| |
| 59 | FANPEZAQ\_CDS\_0059 Locus: [38292:38460](+) Seq: MARSNTPWEHSKQCHAHA... Seq Len: 55 aa MW: 6.1 kDa | 1.4e+00 | 0 | 1.2e-02 | 50.0 | -- | **DA** \|; **IP** \|; **SeS** \|; **StS** \| |
| 60 | FANPEZAQ\_CDS\_0060 Locus: [38713:38818](+) Seq: MTGDKRVTNSLPPCHPFL... Seq Len: 34 aa MW: 3.9 kDa | 2.6e-01 | 0 | 6.9e-02 | 56.4 | 7.6e+00 | **DA** \|; **IP** \|; **SeS** \|; **StS** \| |
| 61 | FANPEZAQ\_CDS\_0061 Locus: [38832:39267](+) Seq: MELPRTVQEIADVIGRER... Seq Len: 144 aa MW: 16.6 kDa | 9.6e-33 | 0 | 1.5e-31 | 84.3 | 8.0e-14 | mor, domain\_containing, transcription, activator, regulator, transcriptional, helix\_turn\_helix, response, dna, hypothetical, binding, and, hth, regulatory, dna\_binding, operon, homeodomain\_like, rna, regulation, middle **DA** \| and, DNA, RNA, nucleotide, metabolism, transcription, transcriptional, regulation, binding, HTH, terminase, head, packaging, small, regulator, activator, endonuclease, late, HNH, repressor; **IP** \|; **SeS** \| transcription, Mor, activator, domain\_containing, regulator, Helix\_turn\_helix, Transcriptional, Response, hypothetical, DNA, regulatory, DNA\_binding, BINDING, HTH, operon, system, Two\_component, Middle, Homeodomain\_like, Fragment; **StS** \| Mor, domain\_containing, transcription, activator, regulator, Transcriptional, Helix\_turn\_helix, Homeodomain\_like, DNA, transposition, HTH, activator\_like, DNA\_binding, HTH\_7, Winged, Middle, operon, Putative, luxR\_type, Sigma70\_r4\_2 |
| 62 | FANPEZAQ\_CDS\_0062 Locus: [39271:39592](+) Seq: MEFFQKFLDNIGWGIAGI... Seq Len: 106 aa MW: 11.5 kDa | 6.3e-25 | 0 | 1.9e-26 | 83.5 | 1.8e-08 | holin, hypothetical, putative, phage, membrane, fragment, transporter, mfs, lysis, peptidase, m48, ste24p, nc\_023006\_p63, p387102, vi\_09157, p363326, vi\_04187, p127786, vi\_03659, antiholin **DA** \| holin, lysis, NC\_023006\_p63, p387102, VI\_09157, p363326, VI\_04187, p127786, VI\_03659; **IP** \|; **SeS** \| Holin, hypothetical, Putative, Phage, membrane, Fragment, transporter, MFS, Peptidase, M48, Ste24p, antiholin, lambda, Holin\_like, II, Conserved, N\_acetylmuramidase, DUF1097, domain\_containing, prophage; **StS** \| MFS, transporter, Contig\_80, whole, genome, shotgun, sequence, Ethanolamine, permease, Holin |
| 63 | FANPEZAQ\_CDS\_0063 Locus: [39591:39870](+) Seq: MLLTVINFISVLVIFCWS... Seq Len: 92 aa MW: 10.4 kDa | 1.2e-35 | 0 | 8.3e-41 | 79.9 | 3.9e-05 | holin, hypothetical, putative, holin\_like, i, membrane, lysis, nc\_019419\_p7, rhs, repeat\_associated, core, domain\_containing, feob\_associated, cys\_rich **DA** \| holin, lysis, NC\_019419\_p7; **IP** \|; **SeS** \| Holin, hypothetical, Putative, holin\_like, I, membrane, RHS, repeat\_associated, core, domain\_containing, FeoB\_associated, Cys\_rich; **StS** \| |
| 64 | FANPEZAQ\_CDS\_0064 Locus: [39878:40358](+) Seq: MKISSNGIAVLKYFEDCH... Seq Len: 159 aa MW: 17.0 kDa | 1.2e-33 | 0 | 1.1e-52 | 96.4 | 1.8e-21 | lysozyme, endolysin, t4, receptor, hydrolase, e, c, glycoside, fragment, chimera, phage, type, muramidase, phage\_related, glucagon\_like, peptide, a, membrane, lysis, b **DA** \| Lysozyme, endolysin, phage, lysis, in, a, yes, the, escherichia, coli, Enterobacteria, Engineered, Organism\_taxid, Gene, Expressed, Expression\_system\_taxid, Mainly, Alpha, ARCH, Orthogonal; **IP** \|; **SeS** \| Lysozyme, Endolysin, E, C, Fragment, hydrolase, receptor, glycoside, Membrane, complex, Baseplate, SO4, T4, B, Calcium, GPCR, RND, Antimicrobial, STRUCTURAL, NAG; **StS** \| Lysozyme, T4, Endolysin, receptor, chimera, hydrolase, Glycoside, type, Glucagon\_like, peptide, phage\_related, GH24, Muramidase, phage, factor, domain\_containing, RrrD, Putative, chemokine, Phosphatidylinositol |
| 65 | FANPEZAQ\_CDS\_0065 Locus: [40354:40876](+) Seq: MMDKYATPVKLIVAIIIA... Seq Len: 173 aa MW: 18.7 kDa | 2.2e-35 | 0 | 3.4e-36 | 92.3 | 8.5e-11 | duf2514, lysis, domain\_containing, fragment, lysozyme, rz, bacteriophage, putative, spanin, endopeptidase, prophage, phage, membrane, rz\_like, system, i\_spanin, assembly, inner, phage\_tail, associated **DA** \| lysis, Rz\_like, spanin, DUF2514, Bacteriophage, Rz, NC\_027388\_p53, KY000219\_p64, p303683, VI\_03517, p139219, VI\_05583, p348110, VI\_03518, NC\_011589\_p12, p236003, VI\_10093, MG897800\_p45; **IP** \|; **SeS** \| DUF2514, lysis, domain\_containing, Fragment, Lysozyme, Rz, Putative, Bacteriophage, Endopeptidase, prophage, Phage, membrane, Spanin, system, i\_spanin, assembly, inner, Phage\_tail, associated, hypothetical; **StS** \| DUF2514, Lysozyme |
| 66 | FANPEZAQ\_CDS\_0066 Locus: [40872:40983](+) Seq: MKIFSRTTQIMLDIDRSR... Seq Len: 36 aa MW: 3.7 kDa | 7.1e-01 | 0 | 5.1e-02 | 69.5 | 8.9e+00 | **DA** \|; **IP** \|; **SeS** \|; **StS** \| |

  


---
